# Phenotypic reconstruction of the last universal common ancestor reveals a complex cell

**DOI:** 10.1101/2020.08.20.260398

**Authors:** Fouad El Baidouri, Chris Venditti, Sei Suzuki, Andrew Meade, Stuart Humphries

## Abstract

A fundamental concept in evolutionary theory is the last universal common ancestor (LUCA) from which all living organisms originated. While hypotheses regarding the phenotype of LUCA do exist, most are at best based on gene presence or absence. However, despite recent attempts to link genes and phenotypic traits in prokaryotes, it is still inherently difficult to predict phenotype based on the presence or absence of genes alone. Here, we apply a novel approach based on phylogenetically informed ancestral character reconstruction with 22 fundamental descriptors of prokaryotic cells to reconstruct LUCA’s phenotype. We infer that the last universal common ancestor of all living organisms was likely an ovoid cell with a large genome, which probably had a cell wall and was actively motile. Potentially obtaining energy from the oxidation of inorganic compounds, tolerant of saltwater and thriving at temperatures above 70°C, LUCA was probably found living freely in anaerobic conditions in water of neutral pH. Our results depict LUCA as likely to be a far more complex cell than has previously been proposed, challenging the evolutionary model of increased complexity through time in prokaryotes. Given current estimates for the emergence of LUCA we suggest that early life very rapidly evolved considerable cellular complexity.

## Introduction

The pervasive perception of LUCA as a ‘primitive’ cell is inherently linked to the slow and uniform gradualist model of evolution. Within this model it is widely assumed that LUCA must have been a very simple cell [1,2] and that life has subsequently increased in complexity through time [3,4]. However, while current thought tends towards a general increase in complexity through time in most Eukaryotes [5–7], there is increasing evidence that bacteria and archaea have undergone considerable genome reduction during their evolution [8,9]. It is now largely accepted that LUCA is the common ancestor of bacteria and archaea [10,11] with eukaryotes originating from within the archaea in the two domains tree of life [12]. This raises the surprising possibility that LUCA may have been a complex cell [13,14].

While the concepts of LUCA and progenote are sometimes used interchangeably in the literature, creating some confusion, here we refer to LUCA as the most recent common ancestral population of cells that diverged into the two primary domains of life, bacteria and archaea. LUCA itself must have evolved from a population of ancestral proto-cellular forms of lower complexity and Woese and Fox [15] introduced an ancestral organism of bacteria and archaea, the “progenote”, as an organism with a lower level of cellular organization than that of a free-living cell [16,17].

For over half a century one of the most challenging aspects of evolutionary biology has been to infer the ancestral habitat and characteristics of LUCA. While much has been learnt about LUCAs habitat from geology and geochemistry [e.g., 18,19], few hypotheses exist about its phenotype and physiology [3,4,20], and most were classically considered best deduced from comparative genomics and gene content. However, despite recent attempts to link genotypes and phenotypes in prokaryotes [21,22], it is still inherently difficult to predict phenotypic traits based on presence or absence of genes alone. Moreover, the methods used to infer ancestral gene content are limited in their ability to accurately identify conserved clusters of orthologous groups (COGs), detect and exclude genes that have undergone Horizontal gene transfer (HGT) from the ancestral pool of genes that trace back to LUCA [13,23,24]. Most methods are therefore conservative and while the latest estimates predict a few hundred genes (∼355) that trace back to LUCA [4] older studies predicted around 1,000 genes [20]. In both cases these estimates are similar to either the average gene content of parasitic or symbiotic bacteria and archaea that lost genes mainly through genetic drift, or free-living prokaryotes that have undergone genome streamlining [9,25].

A LUCA with a relatively complex genome has been explicitly suggested [14,20], but in recent years it has become apparent that many archaeal and bacterial phenotypic traits, as well as their the replicative machineries are similar. The latest evidence suggests that bacterial and archaeal replication and transcription systems have a common origin and that LUCA most likely had complex replication and transcription systems too [26]. Similarly, while archaeal and bacterial membranes are composed of different types of phospholipids (leading to the idea of “non-cellular” LUCA [4,27]) it now seems that LUCA might have possessed a mixed membrane of both bacterial and archaeal phospholipids [28,29]. While the possibility of a wall-less LUCA cannot be excluded similar reasoning can be applied due to the presence of an S-layer (archaeal cell wall) in some bacteria [30]. A particularly contentious debate centred around LUCA’s habitat and physiology is whether it was hyperthermophilic as suggested by Weiss et al. [4,31]. While arguments in favour of a hyperthermophilic common ancestor of archaea exist [32,33] there is no clear consensus for LUCA and is the subject of ongoing debate [4,17,34]. Taken together these studies highlight the conflicting views on the complexity of LUCA and suggest that researchers need to identify new ways to approach the study of cellular origins.

Given these discrepancies and the difficulty in predicting phenotype based on the presence or absence of genes alone, a new approach focused on phylogenetic ancestral state reconstruction of traits could provide new insights and move the discussion onwards. Here we present one new potential approach to understanding the evolution of cells, by using Bayesian Phylogenetic comparative methods [35,36]. After testing for robustness to HGT we infer the phenotypic traits of LUCA using two recently published phylogenetic trees [37,38] and the largest phenotypic dataset of 3,128 bacterial and archaeal species (https://doi.org/10.6084/m9.figshare.12987509.v1) using 18 discrete and 10 continuous traits.

## Results and Discussion

Using phylogenetic comparative methods [39,40] and the largest compilation to-date of bacterial and archaeal phenotypic data (Fig. 1), along with two robust prokaryotic phylogenetic trees [37,38] we are able to infer that the last universal common ancestor of all living organisms was likely to have been a complex cell with at least 22 reconstructed phenotypic traits probably as intricate as those of many modern bacteria and archaea (Fig. 2 and Table 1). Given current estimates for the time of Earth’s formation and the presence of liquid water offering habitable conditions [41] our results suggest that early life must have very quickly (perhaps ∼ 200 Myr [31], but certainly within 500 Myr [18]) evolved considerable cellular complexity. While there are already strong arguments for a trend of increasing biological complexity over time [5,6,42] they point to a general increase in complexity across the whole tree of life and do not address the rapid development of complexity we propose here for the early prokaryotes.

**Table 1.** Bayesian values estimates for continuous traits for LUCA, LBCA and LACA from the large tree. Ancestral state estimates for cell and genome sizes, optimum temperature, NaCl and pH from the best of two competing models - the random walk and the directional models. Phylogenetic signal (λ) is estimated for each trait. Abbreviations: LUCA: last universal common ancestor, LBCA: last bacterial common ancestor, LACA: last archaeal common ancestor. HPD - Highest Posterior Density; 95% PI - 95% Probability Interval.

| | Number of species | $\lambda$ | $\lambda$ HPD range | $\lambda$ 95% PI range | LUCA PP | LUCA 95% PI range | LBCA PP | LBCA 95% PI range | LACA PP | LACA 95% PI range |
| --- | --- | --- | --- | --- | --- | --- | --- | --- | --- | --- |
| **Cell size ($\mu\text{m}$)** |  |  |  |  |  |  |  |  |  |  |
| **Average width** | 1705 | 1.00 | 0.99-1.00 | 0.99-1.00 | 0.53 | 0.43-0.65 | 0.49 | 0.39-0.62 | 0.54 | 0.44-0.67 |
| **Average length** | 1656 | 0.99 | 0.98-0.99 | 0.98-0.99 | 1.72 | 1.24-2.38 | 1.87 | 1.39-2.51 | 1.73 | 1.2-2.49 |
| **Genome** |  |  |  |  |  |  |  |  |  |  |
| **Average genome size (Mb)** | 2978 | 1.00 | 1.00 | 1.00 | 2.48 | 2.13-2.90 | 2.76 | 2.25-3.37 | 2.42 | 2.05-2.85 |
| **Average gene number** | 2965 | 1.00 | 1.00 | 1.00 | 2624 | 2261-3047 | 2717 | 2244-3290 | 2585 | 2225-3003 |
| **Temperature ($^{\circ}\text{C}$)** |  |  |  |  |  |  |  |  |  |  |
| **Optimum lower** | 1448 | 1.00 | 1.00 | 1.00 | 72 | 66.1-78.5 | 51.1 | 44.6-58.4 | 73.1 | 67.0-80.0 |
| **Optimum upper** | 1432 | 1.00 | 1.00 | 1.00 | 73.2 | 67.5-79.4 | 43.3 | 38.6-48.5 | 74 | 68.2-80.3 |
| **pH** |  |  |  |  |  |  |  |  |  |  |
| **Optimum lower** | 896 | 0.97 | 0.97 | 0.97 | 6.9 | 6.6-7.2 | 7.0 | 6.7-7.4 | 6.9 | 6.6-7.2 |
| **Optimum upper** | 879 | 0.96 | 0.96 | 0.96 | 7.1 | 6.8-7.3 | 7.2 | 6.9-7.4 | 7.0 | 6.8-7.3 |
| **NaCl (% w/v)** |  |  |  |  |  |  |  |  |  |  |
| **Optimum lower** | 470 | 0.99 | 0.99 | 0.99 | 2.22 | 1.72-2.81 | 2.30 | 1.69-3.04 | 2.2 | 1.68-2.82 |
| **Optimum upper** | 479 | 0.98 | 0.98 | 0.98 | 2.74 | 1.94-3.73 | 2.85 | 1.92-4.04 | 2.71 | 1.91-3.71 |

**Fig. 1.**
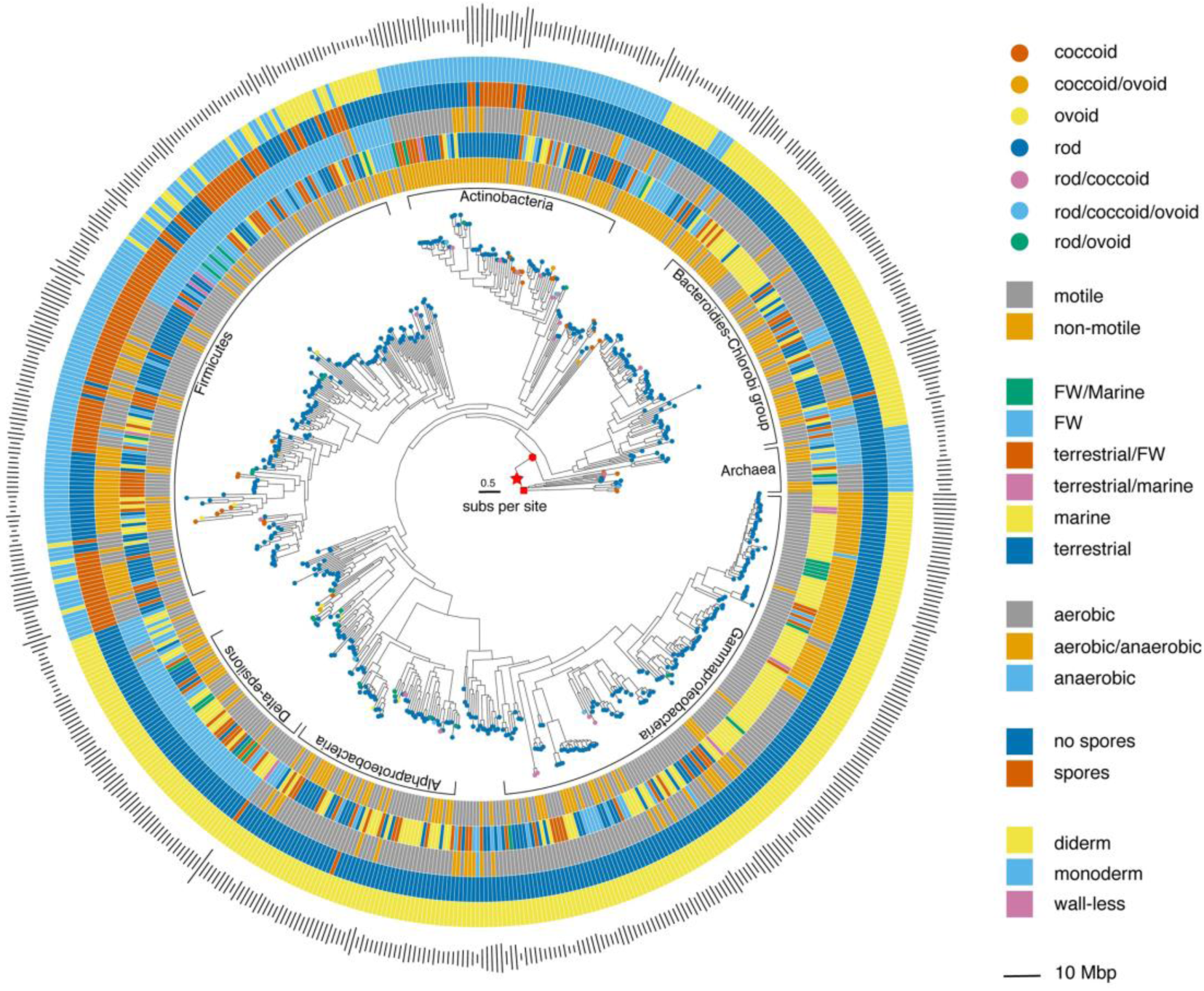
Distribution of phenotypic traits on the larger of the two phylogenetic trees used in the study. To aid clarity only species with complete data for all traits are included (n = 674). Tips on the tree are colour coded according to cell shape, while the **inner** to **outer** rings are coded according to motility status (**inner**), habitat (**second**), oxygen requirement (**third**), spore formation (**fourth**) and cell wall type (**outer**). The outer bars are proportional to genome size in Mbp. Red polygons at the base of the tree indicate the positions of LUCA (star), LACA (square) and LBCA (hexagon).

**Fig. 2.**
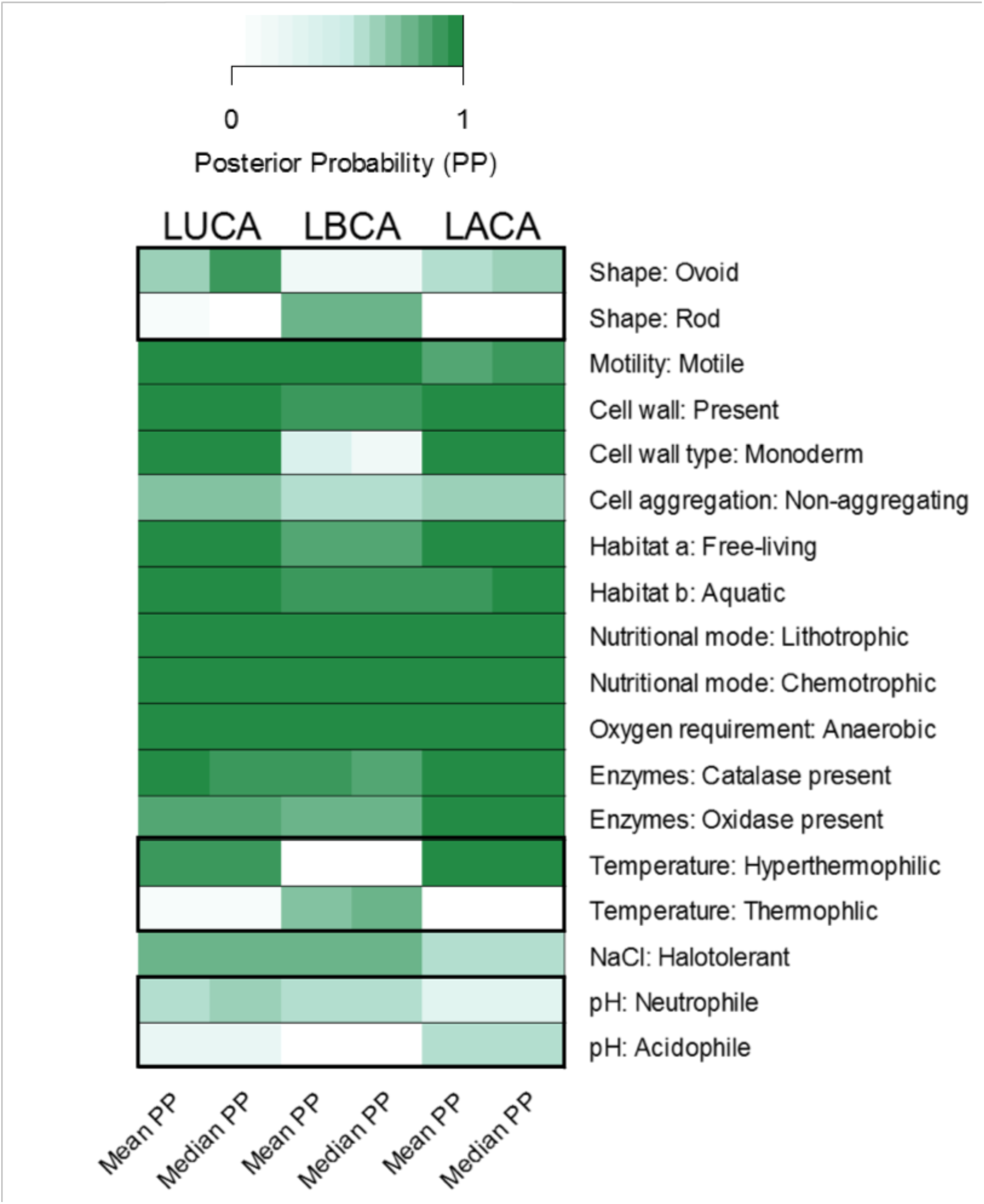
Bayesian estimates of the ancestral state for categorical traits. Mean and median posterior probabilities (PP) for each character state for LUCA, LBCA and LACA with colour intensity proportional to the value of the PP. On the right side the character state with the highest PP for each character is shown (e.g., Motility: Motile). When the ancestral character state differs between LUCA, LBCA or LACA, alternative character states are shown within thick boxes (i.e., shape, temperature and pH). Values are from the large tree, and only consistent results for ancestral character states between the large and the small trees are reported. Supplementary Table 1 contains the full results for all characters. Abbreviations: LUCA - last universal common ancestor; LBCA - last bacterial common ancestor; LACA - last archaeal common ancestor; PP – posterior probability.

Analyses with the two trees showed little qualitative difference and we present median and 95% probability intervals (95% PI) for the highest posterior probabilities (PP) in each analysis based on the larger of the two trees [37], with all estimates of uncertainty available in the SI. Where the two trees differed, this is noted. However, while the median and PI give some indication of the error associated with a given trait reconstruction it is more informative to examine the PP histograms given in the SI as these show the distribution and relationship between all trait value reconstructions, not just the most likely. These distributions often reveal a long tail in the probability distribution of a trait value, indicating a large error bound, but also a clear and narrow distribution of trait values close to a probability of zero for the other possible values.

Our probabilistic Bayesian phylogenetic models show clear phylogenetic signal in all 22 phenotypic traits we investigate (Table 1, Table S3 and Table S6). Where this is easily quantified (five continuous traits – see Methods), there is a minimum value of λ of 0.95 across the two trees (Tables 1 and Table S3). This signal indicates vertical inheritance of complex microbial traits and supports previous findings that trait (as opposed to gene) conservatism is widespread in bacteria and archaea [43,44]. Genetically complex traits, such as motility, shape, and the cell envelope are controlled by multiple genes [45–48] and while unlikely to be HGT free, the extent of this mechanism in the traits we study does not appear strong enough to blur their phylogenetic signal (see below and Methods).

Our analyses suggest that LUCA was likely actively motile (PP 0.98, 95%PI 0.85-0.99) with a cell wall (PP 1.00, 1.00-1.00) and a single cell membrane (monodermic cell plan (PP 1.00, 1.00-1.00, Figs. 2 & 3). Our reconstruction of LUCA with a cell wall is supported by the recent suggestion that it had genes associated with cell wall synthesis [4]. While the presence and nature of a cell membrane in LUCA has been debated [27,49,50], recent evidence points towards a mixed membrane of archaeal- and bacterial-like lipids [28,29,51]. Our results suggest that LUCA was halotolerant (PP 0.75, 0.64-0.81) with a preferred salinity range (2.31% - 2.76%) below that of modern seawater (3.1–3.8 %), hyperthermophilic (existing at temperatures above 70 °C, PP 0.92, 0.52-0.97) and probably living freely in an aquatic environment of neutral pH (6.93-7.05, PP 0.62, 0.17-0.76, next best PP acidophile: 0.24, 0.08-0.76) compared to modern seawater of pH 7.5 to 8.4 (Fig. 2, Tables 1, S1-S3). However, while we provide support for hyperthermophilia there is still debate over the presence or absence of reverse gyrase (a hyperthermophile-specific enzyme) in LUCA [4,34] and indeed whether LUCA was a thermophile [4,13]. There is however, evidence for thermostable ancestral enzymes reaching back between ∼1.4 and ∼4 Billion years [52,53]. We also note that an hyperthermophilic LUCA does not exclude a mesophile or ‘hot cross’ origin for life itself [18]. Most likely a chemolithotroph (chemotroph, PP 1.00, 1.00-1.00; lithotroph, PP 1.00, 0.99-1.00), using inorganic molecules to drive its metabolism, LUCA was probably adapted to anaerobic conditions (PP 1.00, 0.99-1.00).

**Fig. 3.**
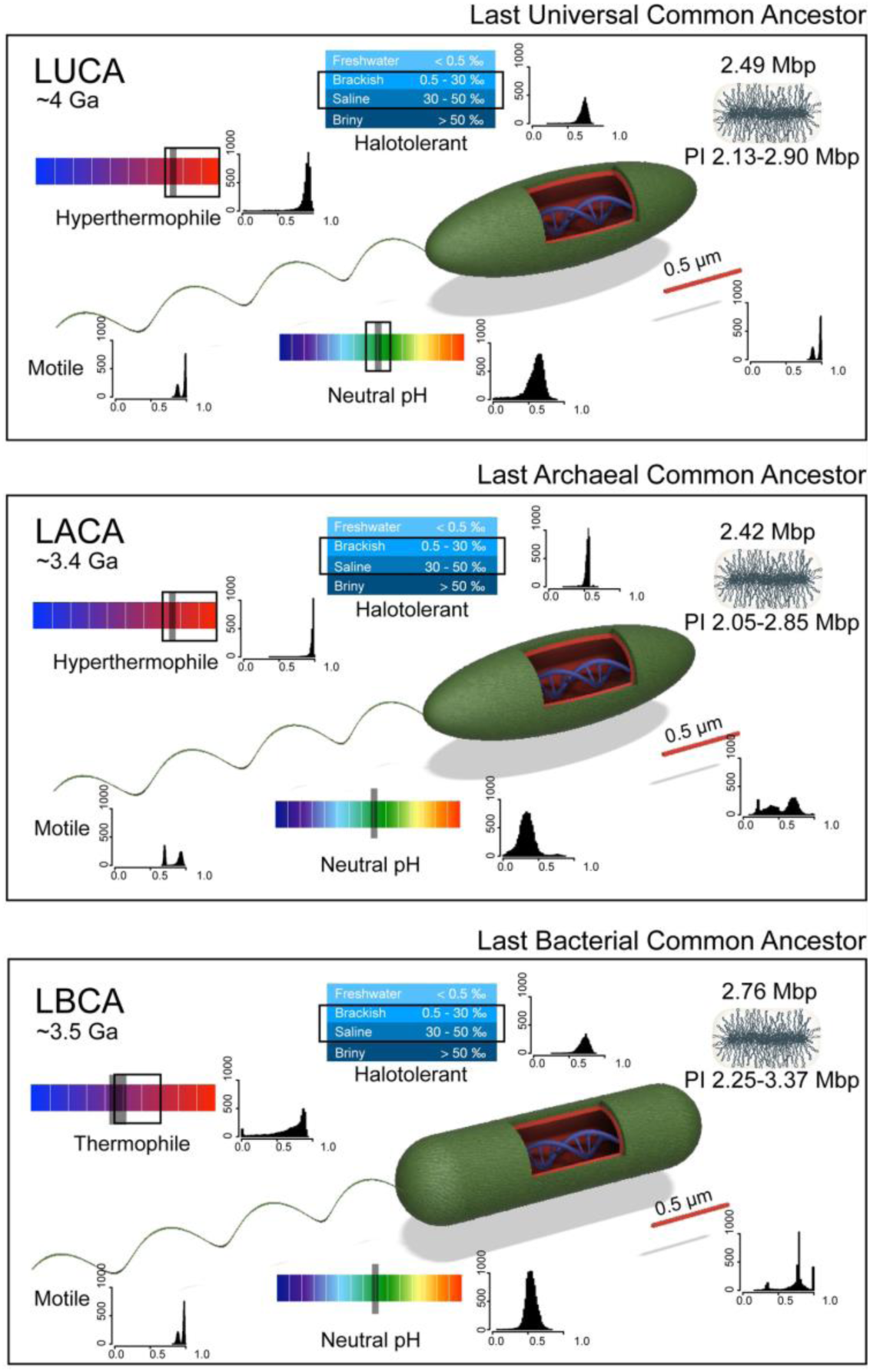
Phenotypic reconstructions of ancestral cells. (A) LUCA, (B) LBCA, (C) LACA. Flagella-like appendages indicate motility, green outer layer a cell wall and red inner layer a cell membrane. Where available, grey shading on habitat categories represents continuous data estimation while black rectangles bracket the categorical estimates. Histograms give posterior probability distributions for the discrete trait values. PI = 95% Probability Intervals.

Many of these reconstructed traits are indicative of the environment around hydrothermal vents, a habitat suggested for both the origins of life and the emergence of LUCA [4,54]. While modern vent systems are not generally of neutral pH [55], recent estimates for the early ocean suggest slightly acidic to neutral pH conditions [56] at the time of LUCA (∼ 4.0 Ga) [19,57] with values similar to our estimates (Table 1). Although an anaerobe, our reconstructions show that LUCA likely possessed both catalase (PP 0.93, 0.89-1.00) and oxidase (PP 0.84, 0.77-0.90) enzymes, suggesting an ability to resist oxidative stress. The presence of these enzymes might seem surprising given current evidence for free oxygen on early earth at concentrations less than 0.001% of those today [58], but is supported by genomic studies of Cytochrome oxidase genes [59], but see [60]. It has been hypothesized that LUCA might have possessed pathways to remove reactive oxygen species (ROS) [61] which is supported by our results, in particular, the presence of catalase [62]. While genes involved in resistance to oxidative stress might have arisen and been horizontally transferred between many ancestors after LUCA, recently Weiss et al. [4] identified eight of their 355 inferred LUCA proteins as enzymes involved in oxygen-dependent reactions.

Although bacteria and archaea exhibit a variety of morphologies, shape is usually well conserved within species [63]. Based on categorical data (‘rod-like’, ‘ovoid’ or ‘coccoid’) we provide evidence that LUCA was probably an ovoid cell (Fig. 2, PP 0.87, 0.08-0.95, Tables 1 & S1), implying that it had a complex cell cycle including an elongation phase (*sensu* [64]). Our estimates of cell dimensions also support an elongated cell with a length of 1.72 μm and width of 0.53 μm (95% PI 1.24-2.38 μm and 0.43-0.65 μm, Table 1). We were unable to draw a clear picture of pleomorphism in LUCA (PP < 0.6), but aggregation of individuals into clusters was likely not frequent (Fig. 2, PP 0.68, 0.55-0.85, Tables S1 & S2). The ability to form spores, while hypothesized as ancestral [65], was unclear (PP 0.01, 0.00-0.01 for the larger tree and PP 0.86, 0.13-0.99 for the smaller tree), likely owing to the uncertainty in the deep phylogenetic branching order of the two trees [37,38] (Tables S1 & S2, Fig. S3-S4). While some evidence exists for HGT of sporulation-related genes within the firmicutes [66], these transfers occurred locally and did not remove the global phylogenetic signal we recover from our data (Table S6).

The complex phenotypic picture we draw of LUCA implies to us that it must also have possessed a complex genome. We estimated LUCA’s ancestral genome size at 2.48 Mbp (95%PI 2.13-2.90 Mbp, Table 1), larger than the phylogenetically corrected average genome size in modern prokaryotes from our dataset (∼1.94 Mbp). Models allowing both gene losses (genome reduction) and gains (HGT) suggest that high levels of HGT lead to ancestral genome size estimates far smaller than modern genomes, while with limited HGT ancestral genome sizes become unrealistically large [67]. Unlike eukaryotes, prokaryotes show a consistent relationship between genome size and gene number [68]. Our estimate of the number of genes (PP 2624, 2261-3047) in LUCA (Table 1), assuming an average gene length for prokaryotes of 1024 bp and 13% of non-coding DNA in prokaryotic genomes [69], implies a genome size of around 2.34 Mbp, compared to our independent reconstruction of 2.48 Mbp (Methods).

We note that our approach differs fundamentally from gene-based studies in that the inherent complexity of the traits we examine acts to isolate them from the effects of HGT. The involvement of multiple genes and gene-clusters in determining the phenotypic values of many traits reduces the likelihood that they have been subject to HGT [70–72] and the close correspondence between the data and the trees (indicated by high λ, Table 1) gives us confidence that comparative phylogenetic methods can be used to understand the evolution of prokaryotic phenotypes and reliably estimate ancestral states as we have previously shown [40]. While HGT could have disrupted any phylogenetic signal in the trees we use, the majority of taxonomic groups are seen to be supported by both trees. Deep branching in the two trees does differ, but our approach of accounting for phylogenetic uncertainty (Methods) gives us confidence that these differences are unlikely to be due to HGT.

Irrespective of any network structure due to HGT, we recover a strong phylogenetic signal in all *traits* from both trees, implying that HGT does not rapidly act to change whole phenotypes or that it occurs mostly between very closely related lineages. While a positive correlation between the frequency of HGTs in distant lineages and genome size has been suggested [73], some studies indicate that HGT in prokaryotes also depends on both physical and genetic proximity [74–76]. Moreover, there is accumulating evidence to suggest that some of the traits that we present such as motility, cell envelope and sporulation, were subject to few or rare HGT events during their evolutionary history in a number of prokaryotic lineages [45,46,48,66,76].

As a further check we also performed simulations of HGT for each trait (See Methods, Fig. S2, SI) showing that our method is robust to HGT biases when estimating ancestral states. We were able to reconstruct ancestral nodes even in the presence of unrealistically high [77] rates of HGT. Thus, the strong phylogenetic signal that we and others [43,44,72,78] find for prokaryote phenotypic traits (rather than genes) demonstrates that the underlying phylogenetic relationships, even into deep time, are often conserved and that HGT does not always act to erase the phylogenetic signal for many complex traits.

Based on all of the above, we suggest that LUCA, the cenancestor, was more than a “half-alive” progenote [16,17], as advocated by Weiss et al. [4], and propose that it be considered a complex “prokaryotic” cell resembling modern archaea and bacteria. The complex phenotypic picture we depict of LUCA implies a complex genome, which is supported by our estimates of genome size and gene numbers. Indeed, establishment of biochemical pathways and complex enzymes over 3.5 Billion years ago has already been suggested [79,80]. Combined, these results seem to challenge the common assumption of increasing complexity through time, suggesting instead that cellular complexity arose near the very beginning of life and was retained or even lost through the evolution of the prokaryote lineage.

LUCA itself gave rise to two diverse and important domains of life, bacteria and archaea, whose ancestral phenotypic characteristics we also reconstructed. Our results suggest that both the last bacterial common ancestor (LBCA) and the last common ancestor of archaea (LACA) were similar to LUCA, in being free-living, motile organisms with a cell wall (LBCA free-living PP 0.82, 0.81-93, motile PP 0.97, 0.86-0.99, cell wall PP 0.93, 0.93-0.93; LACA free-living PP 0.98, 0.97-99, motile PP 0.90, 0.69-0.96, cell wall PP 1.00, 1.00-1.00). We were unable to draw a clear picture of pleomorphism or determine the pH conditions under which LBCA and LACA lived (PP <0.6, Tables 1 & S1) but as with LUCA neither were likely to be able to form aggregates (LBCA PP < 0.6, LACA PP 0.65, 0.55-0.80, Fig. 2, Tables S3 & S4). While LACA was likely non-spore forming (PP 0.99, 0.98-1.00) it is unclear whether LBCA was able to do so (large tree non-spore forming PP 0.95, 0.90-0.97, small tree spore forming PP 0.98, 0.96-0.99), likely owing to the uncertainty in deep phylogenetic branching of bacterial phyla between the two trees, (Tables S1-S2). As for LUCA, while there is some evidence for HGT of sporulation-related genes within the firmicutes, these transfers occurred locally and did not remove the global phylogenetic signal we recover from our data (Table S6).

It seems probable that LBCA was halotolerant (PP 0.75, 0.6-0.84), but evidence for this trait in LACA is very much weaker (Large tree PP <0.6; small tree PP 0.78, 0.00-1.00). Like LUCA, these progenitors of the bacterial and archaeal domains were probably chemolithotrophic, anaerobes with an ability to resist oxidative stress as suggested by the presence of catalase and oxidase (all PP > 0.8, all lower 95% PI >0.62, Table S1 and Fig. 2). In agreement with previous work [81–83] our results suggest that LBCA was a rod-like cell (PP ‘rod’ 0.79, 0.34-1.00 and length vs width in Table 1), implying that it had a complex life cycle with an elongation phase [83]. Our results also support the suggestion [82] that LBCA was gram positive, with a monodermic cell plan (PP 0.97, 0.38-1.00, Fig. 2, Table S1). More recently, and using different methods, LBCA has been reported as an anaerobic, rod-shaped cell utilising glucogenesis [84], and as rod shaped (with a diderm outer membrane), and motile [85], both studies depicting the bacterial ancestor in a very similar way to our own results.

In contrast to LBCA, LACA was potentially a slightly elongated ovoid cell (PP 0.61, 0.20-0.81 and length vs width in Table 1, Figs. 2c & 3), living in extremely hot aquatic environments (PP hyperthermophile 0.98, 0.90-0.99), as has been previously suggested [32,33,86] with optimal temperatures between 73 and 74 °C (95% PI 67-80 and 68.2-80.3, Fig. 2, Table 1). Very similar temperatures (∼73-76 °C) have been recently estimated for LACA based on a different approach [86]. Moreover, the absence of known RNA viruses that infect archaea may-indirectly support the idea of an ancestral hyperthermophilic archaeon [14,87]. As with LUCA, the complex phenotypic characters of both LBCA and LACA suggest to us that they too had large genomes, supported (as before) by our estimates of genome size (LBCA PP 2.76 Mb, 2.25-3.37; LACA PP 2.42 Mb, 2.05-2.85) and gene numbers (LBCA PP 2717, 2244-3290; LACA PP 2585, 2225-3003, Tables 1 and S3). However, Williams et al. [86] using different methods, estimated a genome half the size (∼1090 to 1328 genes) of our estimates for LACA, with a dataset of 62 archaeal genomes compared to 205 archaeal species in our study.

## Conclusion

While LACA, a descendant of LUCA, has been suggested to be a relatively complex organism based on gene number [86,88], our results have the potential to push cellular complexity back to the very beginnings of life. Barring the unlikelihood of panspermia [89], these results imply that complex phenotypic traits arose far earlier in the history of life than previously thought. Accumulating lines of evidence suggest that LUCA appeared very quickly during Earth’s history, perhaps as early as four billion years ago [19,57] during the Hadean or early Archaean eons. Evidence for black shales and banded iron formation in Greenland at ∼3.7 Ga is also suggestive of anoxygenic photosynthesis within 500 Million years of Earth becoming habitable [18]. Given that Earth formed around 4.5 Billion years ago [90], life probably arose in less than 500 million years (perhaps even within 200 million years [18,31]). This indicates that early life may have very quickly evolved considerable cellular complexity. We thus reveal LUCA as a potentially complex cell possessing a genetic code perhaps more intricate than many modern bacteria and archaea.

## Materials and Methods

### Experimental Design

#### 1. Data collection

##### 1.1. Phenotypic data

To reconstruct the ancestral states for LUCA, we collected all the relevant phenotypic data in the species description section from the Bergey’s Manual of Systematic Bacteriology that were consistently reported for many phyla and genera. The phenotypic traits (dataset: doi.org/10.6084/m9.figshare.12987509.v1) that were consistently reported for all the species included shape, cell size, pleomorphism, cell aggregation, motility, habitat, cell wall and spore formation, as well as catalase and oxidase activity, oxygen, temperature, pH and NaCl requirements and nutritional mode for 3,128 prokaryotic species from the phylogenies of Chai et al. [37] (large tree) and Segata et al. [38] (small tree). A breakdown of the number of species scored for each of the 28 traits for each phylogeny is given in table S7 (SI). These two trees were chosen to account for phylogenetic uncertainties as they were reconstructed from two different data sets and have different deep branching orders and different positions of the root. The main phyla in the two trees are monophyletic but the branching of these phyla differs between them. These differences can be explained by the different datasets used (∼400 different proteins for each, see Methods) as well as differences in the parameters used for phylogenetic reconstruction. The larger tree shows the Tenericutes and Proteobacteria as the deep branching phyla while the smaller tree indicates that the deepest branches are for the Thermotogae and Firmicutes. It is because of these differences that we use two different trees as this allows us to account for conflicting signals, especially for deep branching taxa. Despite the conflicting branching our results differ qualitatively only for sporulation and some LBCA – LACA comparisons (Tables S1 and S2). There is strong branch support in the large tree apart from those separating bacteria and archaea, while for the small tree the branch separating archaea from bacteria is supported by 100bp.

A representation of the large tree and the phylogenetic distribution of phenotypic data on it (where all traits were present for a species) are shown in Fig. 1. Data for most species was collated from the five volumes of Bergey’s Manual of Systematic Bacteriology [91– 95], but for species described after their publication data it was collected from the primary literature based on the List of Prokaryotic Names with Standing in Nomenclature (LPSN, Supplementary Text) [96]. Data for outlier strains (potentially misclassified species), were not included in the analysis (for example, *Clostridium difficile* strain P28 did not cluster with members of the same species). Bacterial and archaeal that have been delineated based on genomic data but have not been described phenotypically (not cultured, e.g., bacterial candidate phyla radiation (CPR) or Asgard archaea) were not included in the present study as no phenotypic data was available.

We did not include eukaryotes in our dataset the archaea as their origin within in the two-domain model means that they are somewhat secondary to our main question. The discovery of Asgard archaea, with genomes enriched in proteins previously considered specific to eukaryotes [97] and which branch closer to eukaryotes than other archaea in phylogenomic studies, imposes new constraints on eukaryogenesis and tree of life in general. Furthermore, the underlying complexity of the evolution of eukaryotes as a symbiotic union presents a number of very challenging obstacles to including them in any phylogenetic study of this type.

##### 1.2. Genomic data

Genome size and gene number data were collected for completely sequenced bacterial and archaeal genomes from the GOLD database [98, accessed March 2018]. Average genome size and average gene number per species were calculated from all the available strains of a given species. To independently estimate LUCA’s genome size we multiplied our estimated number of genes for LUCA (2629) by the estimated average gene length for prokaryotes from our data (1024 bp). Average gene length was estimated by regressing the genome size (Mbp) on the number of genes using Phylogenetic generalized least-squares (PGLS) implemented in the CAPER package to account for species non-independence due to shared ancestry [99]. Estimated genome size (gene number × gene length = 2.69 Mbp) was multiplied by the average ratio of coding versus non-coding sequences in prokaryotic genomes (0.87) based on Xu [69] to correct for non-coding sequences in the genome.

We note that our estimates do not include CPR, known for their small genome size [100], as no phenotypically described species existed for this group at the time of data collection. However, the position of the CPR in the bacteria tree of life) [101] suggests that they are mostly symbiotic and are derived (in particular in reference to their small genome) and therefore unlikely to influence the results of our reconstructions.

#### 2. Phenotypic characterization

##### 2.1. Cell morphology

Shape characterization in the species description sections from Bergey’s Manual of Systematic Bacteriology and the primary literature is qualitative, often subjective, and is not geometrically precise. A more reliable description would ideally be based on size measurements from individual cells. However, generally only cell length and width (diameter for coccoid cells) are available in the literature. These dimensions do however allow calculation of cell aspect ratio (AR) as length divided by width (Supplementary Text). While AR can be used to distinguish between coccoid and non-coccoid species, it cannot be used to distinguish categories within elongated, cylindrical cells (e.g. rod versus ovoid). Based on qualitative descriptions of cell morphology, we defined three broad shape categories for individual cells – rod, ovoid and coccoid (see Supplementary Text for a detailed description). For all the species with one shape category for which AR was available (2005 species) the delineation between coccoid and non-coccoid species based on qualitative descriptions was largely congruent with the AR data (Fig. S1, Supplementary Information).

##### 2.2. Pleomorphism

Pleomorphism describes the ability of some prokaryotes to alter their shape or size in response to environmental conditions [102]. To investigate the extent and evolution of pleomorphism we classified bacteria and archaea as either monomorphic or non-monomorphic.

##### 2.3. Motility

Motility was defined for the purpose of this study as the ability to move. Species were classified as motile or non-motile regardless of motility type (i.e. swarming, swimming, gliding or twitching motility).

##### 2.4. Cell aggregation

We classified species as either aggregating or non-aggregating, with cell aggregation defined as the ability to form an association between two or more cells. Species with cells occurring in pairs, tetrads, chains, sarcina, clusters or V-form, Y-form, T-form and Palisade groups or species able to form multicellular structures (e.g. fruiting bodies) were considered as aggregating. Species occurring only as single cells were coded as non-aggregating, while those occurring as single cells but with the ability to associate were coded as aggregating.

##### 2.5. Habitat

Due to limited availability of information on microenvironments we used three broad categorizations [40] of habitat types based on macro-environmental descriptions (i.e. the different locations where the organism naturally lives and grows and from which it could be recovered and isolated). Where habitat was not known, the first isolation site was used (e.g., human tissue, soil, shallow water, etc.). The first categorization set (habitat a) was a simple division into free-living (i.e., living independently in the environment) and non-free-living species (i.e., those associated with a host, manufactured products or isolated from industrial environments). Both free and non-free living species were further classified as Terrestrial or Aquatic (habitat b). For instance, species living in soil, fields or isolated from terrestrial sediments, were coded as Terrestrial, while an aquatic habitat was defined by living in marine or freshwater environments. Finally, we used a third habitat classification (habitat c) that included three categories: Terrestrial, Freshwater or Marine (see Supplementary Text for further details).

##### 2.6. Cell envelope

To test whether ancestral forms had a cell wall we classified species based on cell envelope type. Species with a description of a cell wall ultrastructure and/or described as Gram positive or Gram negative were all coded as having a cell wall. However, wall-less species (e.g., *Mycoplasma bovis*, see Supplementary Text) were considered as lacking a cell wall regardless of Gram stain. To investigate the ancestral state of cell plan organization, we further classified bacteria and archaea based on two cell types: monoderm (single membrane) and diderm (inner and outer membranes) based on information on cell envelope type and Gram reaction. Under this classification scheme species described as Gram positive or Gram negative were coded as having monodermic and didermic cell plans respectively, while those described as having a variable Gram reaction or staining negative but having a positive cell wall structure were coded as monoderms. Wall-less species were coded as monoderm as they possess a cell membrane. Species displaying a variable Gram reaction and without information on cell wall structure were not included in the analysis (see Supplementary Text for further details).

##### 2.7. Spore formation

We classified bacterial species as spore-forming or non-spore-forming based on clear statements in the species description and/or with a description of the spore type (e.g. “coccoid spores are formed centrally”). Archaea were classified as non-spore forming, and those species without information on spore-formation were not included in the analysis (see Supplementary Text for further details).

##### 2.8. Catalase, oxidase and oxygen requirements

We classified bacteria and archaea based on their ability to produce catalase and oxidase enzymes. Species described as having variable catalase or oxidase activities (i.e. positive or negative catalase or oxidase tests, 14 species in total) and those for which catalase activity was described as ‘weakly positive’ were excluded from the analysis. Species described as not usually producing an enzyme were recorded as not producing it rather than excluded from the analysis.

Species were recorded according to their ability to use oxygen for metabolic processes as aerobic, anaerobic, or both. Facultative aerobes or anaerobes were coded as both aerobic and anaerobic. Microaerophylic species (requiring oxygen to metabolize energy sources but not able to tolerate high concentrations) and microaerotolerant (tolerant of oxygen but unable to use it for growth) were not considered as a distinct category and were excluded from the analyses (42 species).

##### 2.9. Metabolism

The source of energy and electron donor that bacterial and archaeal species exploit was used to classify their metabolism. Data were extracted from species descriptions in the form of, for example, ‘chemoorganotroph’, ‘chemolithotroph’ or ‘phototroph’. Use of the photo- or chemo-prefix classified species phototrophs or chemotrophs respectively. For analysis of ancestral electron donor sources species described with words such as litho- or organo-were classified as lithotrophs or organotrophs respectively.

#### 3. Physicochemical parameters

For all species data on optimal temperature, pH and NaCl ranges were collected (lower and upper values) and used to estimate ancestral values based on a variable rate model (see Continuous traits). To assess whether these estimates were reliable we used a different approach based on categorical data (see Categorical traits) employing four key categories for temperature, pH and NaCl defined using optimal ranges (See dataset: doi.org/10.6084/m9.figshare.12987509.v1).

Species growing optimally in a temperature range of 5-20 ºC were classified as psychrotolerant, and those at temperatures between 20 to 45 ºC as mesophilic. Thermophilic species were defined as growing optimally at temperatures between 45 to 70 ºC, while hyperthermophiles were those with optimal growth temperatures above 70 ºC (see Supplementary Text). Optimal pH ranges were used to define neutrophiles (pH 5.5-7.5), acidophiles (pH 3-5.5), hyperacidophiles (pH 0-3) and alkaliphiles (pH 7.5-13). Optimal NaCl range was used to define non-halophiles (0-1% w/v), halotolerant (1-5% w/v), halophiles (5-20% w/v) and extreme-halophiles (20-32% w/v). Results from both approaches were compared to assess whether optimal lower and upper estimated ancestral values for temperature, pH and NaCl fell within the corresponding ancestrally reconstructed category (e.g., if the estimated lower and upper optimal temperature values were 72 ºC and 73 ºC, the ancestral reconstructed temperature category was hyperthermophile).

## Statistical Analysis

### Summary

Each phenotypic trait was analysed separately on each of the two independent trees, and these trees were pruned for each analysis to include only those species for which we had data for a specific trait. Bayesian methods using MCMC were used to estimate the parameters of evolutionary models that describe the evolution of a given trait along the branches of the phylogenetic tree. From this we derived a posterior distribution of ancestral states (sampled in proportion to their probability) that incorporates uncertainty in both the tree and the data. We were thus able to capture the error associated with the estimated ancestral state. We note that the MCMC models allow some relaxation of assumptions about trait evolution and do not necessarily assume evolution is gradual (see below).

#### 1. Ancestral state reconstruction

##### 1.2. Categorical traits

To allow us to account for model and parameter uncertainty a Bayesian rjMCMC [35] approach was used to reconstruct LUCAs ancestral states for all categorical traits (Table S1) using the Multistate method in the multicore version of BayesTraits V3.0.1 [103]. For the categorical data, species exhibiting more than one state of the same character were coded as having two states or more in BayesTraits (e.g. if a species exhibited both rod ‘R’ and coccoid ‘C’ forms it was coded as RC, see Cell morphology and dataset: doi.org/10.6084/m9.figshare.12987509.v1). For all the categorical data species with missing information (NAs, dataset) were excluded from the analyses.

The ‘AddMRCA’ command was used to estimate the ancestral states of the LBCA and LACA. All rjMCMC analyses were run using both the large [37] and small [38] Maximum Likelihood trees, reconstructed using 400 conserved proteins among 14,727 and 3,737 prokaryotic genomes respectively. All rjMCMC analyses were run in BayesTraits using the same parameters with all priors set to an exponential with a mean of 10. rjMCMC chains were run for 5,000,000 iterations with the first 10% as burn-in and sampling every 1,000 iterations after convergence. The exception was the analysis for pH, which was run for 20 million iterations with the first two million iterations as burn-in and a sampling interval of 2,000 to ensure chain convergence. MCMC chains were run three times per tree and chain mixing and convergence were assessed using Tracer v1.7 [104]. For all the analyses we report the mean and median posterior probability for the root (LUCA), LBCA and LACA, as well as the lower and upper HPD and 95% PI for the probability of the character state.

##### 1.2. Continuous traits

Ancestral values for cell size, genome size, and gene number, as well as optimal pH, NaCl and temperature were estimated for LUCA, LBCA and LACA using a variable rates model [105], implemented in BayesTraits, that detects heterogeneous rates of evolutionary changes along the tree within a Bayesian framework. This method can be used to detect evolutionary trends and has been shown to estimate ancestral states more accurately than other methods [106].

Following the methodology in Baker et al. [106] we ran the variable-rates model for each continuous trait (the MCMC chain was run for 1 billion iterations, sampling every 500,000 iterations after a burn-in period of 500 million iterations). This approach detects branches or clades of the tree that have undergone especially fast or slow rates of change, it stretches (increased rate) or compresses (decreased rate) branch lengths by an amount reflecting the inferred rate of evolution in that branch. Specifically, the method relaxes the BM model assumptions of gradual evolution [105].

We summed all the rate-scaled branches along the phylogenetic path of each species from the root of the tree to the terminal branch (root-to-tip rate) – we use the median rate from the posterior distribution to rate-scale the branches. To test for a trend, we regressed the value of the continuous trait under investigation with root-to-tip rate using a Bayesian MCMC phylogenetic generalized least squares (GLS) regression [99,107] (the MCMC chain was run for 1 billion iterations with a burn-in of 500 million iterations and a sampling interval of 500,000 iterations). We used the proportion of time (Px) the regression coefficient determined the crossed zero to determine significance where Px < 0.05 we declared a significant trend.

We estimate the ancestral values for LUCA, LACA and LBCA for all continuous traits using the phylogenetic prediction method originally described in Organ et al. [108], where we find a significant trend we account for that as in Baker et al. [106]. Each of our MCMC analyses was repeated three times to ensure convergence was achieved – we report the results from one randomly chosen chain for each analysis.

As recommended in Venditti et al. [105], we randomly removed all except one species in a clade where the continuous data were identical. We do this, as from a statistical modelling point of view, the best fit to such a group of species with exactly the same size would be to infer that no evolutionary change occurred in that group. This is an unrealistic scenario. Using this reduced set of species, we also repeated our Categorical analyses for pH, temperature and NaCl, finding concordant results (Tables S4-S5). Continuous data were log10 transformed prior to analysis, except NaCl which were cube-root transformed owing to the presence of zeros, and pH where the original values were used as the scale is logarithmic.

In contrast to rod or ovoid shape species, coccoids are evolutionarily constrained and thus can only divide and not elongate [64]. We therefore excluded data from coccoid species with an Aspect Ratio (AR) = 1 (see Cell morphology and Supplementary Text) prior to estimation of ancestral cell dimensions for LUCA, LACA and LBCA.

### Phylogenetic signal

Phylogenetic signal is a measure of the statistical dependence between trait values as a result of phylogenetic relationships between species. To evaluate the extent of the phylogenetic signal in our data for continuous traits the phylogenetic signal was estimated using Maximum likelihood and the variable rates model and λ values [99] were selected from the best of two competing models - one with λ set to zero and one for which λ was estimated. Estimation of λ gives a transformation of a phylogeny that offers the best fit of the trait data to a Brownian Motion (BM) model of evolution. Thus, if λ = 0 we infer no phylogenetic signal in the trait data, if λ = 1 this corresponds to a BM model, with intermediate values possible.

For discrete traits we reconstructed the ancestral states for each trait using a Markov transition model in a MCMC Bayesian context [109] (Table S6). This model estimates instantaneous transition rates and so allows traits to change at any point on the branches of the phylogeny. We then compared the posterior probabilities with a randomized data set from each of the two trees. We concluded that there is phylogenetic signal if we were able to discriminate between the states of a given trait (e.g. P(0) = 0.99 and P(1) = 0.01) in our data while simultaneously obtaining equal posterior probabilities for these states of the same trait from the randomized data set (Table S6).

#### 2. Horizontal trait transfer simulations

While HGT plays a role in prokaryotic evolution, often seen as obscuring the assumption of vertical descent with modification, the traits under investigation in our study are multi-gene entities and as such our ancestral reconstructions are less likely to be influenced by HGT [70–72]. Furthermore, a number of studies show that HGT occurs predominantly among near phylogenetic relatives and closely interacting species [110–112] and is far less common between distantly related organisms (but see [73,113]). To test the accuracy of our reconstruction methods to cases of horizontal trait transfer (HTT) we conduct two sets of simulations. We call our simulation process HTT rather than HGT as we simulate the extreme case where the trait itself is transferred between phylogenetic branches – this represents a suite of functional genes associated with a particular trait transferred and instantly fixed in the population. HTT is thus a much more severe test of HGT than a gradual replacement model as it implies the transfer of an entire operon or the entire set of genes coding for the trait. We note that similar conclusions have been drawn for both continuous and discrete traits in human language [114,115].

Using the Chai et al. [37] (i.e., larger) phylogenetic tree we randomly identified between 0 and 1000 (in 200 increments, see Fig. S2) points along the branches. We simulated a trait via Brownian motion (variance = 1) along the branches. When the simulation reached one of the randomly chosen points, an HTT event occurred from the ‘donor’ (red points in Fig. S2 a and b) to one of the potential ‘recipients’ (yellow squares in Fig. S2 a and b). We ran simulations for two types of HTT events. Firstly, a *Local* version, where HTT occurred between near phylogenetic relatives (Fig. S2a) – here, at the HTT point (and example is provided as a red point in Fig. S2) we selected the recipient branch from the clade defined by the most recent common ancestor (i.e., one phylogenetic node back) of the HTT event (in our example the potential recipient branches are shown with a yellow squares). Where there is more than one recipient branch (as in Fig. S2a) we randomly select one to receive the horizontally transferred trait. At the point of transfer the recipient branch and the donor have the same trait value – following this the Brownian motion process continues. We also simulate a very severe *Global* process which is identical to the *Local* simulation apart from the fact that at the HTT point the potential recipient points can be chosen from all branches at that time point (i.e. from the root of the tree rather than the most recent common ancestor), this is illustrated in Fig. S2b. Through the simulation process we record the trait value at each node of the tree – these are the ‘known’ ancestral states. Following the simulation process, we used the data at the terminal branches (the end point of the simulation at tips of the tree) to infer the ancestral states at each node in the phylogeny. We compare these with the known ancestral states for each simulation condition. We repeat the process 1000 times for each condition for both the global and the local set – a total of 12000 simulations.

We use linear regression to compare the known to inferred ancestral state values. We expect, where the reconstructions are perfectly accurate, to see a slope (regression coefficient) equal to one, an intercept equal to zero, and a coefficient of determination near one. We also recorded the estimated Brownian motion variance from the inference stage – the simulated variance was one. The results show that *Local* HTT does not significantly influence our ability to accurately infer ancestral states even up to a point where there are 1000 such events in the tree (Fig. S2c-f). As expected, the accuracy of the unrealistically severe *Global* simulations reduces with the number of HTT events – specifically the variance in the intercept increases and the variance and magnitude of the Brownian motion variance increases.

## ACKNOWLEDGMENTS

We thank, O. Guadayol Roig, M. El Baidouri and R. Schuech, Ciara O’Donovan, Manabu Sakamoto and Joanna Baker for helpful comments and advice on the draft MS; Comments from Norman Sleep, Hervé Sauquet and three anonymous reviewers greatly improved the manuscript. Funding: This work is financially supported by a Leverhulme Trust Research Leadership Award (RL-2012-022), granted to S.H. and a Leverhulme Trust Research Project Grant (RPG-2013-185) to C.V.

## Author contributions

F.E.B., C.V. and S.H designed the study; F.E.B., S.S, and S.H. developed the protocol for the data collection; F.E.B. and S.S. collected the data and F.E.B., C.V., S.S., A.M. and S.H. analysed the data. F.E.B., C.V., and S.H. wrote the first draft of the manuscript, and all authors contributed substantially to revisions.

## References

1. Doolittle WF. The nature of the universal ancestor and the evolution of the proteome. Curr Opin Struct Biol. 2000;10: 355–358. doi:10.1016/S0959-440X(00)00096-8

2. Fitch WM, Ayala FJ, Doolittle WF, Brown JR. “Tempo and Mode in Evoluton” organized Tempo, mode, the progenote, and the universal root. 1994;91: 6721–6728.

3. Koonin E V. Comparative genomics, minimal gene-sets and the last universal common ancestor. Nat Rev Microbiol. 2003;1: 127–136. doi:10.1038/nrmicro751

4. Weiss MC, Sousa FL, Mrnjavac N, Neukirchen S, Roettger M, Nelson-Sathi S, et al. The physiology and habitat of the last universal common ancestor. Nat Microbiol. 2016;1: 16116. doi:10.1038/nmicrobiol.2016.116

5. Carroll SB. Chance and necessity: The evolution of morphological complexity and diversity. Nature. 2001;409: 1102–1109. doi:10.1038/35059227

6. Hedges SB, Blair JE, Venturi ML, Shoe JL. A molecular timescale of eukaryote evolution and the rise of complex multicellular life. BMC Evol Biol. 2004;4: 2. doi:10.1186/1471-2148-4-2

7. Valentine JW, Collins AG, Meyer CP. Morphological complexity increase in metazoans. Paleobiology. 1994;20: 131–142. doi:10.1017/S0094837300012641

8. Wolf YI, Koonin E V. Genome reduction as the dominant mode of evolution. BioEssays. 2013;35: 829–837. doi:10.1002/bies.201300037

9. Giovannoni SJ, Cameron Thrash J, Temperton B. Implications of streamlining theory for microbial ecology. ISME Journal. 2014. pp. 1553–1565. doi:10.1038/ismej.2014.60

10. Williams TA, Foster PG, Cox CJ, Embley TM. An archaeal origin of eukaryotes supports only two primary domains of life. Nature. 2013. pp. 231–236. doi:10.1038/nature12779

11. Spang A, Saw JH, Jørgensen SL, Zaremba-Niedzwiedzka K, Martijn J, Lind AE, et al. Complex archaea that bridge the gap between prokaryotes and eukaryotes. Nature. 2015;521: 173–179. doi:10.1038/nature14447

12. Williams TA, Cox CJ, Foster PG, Szöllősi GJ, Embley TM. Phylogenomics provides robust support for a two-domains tree of life. Nat Ecol Evol. 2020;4: 138–147. doi:10.1038/s41559-019-1040-x

13. Berkemer SJ, McGlynn SE. A New Analysis of Archaea-Bacteria Domain Separation: Variable Phylogenetic Distance and the Tempo of Early Evolution. Mol Biol Evol. 2020. doi:10.1093/molbev/msaa089

14. Krupovic M, Dolja V V., Koonin E V. The LUCA and its complex virome. Nat Rev Microbiol. 2020;18: 661–670. doi:10.1038/s41579-020-0408-x

15. Woese CR, Fox GE. Phylogenetic structure of the prokaryotic domain: The primary kingdoms (archaebacteria/eubacteria/urkaryote/16S ribosomal RNA/molecular phylogeny). 1977.

16. Woese CR, Fox GE. The concept of cellular evolution. J Mol Evol. 1977. doi:10.1007/BF01796132

17. Gogarten JP, Deamer D. Is LUCA a thermophilic progenote ? Nat Publ Gr. 2016;1: 1–2. doi:10.1038/nmicrobiol.2016.229

18. Sleep NH. Geological and geochemical constraints on the origin and evolution of life. Astrobiology. 2018;18: 1199–1219. doi:10.1089/ast.2017.1778

19. Betts HC, Puttick MN, Clark JW, Williams TA, Donoghue PCJ, Pisani D. Integrated genomic and fossil evidence illuminates life’s early evolution and eukaryote origin. Nat Ecol Evol. 2018;2: 1556–1562. doi:10.1038/s41559-018-0644-x

20. Ouzounis CA, Kunin V, Darzentas N, Goldovsky L. A minimal estimate for the gene content of the last universal common ancestor - Exobiology from a terrestrial perspective. Res Microbiol. 2006;157: 57–68. doi:10.1016/j.resmic.2005.06.015

21. Brbić M, Piškorec M, Vidulin V, Kriško A, Šmuc T, Supek F. The landscape of microbial phenotypic traits and associated genes. Nucleic Acids Res. 2016;44: 10074–10090. doi:10.1093/nar/gkw964

22. Edgar RC. SINAPS: Prediction of microbial traits from marker gene sequences. bioRxiv. 2017. doi:doi.org/10.1101/124156

23. Doolittle WF. Uprooting the tree of life. Scientific American. 2000. doi:10.1038/scientificamerican0200-90

24. Puigb P, Wolf YI, Koonin E V. Search for a “tree of Life” in the thicket of the phylogenetic forest. J Biol. 2009. doi:10.1186/jbiol159

25. Kellner S, Spang A, Offre P, Szöllősi GJ, Petitjean C, Williams TA. Genome size evolution in the Archaea. Emerg Top Life Sci. 2018. doi:10.1042/etls20180021

26. Koonin E V, Krupovic M, Ishino S, Ishino Y. The replication machinery of LUCA: common origin of DNA replication and transcription. BMC Biol. 2020;18: 61. doi:10.1186/s12915-020-00800-9

27. Koonin E V., Martin W. On the origin of genomes and cells within inorganic compartments. Trends Genet. 2005;21: 647–654. doi:10.1016/j.tig.2005.09.006

28. Villanueva L, von Meijenfeldt FAB, Westbye AB, Hopmans EC, Dutilh BE, Sinninghe Damste JS. Bridging the divide: bacteria synthesizing archaeal membrane lipids. bioRxiv. 2018; 448035. doi:https://doi.org/10.1101/448035

29. Caforio A, Siliakus MF, Exterkate M, Jain S, Jumde VR, Andringa RLH, et al. Converting Escherichia coli into an archaebacterium with a hybrid heterochiral membrane. Proc Natl Acad Sci U S A. 2018;115: 3704–3709. doi:10.1073/pnas.1721604115

30. Sleytr UB, Schuster B, Egelseer EM, Pum D. S-layers: Principles and applications. FEMS Microbiol Rev. 2014. doi:10.1111/1574-6976.12063

31. Weiss MC, Preiner M, Xavier JC, Zimorski V, Martin WF. The last universal common ancestor between ancient Earth chemistry and the onset of genetics. Achtman M, editor. PLOS Genet. 2018;14: e1007518. doi:10.1371/journal.pgen.1007518

32. Boussau B, Blanquart S, Necsulea A, Lartillot N, Gouy M. Parallel adaptations to high temperatures in the Archaean eon. Nature. 2008. doi:10.1038/nature07393

33. Groussin M, Gouy M. Adaptation to environmental temperature is a major determinant of molecular evolutionary rates in archaea. Mol Biol Evol. 2011. doi:10.1093/molbev/msr098

34. Catchpole RJ, Forterre P. The Evolution of Reverse Gyrase Suggests a Nonhyperthermophilic Last Universal Common Ancestor. Mol Biol Evol. 2019;36: 2737–2747. doi:10.1093/molbev/msz180

35. Pagel M, Meade A, Barker D. Bayesian estimation of ancestral character states on phylogenies. Syst Biol. 2004;53: 673–684. doi:10.1080/10635150490522232

36. Pagel M. Bayesian analysis of correlated evolution of discrete characters by reversible-jump Markov chain Monte Carlo. Am Nat. 2006;167: 808–825. doi:10.1086/503444

37. Chai J, Kora G, Ahn TH, Hyatt D, Pan C. Functional phylogenomics analysis of bacteria and archaea using consistent genome annotation with UniFam. BMC Evol Biol. 2014;14: 1–13. doi:10.1186/s12862-014-0207-y

38. Segata N, Börnigen D, Morgan XC, Huttenhower C. PhyloPhlAn is a new method for improved phylogenetic and taxonomic placement of microbes. Nat Commun. 2013;4: 2304. doi:10.1038/ncomms3304

39. Cornwell W, Nakagawa S. Phylogenetic comparative methods. Current Biology. 2017. pp. R333–R336. doi:10.1016/j.cub.2017.03.049

40. El Baidouri F, Venditti C, Humphries S. Independent evolution of shape and motility allows evolutionary flexibility in Firmicutes bacteria. Nat Ecol Evol. 2016;1: 0009. doi:10.1038/s41559-016-0009

41. Camprubí E, de Leeuw JW, House CH, Raulin F, Russell MJ, Spang A, et al. The Emergence of Life. Space Sci Rev. 2019;215: 56. doi:10.1007/s11214-019-0624-8

42. McShea DW. The evolution of complexity without natural selection, a possible large-scale trend of the fourth kind. Paleobiology. 2005;31: 146–156. doi:10.1666/0094-8373(2005)031[0146:TEOCWN]2.0.CO;2

43. Martiny AC, Treseder K, Pusch G. Phylogenetic conservatism of functional traits in microorganisms. ISME J. 2013;7: 830–838. doi:10.1038/ismej.2012.160

44. Goberna M, Verdú M. Predicting microbial traits with phylogenies. ISME J. 2016;10: 959–967. doi:10.1038/ismej.2015.171

45. Liu R, Ochman H. Stepwise formation of the bacterial flagellar system. Proc Natl Acad Sci U S A. 2007;104: 7116–7121. doi:10.1073/pnas.0700266104

46. Desmond E, Brochier-Armanet C, Gribaldo S. Phylogenomics of the archaeal flagellum: rare horizontal gene transfer in a unique motility structure. BMC Evol Biol. 2007;7: 106. doi:10.1186/1471-2148-7-106

47. Jiang C, Caccamo PD, Brun Y V. Mechanisms of bacterial morphogenesis: Evolutionary cell biology approaches provide new insights. BioEssays. 2015;37: 413–425. doi:10.1002/bies.201400098

48. Antunes LCS, Poppleton D, Klingl A, Criscuolo A, Dupuy B, Brochier-Armanet C, et al. Phylogenomic analysis supports the ancestral presence of LPS-outer membranes in the Firmicutes. Elife. 2016;5. doi:10.7554/eLife.14589

49. Koga Y, Kyuragi T, Nishihara M, Sone N. Did archaeal and bacterial cells arise independently from noncellular precursors? A hypothesis stating that the advent of membrane phospholipid with enantiomeric glycerophosphate backbones caused the separation of the two lines of descent. J Mol Evol. 1998;46: 54–63. doi:10.1007/PL00006283

50. Peretó J, López-García PN, Moreira D, López-García P, Moreira D. Ancestral lipid biosynthesis and early membrane evolution. Trends Biochem Sci. 2004;29: 469–477. doi:10.1016/j.tibs.2004.07.002

51. Harris AJ, Goldman AD. The very early evolution of protein translocation across membranes. PLoS Comput Biol. 2021;17. doi:10.1371/journal.pcbi.1008623

52. Ingles-Prieto A, Ibarra-Molero B, Delgado-Delgado A, Perez-Jimenez R, Fernandez JM, Gaucher EA, et al. Conservation of Protein Structure over Four Billion Years. Structure. 2013;21: 1690–1697. doi:10.1016/J.STR.2013.06.020

53. Perez-Jimenez R, Inglés-Prieto A, Zhao Z, Sanchez-Romero I, Alegre-Cebollada J, Kosuri P, et al. Single-molecule paleoenzymology probes the chemistry of resurrected enzymes. Nat Struct Mol Biol. 2011;18: 592. doi:10.1038/NSMB.2020

54. Baross JA, Hoffman SE. Submarine hydrothermal vents and associated gradient environments as sites for the origin and evolution of life. Orig Life Evol Biosph. 1985;15: 327–345. doi:10.1007/BF01808177

55. Ding K, Seyfried WE, Zhang Z, Tivey MK, Von Damm KL, Bradley AM. The in situ pH of hydrothermal fluids at mid-ocean ridges. Earth Planet Sci Lett. 2005;237: 167–174. doi:10.1016/j.epsl.2005.04.041

56. Krissansen-Totton J, Arney GN, Catling DC. Constraining the climate and ocean pH of the early Earth with a geological carbon cycle model. Proc Natl Acad Sci U S A. 2018;115: 4105–4110. doi:10.1073/pnas.1721296115

57. Tashiro T, Ishida A, Hori M, Igisu M, Koike M, Méjean P, et al. Early trace of life from 3.95 Ga sedimentary rocks in Labrador, Canada. Nature. 2017;549: 516–518. doi:10.1038/nature24019

58. Lyons TW, Reinhard CT, Planavsky NJ. The rise of oxygen in Earth’s early ocean and atmosphere. Nature. 2014. pp. 307–15. doi:10.1038/nature13068

59. Castresana J, Lübben M, Saraste M, Higgins DG. Evolution of cytochrome oxidase, an enzyme older than atmospheric oxygen. EMBO J. 1994;13: 2516–2525. doi:10.1002/j.1460-2075.1994.tb06541.x

60. Dibrova D V., Shalaeva DN, Galperin MY, Mulkidjanian AY. Emergence of cytochrome bc complexes in the context of photosynthesis. Physiol Plant. 2017;161: 150–170. doi:10.1111/ppl.12586

61. Slesak I, Slesak H, Kruk J. Oxygen and hydrogen peroxide in the early evolution of life on earth: in silico comparative analysis of biochemical pathways. Astrobiology. 2012;12: 775–784. doi:10.1089/ast.2011.0704

62. Heck DE, Shakarjian M, Kim HD, Laskin JD, Vetrano AM. Mechanisms of oxidant generation by catalase. Ann N Y Acad Sci. 2010;1203: 120–125. doi:10.1111/j.1749-6632.2010.05603.x

63. Young KD. The Selective Value of Bacterial Shape. Microbiol Mol Biol Rev. 2006;70: 660–703. doi:10.1128/mmbr.00001-06

64. Philippe J, Vernet T, Zapun A. The elongation of ovococci. Microbial Drug Resistance. 2014. pp. 215–221. doi:10.1089/mdr.2014.0032

65. Tocheva EI, Ortega DR, Jensen GJ. Sporulation, bacterial cell envelopes and the origin of life. Nature Reviews Microbiology. 2016. pp. 535–542. doi:10.1038/nrmicro.2016.85

66. Ramos-Silva P, Serrano M, Henriques AO. From Root to Tips: Sporulation Evolution and Specialization in Bacillus subtilis and the Intestinal Pathogen Clostridioides difficile. Mol Biol Evol. 2019;36: 2714–2736. doi:10.1093/molbev/msz175

67. Dagan T, Martin W. Ancestral genome sizes specify the minimum rate of lateral gene transfer during prokaryote evolution. Proc Natl Acad Sci. 2007;104: 870–875. doi:10.1073/pnas.0606318104

68. Konstantinidis KT, Tiedje JM. Trends between gene content and genome size in prokaryotic species with larger genomes. Proc Natl Acad Sci. 2004;101: 3160–3165. doi:10.1073/pnas.0308653100

69. Xu L, Chen H, Hu X, Zhang R, Zhang Z, Luo ZW. Average gene length is highly conserved in prokaryotes and eukaryotes and diverges only between the two kingdoms. Mol Biol Evol. 2006;23: 1107–1108. doi:10.1093/molbev/msk019

70. Jain R, Rivera MC, Lake JA. Horizontal gene transfer among genomes: The complexity hypothesis. Proc Natl Acad Sci U S A. 1999. doi:10.1073/pnas.96.7.3801

71. Wellner A, Lurie MN, Gophna U. Complexity, connectivity, and duplicability as barriers to lateral gene transfer. Genome Biol. 2007;8: R156. doi:10.1186/gb-2007-8-8-r156

72. Martiny JBH, Jones SE, Lennon JT, Martiny AC. Microbiomes in light of traits: A phylogenetic perspective. Science (80-). 2015;350: aac9323–aac9323. doi:10.1126/science.aac9323

73. Cordero OX, Hogeweg P. The impact of long-distance horizontal gene transfer on prokaryotic genome size. Proc Natl Acad Sci U S A. 2009;106: 21748–21753. doi:10.1073/pnas.0907584106

74. Kloesges T, Popa O, Martin W, Dagan T. Networks of gene sharing among 329 proteobacterial genomes reveal differences in lateral gene transfer frequency at different phylogenetic depths. Mol Biol Evol. 2011;28: 1057–1074. doi:10.1093/molbev/msq297

75. Williams D, Gogarten JP, Papke RT. Quantifying homologous replacement of loci between haloarchaeal species. Genome Biol Evol. 2012;4: 1223–1244. doi:10.1093/gbe/evs098

76. Jeong H, Arif B, Caetano-Anollés G, Kim KM, Nasir A. Horizontal gene transfer in human-associated microorganisms inferred by phylogenetic reconstruction and reconciliation. Sci Rep. 2019;9: 5953. doi:10.1038/s41598-019-42227-5

77. Cohen O, Gophna U, Pupko T. The Complexity Hypothesis Revisited: Connectivity Rather Than Function Constitutes a Barrier to Horizontal Gene Transfer. Mol Biol Evol. 2011;28: 1481–1489. doi:10.1093/molbev/msq333

78. Barberán A, Caceres Velazquez H, Jones S, Fierer N. Hiding in Plain Sight: Mining Bacterial Species Records for Phenotypic Trait Information. mSphere. 2017;2: e00237–17. doi:10.1128/msphere.00237-17

79. Perez-Jimenez R, Inglés-Prieto A, Zhao Z-M, Sanchez-Romero I, Alegre-Cebollada J, Kosuri P, et al. Single-molecule paleoenzymology probes the chemistry of resurrected enzymes. Nat Struct Mol Biol 2011 185. 2011;18: 592–596. doi:10.1038/nsmb.2020

80. Nisbet EG, Sleep NH. The habitat and nature of early life. Nature. Nature; 2001. pp. 1083–1091. doi:10.1038/35059210

81. Siefert JL, Fox GE. Phylogenetic mapping of bacterial morphology. Microbiology. 1998;144: 2803–2808. doi:10.1099/00221287-144-10-2803

82. Koch AL. Were Gram-positive rods the first bacteria? Trends Microbiol. 2003;11: 166–170. doi:10.1016/S0966-842X(03)00063-5

83. Yulo PRJ, Hendrickson HL. The evolution of spherical cell shape; progress and perspective. Biochemical Society Transactions. 2019. pp. 1621–1634. doi:10.1042/BST20180634

84. Xavier JC, Gerhards RE, Wimmer JLE, Brueckner J, Tria FDK, Martin WF. The metabolic network of the last bacterial common ancestor. Commun Biol. 2021;4: 413. doi:10.1038/s42003-021-01918-4

85. Coleman GA, Davín AA, Mahendrarajah TA, Szánthó LL, Spang A, Hugenholtz P, et al. A rooted phylogeny resolves early bacterial evolution. Science (80-). 2021;372: eabe0511. doi:10.1126/science.abe0511

86. Williams TA, Szöllosi GJ, Spang A, Foster PG, Heaps SE, Boussau B, et al. Integrative modeling of gene and genome evolution roots the archaeal tree of life. Proc Natl Acad Sci U S A. 2017;114: E4602–E4611. doi:10.1073/pnas.1618463114

87. Forterre P. The common ancestor of archaea and eukarya was not an archaeon. Archaea. 2013. doi:10.1155/2013/372396

88. Wolf YI, Makarova KS, Yutin N, Koonin E V. Updated clusters of orthologous genes for Archaea: a complex ancestor of the Archaea and the byways of horizontal gene transfer. Biol Direct. 2012;7: 46. doi:10.1186/1745-6150-7-46

89. Wesson PS. Panspermia, past and present: Astrophysical and biophysical conditions for the dissemination of life in space. Space Sci Rev. 2010;156: 239–252. doi:10.1007/s11214-010-9671-x

90. Jackson MG, Carlson RW, Kurz MD, Kempton PD, Francis D, Blusztajn J. Evidence for the survival of the oldest terrestrial mantle reservoir. Nature. 2010;466: 853–856. doi:10.1038/nature09287

91. Kuever J, Rainey FA, Widdel F. Bergey’s Manual® of Systematic Bacteriology. Bergey’s Manual® of Systematic Bacteriology. 2001. doi:10.1007/978-0-387-21609-6

92. Brenner DJ, Krieg NR, Staley JT. Bergey’s Manual® of Systematic Bacteriology. Bergey’s Manual® of Systematic Bacteriology. 2005. doi:10.1007/0-387-29298-5

93. De Vos P, Garrity GM, Jones D, Krieg NR, Ludwig W, Rainey FA, et al. Bergey’s manual of systematic bacteriology Volume Three The Firmicutes. Bergey’s Manual of Systematic Bacteriology. Springer Science & Business Media; 2009. doi:10.1007/978-0-387-68489-5

94. Krieg NR, Staley JT, Brown DR, Hedlund BP, Paster BJ, Ward NL, et al. Bergey’s Manual® of Systematic Bacteriology. Bergey’s Manual® of Systematic Bacteriology. 2010. doi:10.1007/978-0-387-68572-4

95. Bornstein BT, Barker HA. The Nutrition of Clostridium kluyveri. J Bacteriol. 1948;55: 223–230. doi:10.1007/978-0-387-68233-4

96. Parte AC. LPSN - List of prokaryotic names with standing in nomenclature (Bacterio.net), 20 years on. International Journal of Systematic and Evolutionary Microbiology. 2018. pp. 1825–1829. doi:10.1099/ijsem.0.002786

97. Zaremba-Niedzwiedzka K, Caceres EF, Saw JH, Bäckström D, Juzokaite L, Vancaester E, et al. Asgard archaea illuminate the origin of eukaryotic cellular complexity. Nat 2017 5417637. 2017;541: 353–358. doi:10.1038/nature21031

98. Mukherjee S, Stamatis D, Bertsch J, Ovchinnikova G, Verezemska O, Isbandi M, et al. Genomes OnLine Database (GOLD) v.6: Data updates and feature enhancements. Nucleic Acids Res. 2017;45: D446–D456. doi:10.1093/nar/gkw992

99. Pagel M. Inferring the historical patterns of biological evolution. Nature. 1999;401: 877–884. doi:10.1038/44766

100. Brown CT, Hug LA, Thomas BC, Sharon I, Castelle CJ, Singh A, et al. Unusual biology across a group comprising more than 15% of domain Bacteria. Nature. 2015;523: 208–211. doi:10.1038/nature14486

101. Castelle CJ, Banfield JF. Major New Microbial Groups Expand Diversity and Alter our Understanding of the Tree of Life. Cell. 2018;172: 1181–1197. doi:10.1016/j.cell.2018.02.016

102. Justice SS, Hunstad DA, Cegelski L, Hultgren SJ. Morphological plasticity as a bacterial survival strategy. Nat Rev Microbiol. 2008;6: 162–168. doi:10.1038/nrmicro1820

103. Pagel M, Meade A. Bayesian analysis of correlated evolution of discrete characters by reversible-jump Markov chain Monte Carlo. Am Nat. 2006;167: 808–825. doi:10.1086/503444

104. Rambaut A, Drummond AJ. Tracer v1.6. 2013.

105. Venditti C, Meade A, Pagel M. Multiple routes to mammalian diversity. Nature. 2011;479: 393–396. doi:10.1038/nature10516

106. Baker J, Meade A, Pagel M, Venditti C. Adaptive evolution toward larger size in mammals. Proc Natl Acad Sci. 2015;112: 5093–5098. doi:10.1073/pnas.1419823112

107. Freckleton, Harvey, Pagel. Phylogenetic Analysis and Comparative Data: A Test and Review of Evidence. Am Nat. 2017;160: 712. doi:10.2307/3078855

108. Organ CL, Shedlock AM, Meade A, Pagel M, Edwards S V. Origin of avian genome size and structure in non-avian dinosaurs. Nature. 2007;446: 180–184. doi:10.1038/nature05621

109. Pagel M, Meade A, Barker D. Bayesian Estimation of Ancestral Character States on Phylogenies. Syst Biol. 2004;53: 673–684. doi:10.1080/10635150490522232

110. Soucy SM, Huang J, Gogarten JP. Horizontal gene transfer: building the web of life. Nat Rev Genet. 2015;16: 472.

111. Wiedenbeck J, Cohan FM. Origins of bacterial diversity through horizontal genetic transfer and adaptation to new ecological niches. FEMS Microbiol Rev. 2011;35: 957–976. doi:10.1111/j.1574-6976.2011.00292.x

112. Caro-Quintero A, Konstantinidis KT. Inter-phylum HGT has shaped the metabolism of many mesophilic and anaerobic bacteria. ISME J. 2015;9: 958–967. doi:10.1038/ismej.2014.193

113. Zeng Y, Feng F, Medová H, Dean J, Koblí Zek M, Blankenship RE. Functional type 2 photosynthetic reaction centers found in the rare bacterial phylum Gemmatimonadetes. doi:10.1073/pnas.1400295111

114. Greenhill SJ, Currie TE, Gray RD. Does horizontal transmission invalidate cultural phylogenies? Proc R Soc B Biol Sci. 2009;276: 2299–2306. doi:10.1098/rspb.2008.1944

115. Currie TE, Greenhill SJ, MacE R. Is horizontal transmission really a problem for phylogenetic comparative methods? A simulation study using continuous cultural traits. Philos Trans R Soc B Biol Sci. 2010;365: 3903–3912. doi:10.1098/rstb.2010.0014

