## Supplementary information for "Phenotypic reconstruction of the last universal common ancestor reveals a complex cell"

**This PDF file includes:**

Supplementary Text

Figs. S1 to S3.

Tables S1 to S6 and S6.1

References

**Other Supplementary Materials for this manuscript include the following:**

The Bacterial and Archaeal Phenotypic Database (BAPdb) can be accessed at: https://doi.org/10.6084/m9.figshare.12987509.v1

**Supplementary text.**

1. Data collection: avoiding sampling bias

While an excellent resource, due to an understandable focus on disease-causing organisms in general Bergey’s manual may reproduce this bias. To avoid potential over-representation of host-associated species, we selected those for our analyses based on their presence on the phylogeny of Chai et al. ^1^. (the ‘large’ tree) and the Bergey’s manuals ^2–6^ (i.e. their intersection) used as a source of data for these species. Data for species described after the publication of the Bergey’s manuals were collected from the primary literature based on the List of Prokaryotic Names with Standing in Nomenclature ^7^ (LPSN). Data were not reported for taxa without a species description (e.g., *Bacillus sp*. 1NLA3E, *Streptococcus sp*. GMD2S). To account for phylogenetic uncertainty for deep branching taxa we ran the same analyses on the independently reconstructed phylogeny of Segata et al. ^8^ (the ‘small’ tree) using the data collected for the large tree.

2. Detailed description of phenotypic traits

*2.1 Cell morphology*

We defined three broad categories of shapes, rod, ovoid and coccoid (Materials and Methods). Any cell described with words such as “coccoid”, “coccus”, “regular or irregular cocci” “spheroid” and “round cells” was coded as coccoid. Cells described as a mycelium that fragments into coccoid or coccoid-like elements were not considered as coccoid. Species described with words and phrases such as “rods”, “rod-like”, “bacillus”, “rods with rounded ends”, “straight rods”, and “curved rods”, “slightly curved rods”, “rods with tapered ends”, “straight to slightly curved rods”, “cylindrical cells”, “oval rods”, and “elliptical rods” or “coccoid rods” were reported in the “Rod” shape category. Species described as “coccobacillus”, were also considered as rods. The curvature and the end type of the cells were not considered (e.g. straight or curved rods, and rounded or tapered ends). Species usually described as vibroid, filamentous, helical or branched were also considered as rods (see criteria below) as their underlying form includes elongated individual cells. Vibroid cells were those described as “vibrio”, “coryneform”, “diphteroidal” and also “S-shaped”, “C-shaped”,” sausage shape”, “ring-like or horseshoe-shaped”, “drumstick” or “trapezoid in shape”. Helical cells were defined as described by, “helical”, “spiral”, “spirillum”, “spirochete”, “corkscrew”, “tightly coiled helical rods”, “rods, with a regular helical coiling” and “coiled or helical filaments”. Descriptions like “spiral bodies” or “spiral chains” were not recorded as helical cells as it was revealed that these bodies are formed by self-aggregation of flagella ^9^ while spiral chains refer to the shape of aggregating but not individual cells. Filamentous cells were defined as those described as “filamentous”, “threadlike”, “threadlike rod”, “threadlike bacilli”, “cylindrical filaments”, “twisted filaments”, “trichomes” or “Trichomes displaying false branching”. Cells described as “very long cells” were not considered as filamentous if this was due to change in cultural conditions. Branched species were described as “branched”, “branched diphtheroids”, “branched filaments”, “mycelium”, “hyphae”, “and spiral aerial hyphae” or “actinomycete”. Finally, “Ovoid” encompasses any cell described as “ovoid”, “ovococcoid”, “ovococcus”, “elliptical”, “teardrop-shaped”, “oval”, “fusiform”, “drop-like shape”, “eye-shaped”, or “tuber-shaped”.

Species displaying more than one morphology among the three (rod, ovoid and coccoid) were coded as having both morphologies (e.g. terms such as ‘coccoid or rod’ were coded as both coccoid and rod). However, if additional morphologies were recorded they were not considered if they were described with adverbs of frequency for scarcity such as “sometimes”, “occasionally”, “rarely”, “seldom” or “sometimes”, etc. (e.g. cells are straight rods, sometimes coccoid cells occur). Additional morphologies of a given species were not considered if they were described with adjectives, words and sentences such as “few”, “some cells”, “in specific growth conditions”, “may” or “may be”, “can be”, “in the stationary phase of growth”, “occur in old cultures”, “X forms are rare”, “in ageing cultures”, “eventually”, “may or may not”.

To assess whether the delineation of coccoid from non-coccoid species based on our qualitative categorization of shape was consistent with cell size measurement, we calculated the aspect ratio of the cell (AR, as length divided by width, Fig. S1.). For species where minimum and maximum length and/or width values were available, an average length and width was calculated as their sum divided by two. If multiple cell size values were reported due to changes in culture conditions, for old cultures, or during the stationary phase of growth, these were not considered. More generally, additional cell size values described with adverbs of frequency such as “sometimes”, “occasionally” or “rarely” were not considered.

*Pleomorphism*

Any species displaying a single morphology among the three shape categories was described as monomorphic, unless alteration of shape was indicated (e.g. “rods tend to pleomorphism”). Species displaying two or three morphologies or described with words and sentences such as “pleomorphic”, “variable”, “morphologically variable”, “cells highly irregular in shape” or “different or various shapes” were described as non-monomorphic.

*Motility*

Motility was defined for the purpose of this study as the ability to move regardless of the method of locomotion (i.e. flagella, archaella, pilli or fimbriae). Therefore, species described as motile with or without a description of the motility appendage were considered as motile. Species described as being motile with adverbs of frequency such as “sometimes”, “occasionally”, “rarely”, “seldom”, or with “few”, “maybe” or “can be” were considered as motile according to our definition. Species for which motility information was extracted from the genus description were considered as missing (NA) for motility and excluded from the analysis if the genus description contained sentences such as “motile or non-motile species” as motility will vary between species in the genus. Species exhibiting changes in motility status dependent on growth conditions were recorded as being motile. Species described as having flagella but were not motile or without a motility description were recorded as missing for motility (three species) and were excluded from the motility analysis.

*Cell aggregation*

Species occurring only as single cells were coded as non-aggregating, while those occurring as single cells but with the ability to associate were coded as aggregating.

*Habitat*

Information on habitat (i.e. the different locations where the organism naturally lives and grows and from which it could be recovered and isolated) were extracted from the species description section in the Bergey’s manuals and the primary literature. The Freshwater category included all species described as living or isolated from freshwater sediments, from rivers or lakes, wells, groundwater, flooded rice-fields, or terrestrial hot-springs, etc. The Marine category included any species isolated from or living in the sea, ocean or coastal waters, hydrothermal vents or marine sediments. The species categorized under Marine or Freshwater were coded together as aquatic in our secondary habitat listing (habitat b, Materials and Methods). Species isolated from or living in soil, woodland, or isolated from root nodules, plants or any terrestrial host were considered as Terrestrial. Species found in more than one habitat and described with sentences like “isolated from a variety of thermal habitats, both terrestrial and aquatic” were recorded as both Terrestrial and Aquatic.

*Cell envelope*

Species were classified as having a mondermic or didermic cell plan regardless of the cell wall structure. In addition to the description in the Materials and Methods species described with sentences such “Gram stain negative but have a gram-positive cell wall structure”, “typical cell-wall structure of Gram-positive bacteria without an outer membrane but Gram-negative staining”, or “Gram-variable but have a Gram positive cell wall”, “Gram-stain variable, but has a Gram-positive ultrastructure”, or “Gram positive, but staining properties can be lost with age” were all considered as monodermic. Species for which the Gram-stain was described with words and sentences such as “variable”, “negative or positive”, “variable to negative”, “variable to positive”, or “Gram-negative and Gram-variable”, and “negative, but staining is irregular” without additional information were excluded from the analysis.

*Spore formation*

Species with description of the spore shape and/or location or position were recorded as spore forming (e.g. “coccoid spores are formed centrally”). To date, spore formation has not been observed in archaea and we therefore coded all the archaea as non-spore forming.

*Physicochemical parameters*

In addition to optimal temperature, pH and NaCl ranges, qualitative description of the corresponding category was often provided in the species description (e.g. thermophile, neutrophile, halophilic). However, the qualitative description is subjective as there is no strict consensus on how to define these categories. For instance, a species growing in an optimal temperature of 70 ºC might be considered as thermophile or hyperthermophile depending on the authors. We therefore defined our physicochemical parameters categories on a consensus based on lower and upper optimal values.

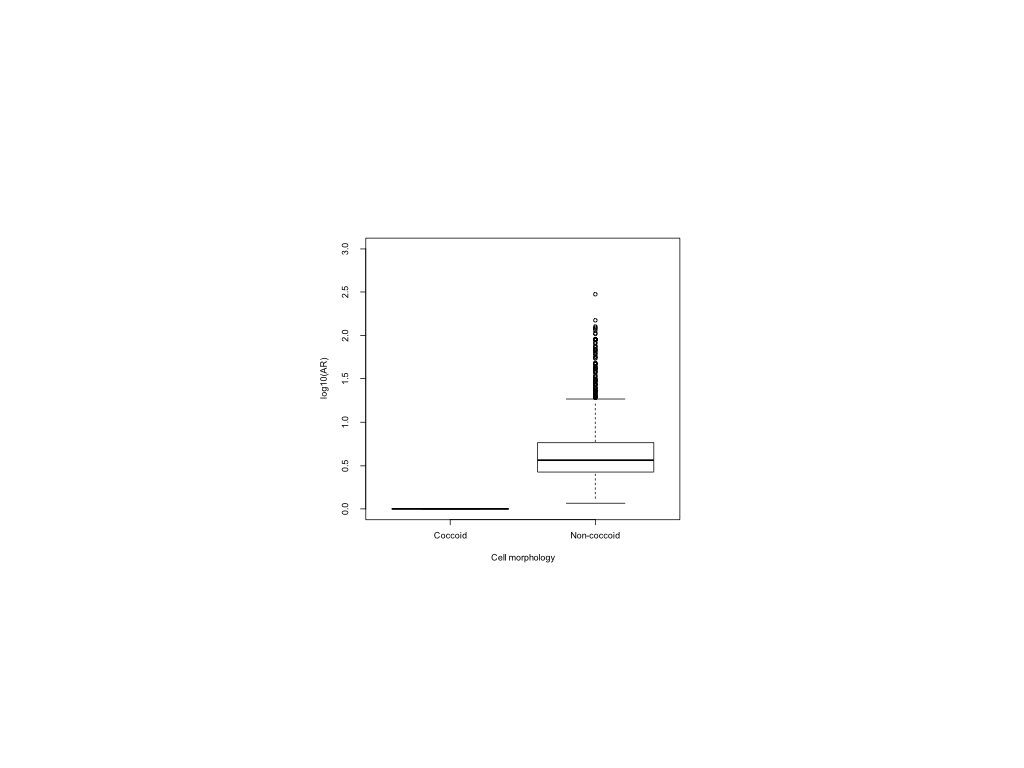

Fig. S1. Delineation between coccoid and non-coccoid species based on Aspect Ratio (AR).

The distribution of log10 AR in coccoid and non-coccoid (rod and ovoid) species. All non-coccoid species have a log10 (AR) > 1. Center line, median; box limits, upper and lower quartiles; whiskers, 1.5× interquartile range; points, outliers.

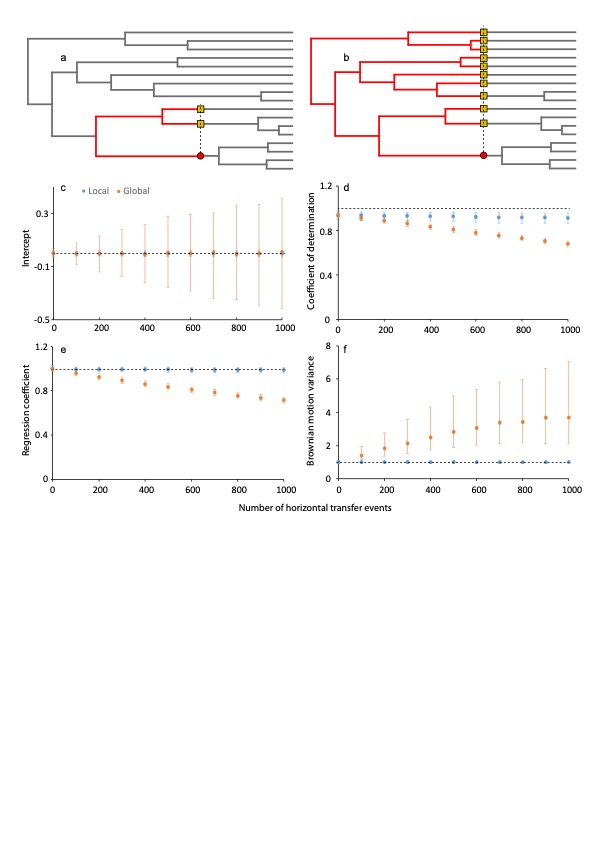

**Fig. S2: Horizontal trait transfer simulations**. (a) An example of how the Local HTT simulations were conducted. The red point is the donor and the yellow squares are the potential recipients. The red branches show the *Local* clade defined by the most recent common ancestor of the HTT event. (b) An example of how the *Global* HTT simulations were conducted. The red point is the donor and the yellow squares are the potential recipients. The red branches show the clade defined by the root (i.e. all the branches in the tree at that point have an opportunity to be a recipient. (c), (d) and (e) show the median (2.5^th^ and 97.5^th^ percentiles) of the intercept, coefficient of determination and the regression coefficient respectively of the 100 simulation for each simulation set. (f) shows the median (2.5^th^ and 97.5^th^ percentiles) of the inferred Brownian motion variance. Orange points are those associated with the global simulation and the blue point the local simulation.

**Fig. S3. Posterior probability histograms for all the characters from Chai et al’s tree.**

Posterior probabilities for all the characters from Chai et al’s tree are shown as histograms for a) LUCA, b) LBCA and c) LACA. Abbreviations: LUCA - last universal common ancestor; LBCA -last bacterial common ancestor; LACA - last archaeal common ancestor;

**a) LUCA**

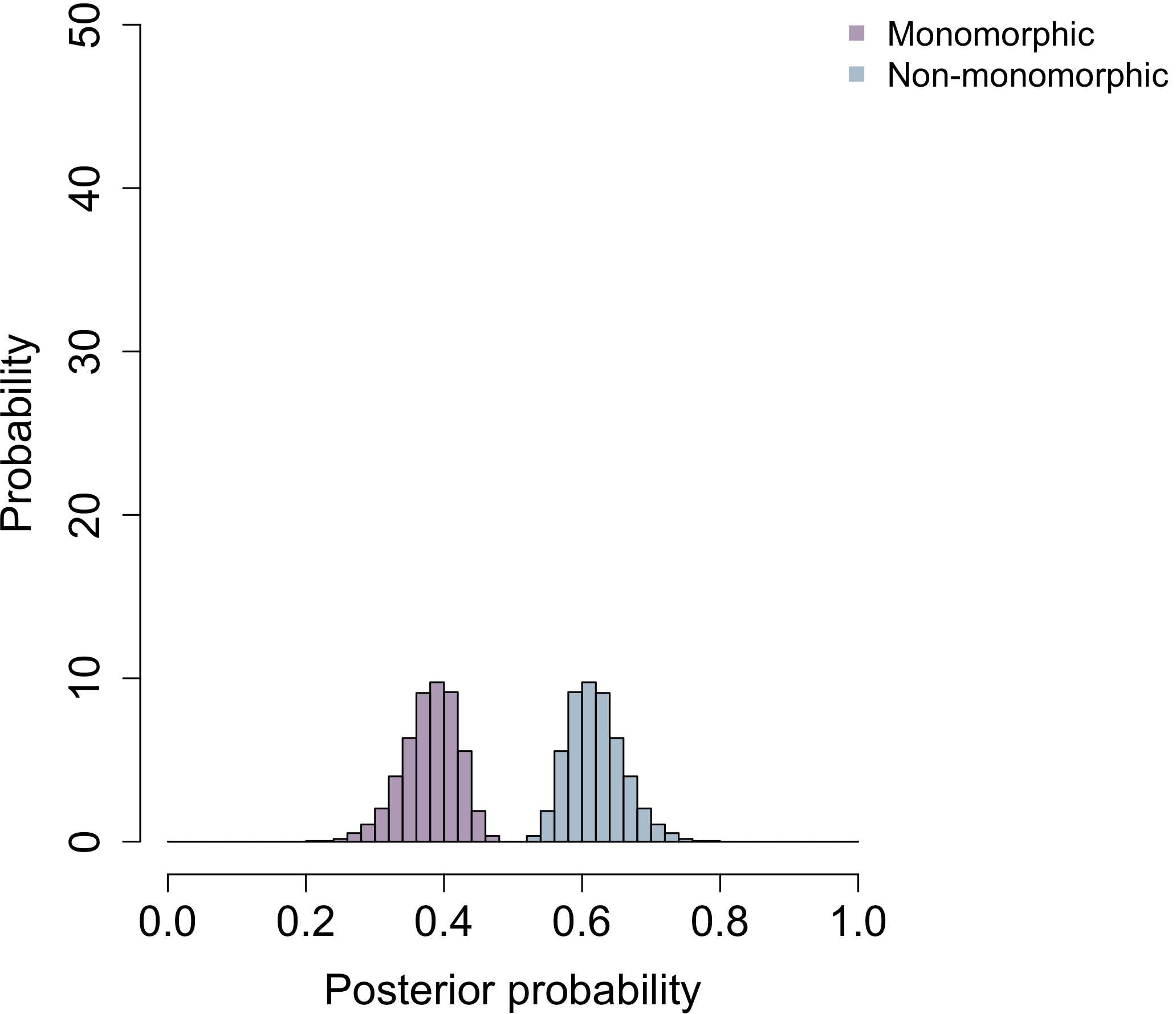

**Pleomorphism**

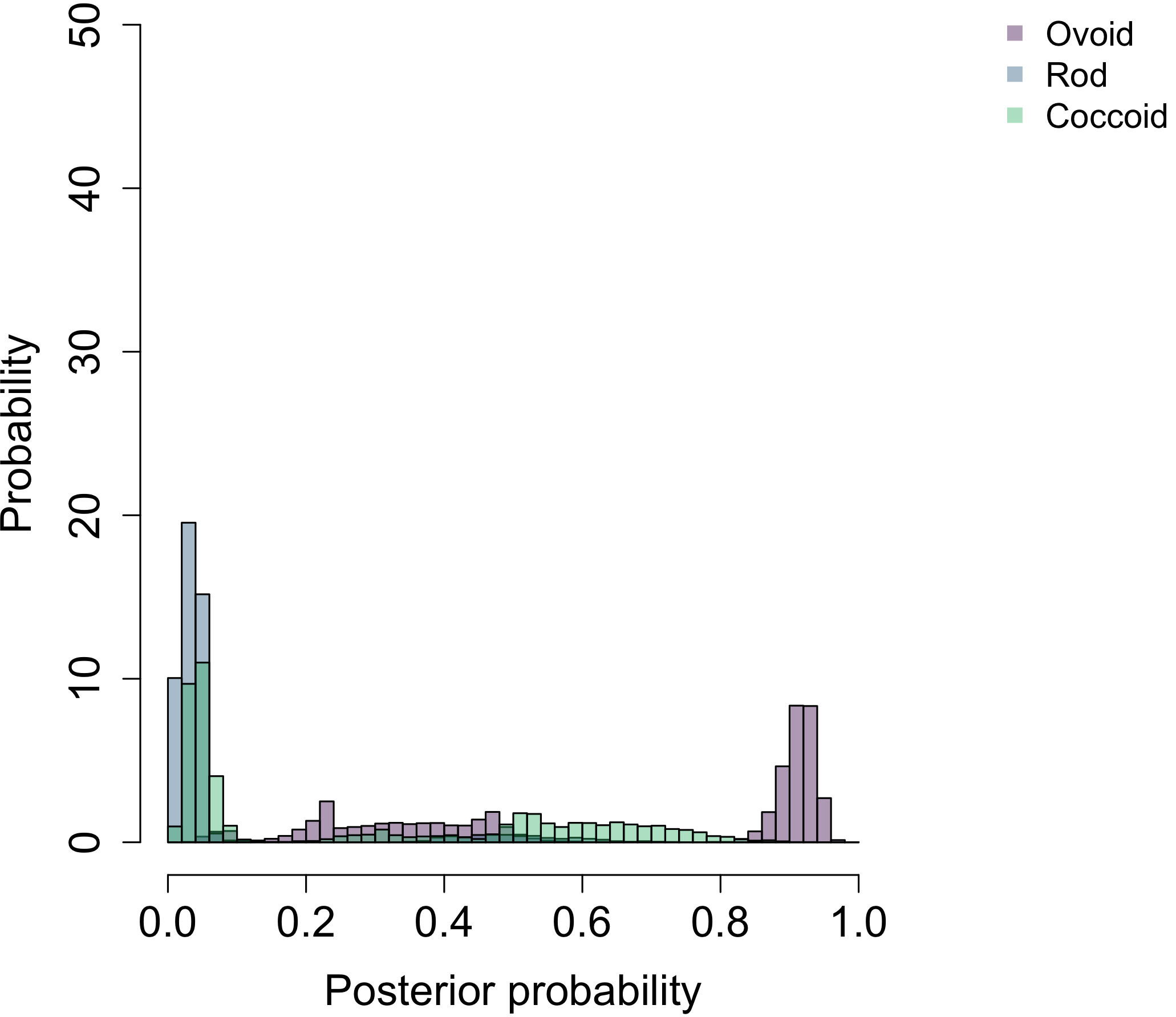

**Shape**

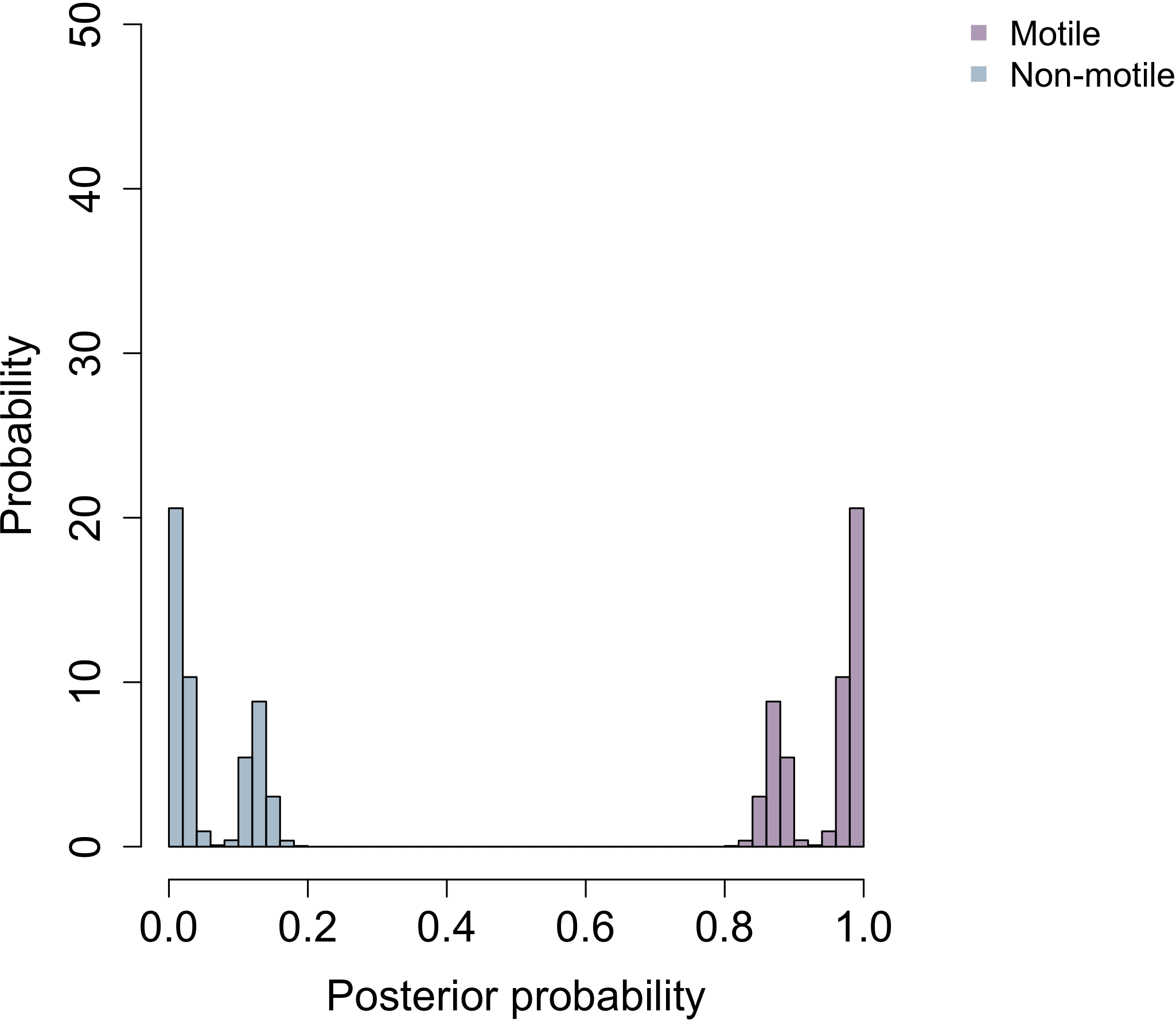

**Motility**

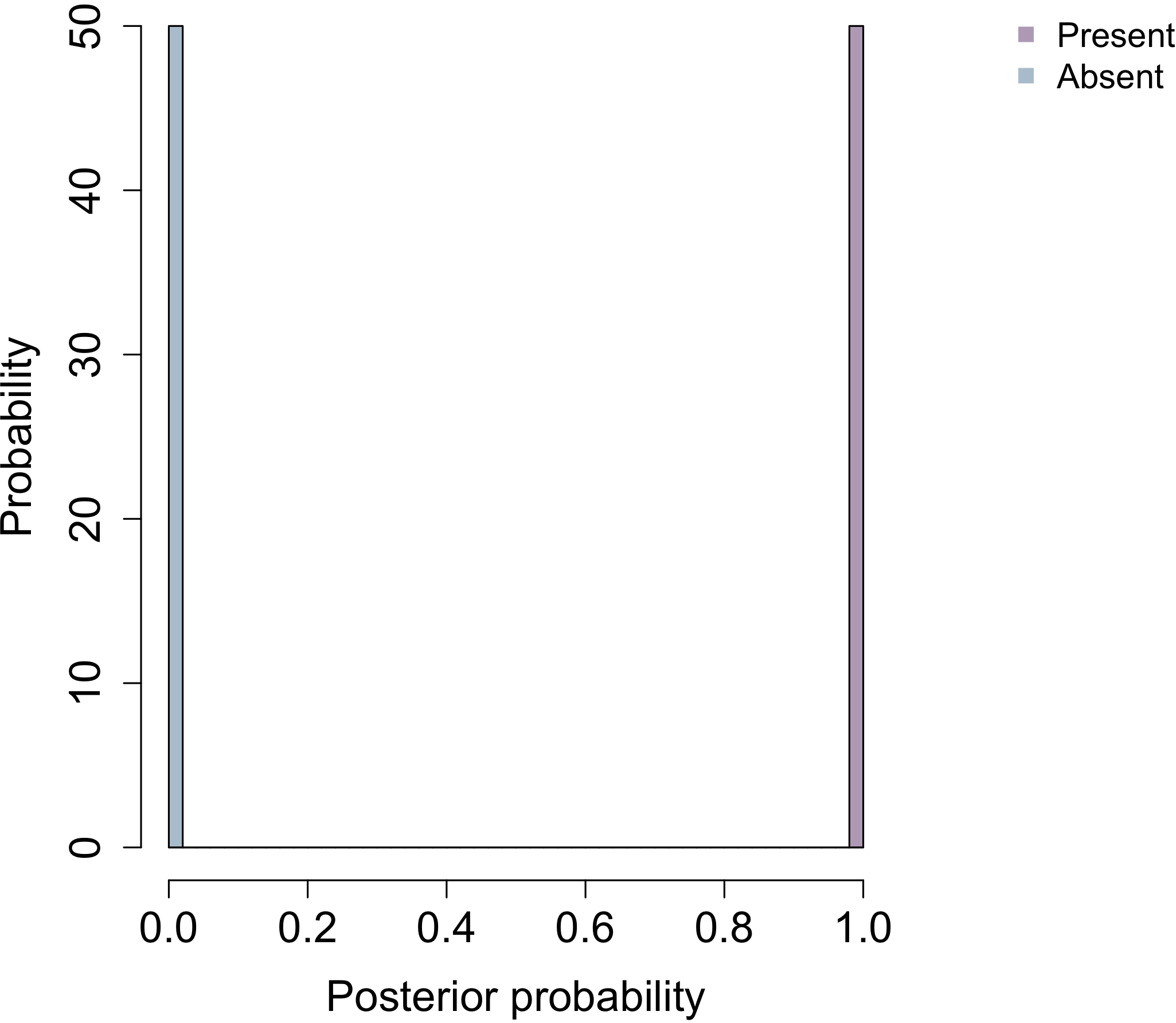

**Cell wall**

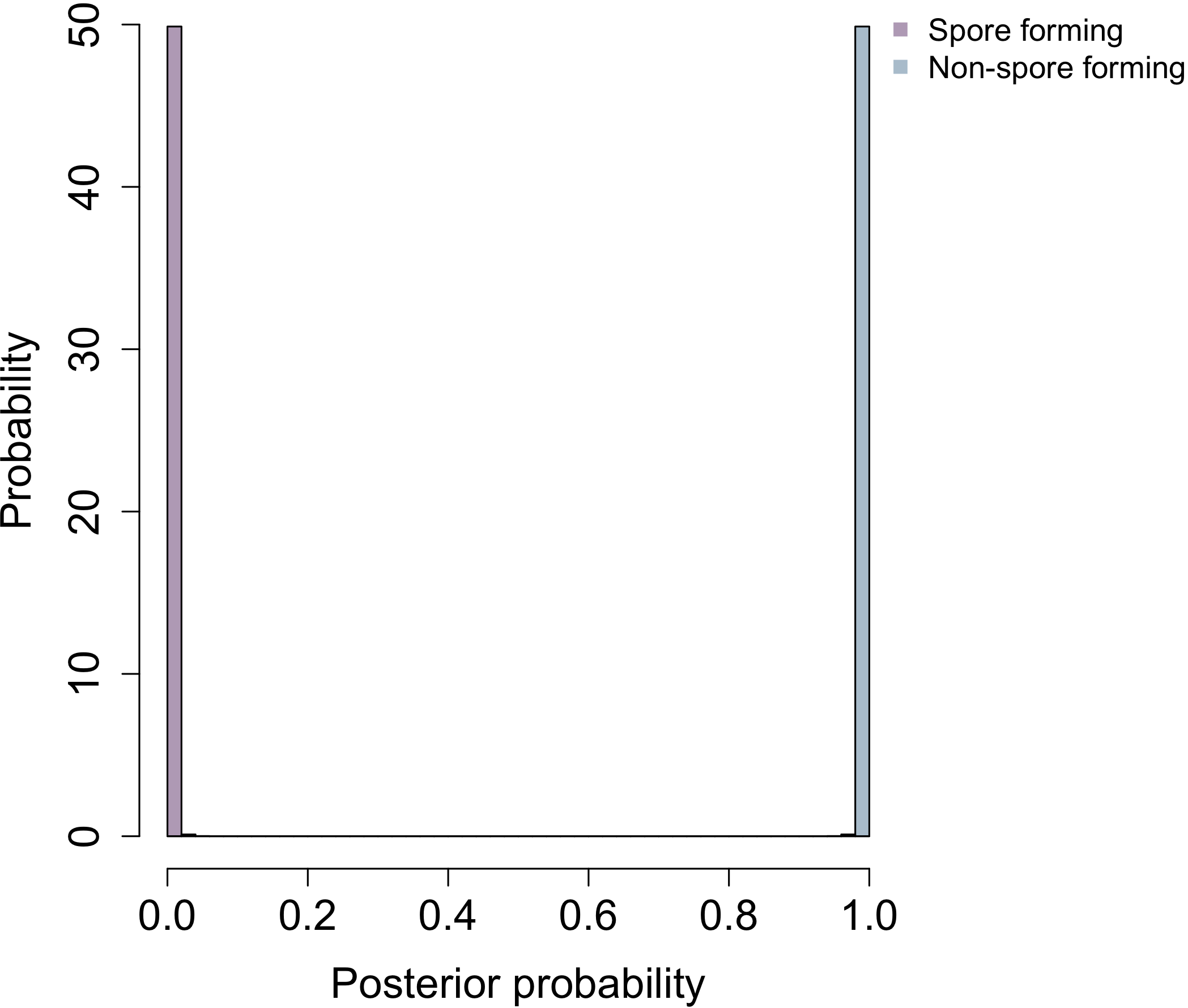

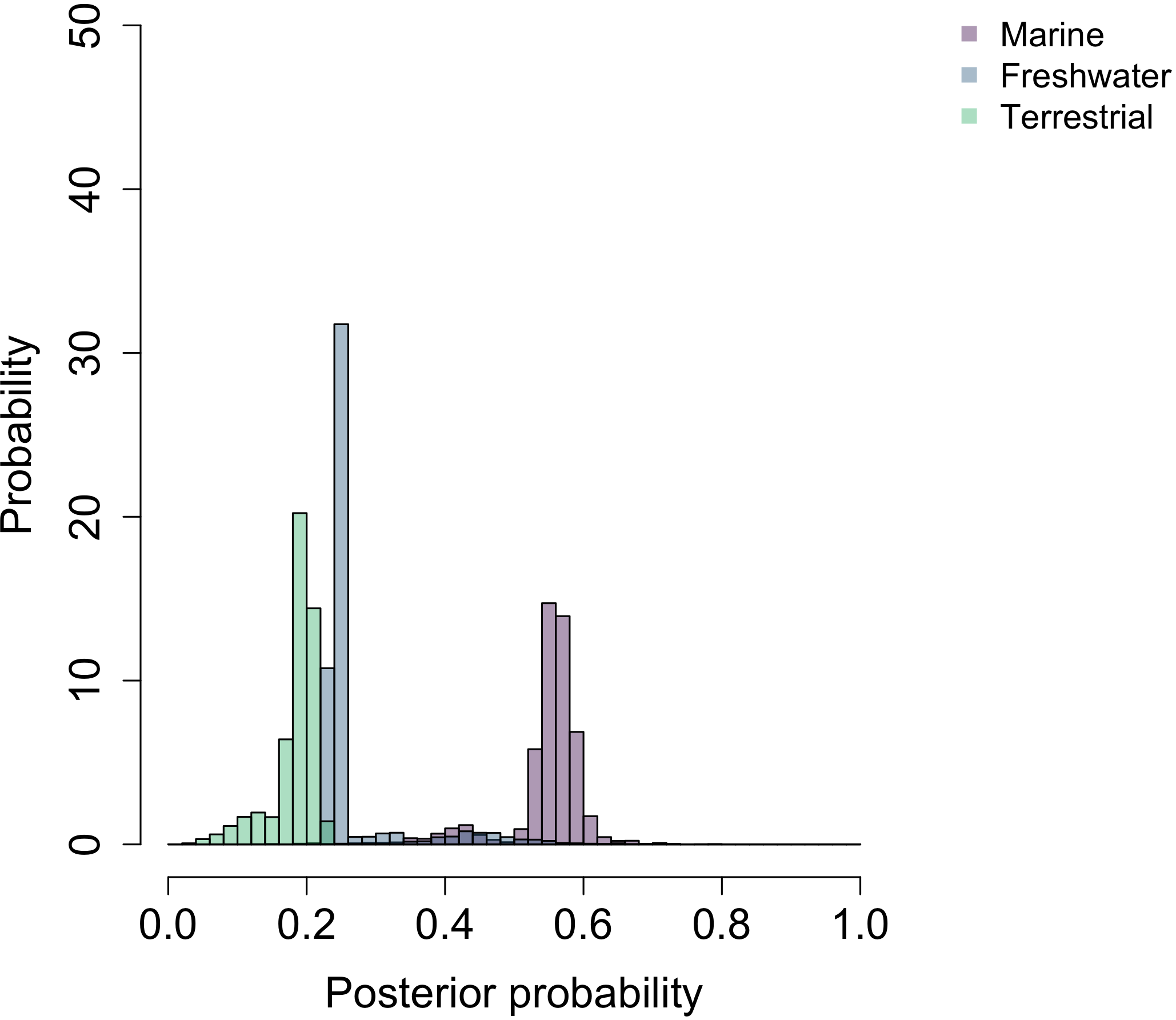

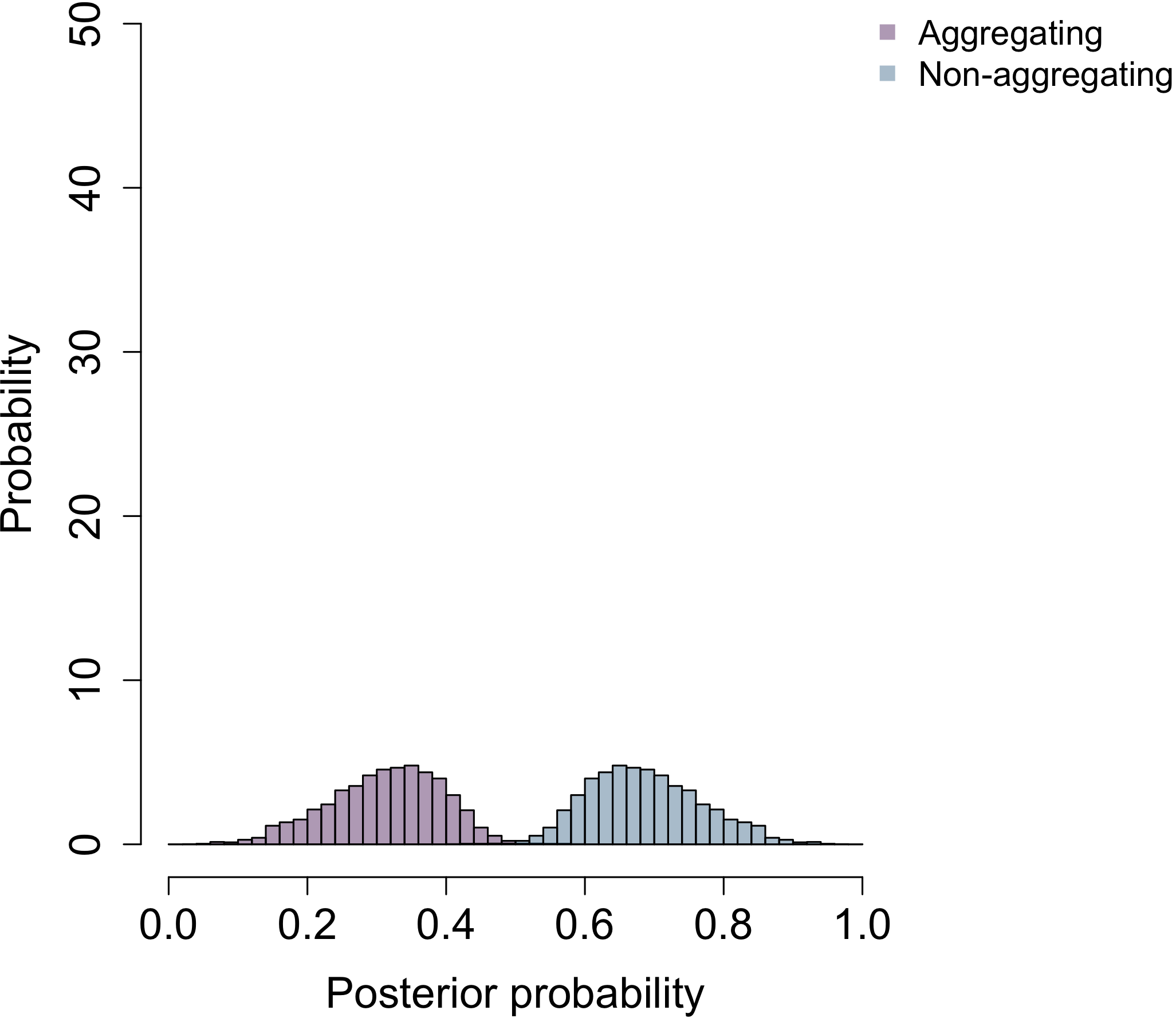

**Cell aggregation**

**Sporulation**

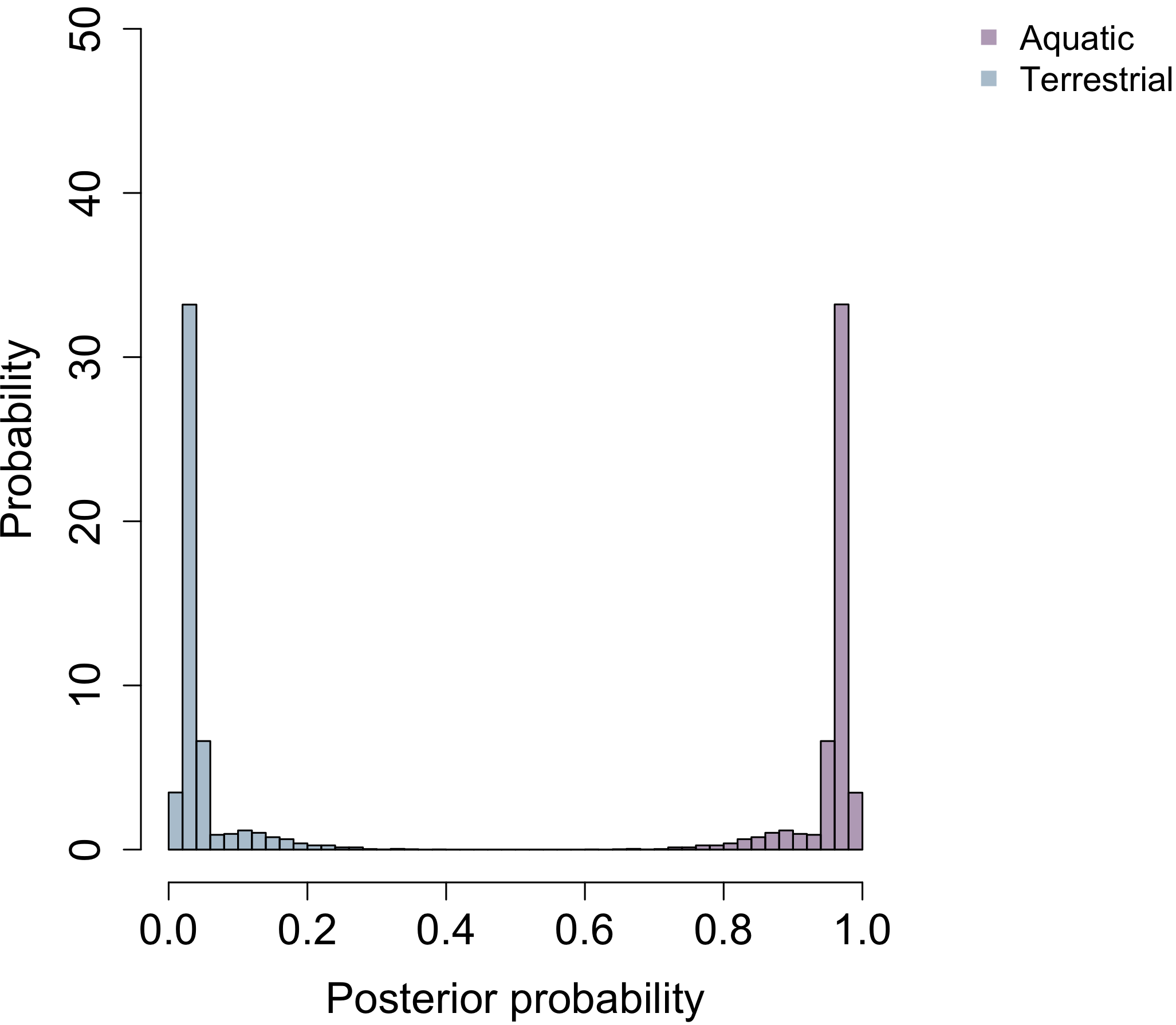

**Habitat b**

**Habitat c**

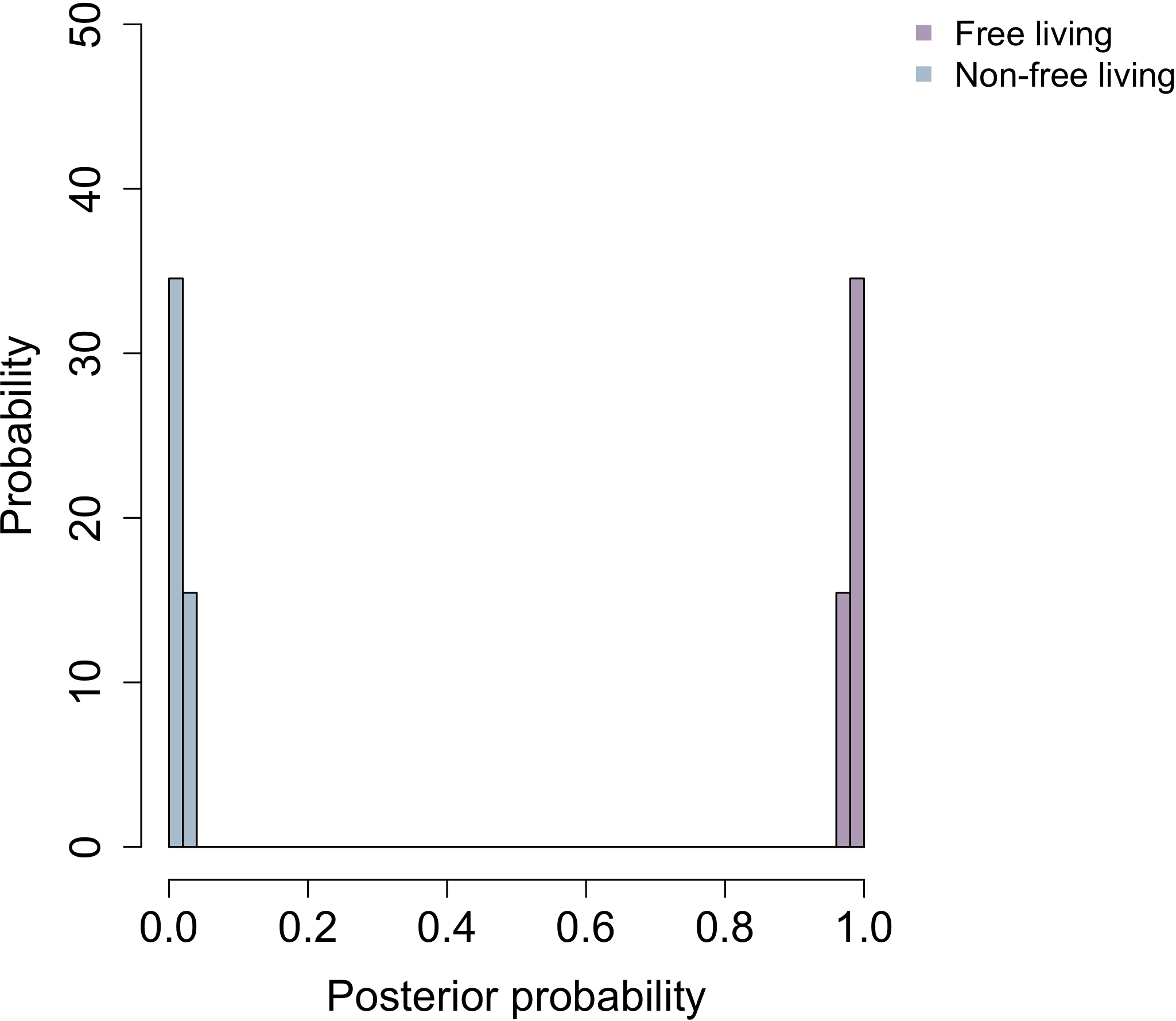

**Habitat a**

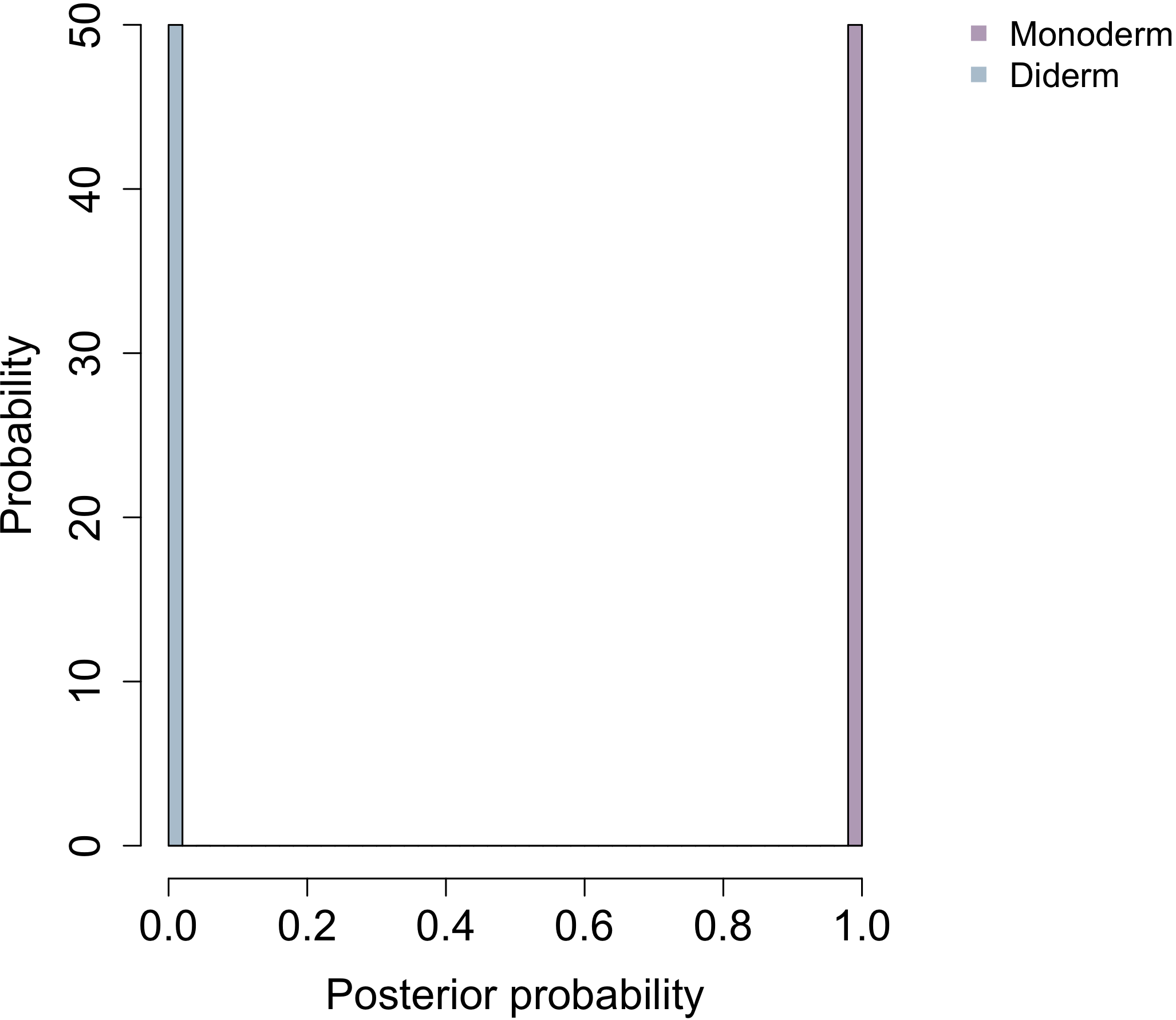

**Cell plan**

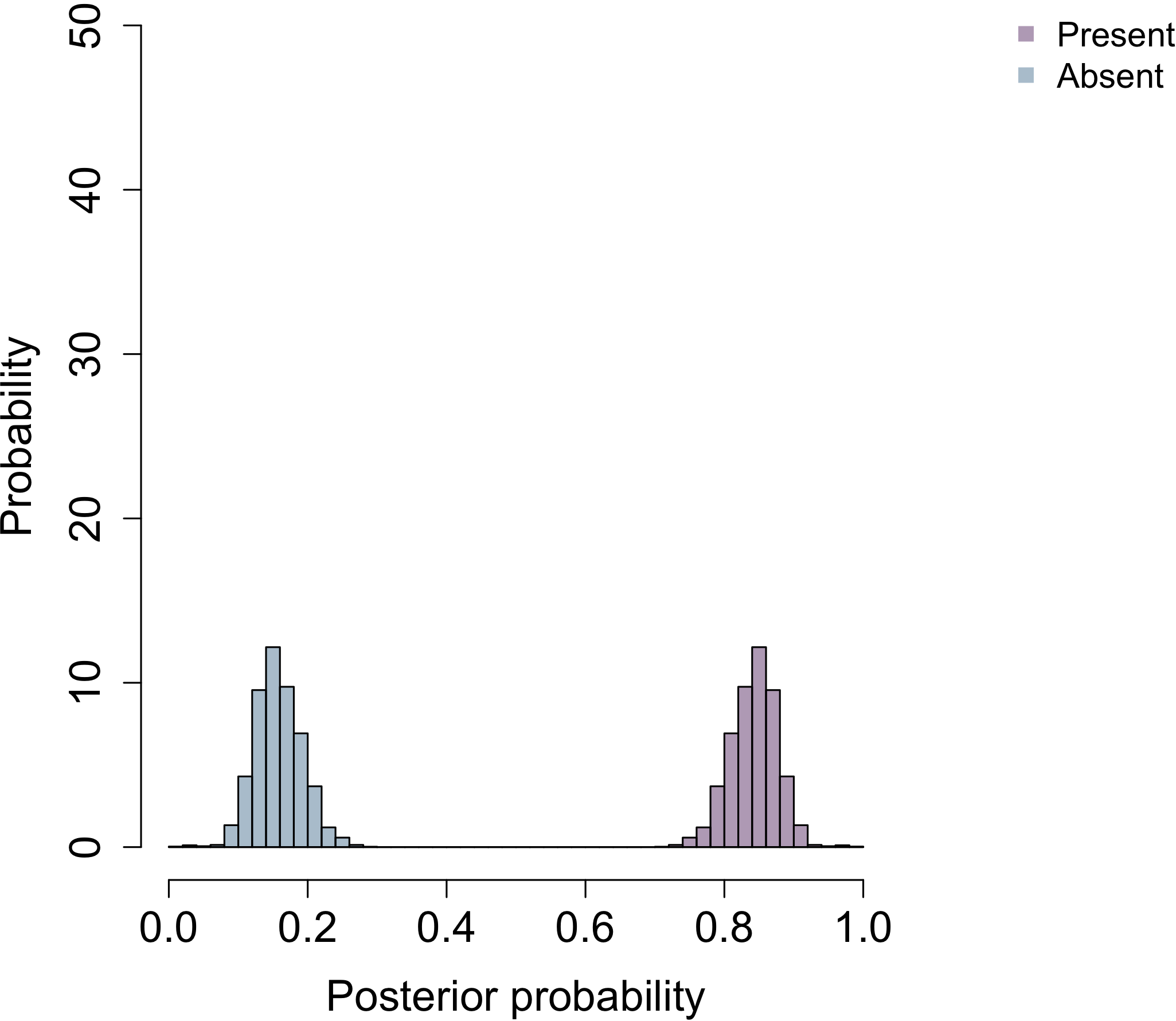

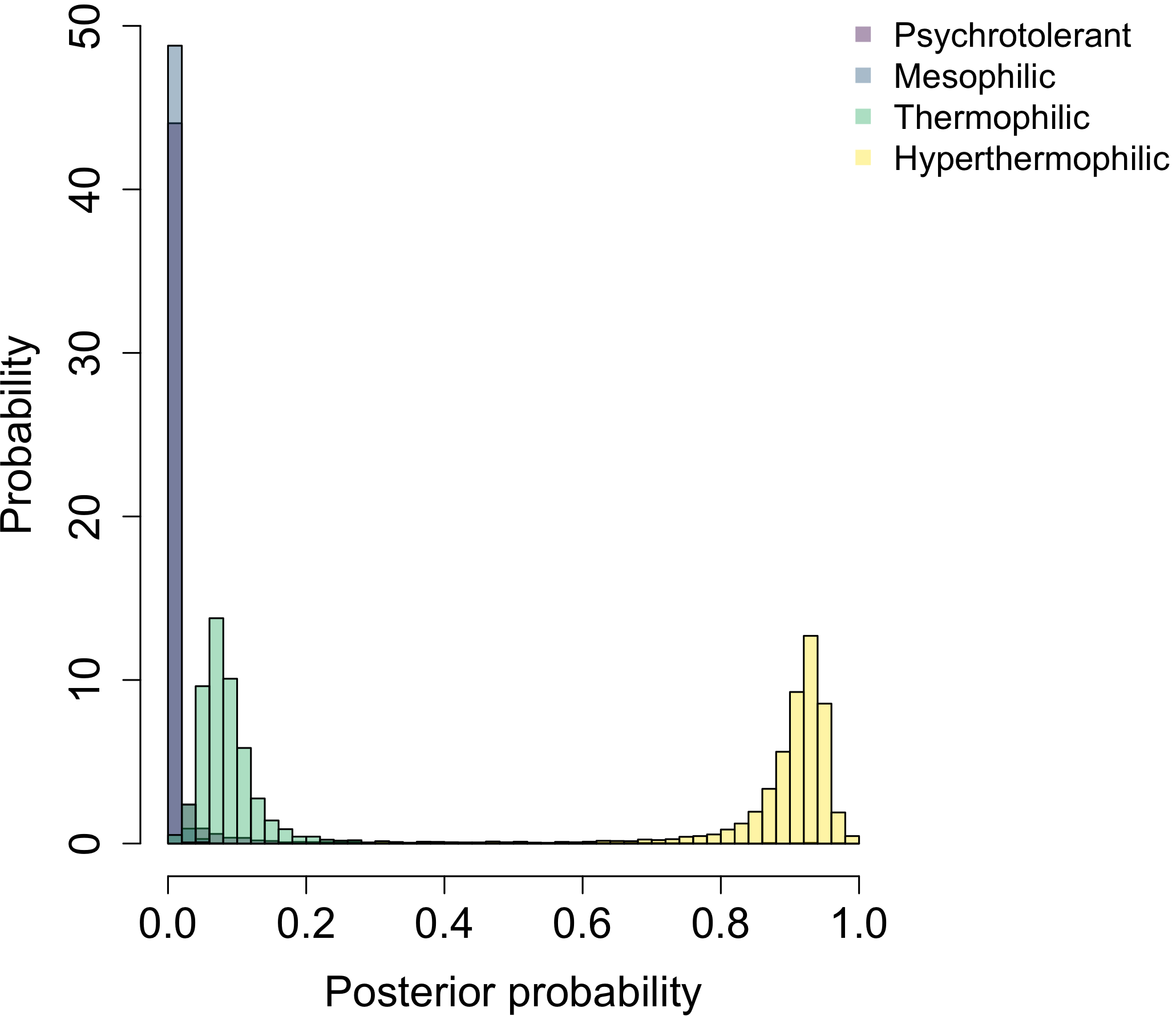

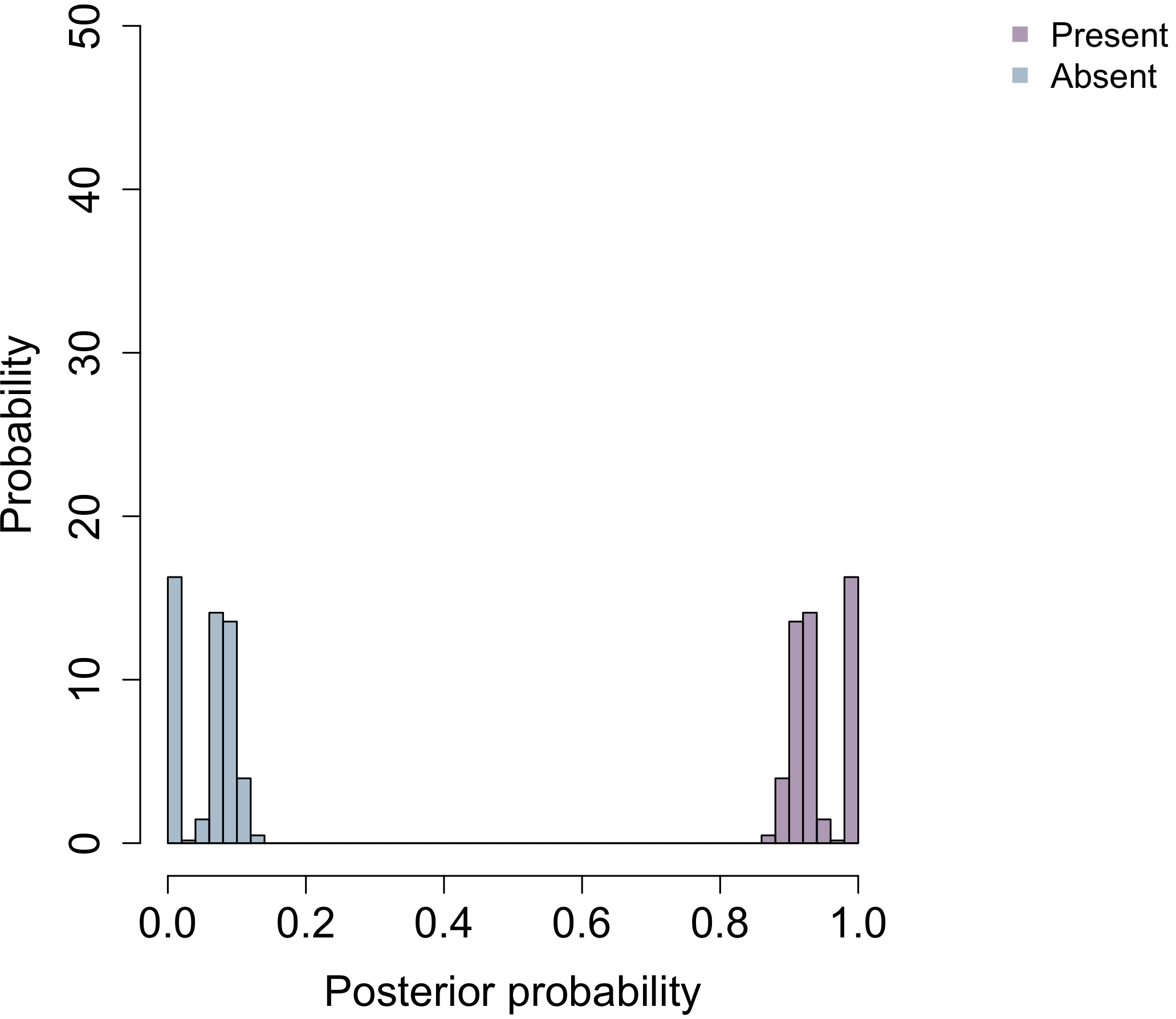

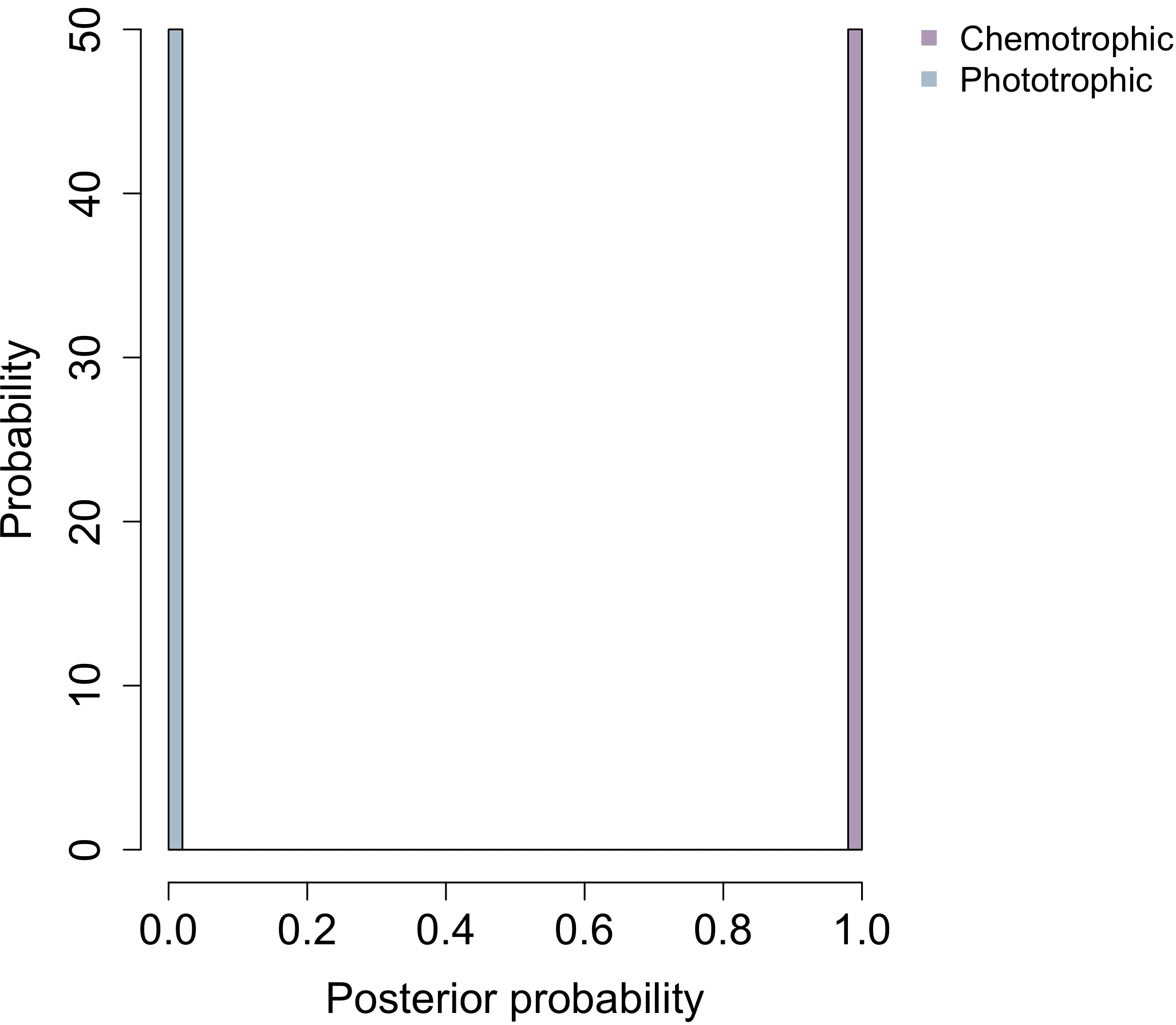

**Energy source**

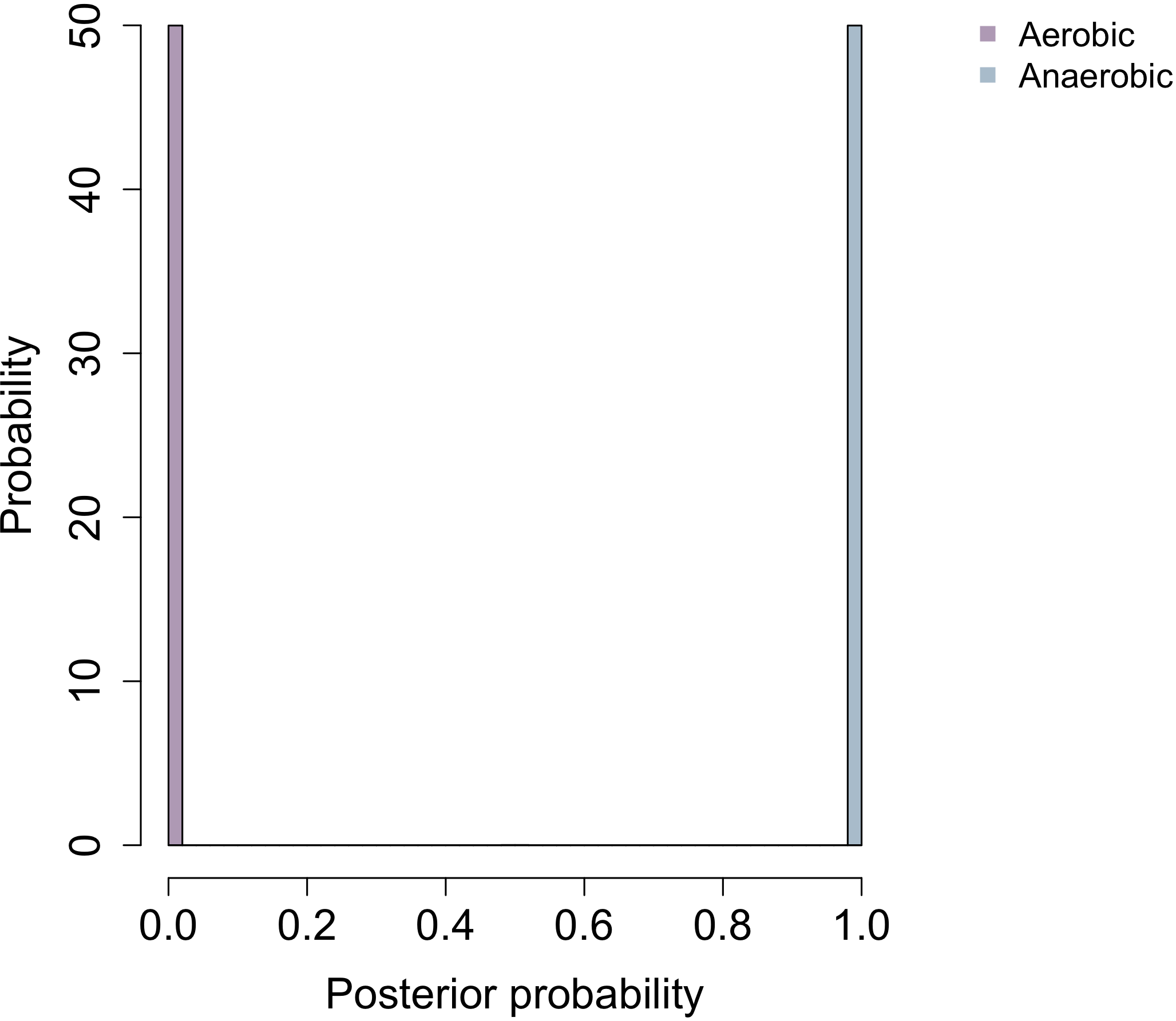

**Oxygen requirement**

**Oxidase**

**Temperature**

**Catalase**

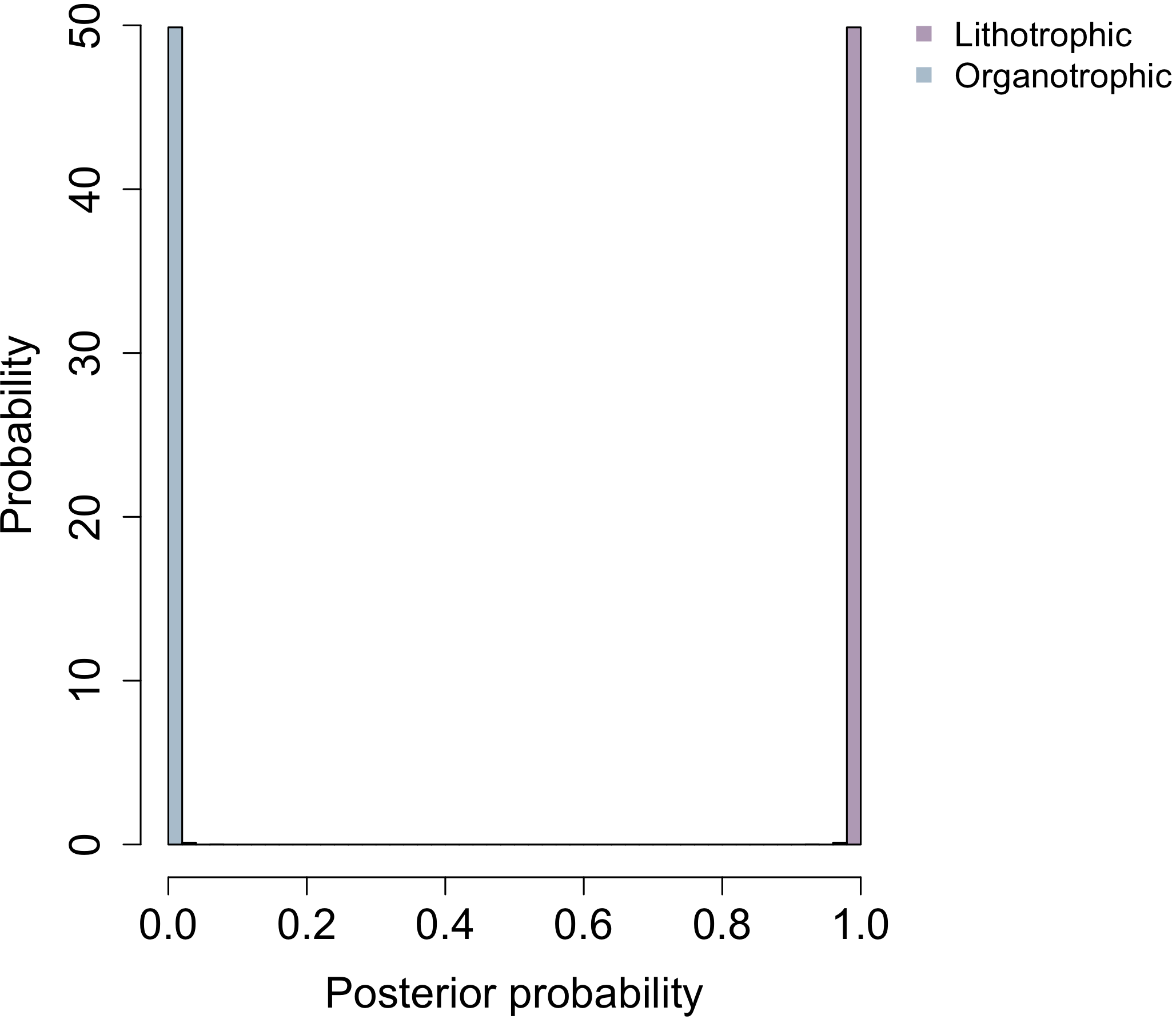

**Electron donor**

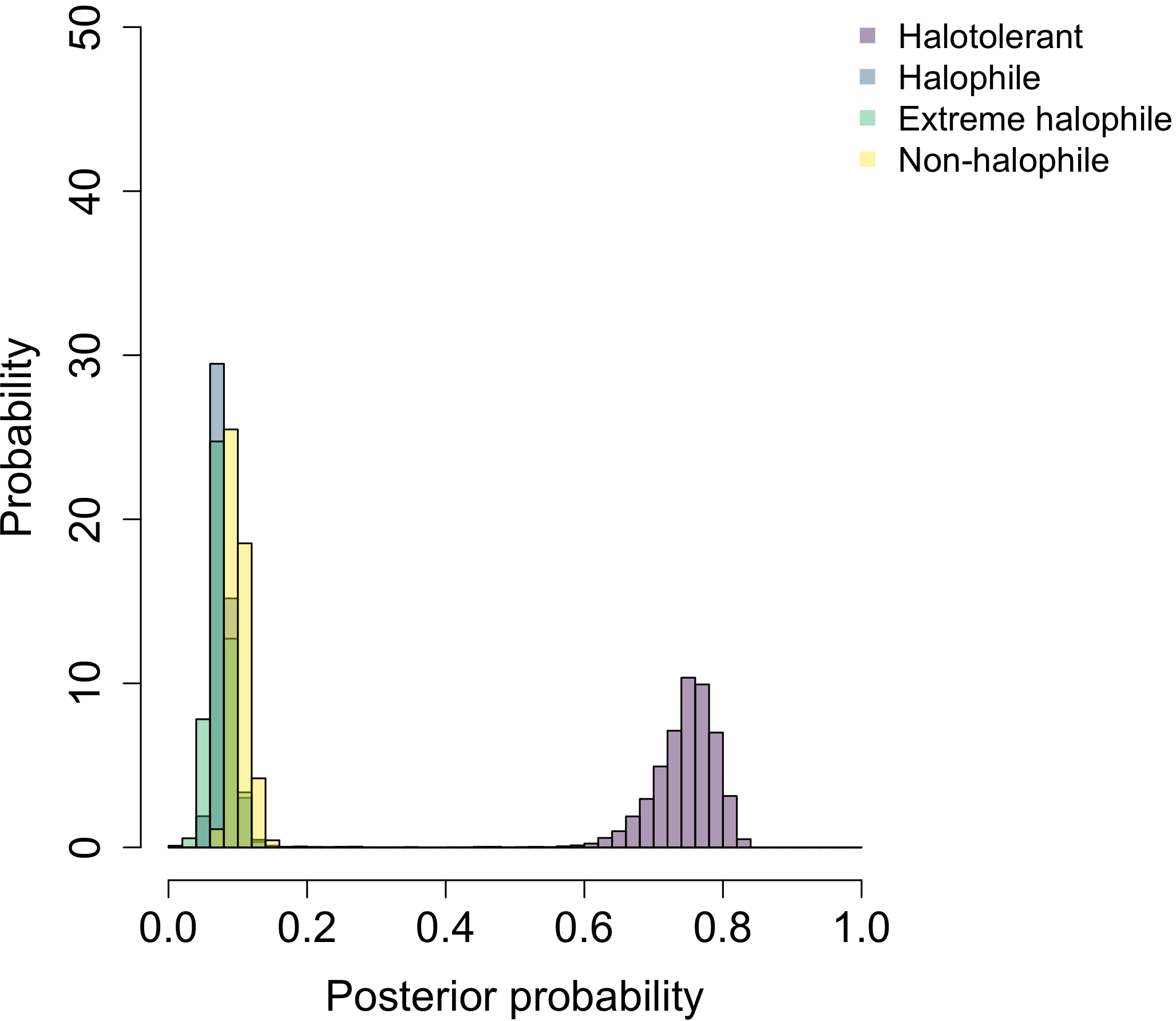

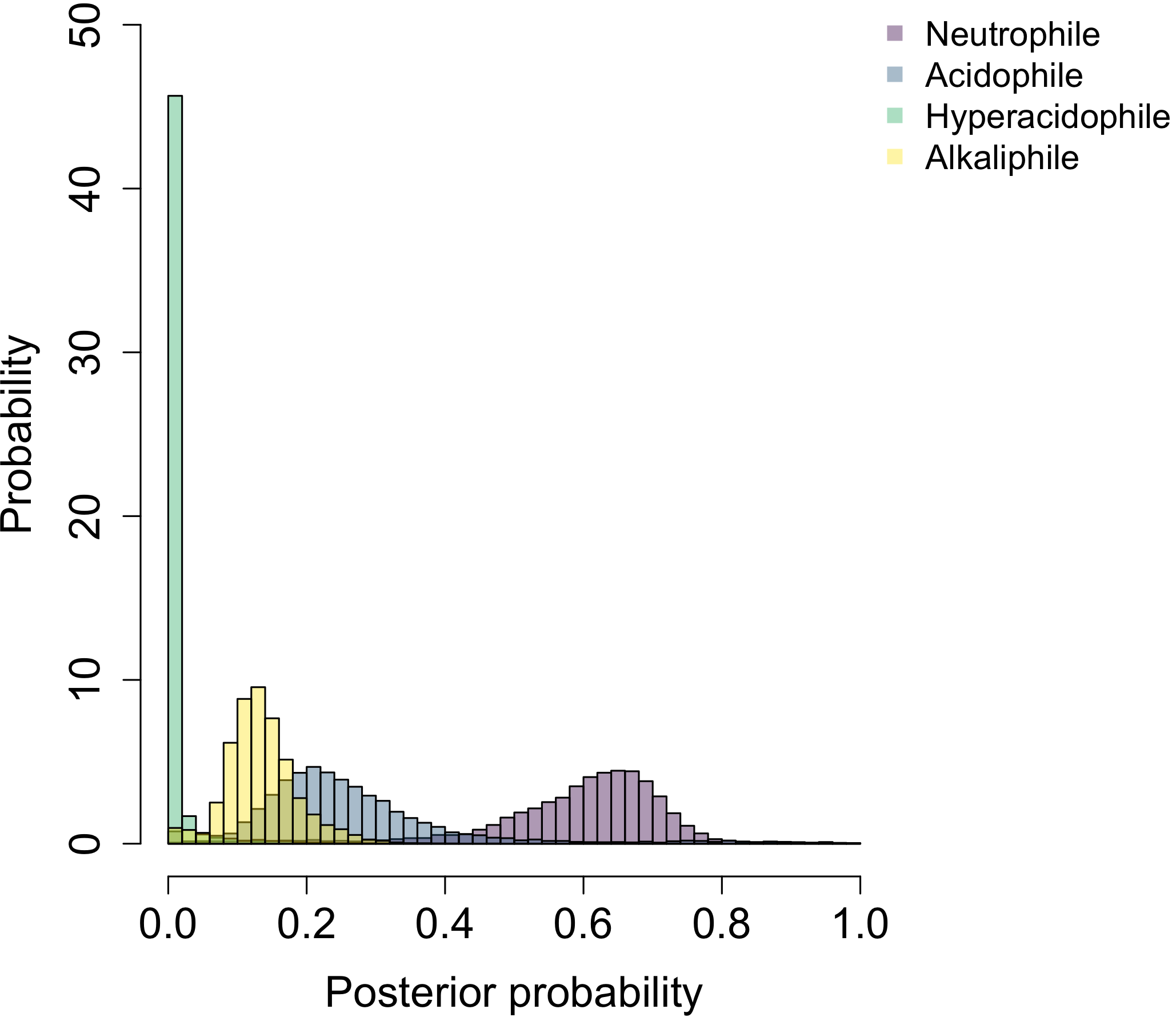

**NaCl**

**pH**

**b) LBCA**

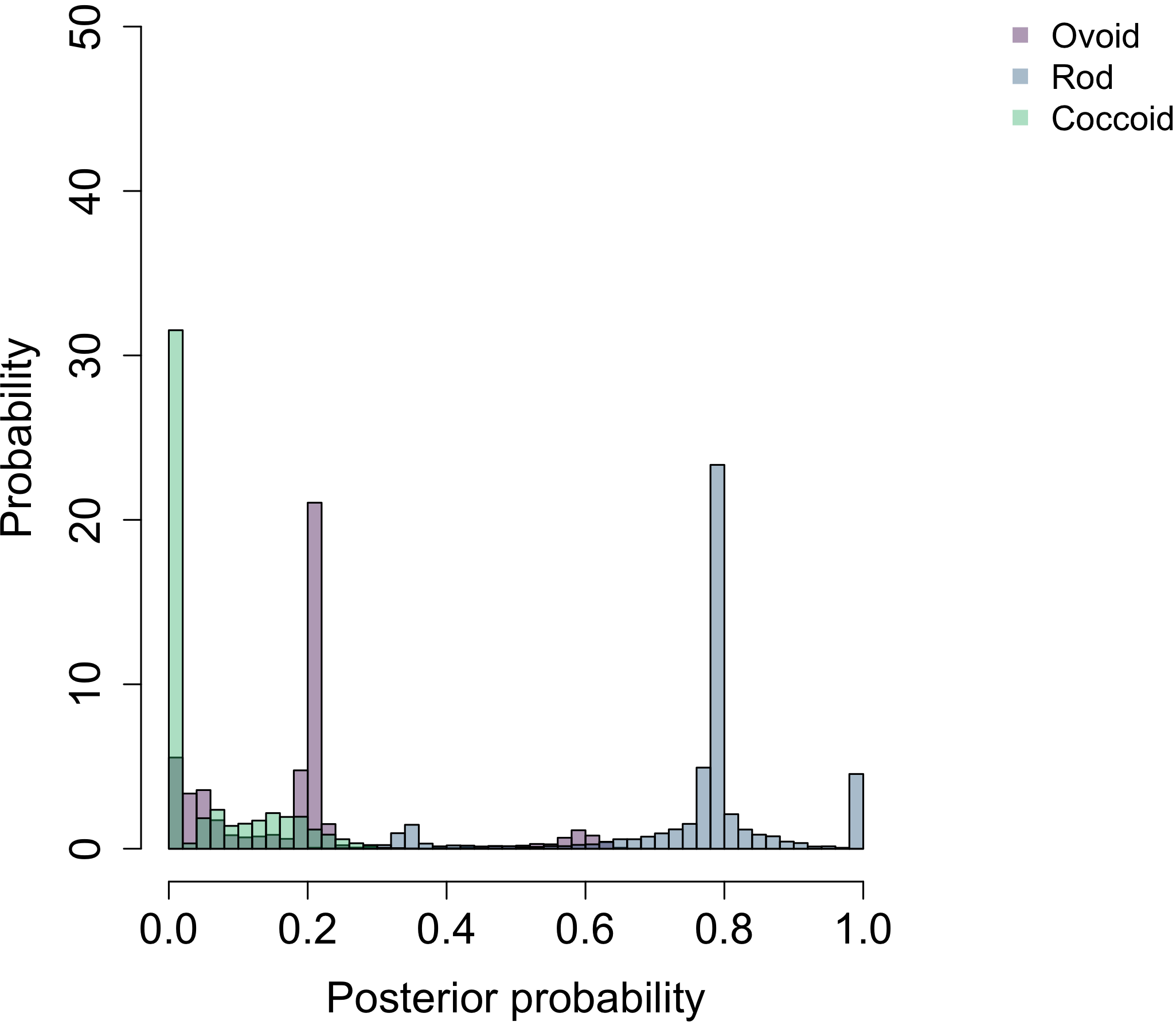

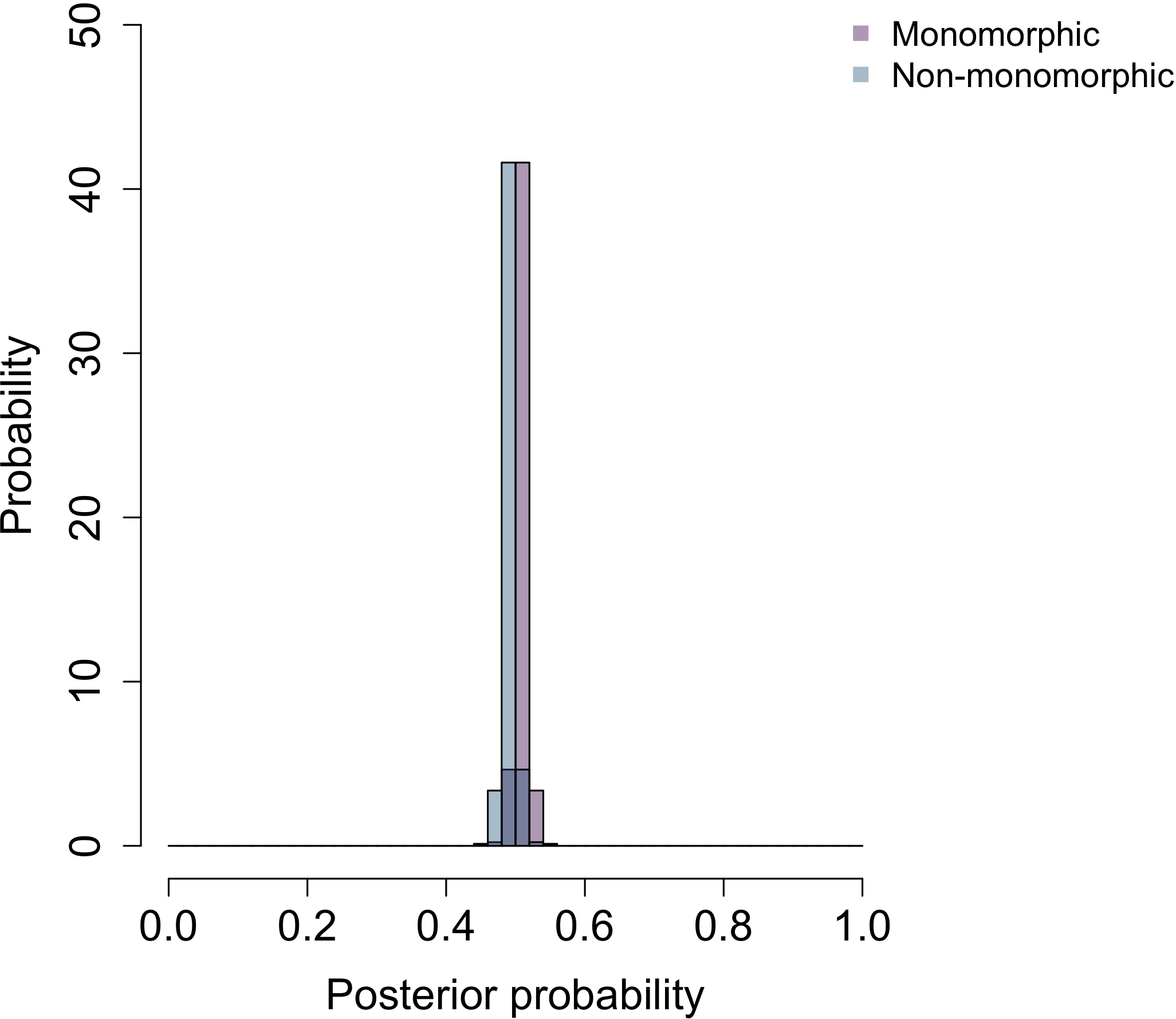

**Pleomorphism**

**Shape**

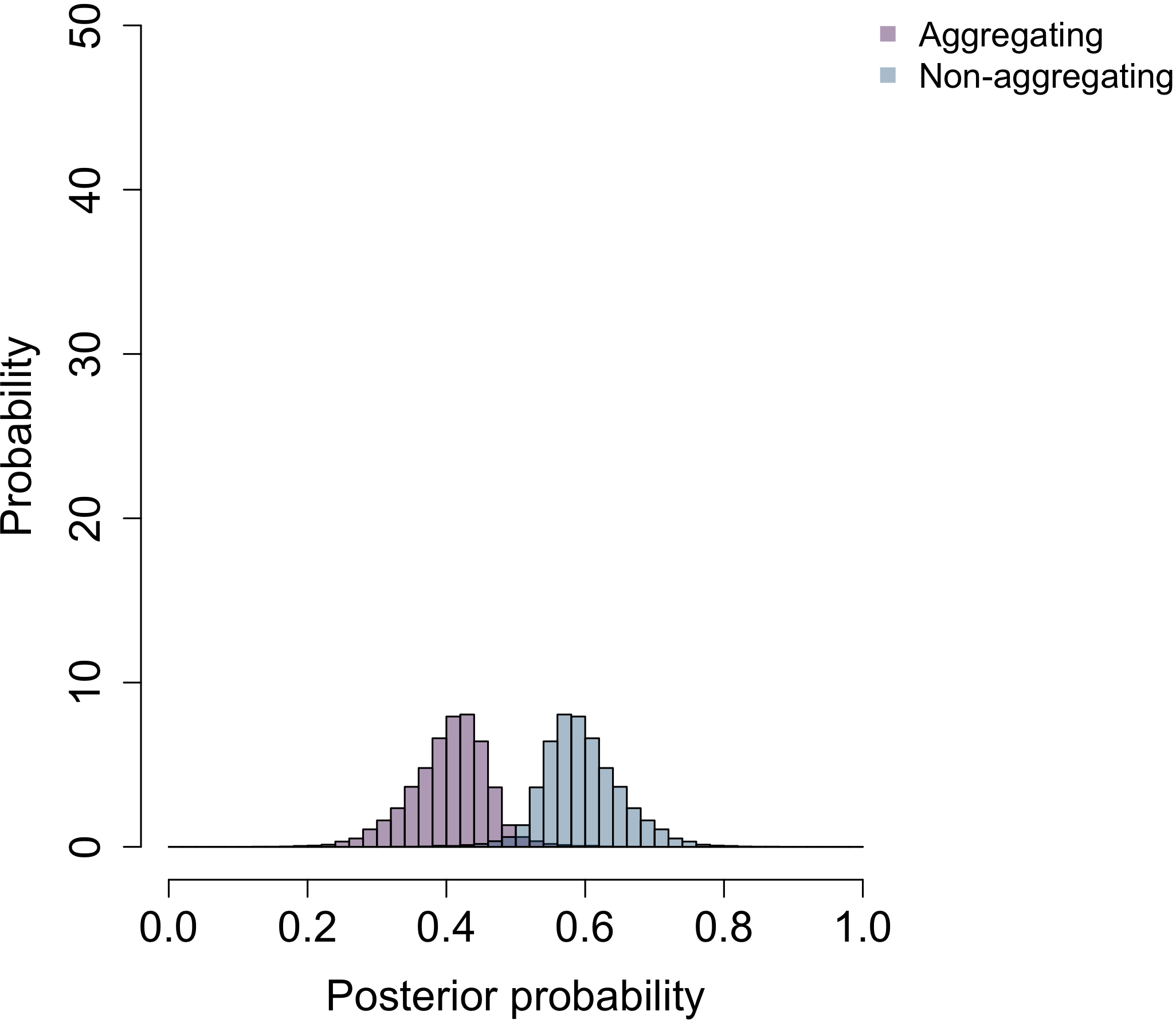

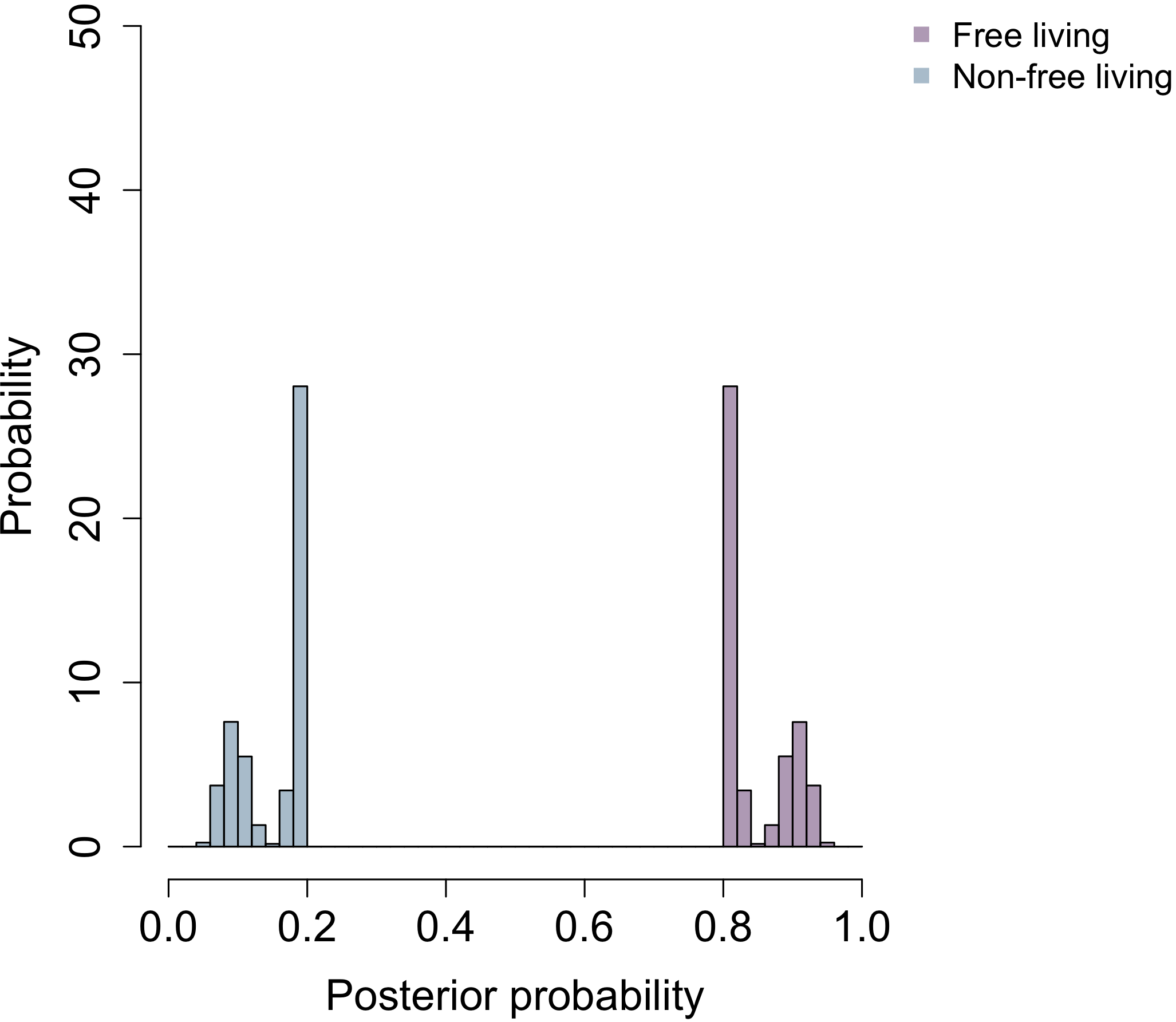

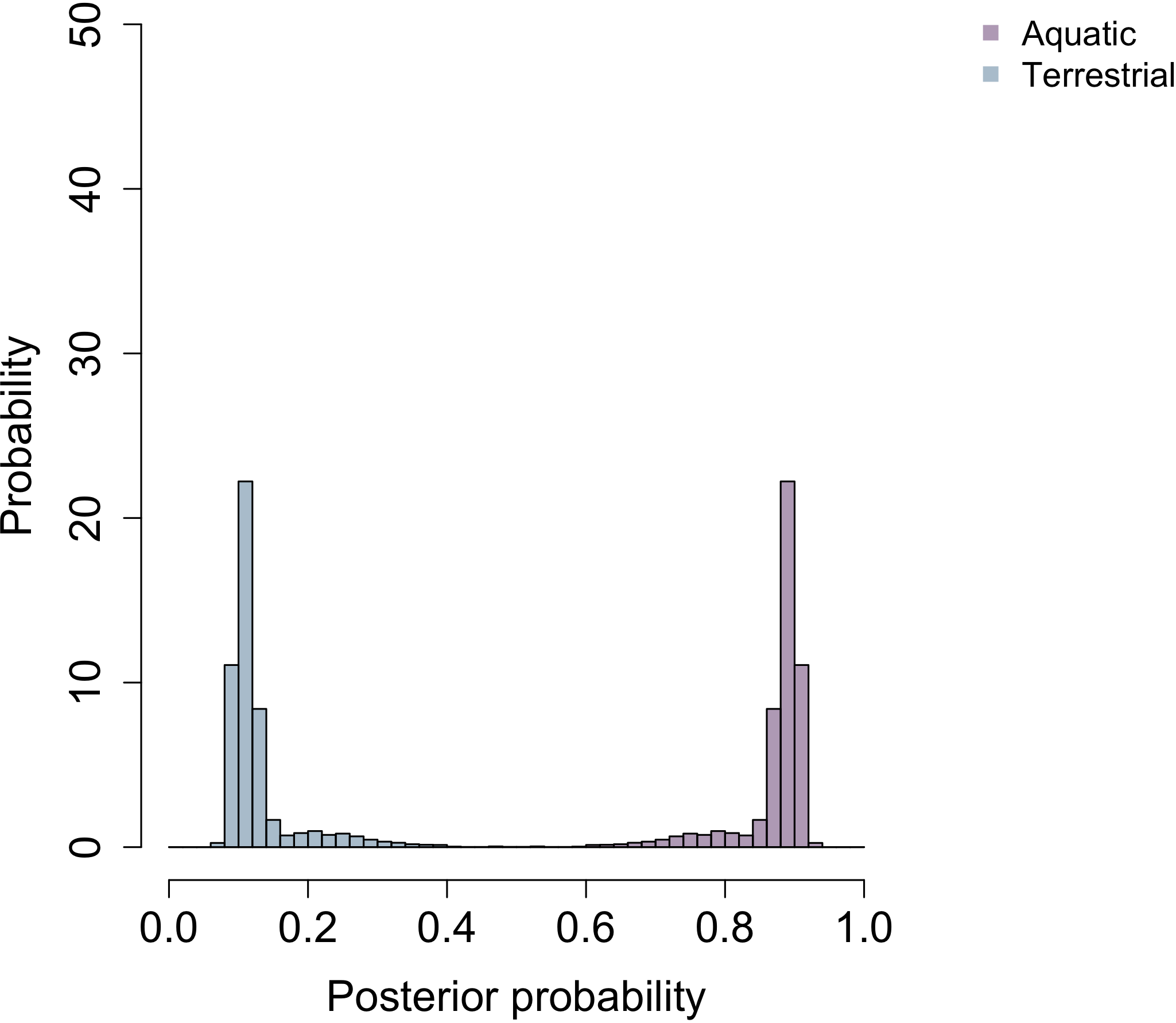

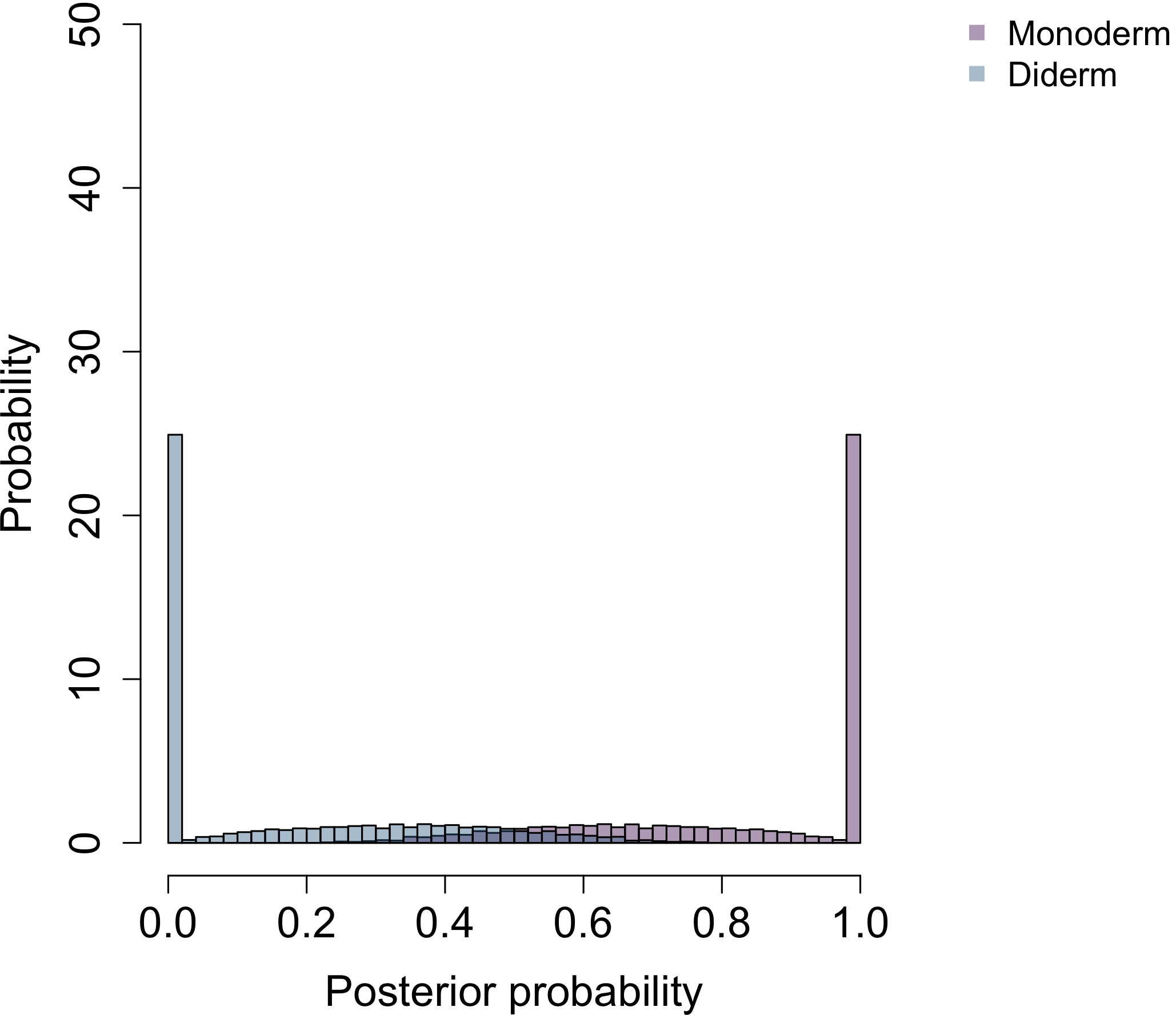

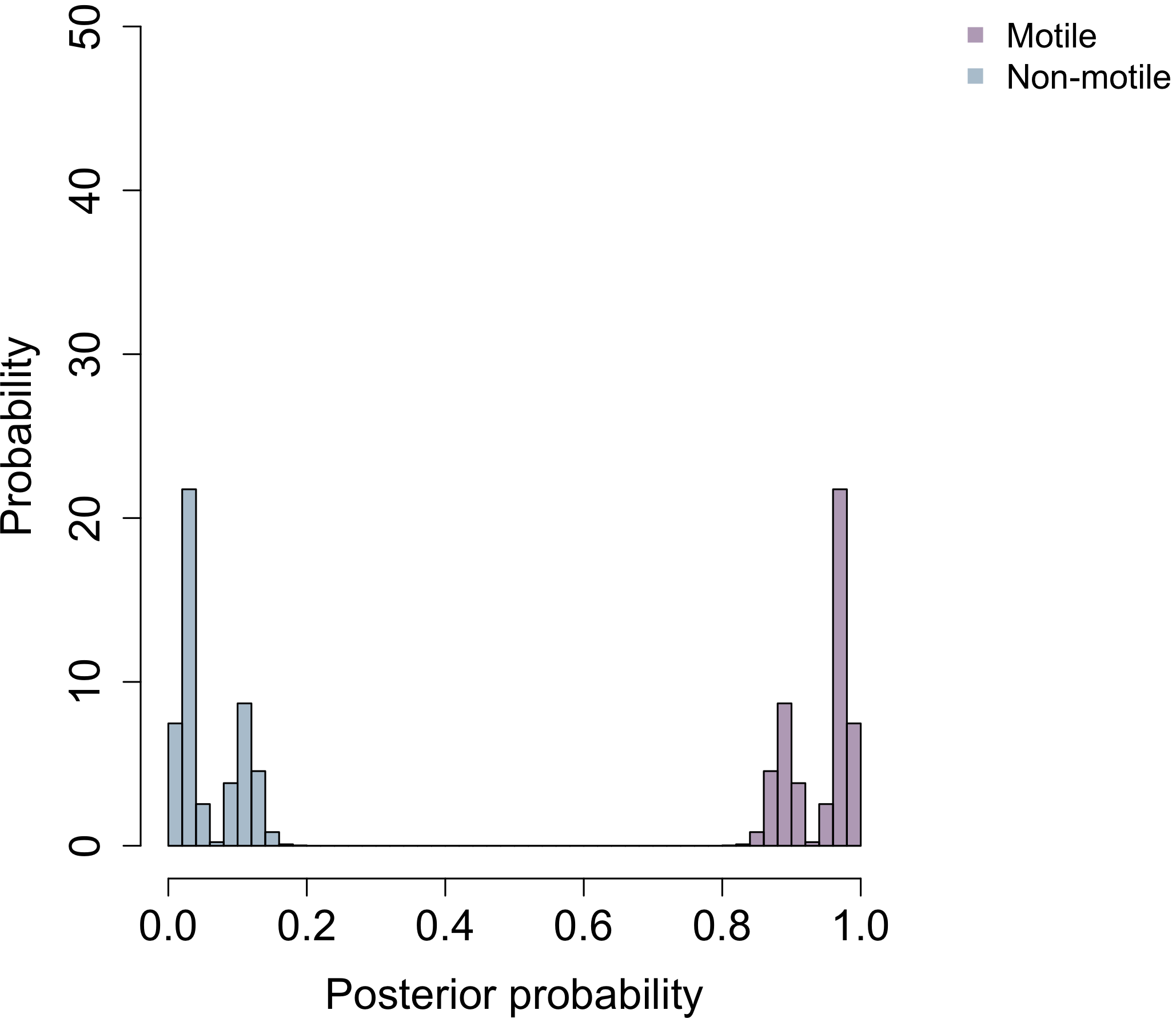

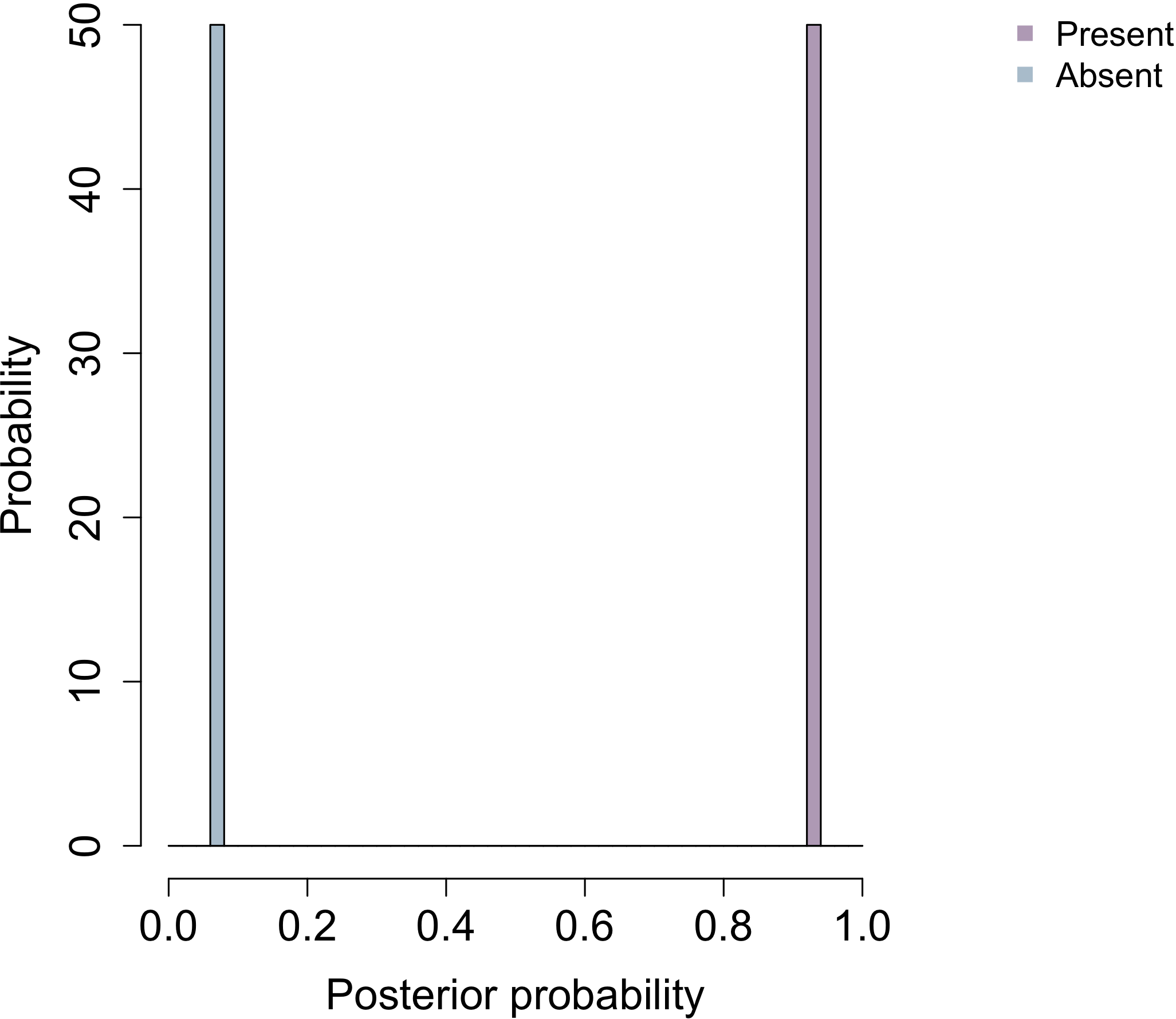

**Cell aggregation**

**Habitat b**

**Habitat a**

**Cell plan**

**Cell wall**

**Motility**

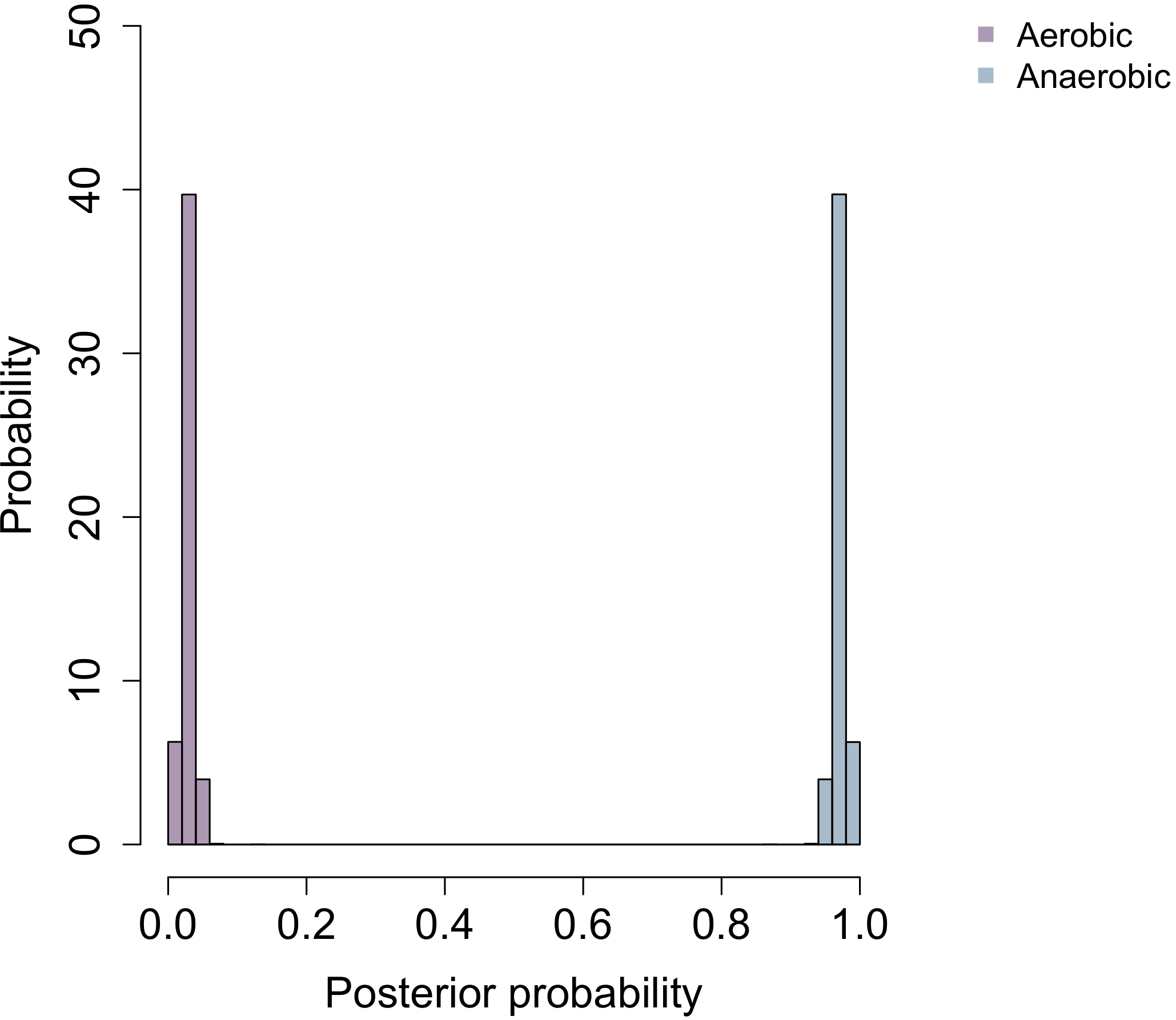

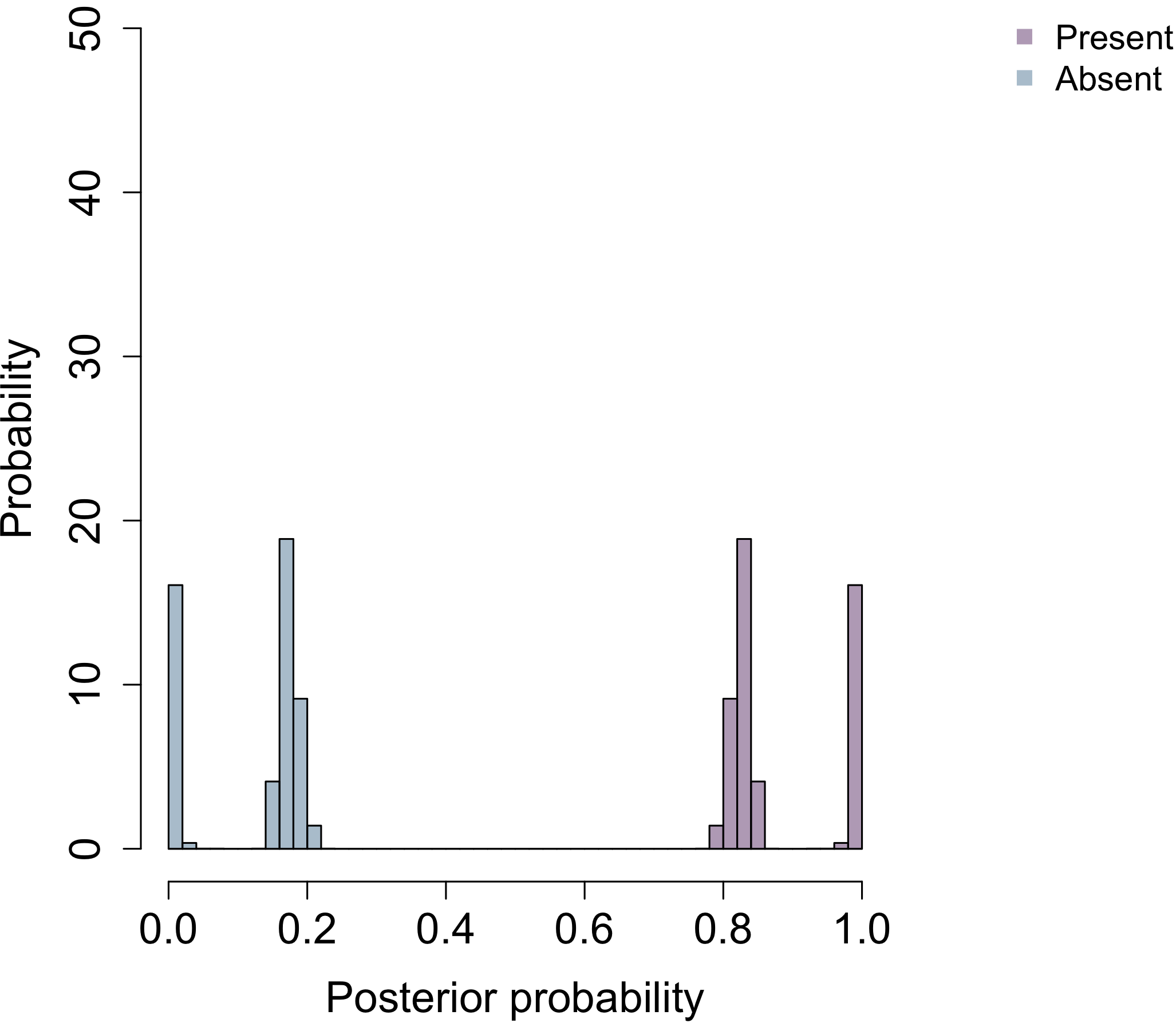

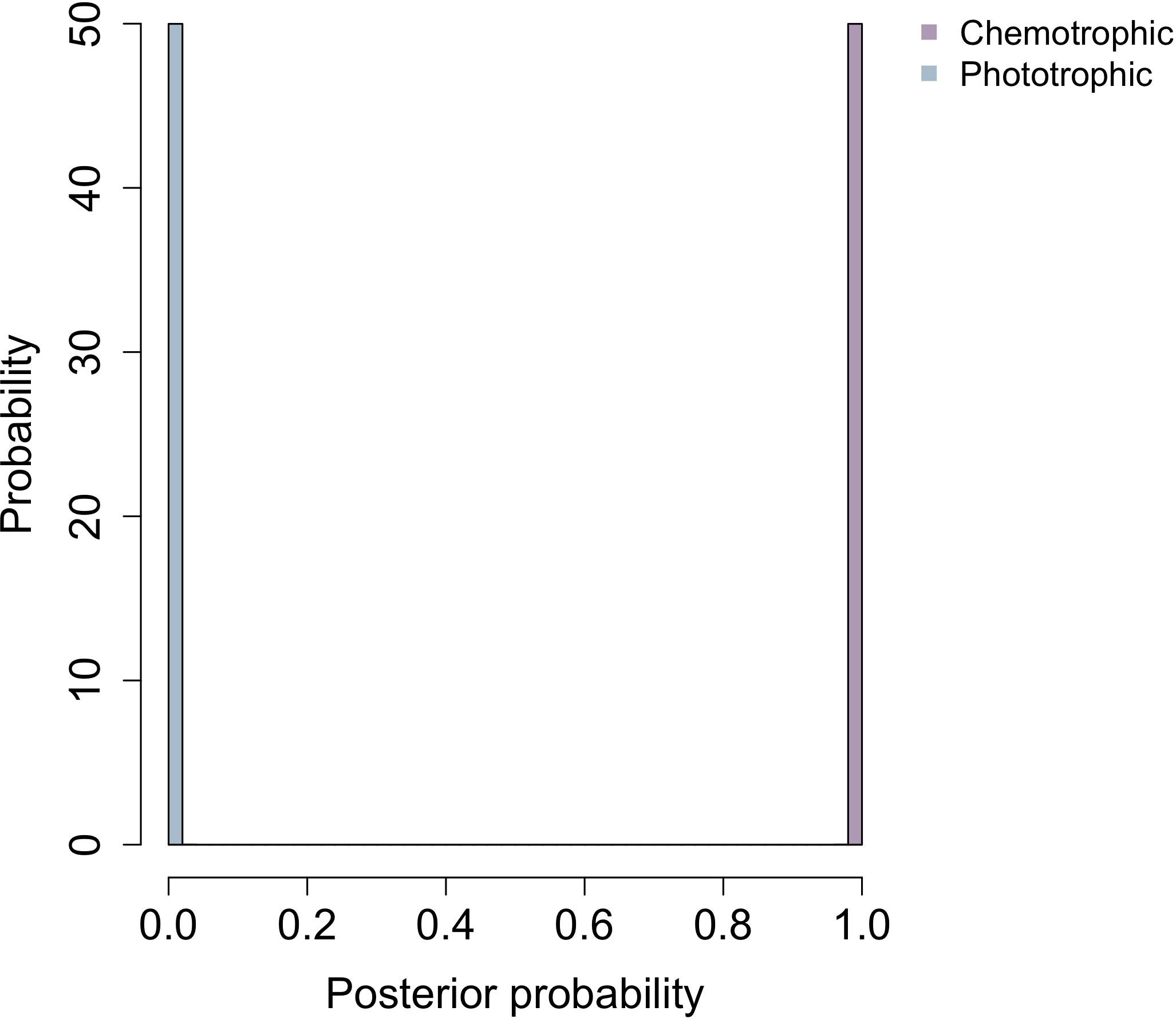

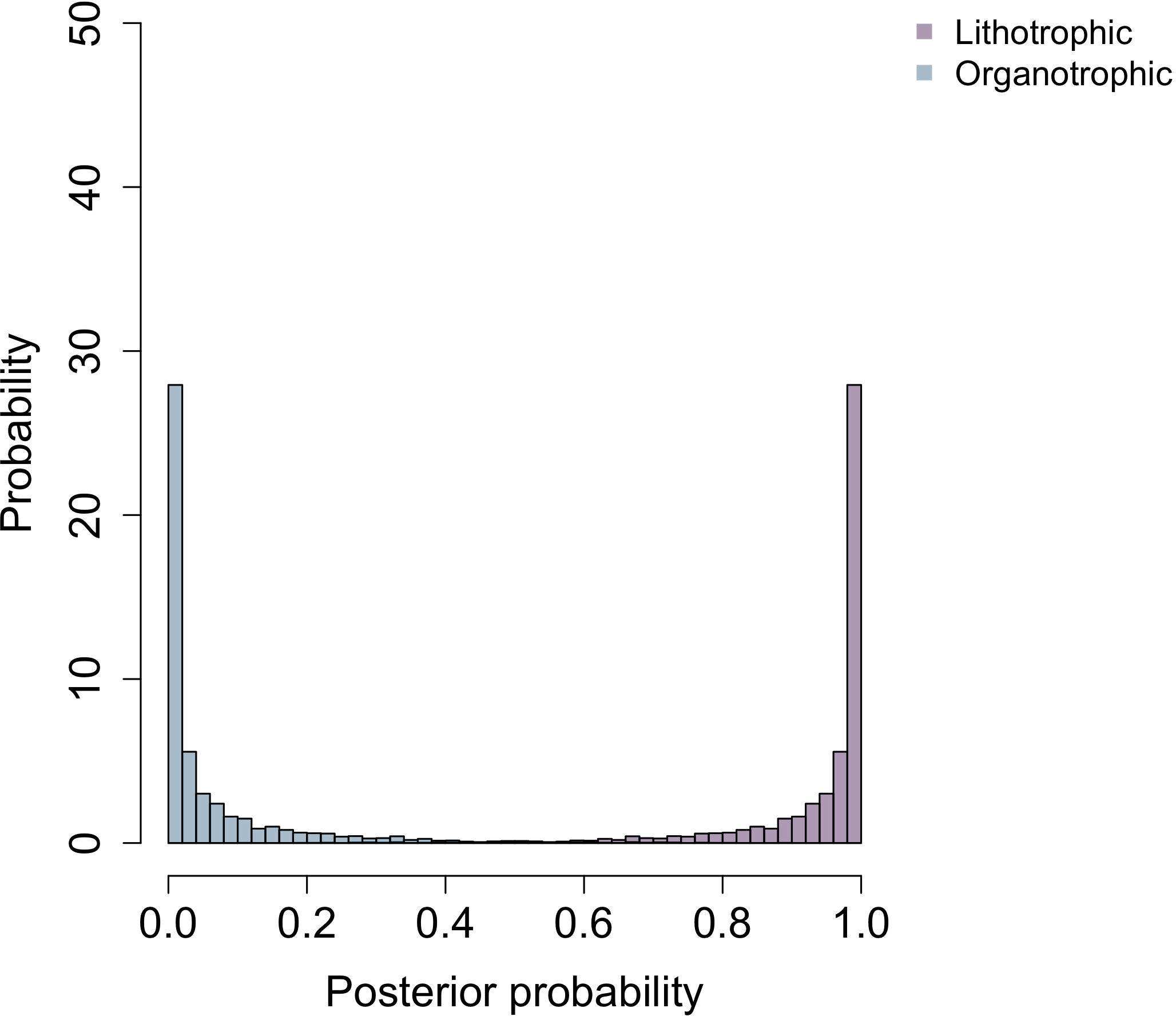

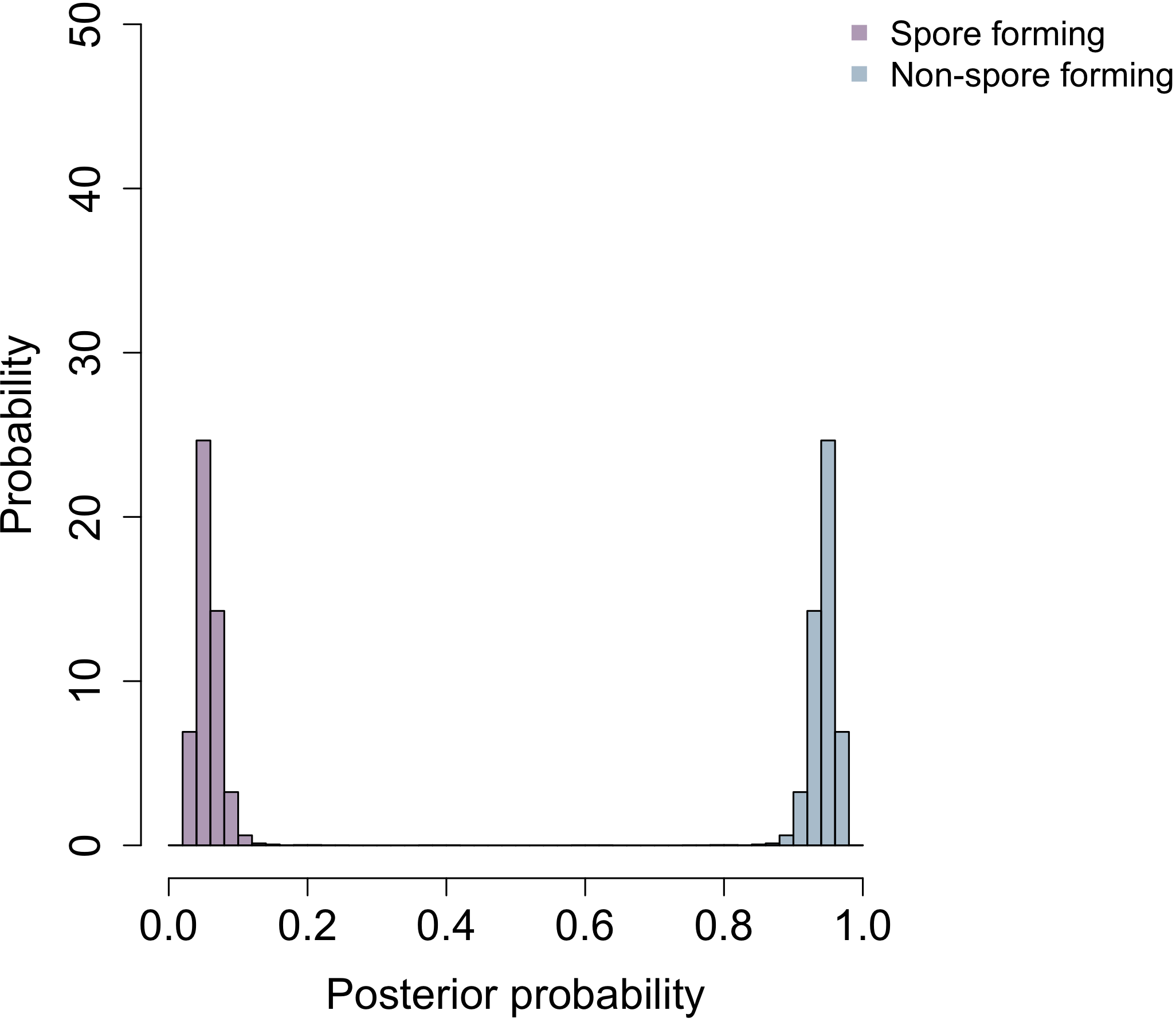

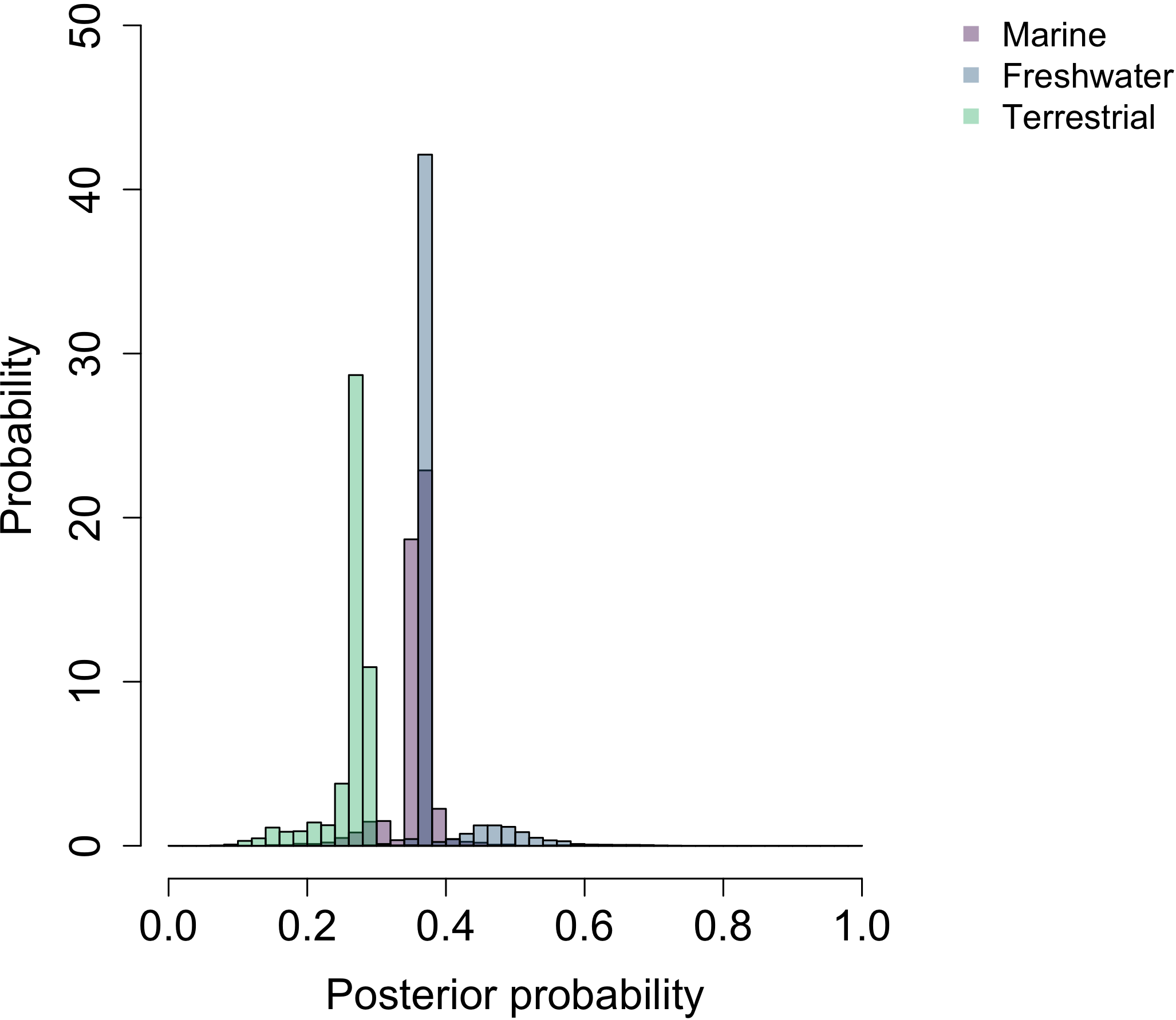

**Oxygen requirement**

**Energy source**

**Catalase**

**Electron donor**

**Sporulation**

**Habitat c**

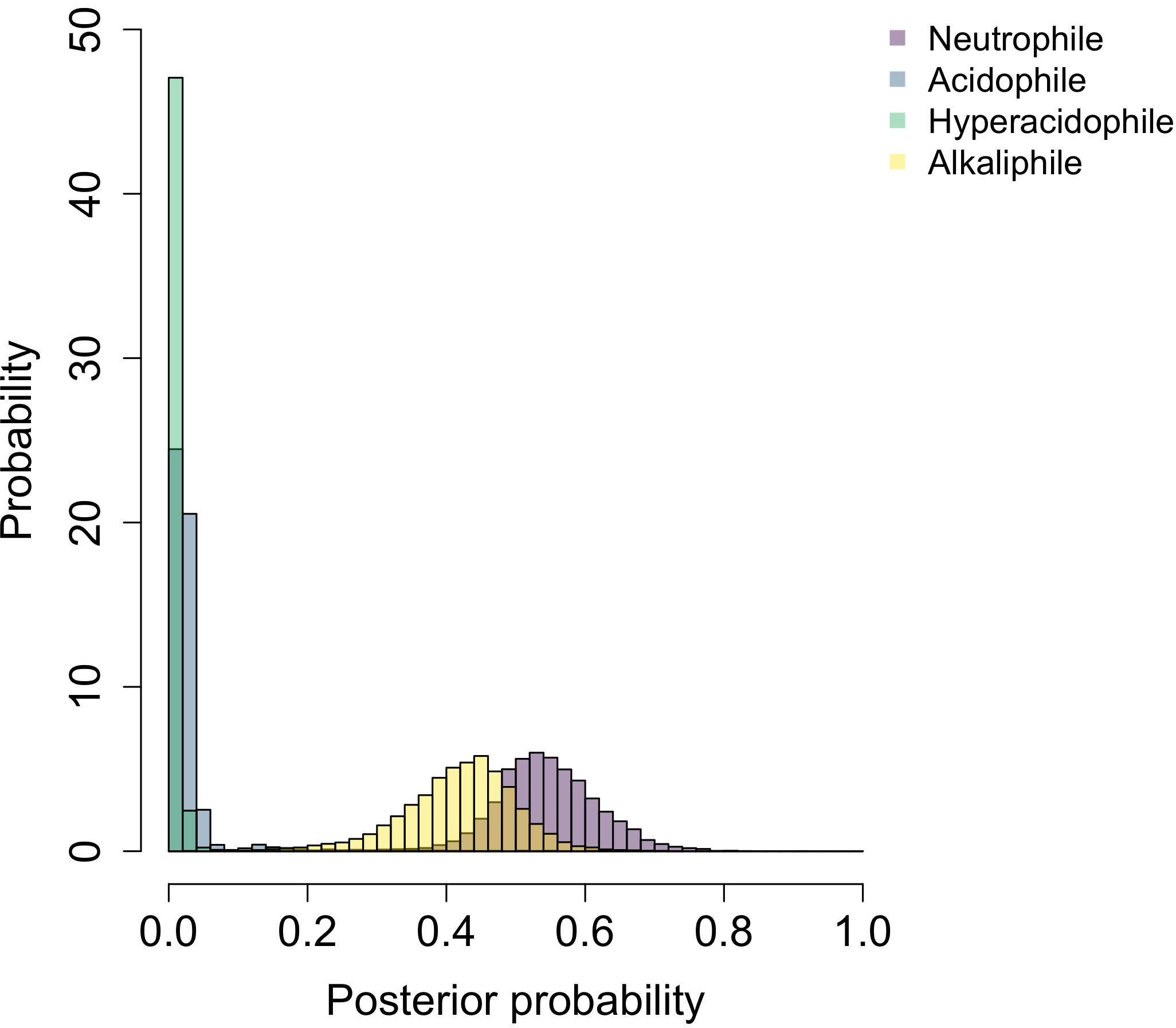

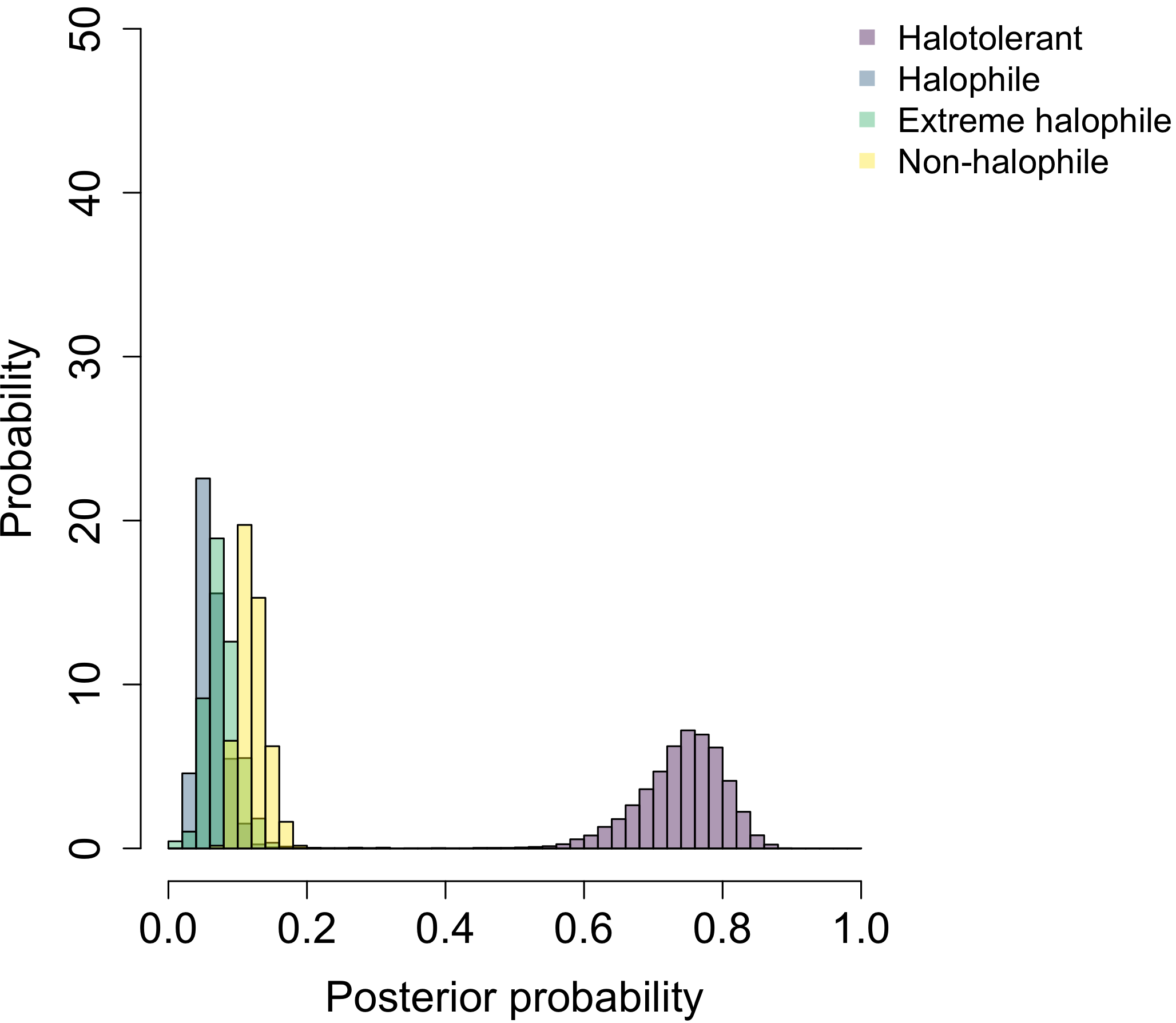

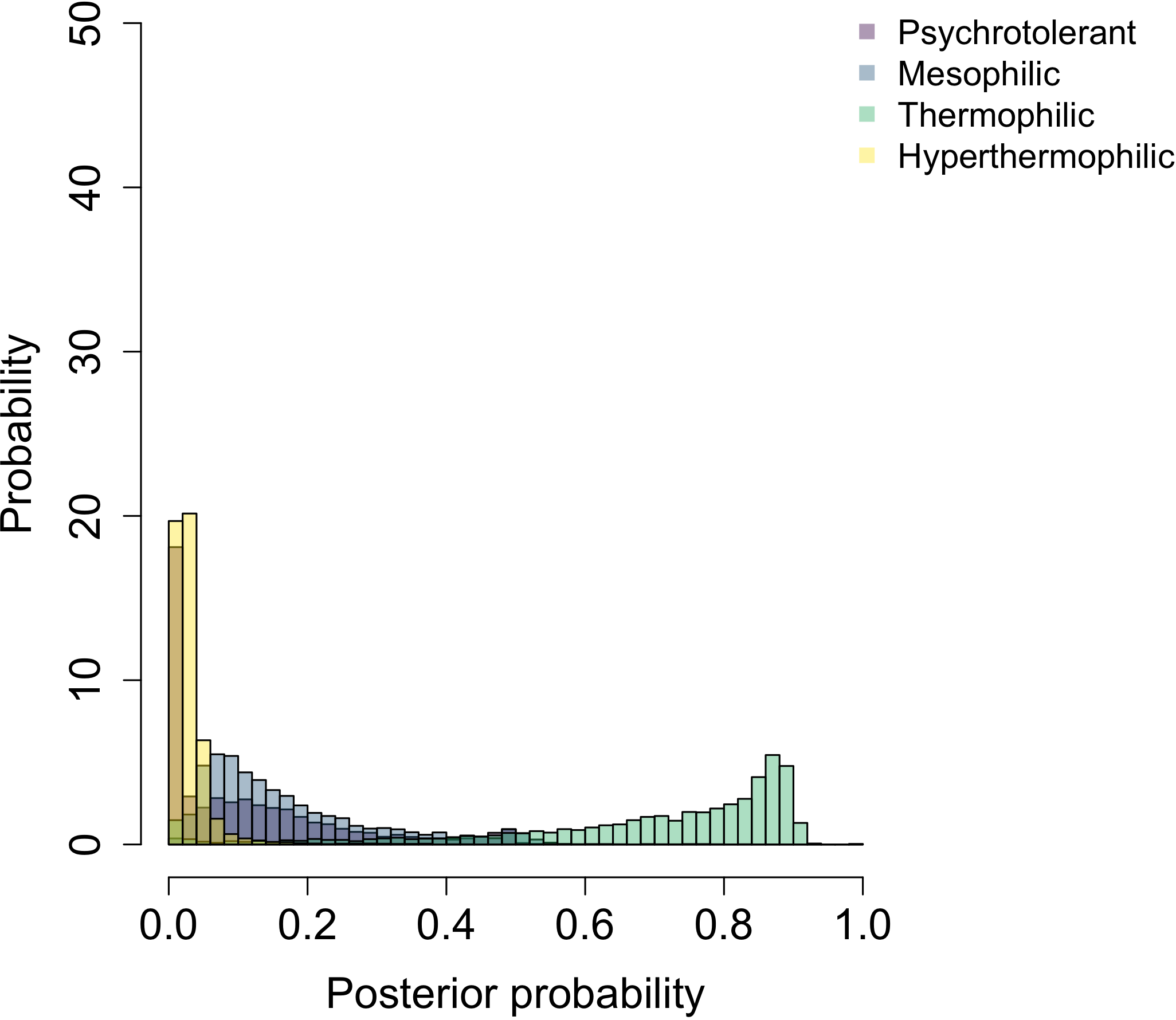

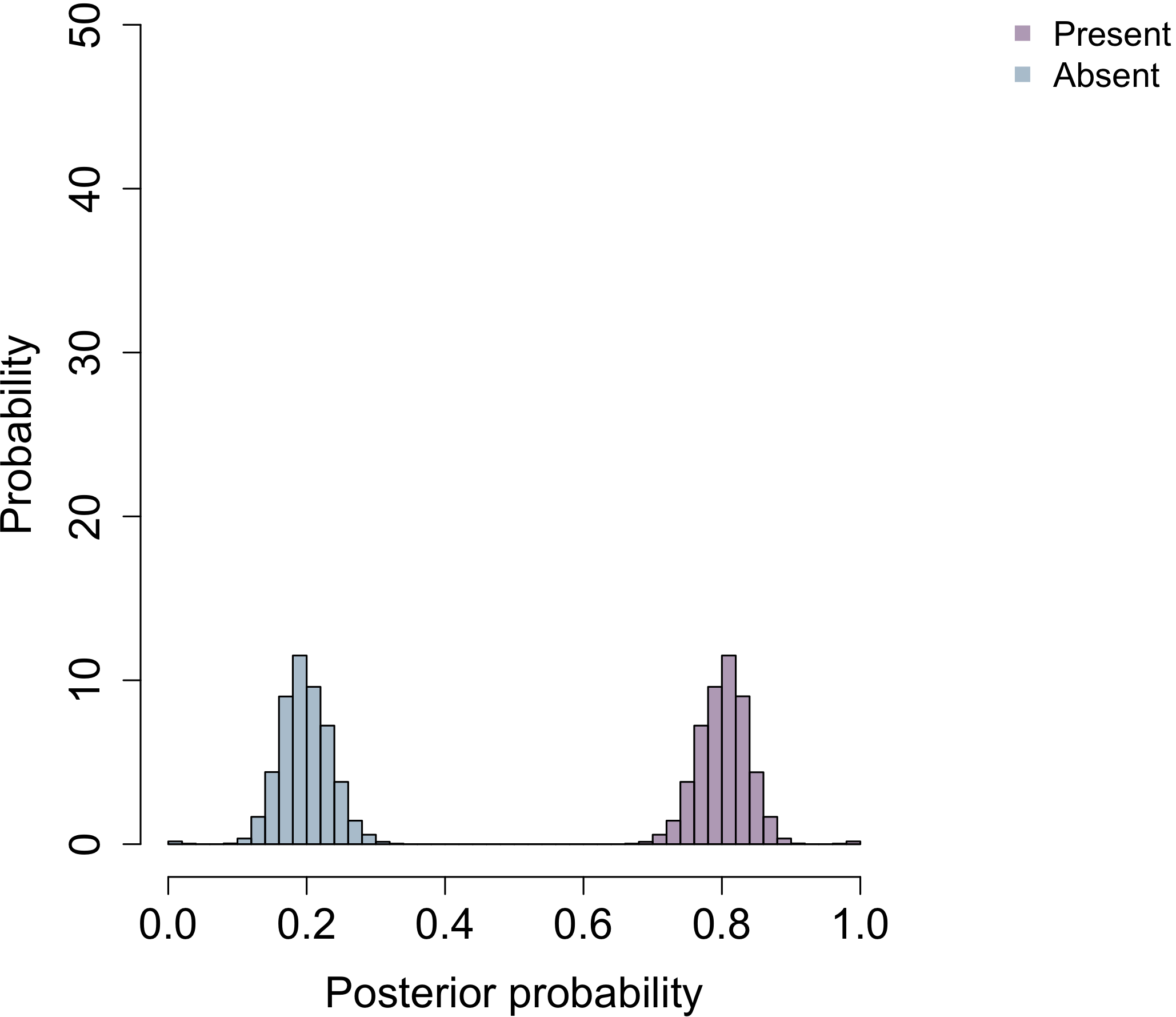

**NaCl**

**pH**

**Temperature**

**Oxidase**

**c) LACA**

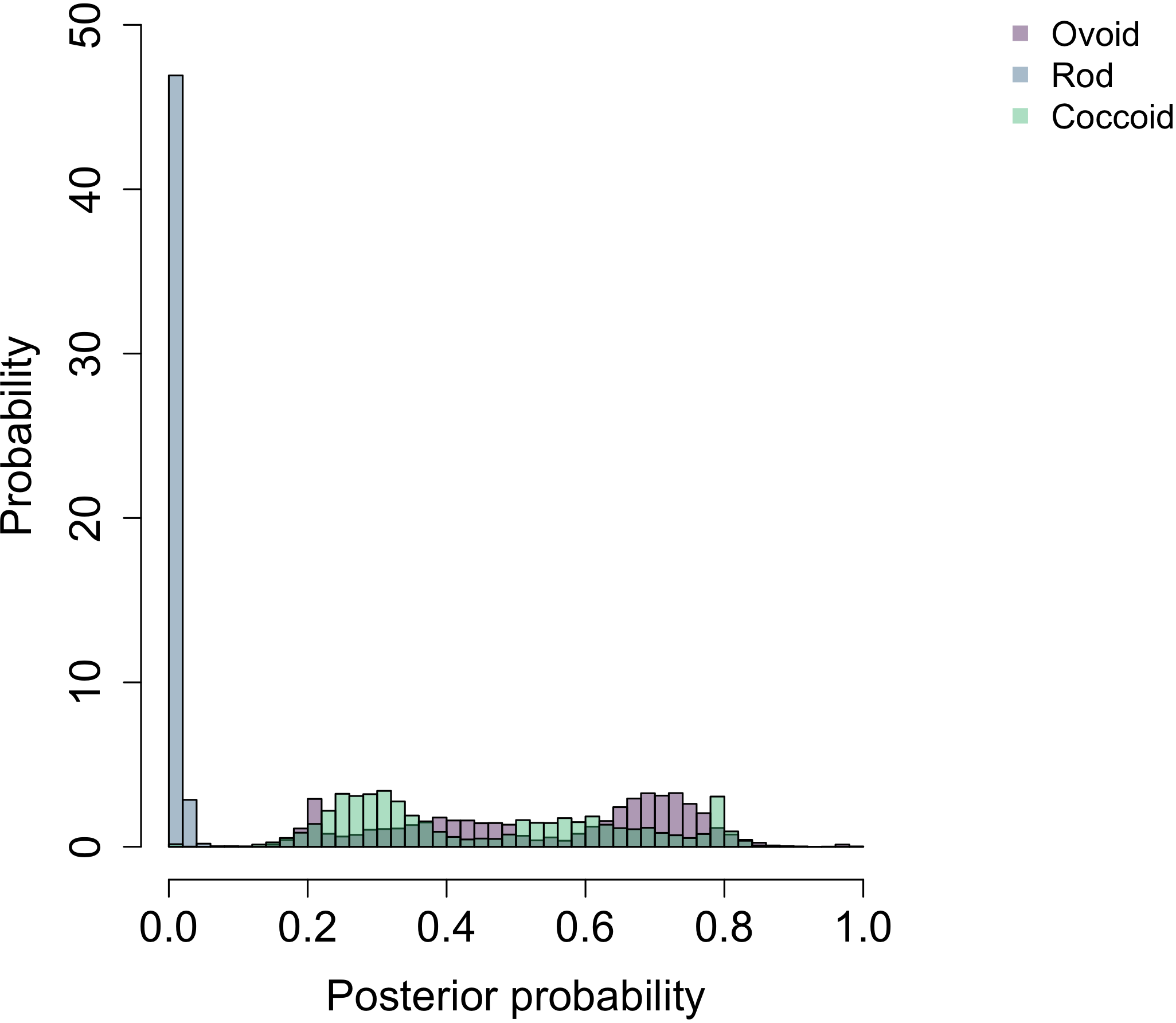

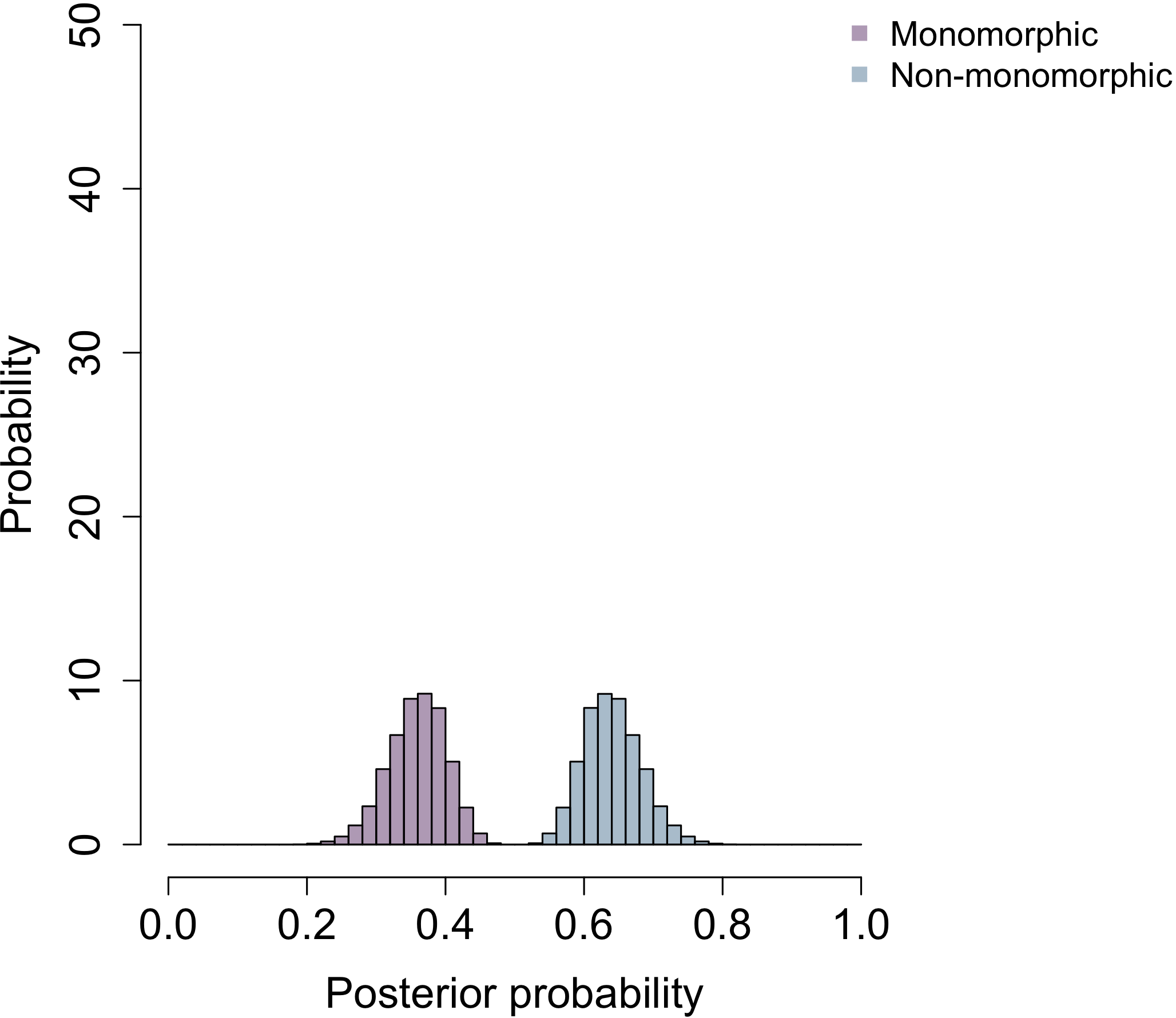

**Shape**

**Pleomorphism**

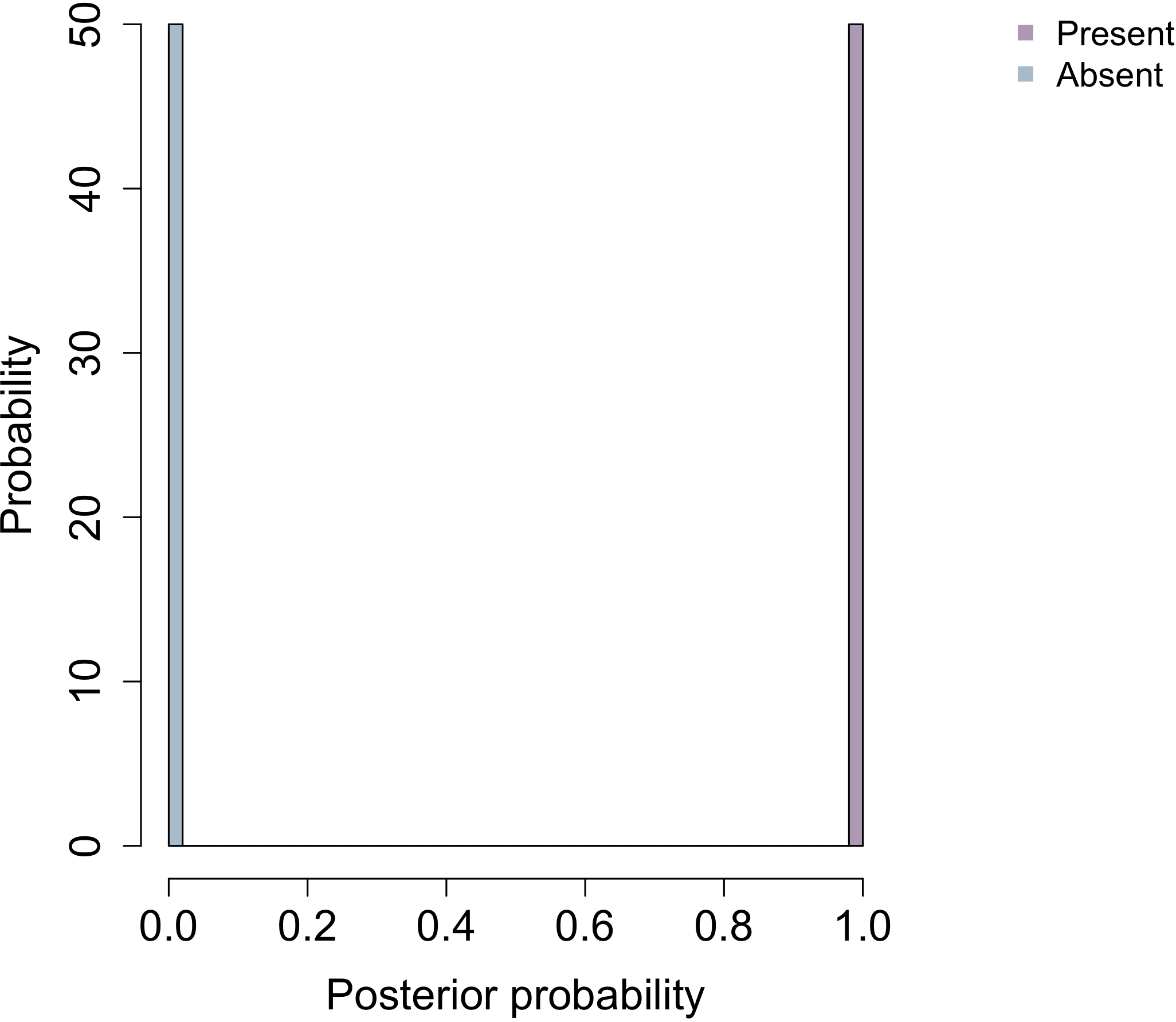

**Motility**

**Cell wall**

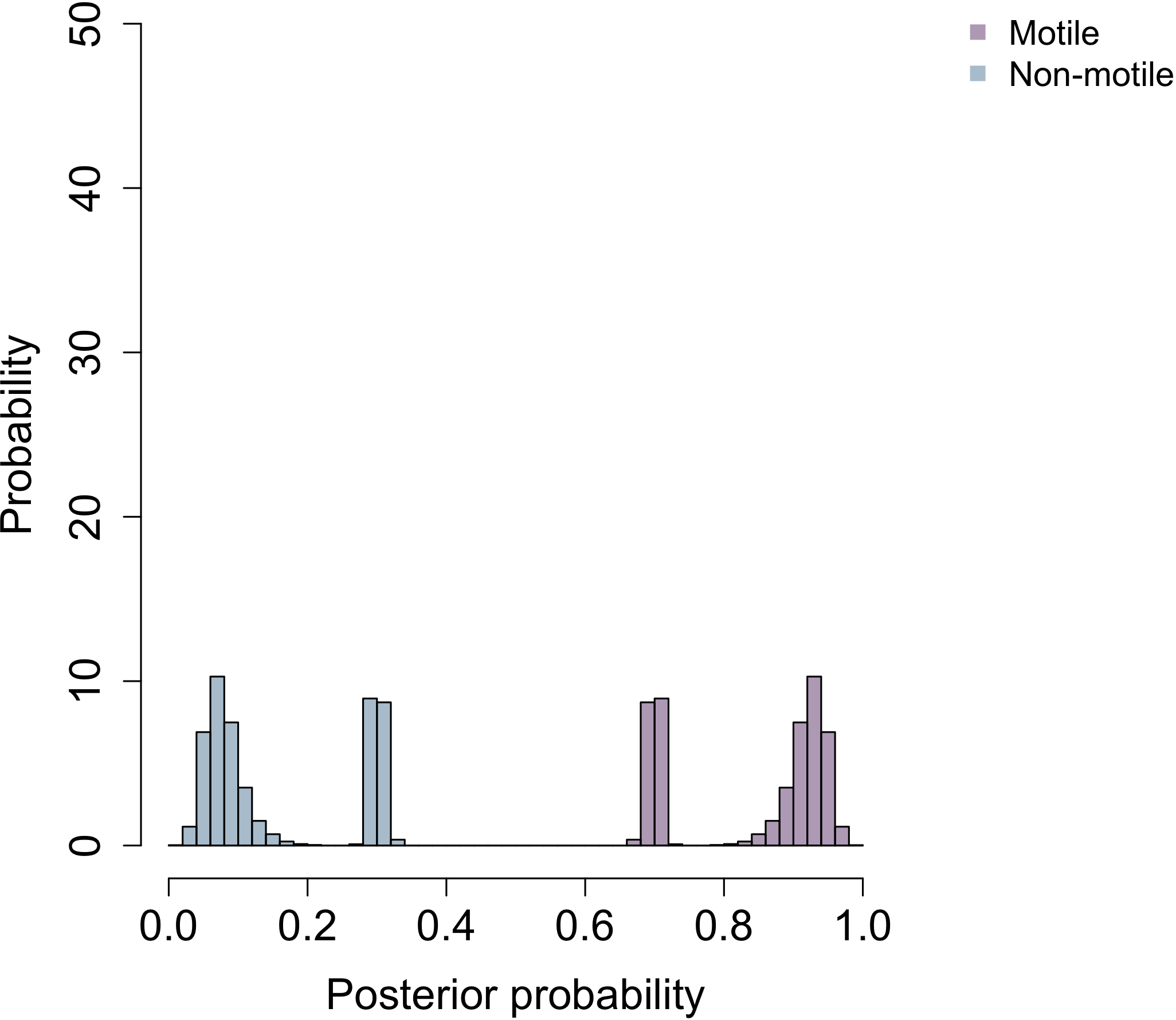

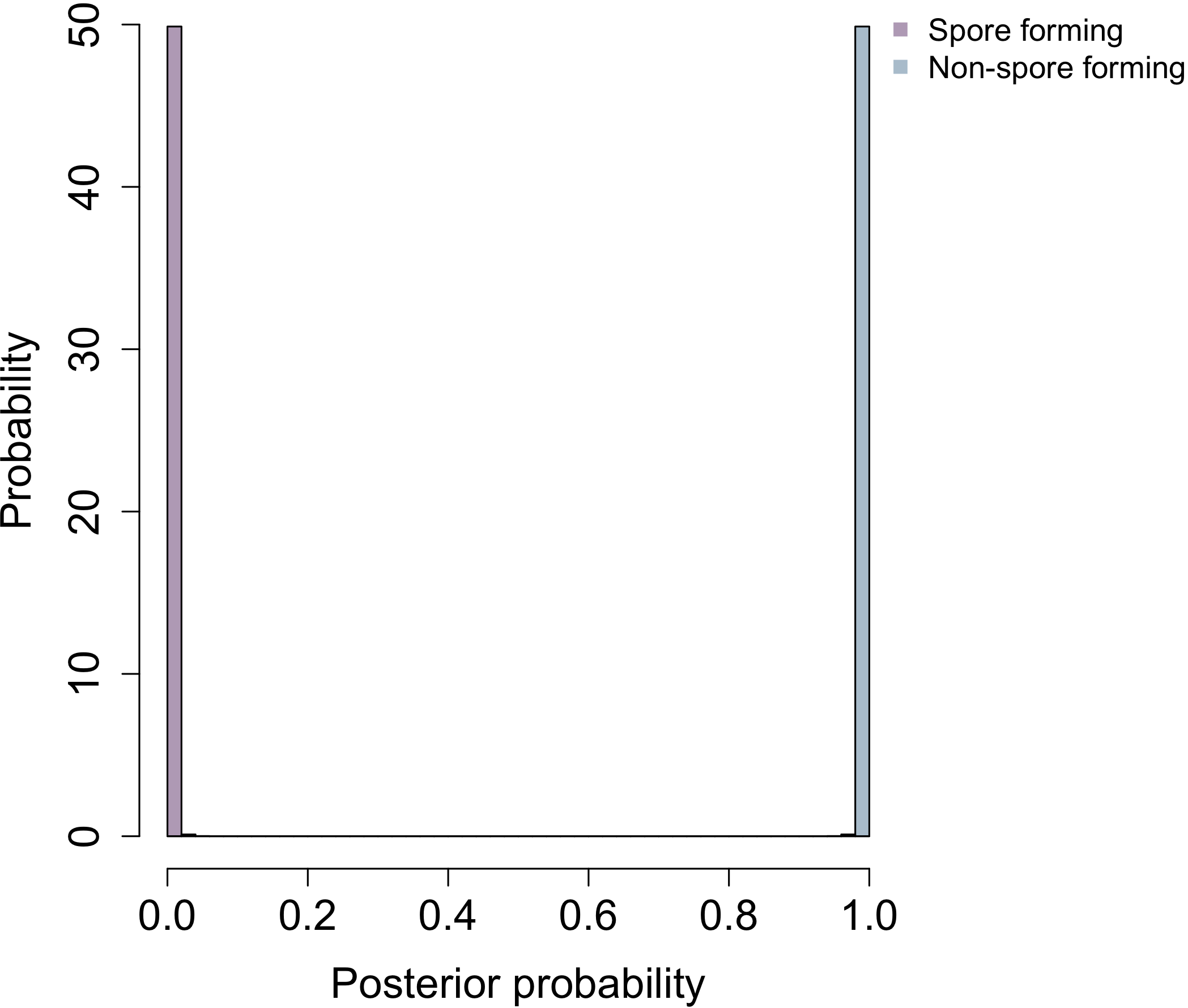

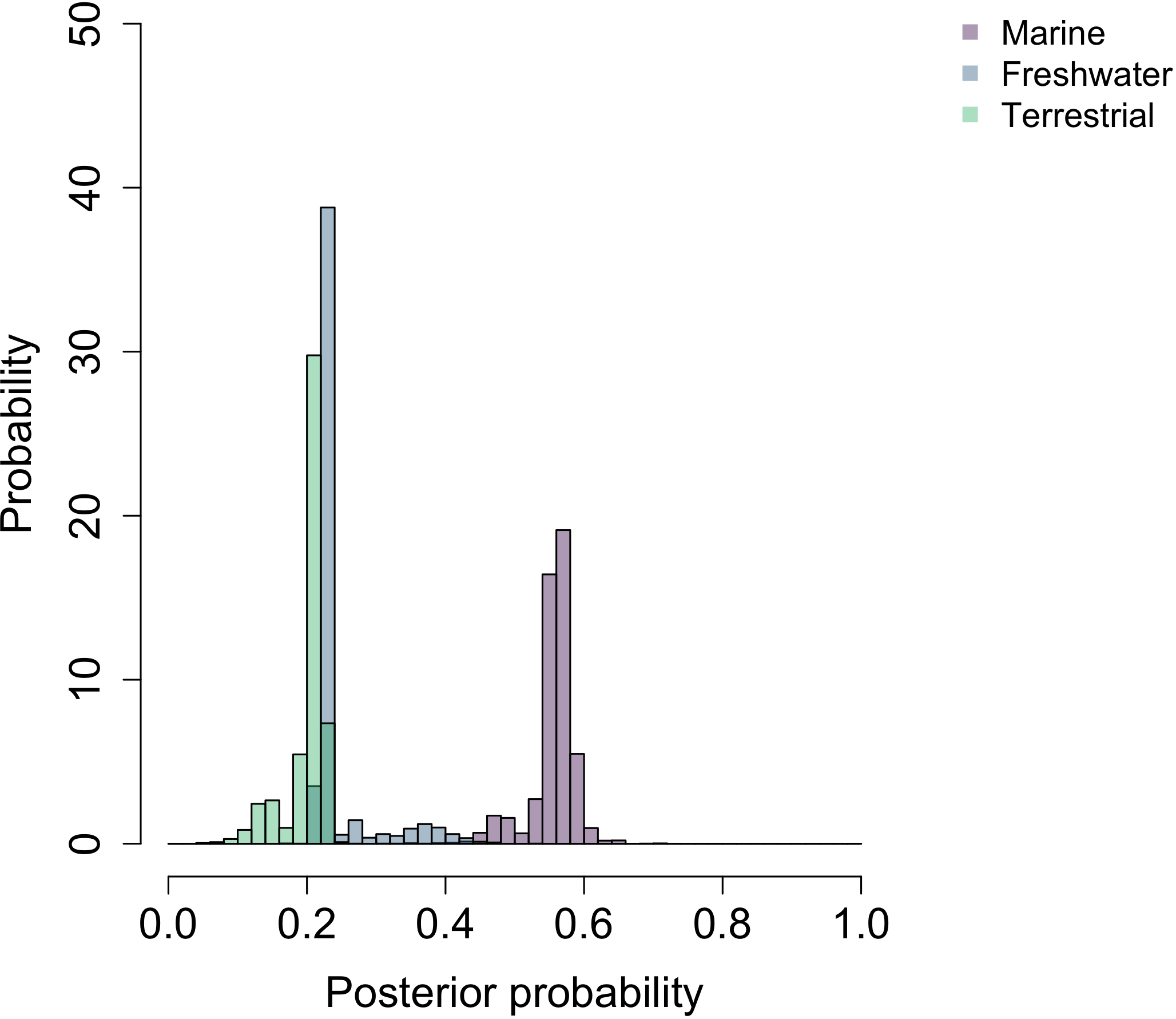

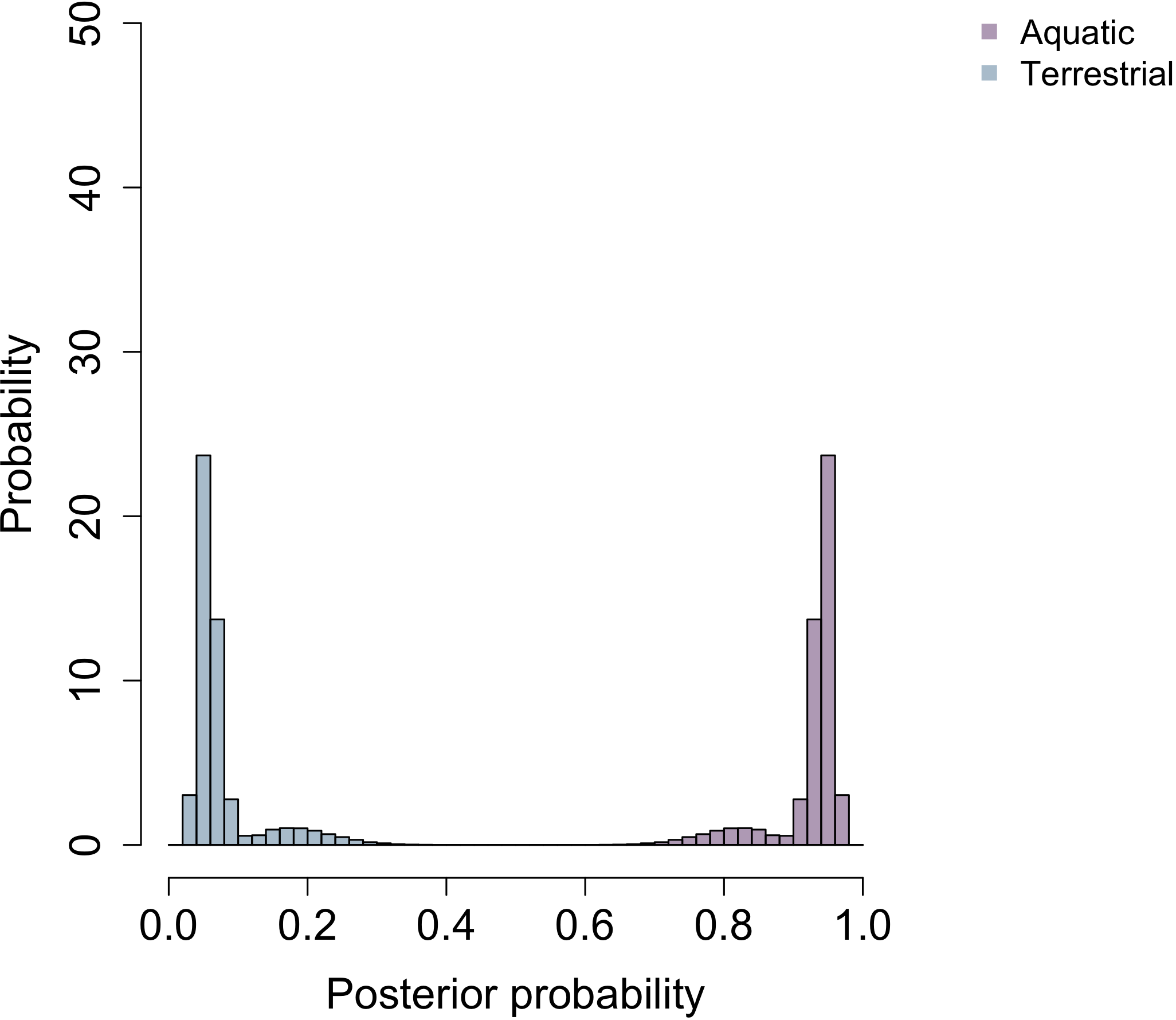

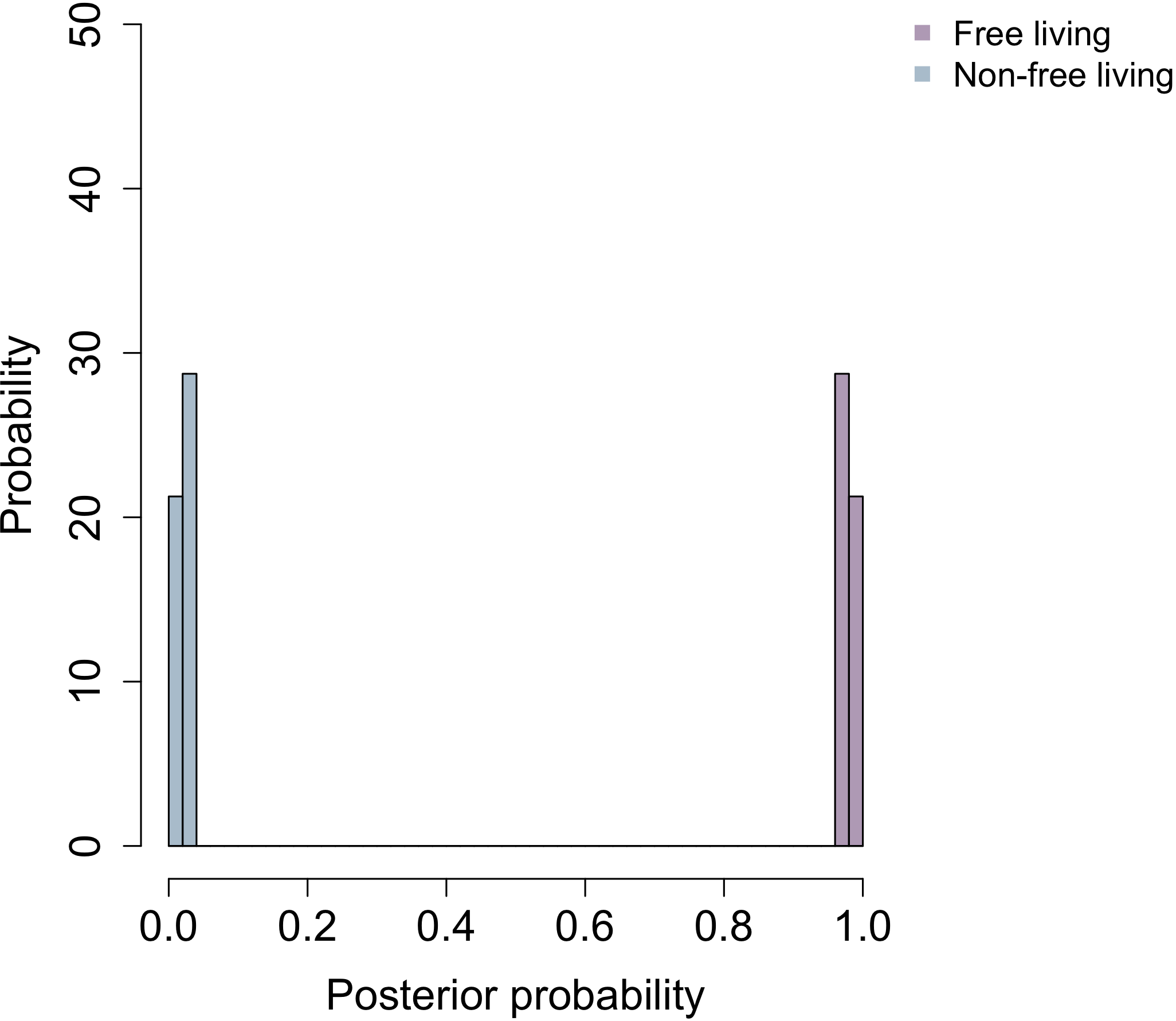

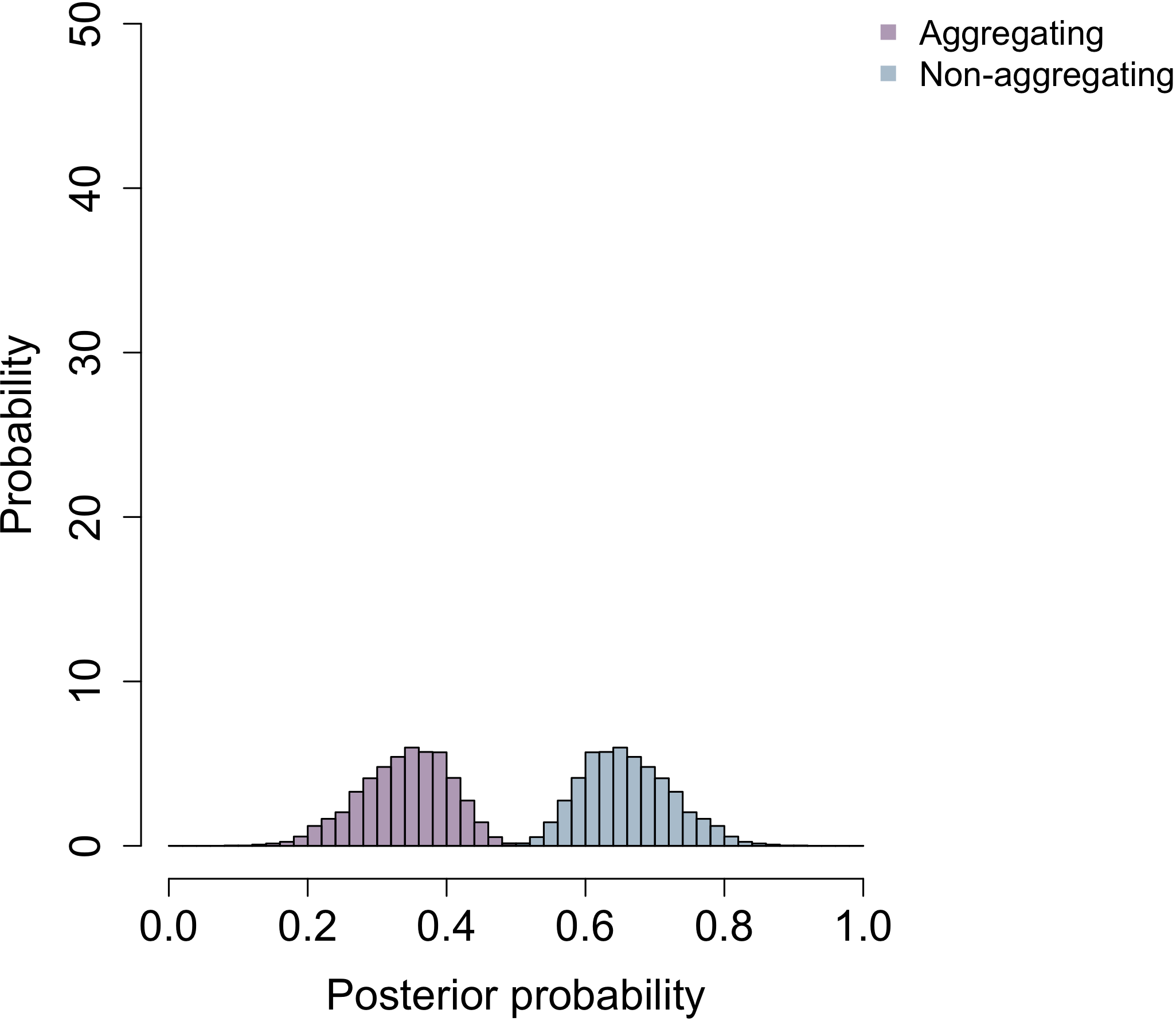

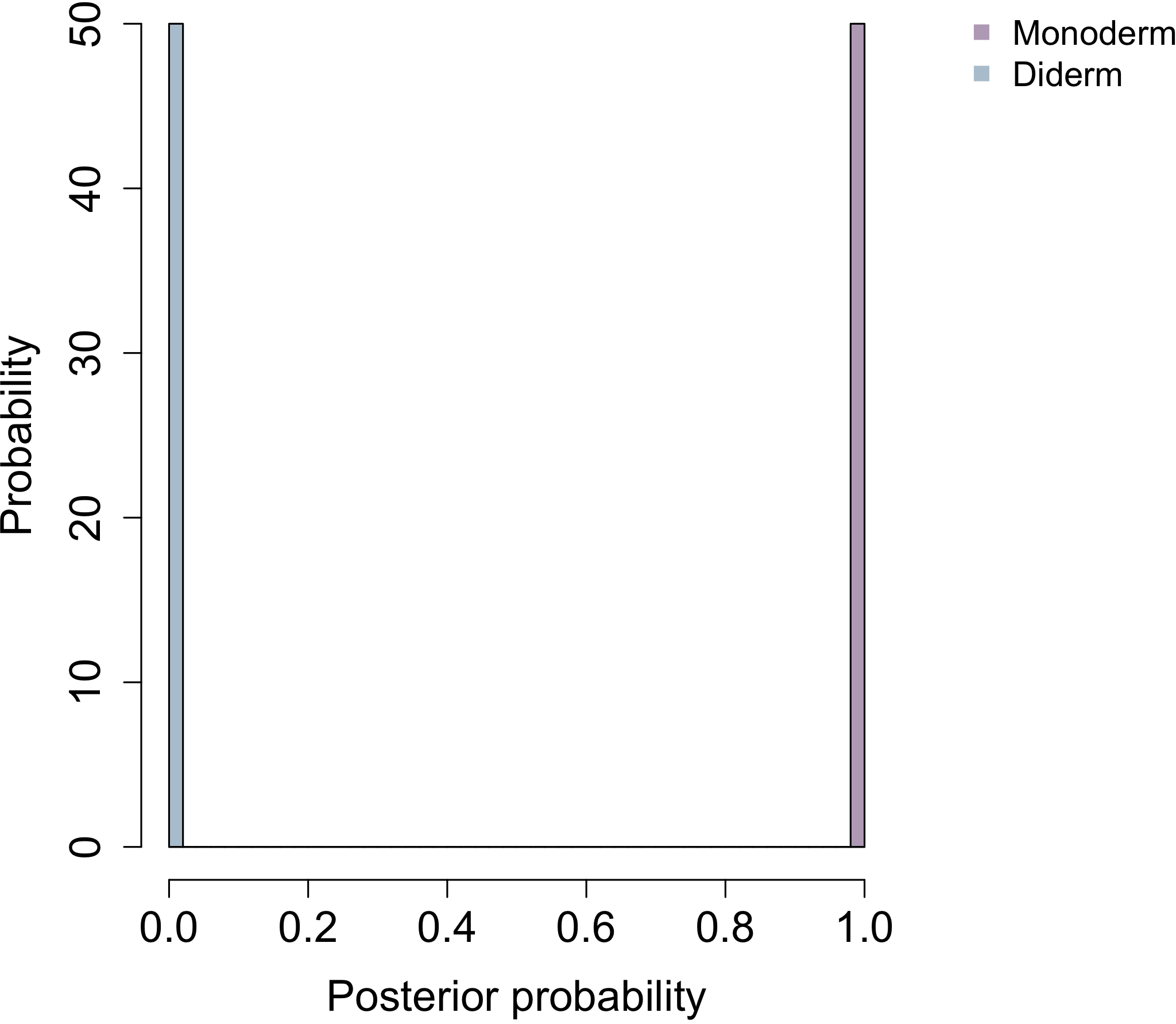

**Sporulation**

**Habitat b**

**Habitat c**

**Habitat a**

**Cell plan**

**Cell aggregation**

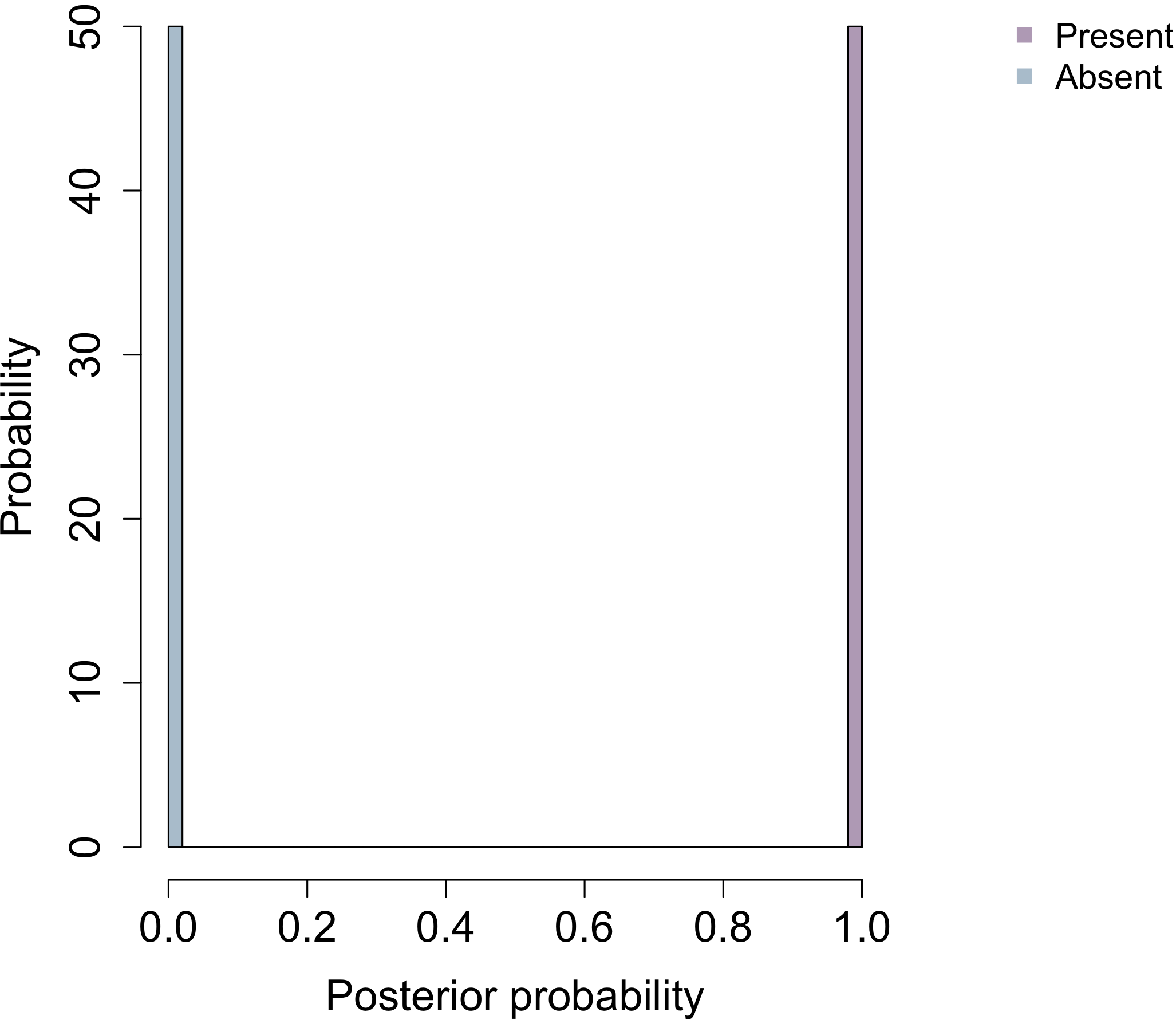

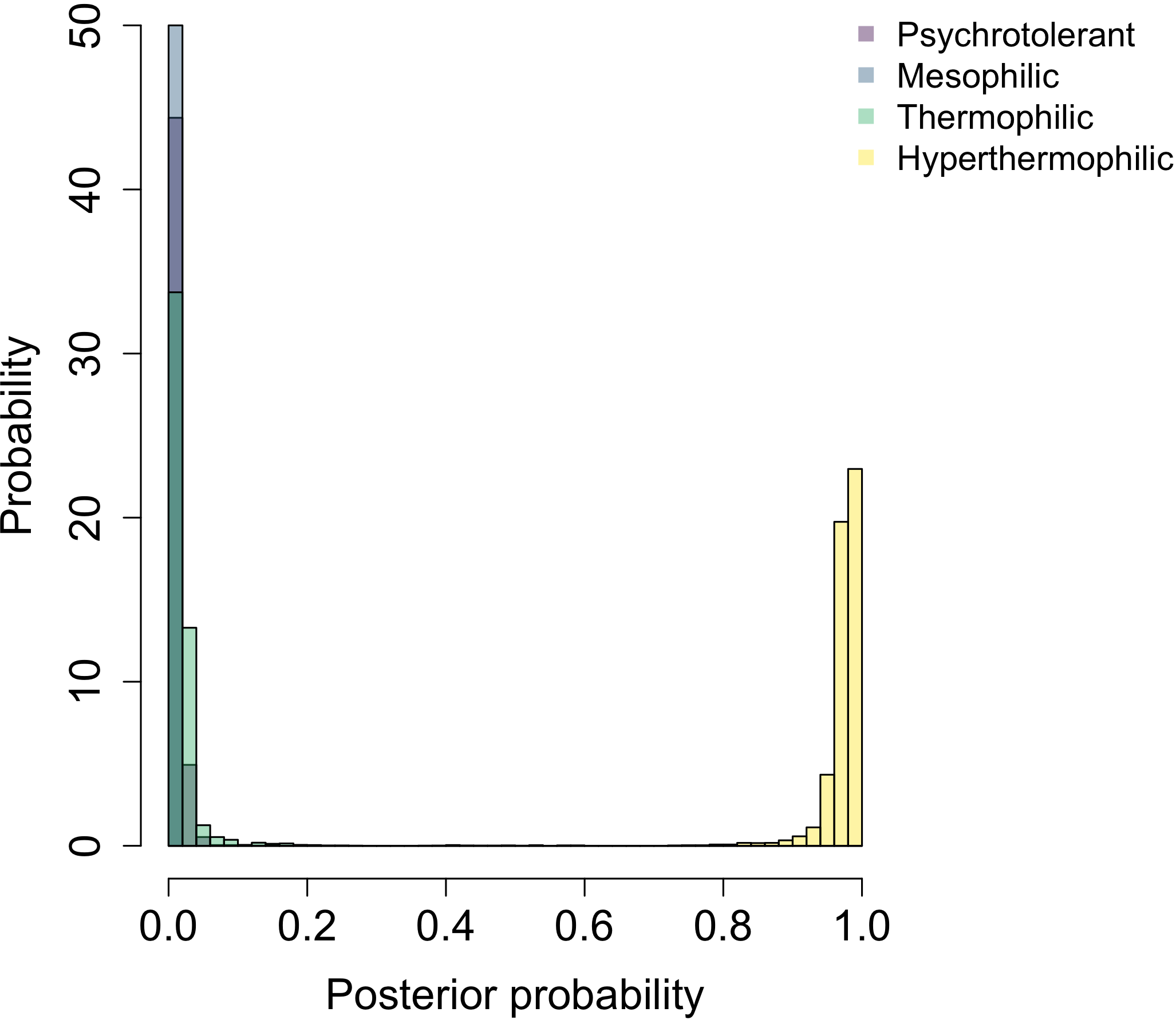

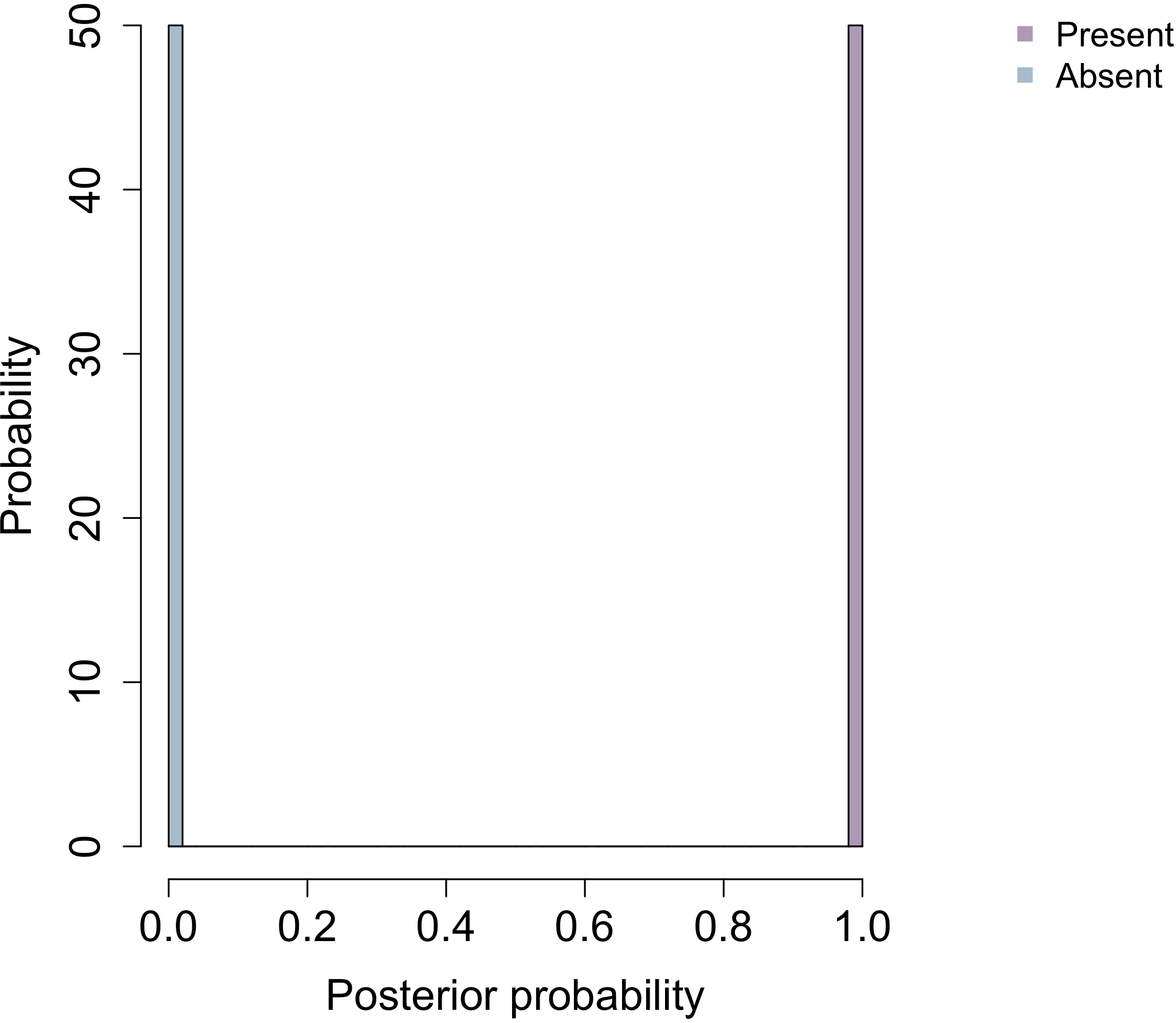

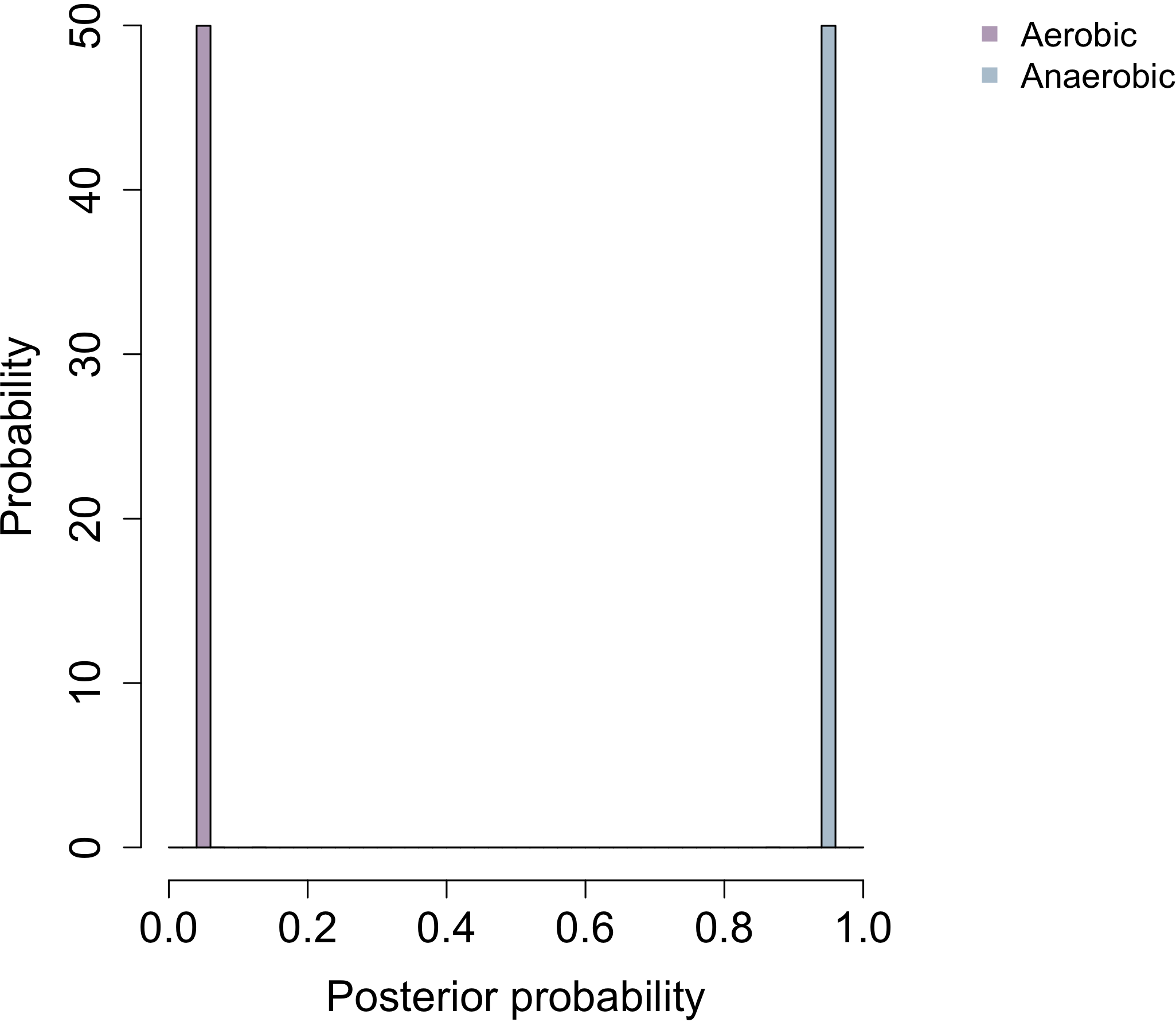

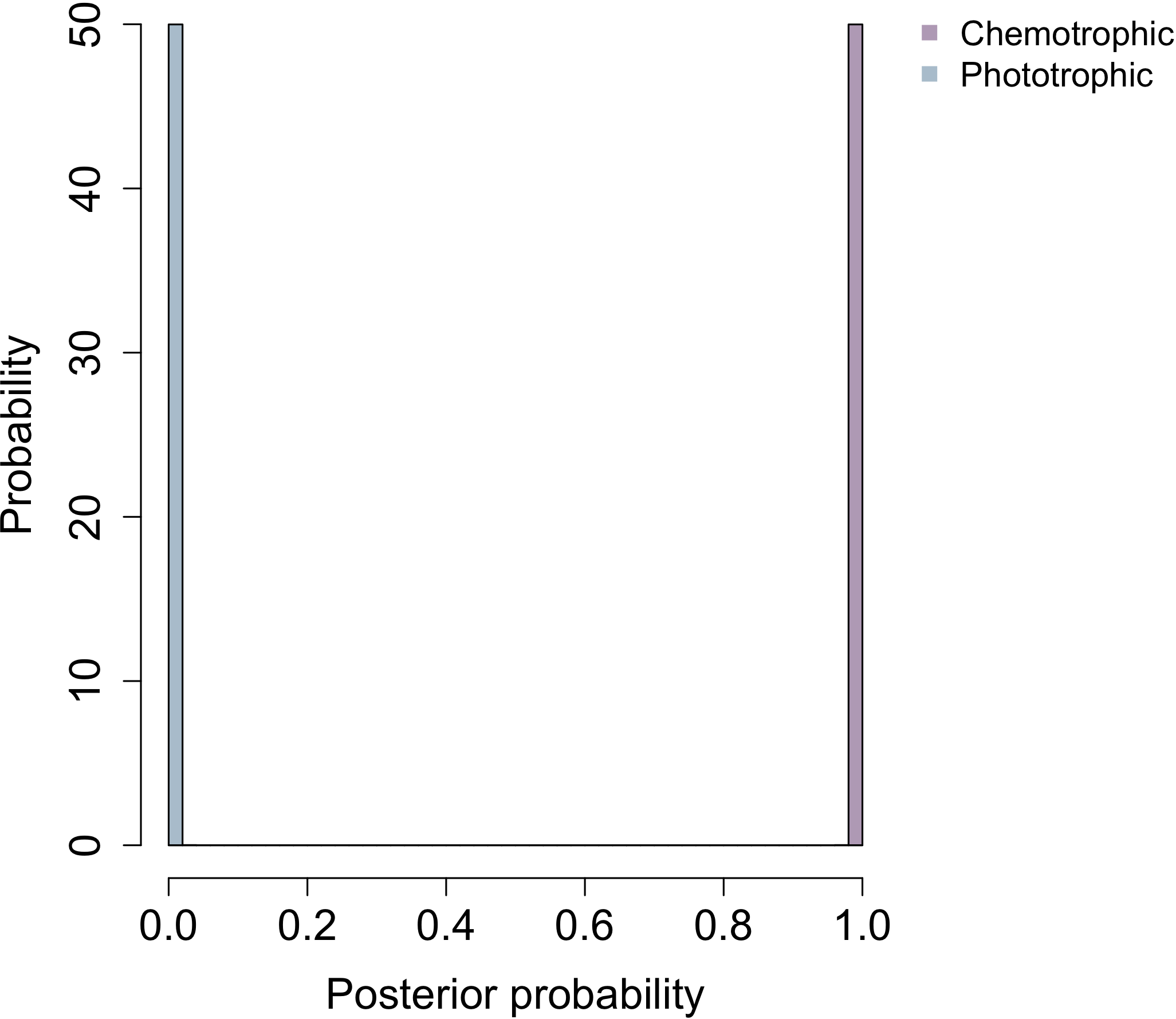

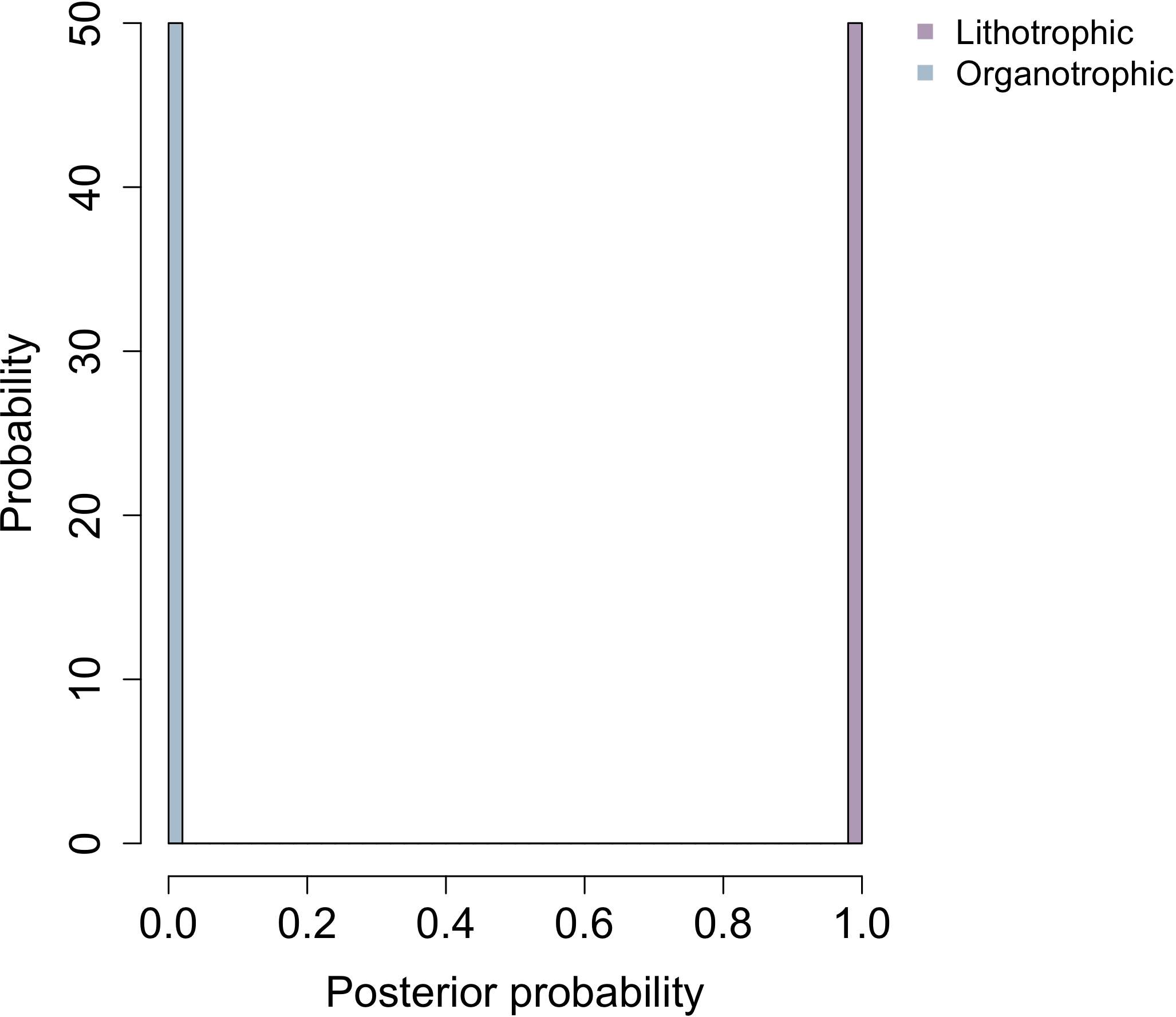

**Temperature**

**Oxidase**

**Catalase**

**Oxygen requirement**

**Energy source**

**Electron donor**

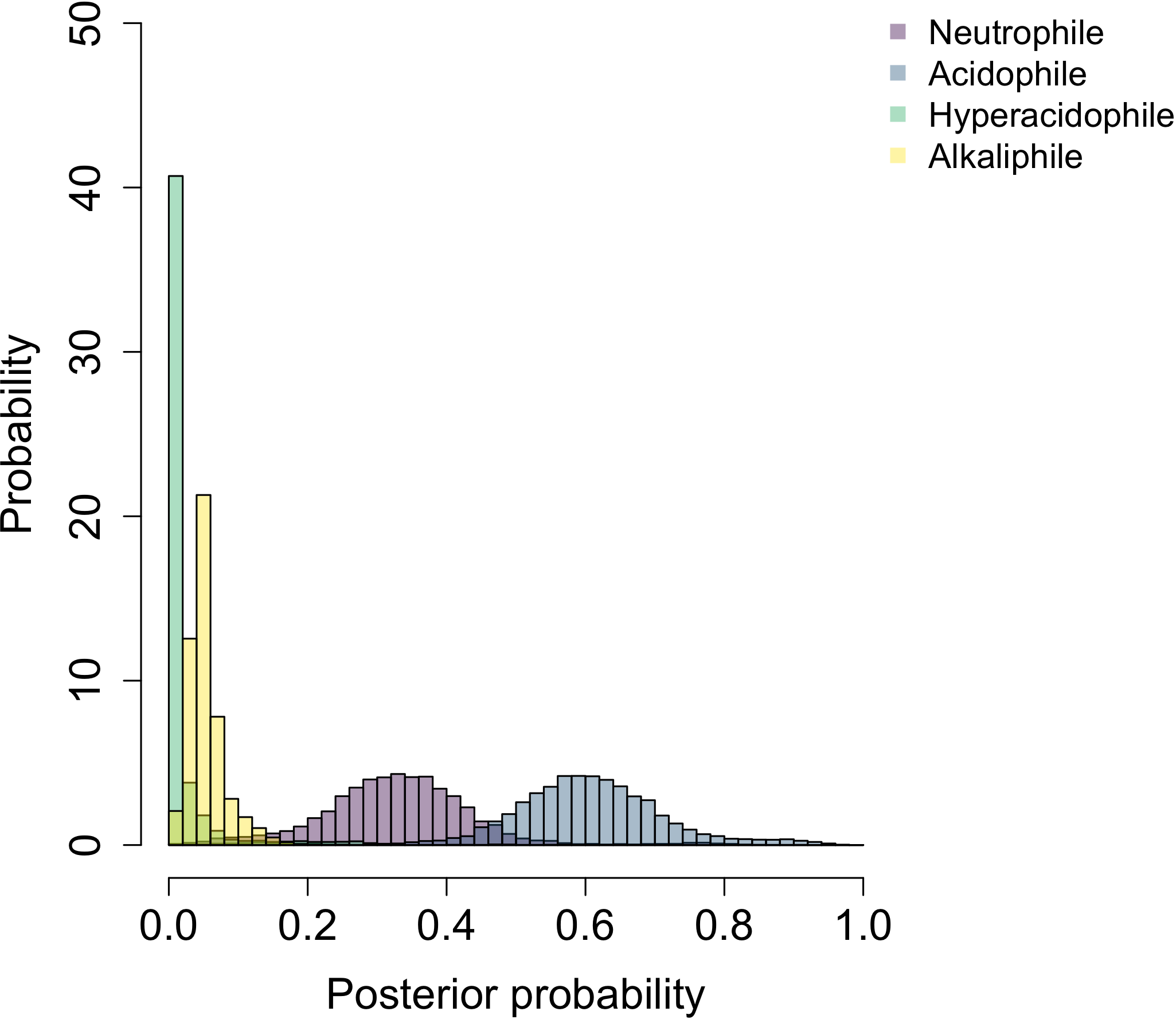

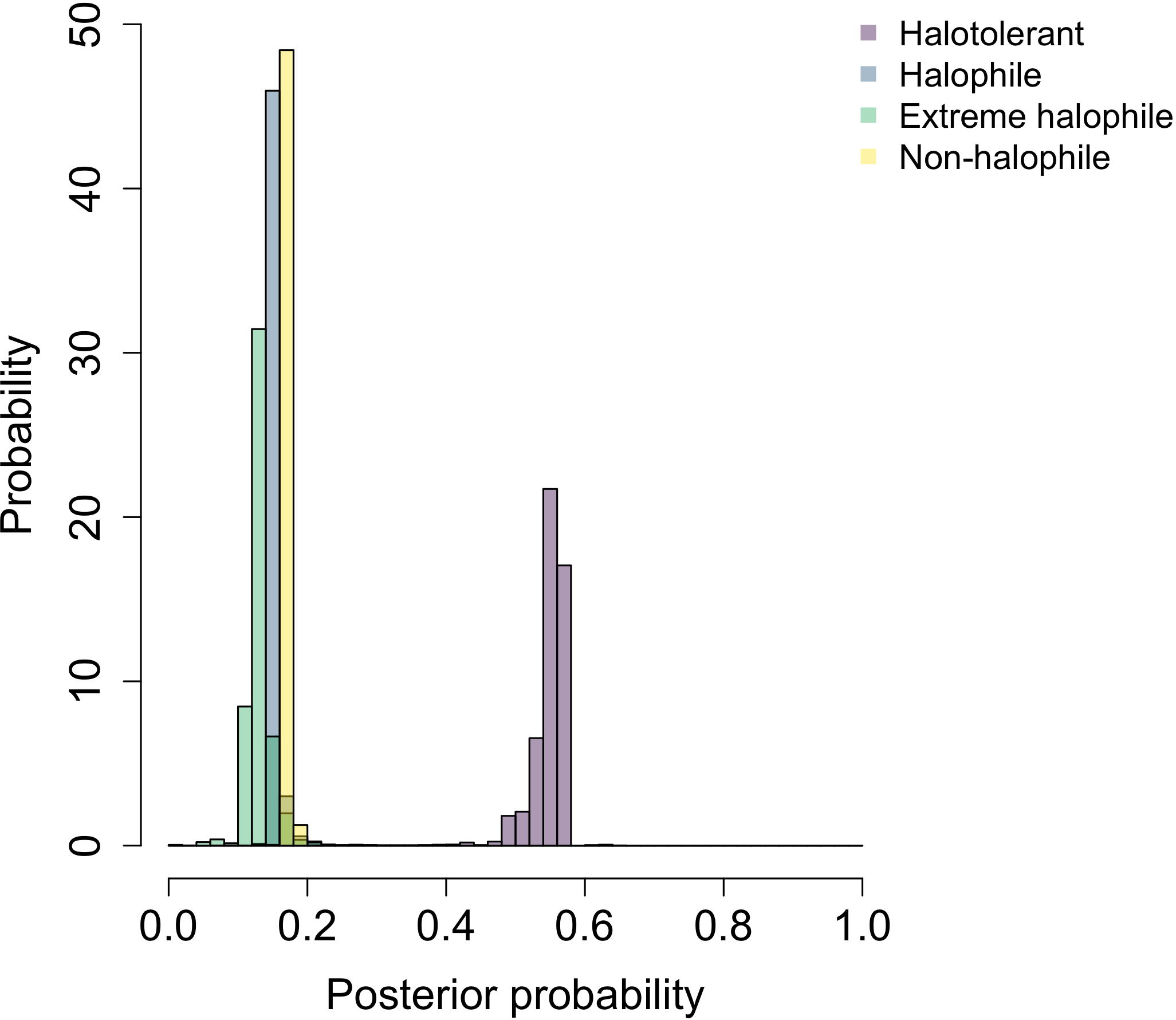

**pH**

**NaCl**

**Fig. S4. Posterior probability histograms for all the characters from Segata et al’s tree.**

Posterior probabilities for all the characters from Segata et al’s tree are shown as histograms for a) LUCA, b) LBCA and c) LACA. Abbreviations: LUCA - last universal common ancestor; LBCA -last bacterial common ancestor; LACA - last archaeal common ancestor;

**a) LUCA**

**Pleomorphism**

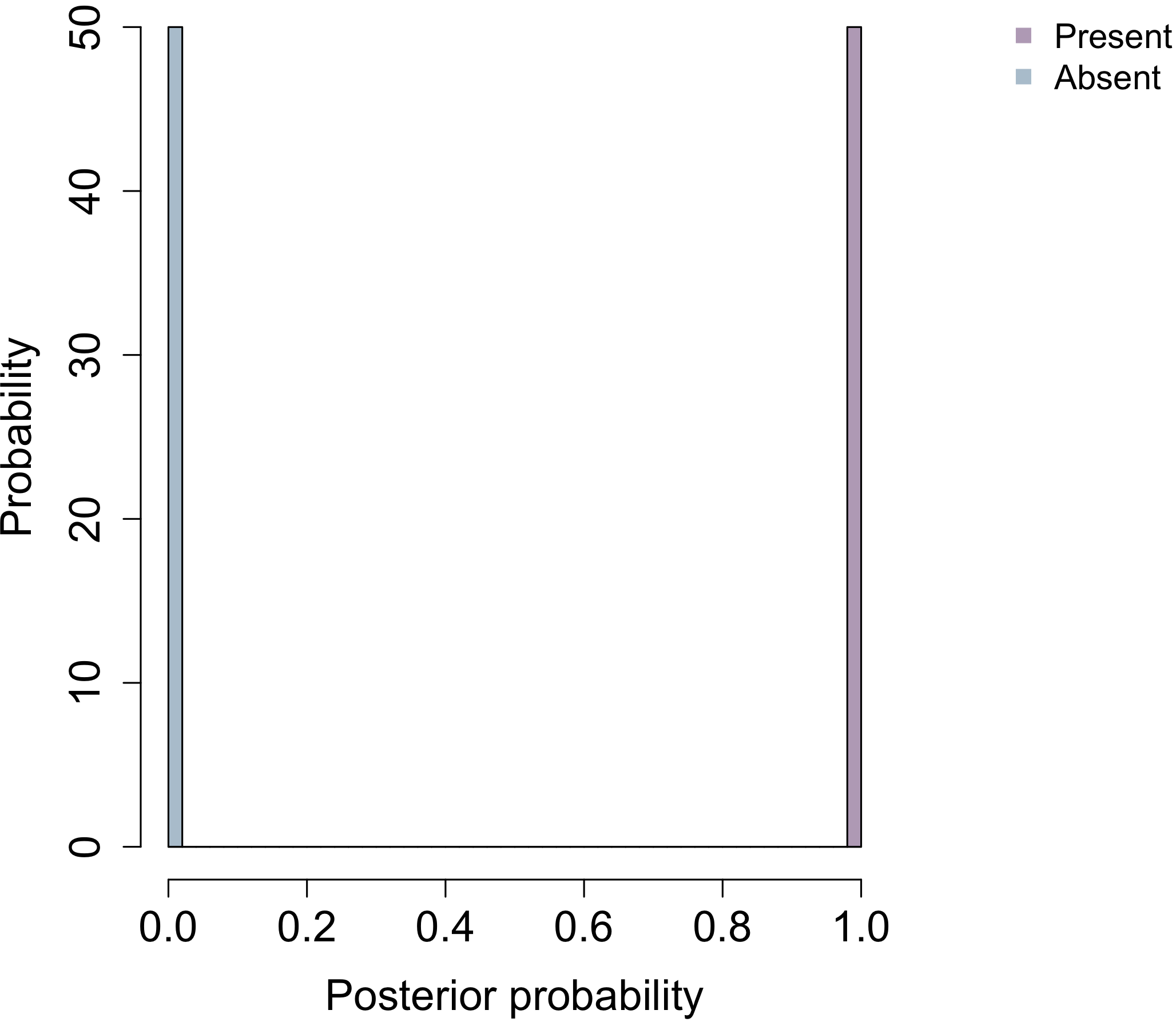

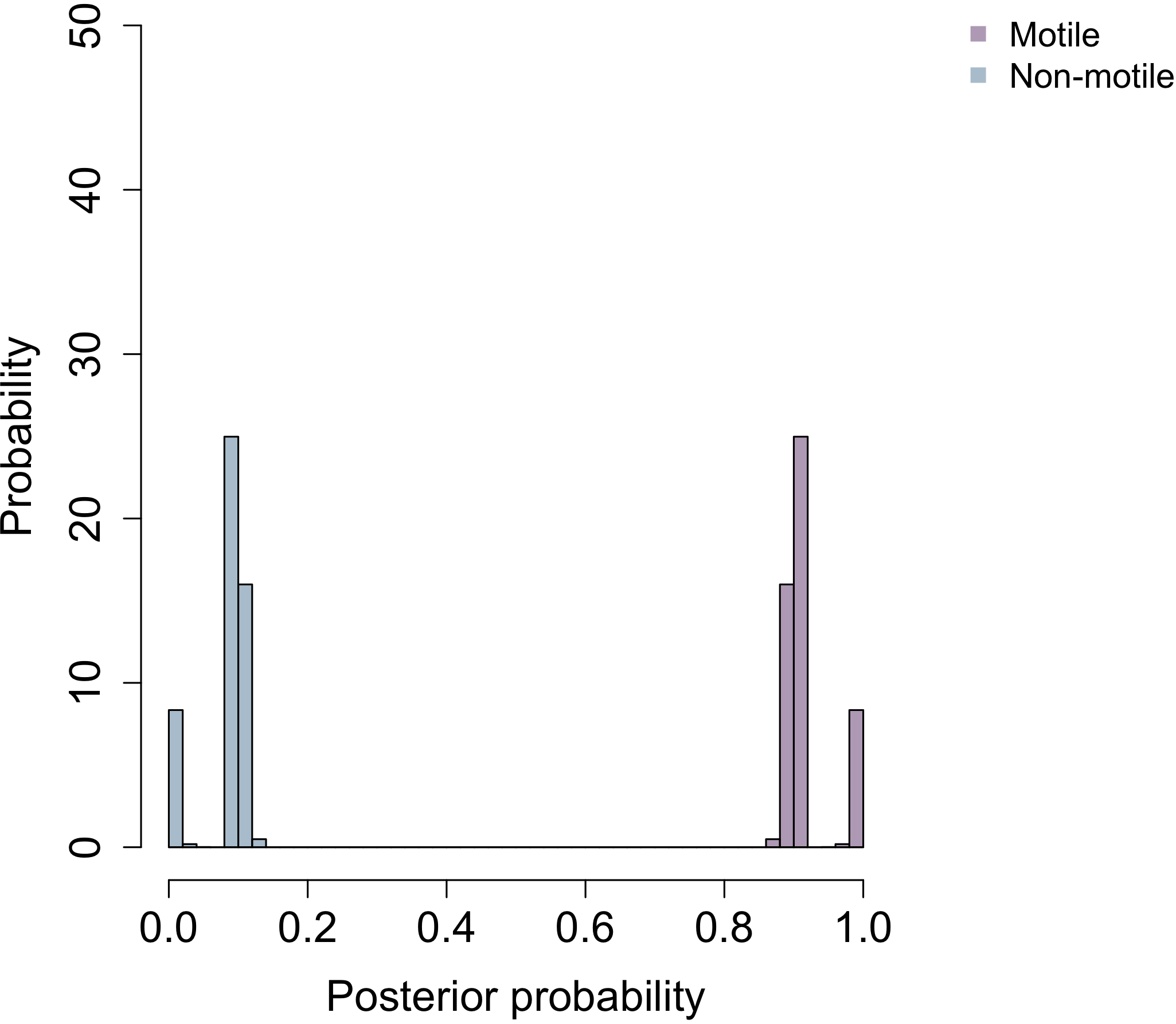

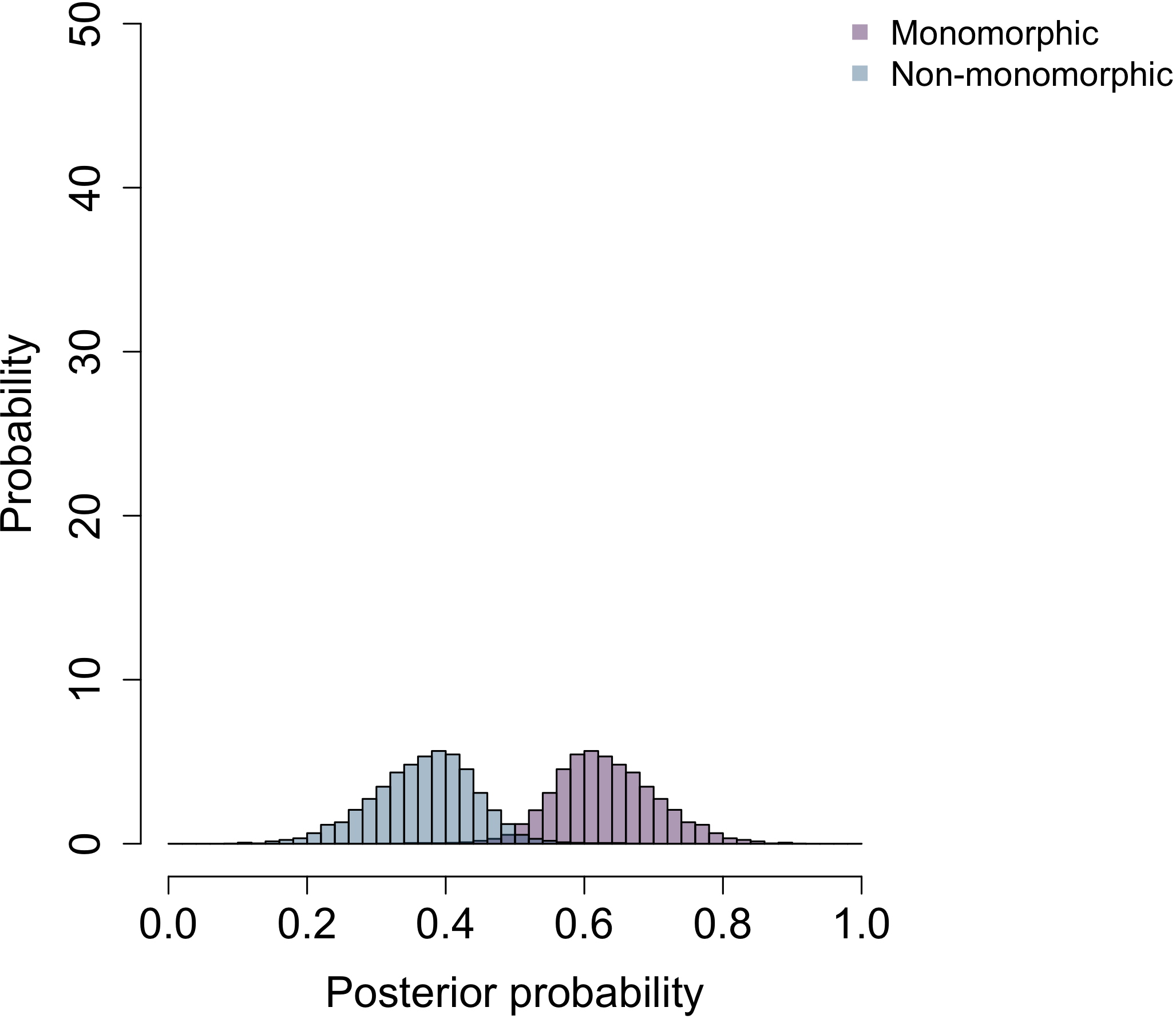

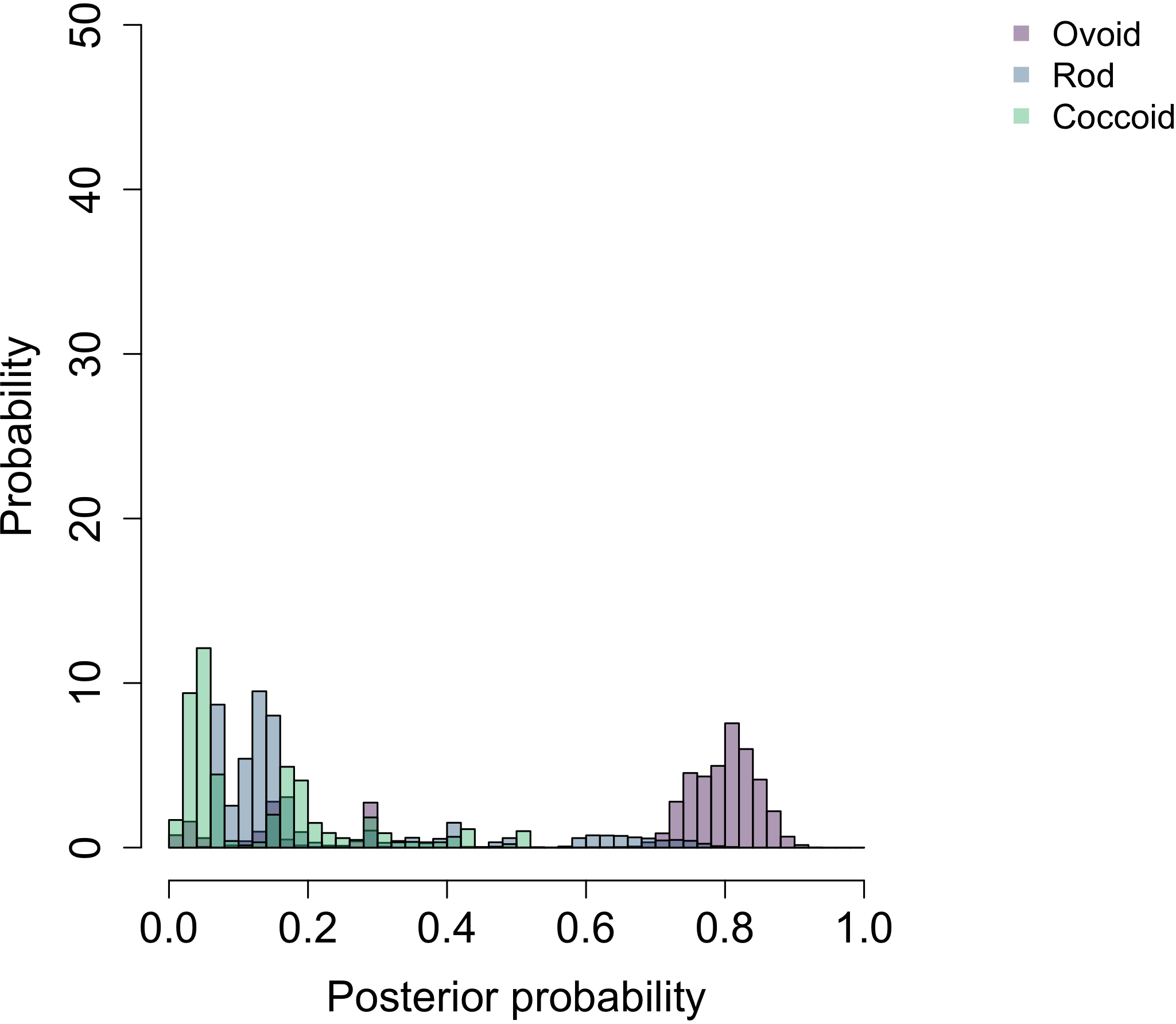

**Cell wall**

**Motile**

**Shape**

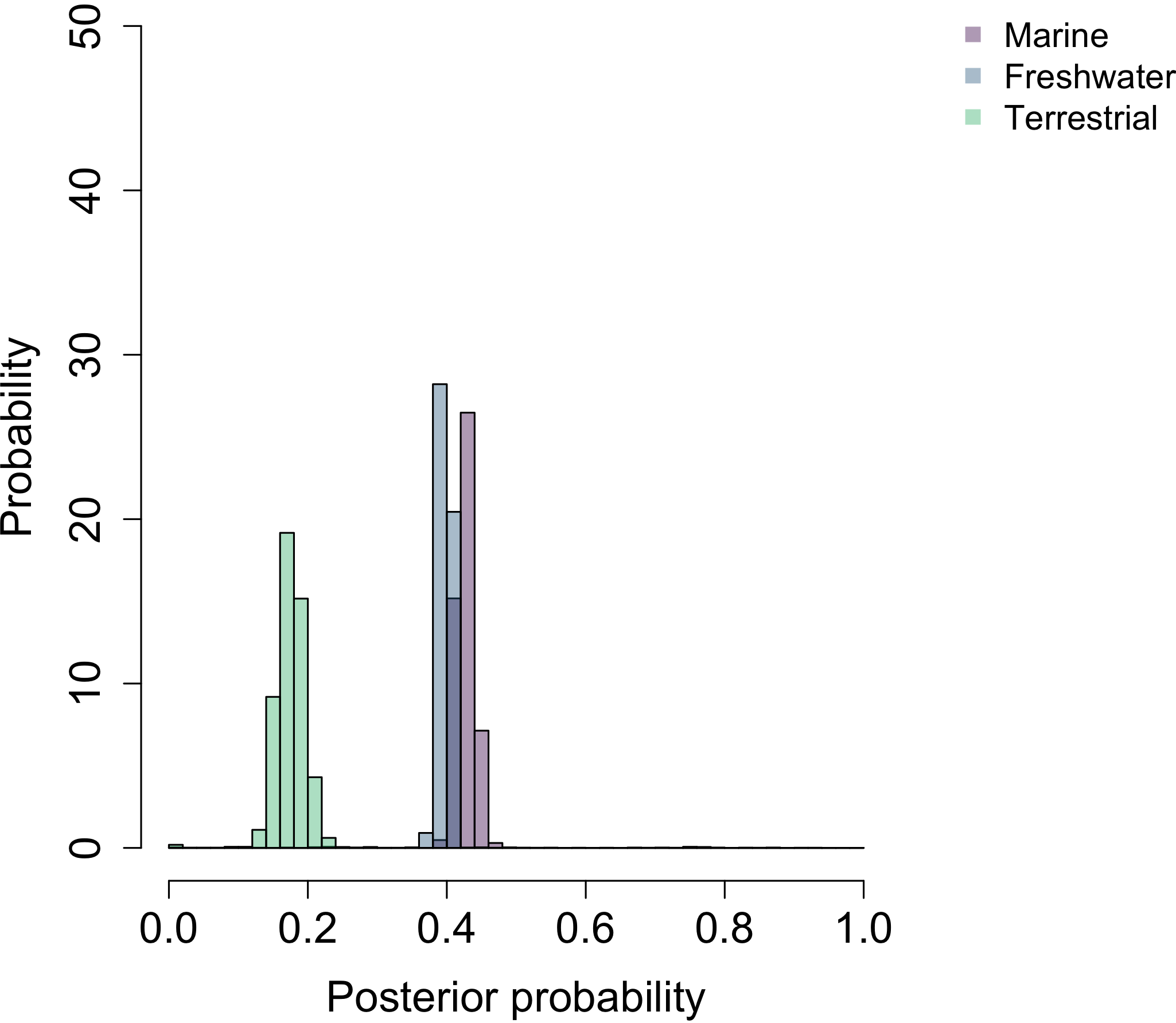

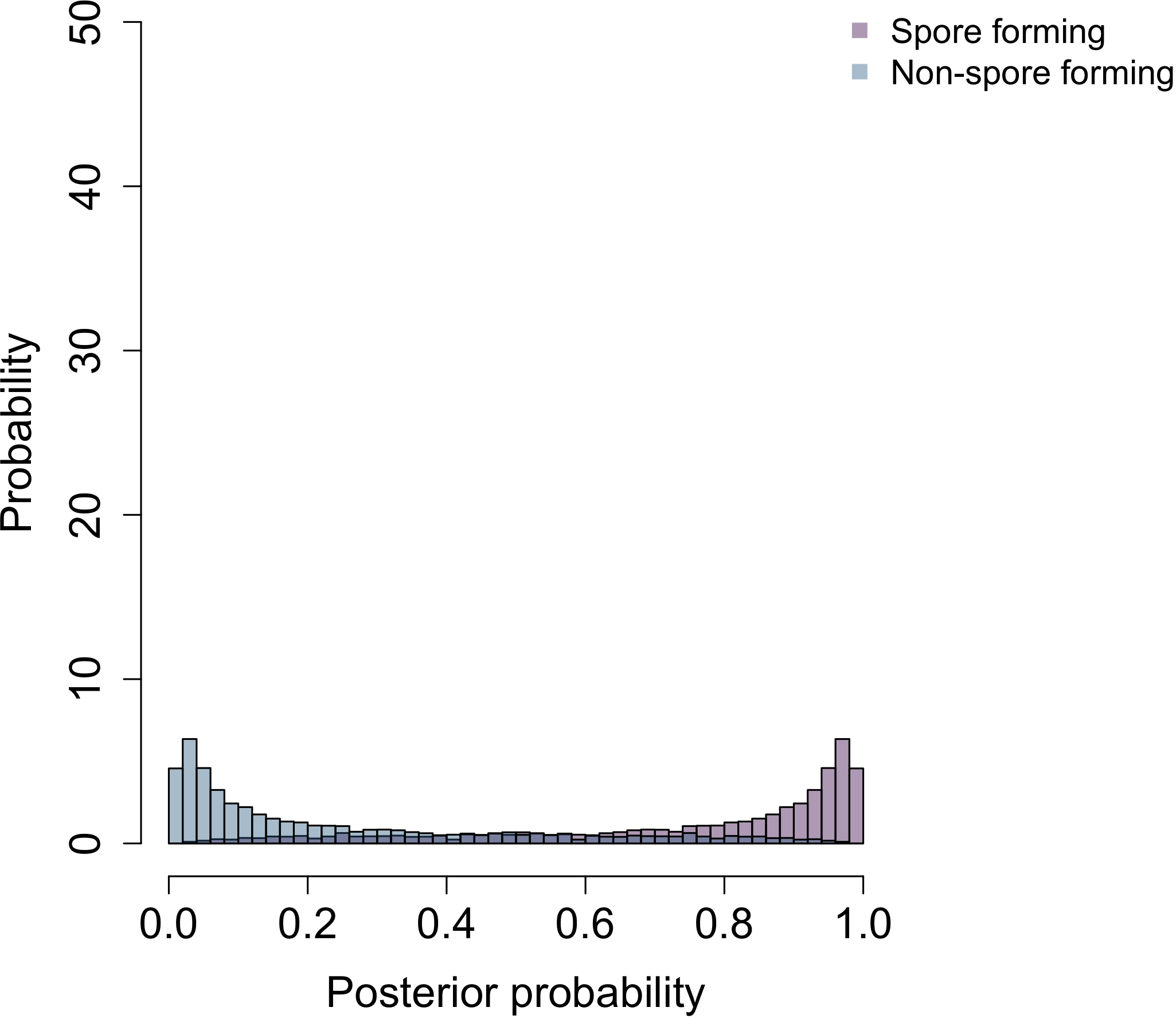

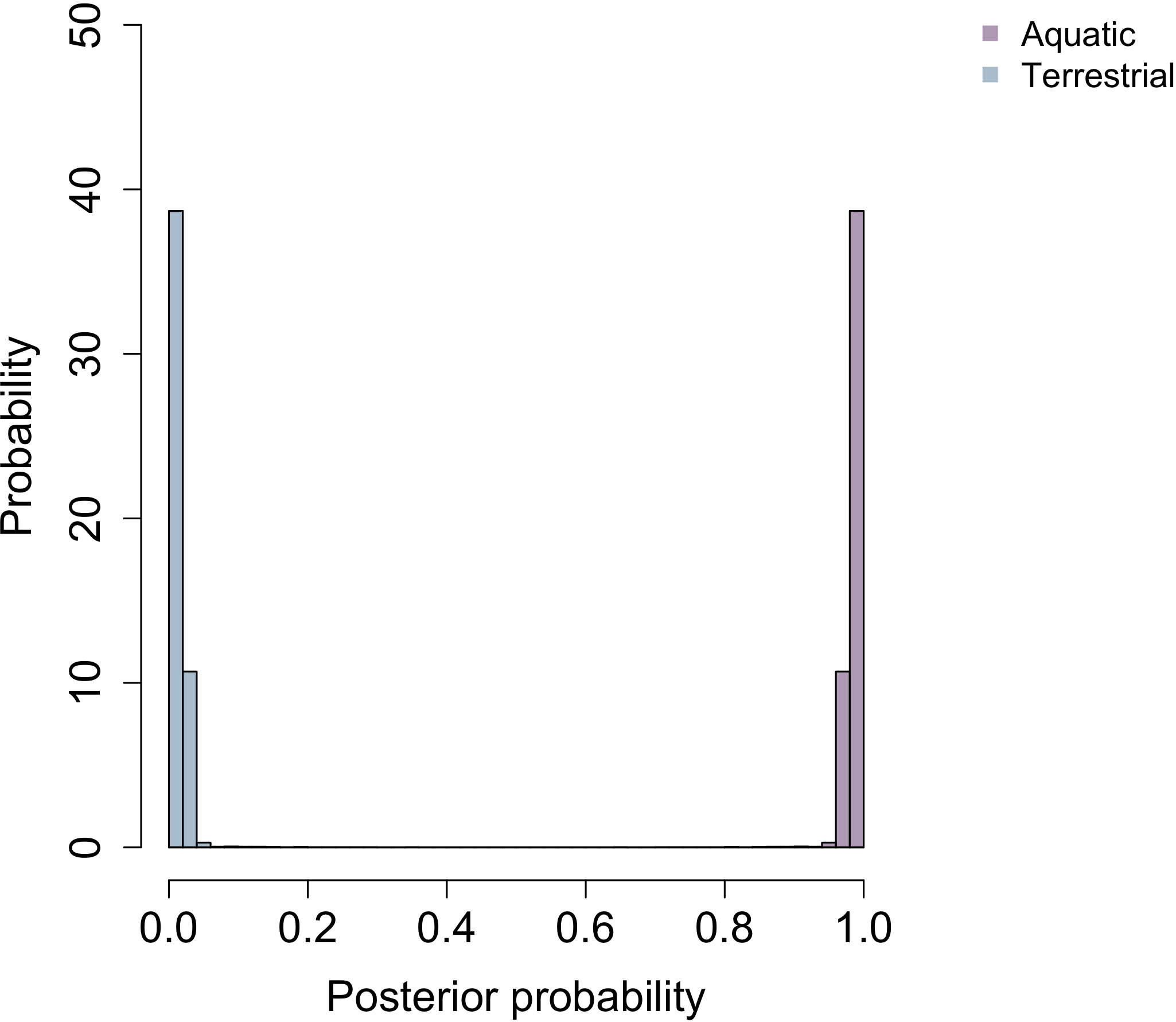

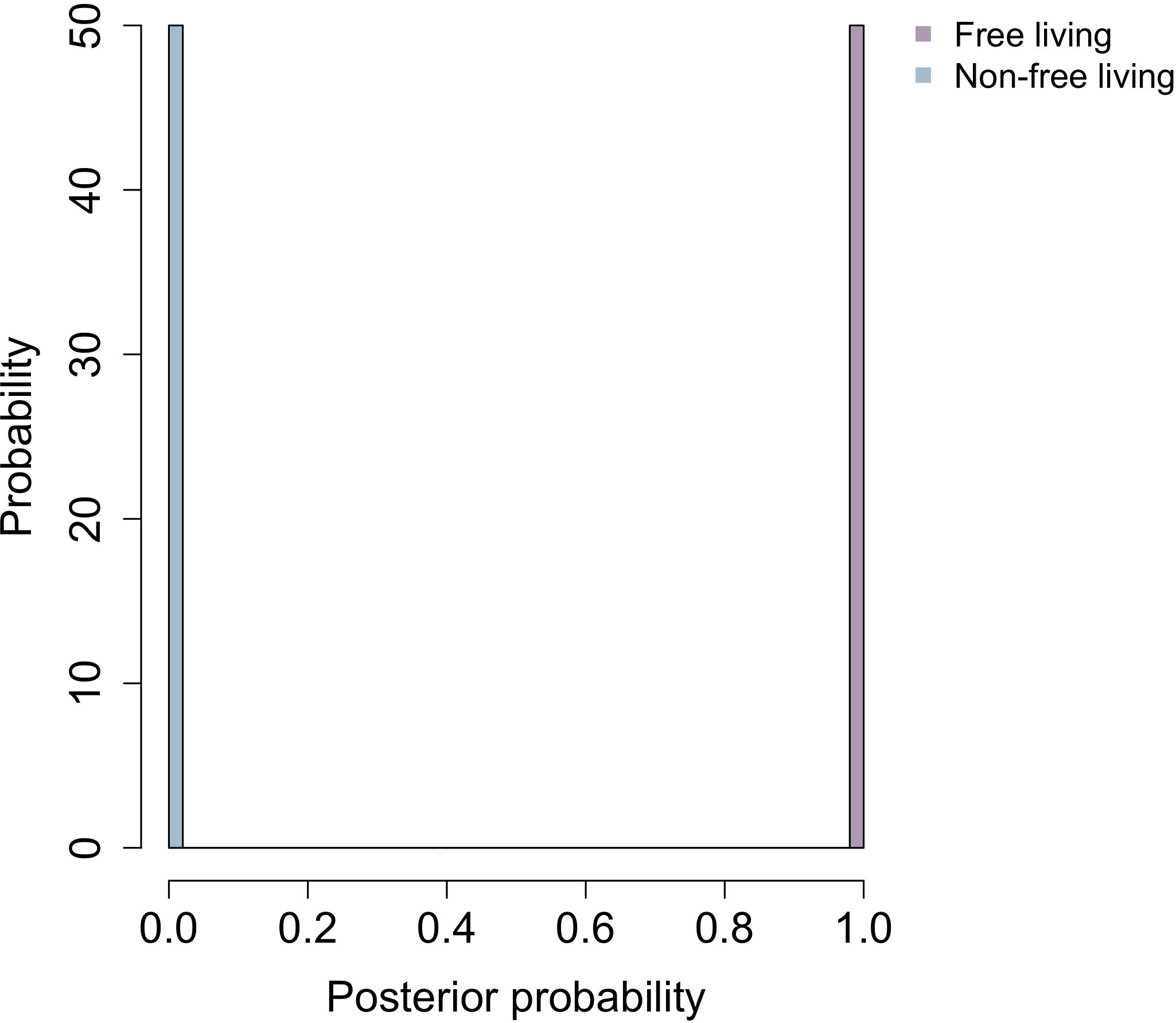

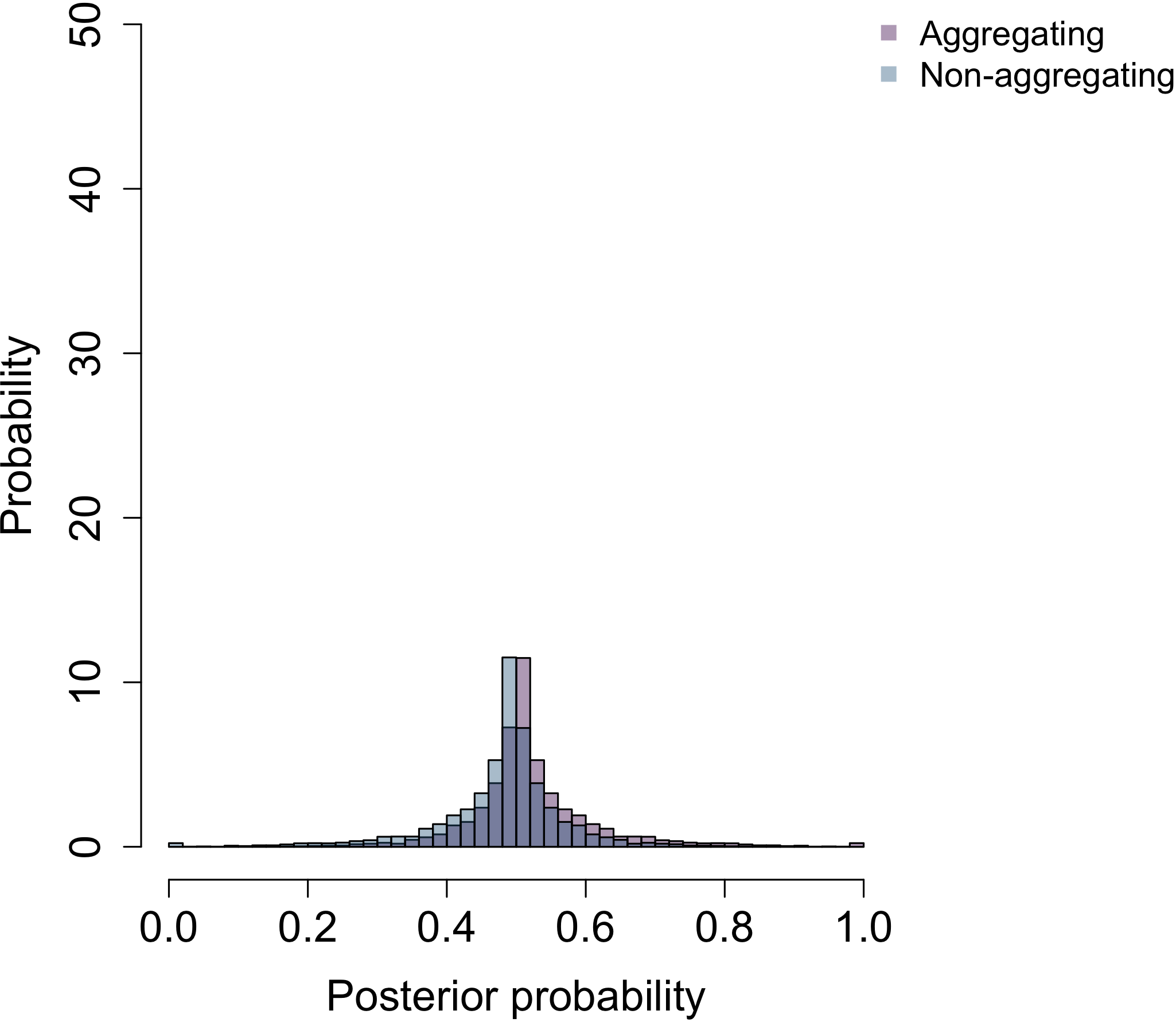

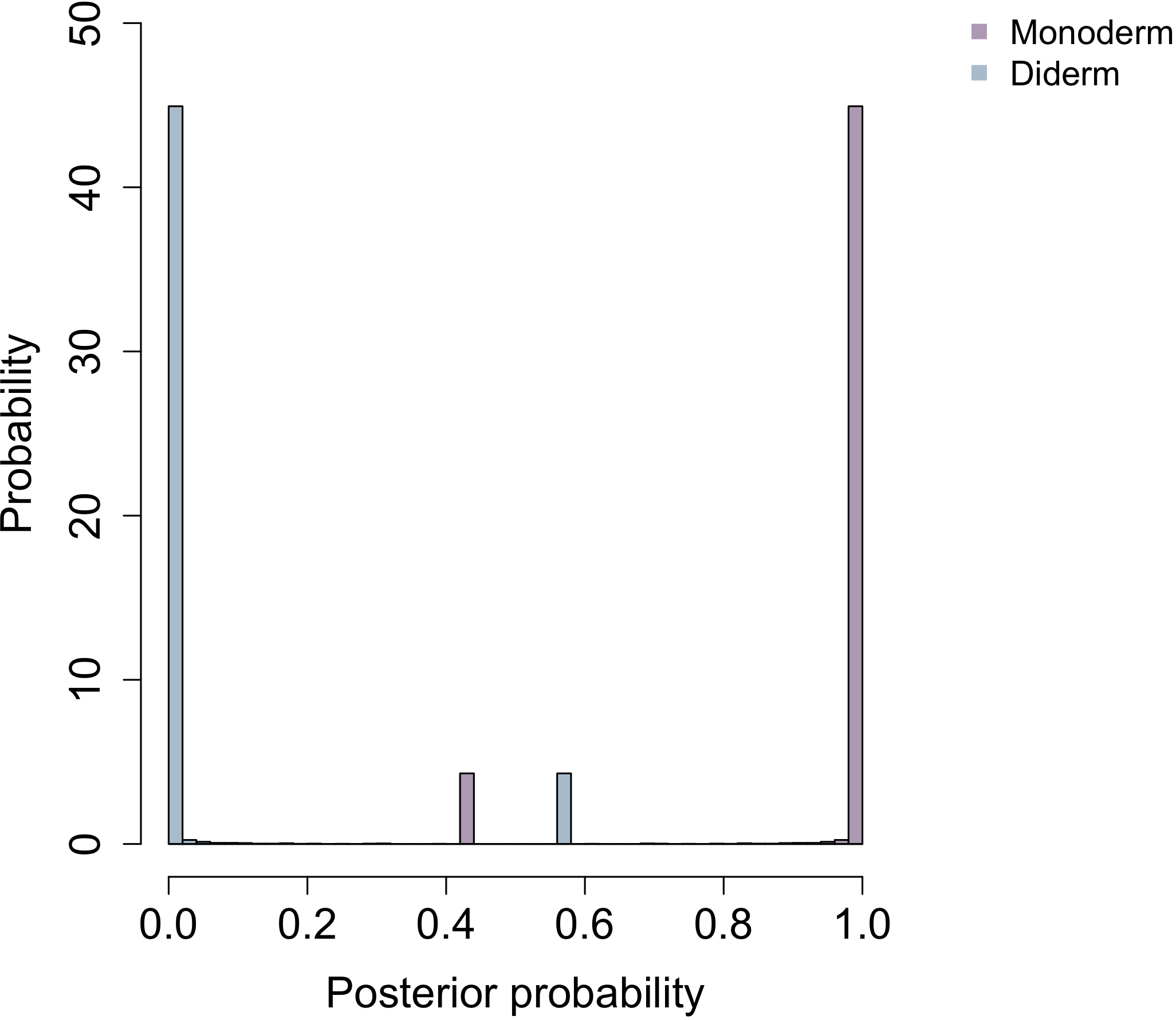

**Habitat c**

**Sporulation**

**Habitat b**

**Habitat a**

**Cell aggregation**

**Cell plan**

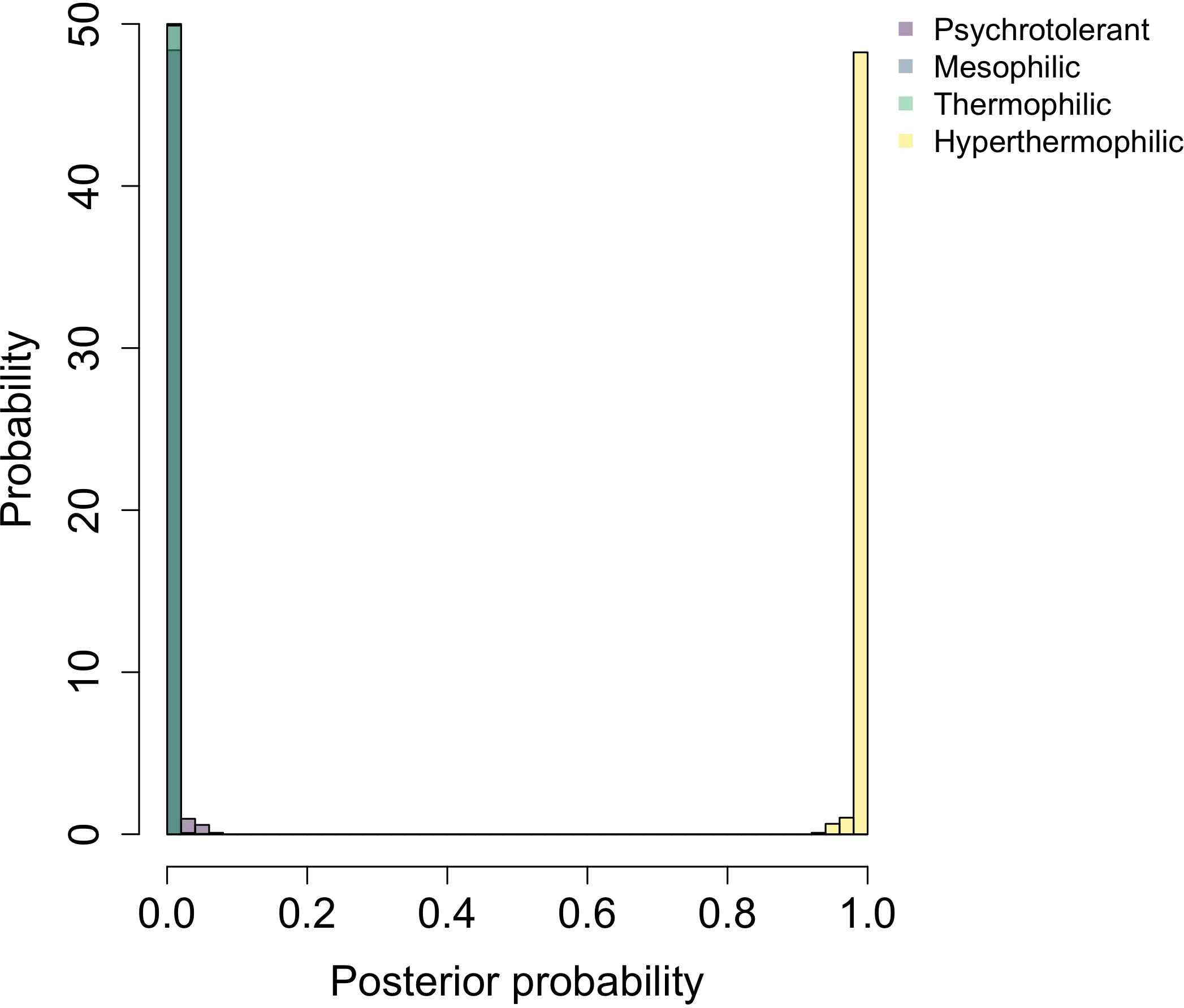

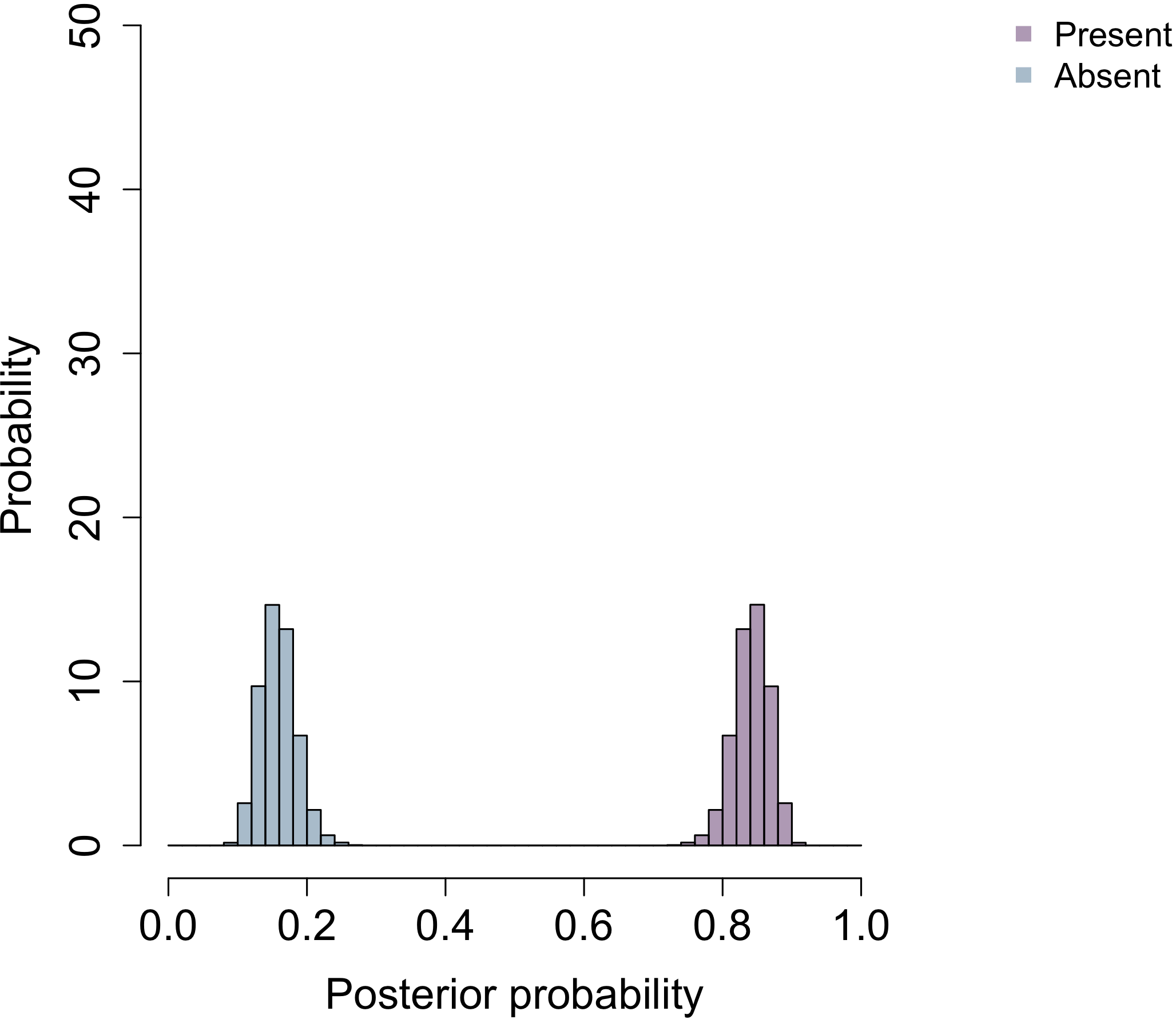

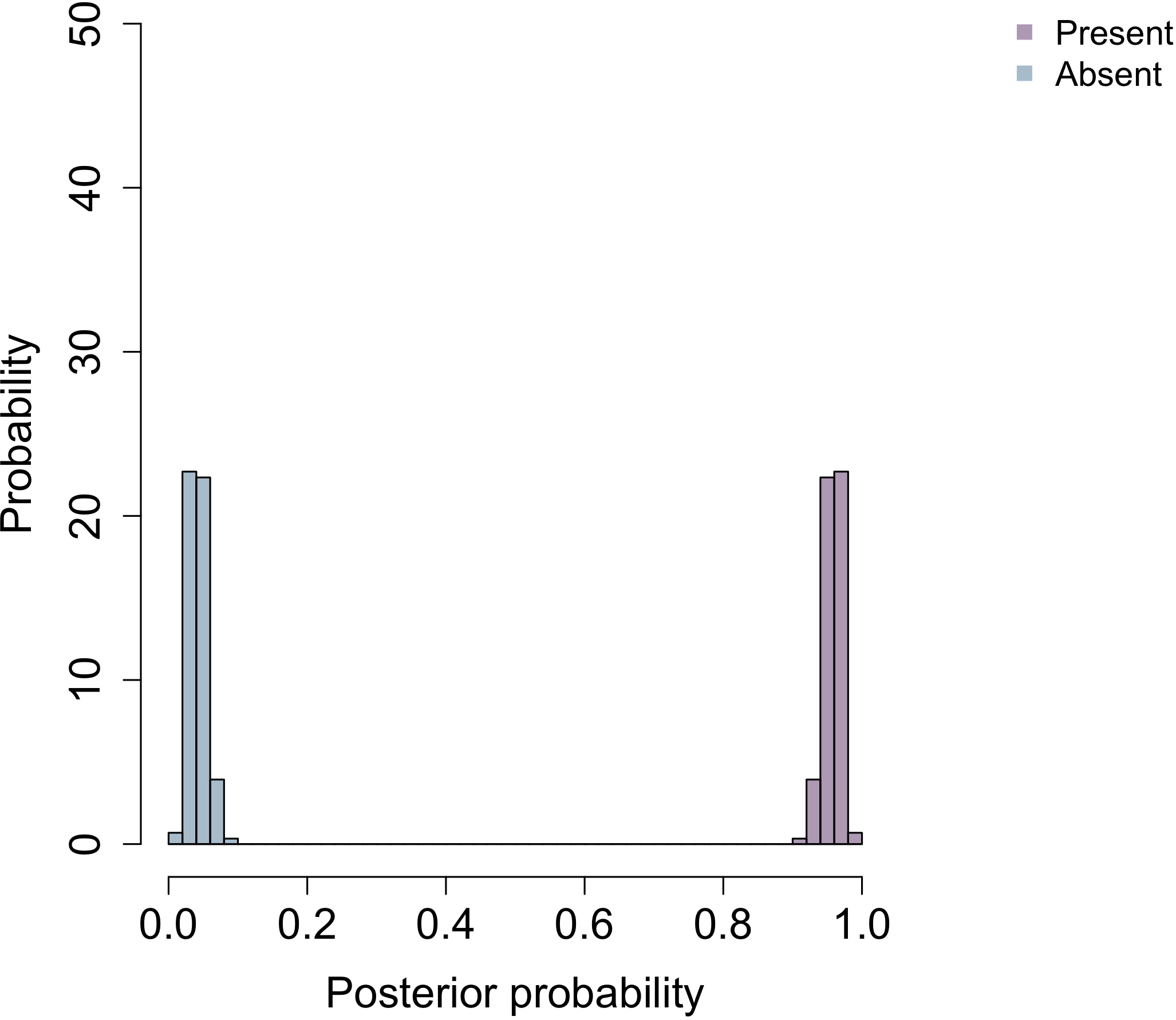

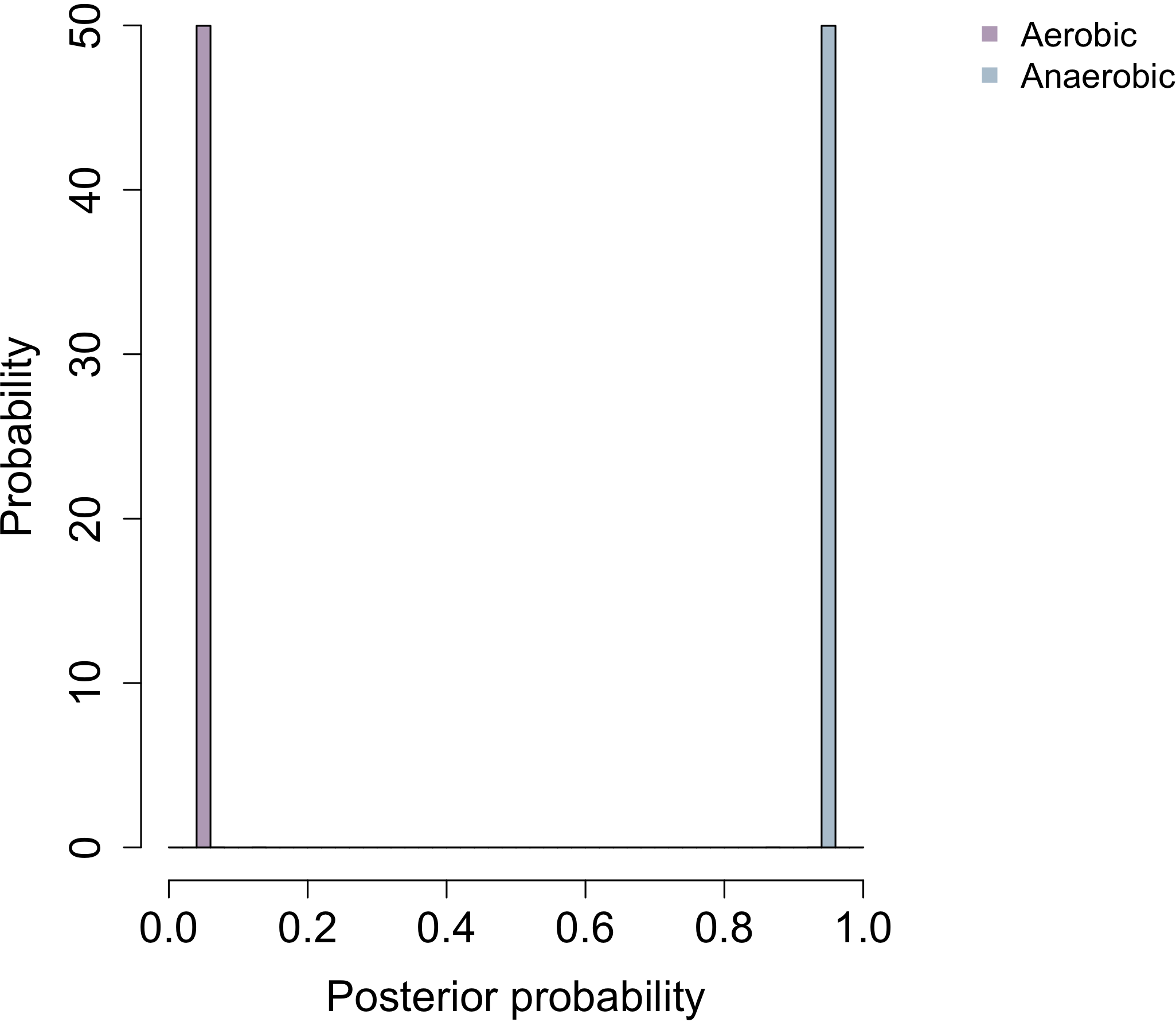

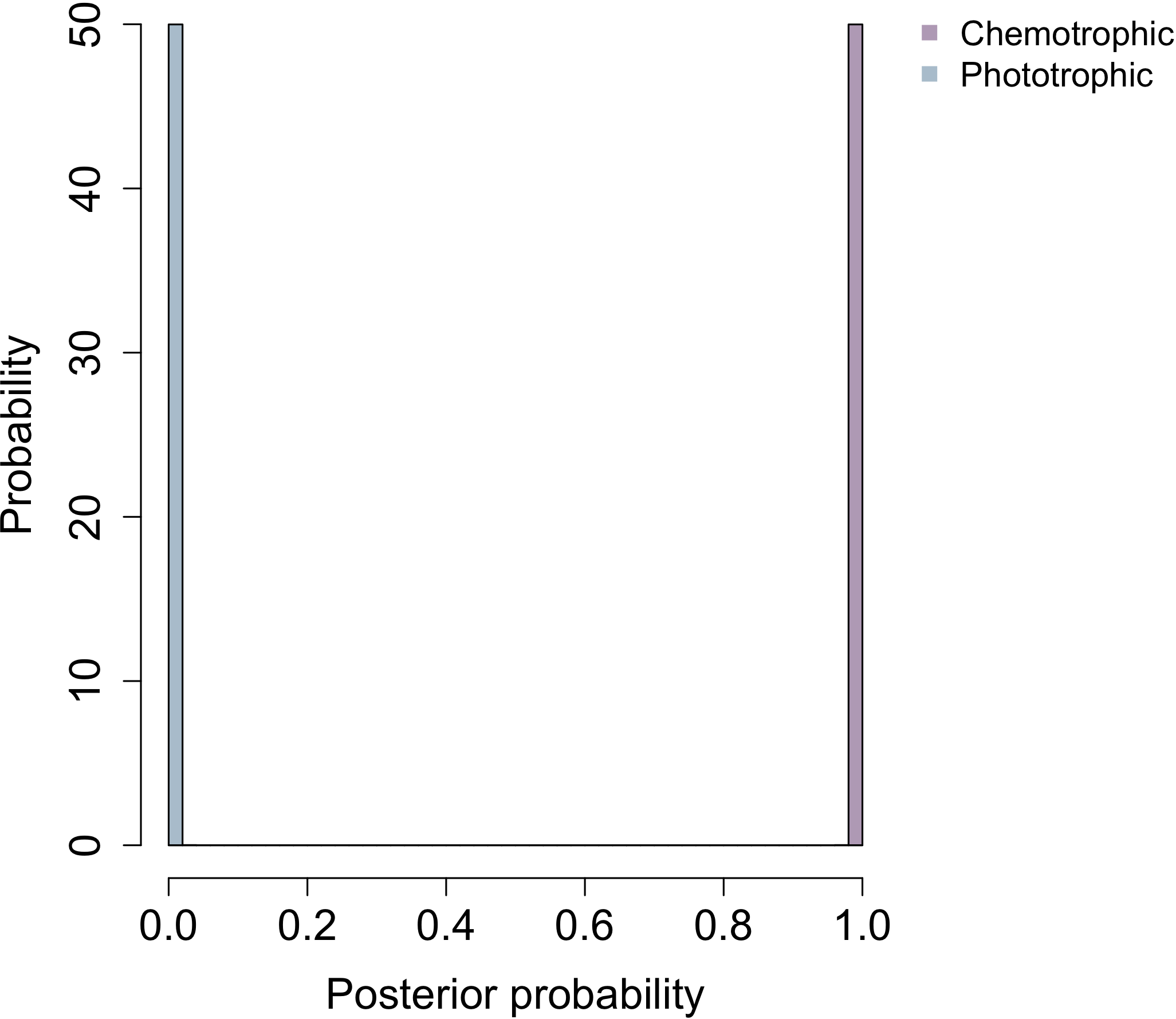

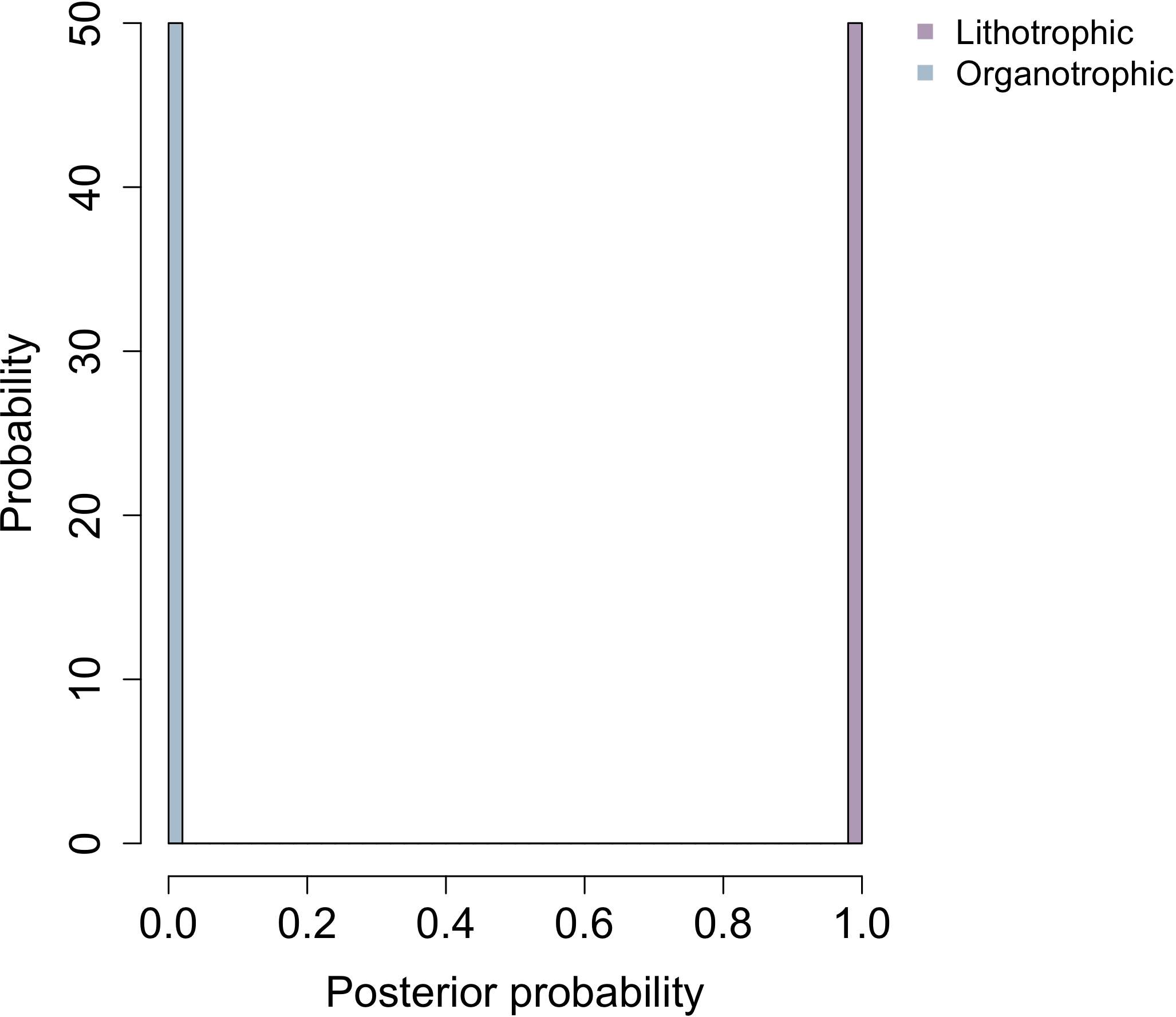

**Oxidase**

**Temperature**

**Catalase**

**Oxygen requirement**

**Electron source**

**Electron donor**

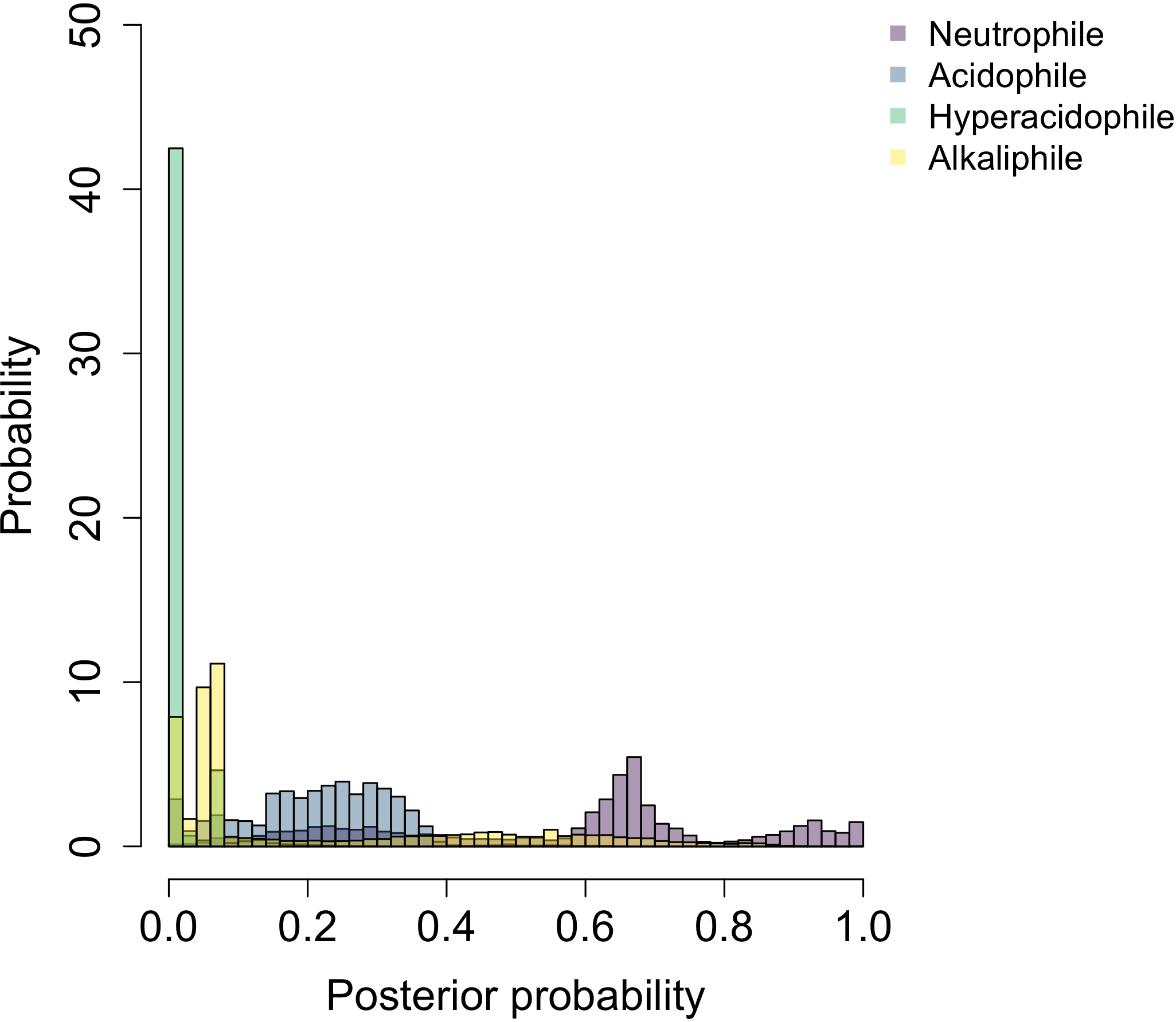

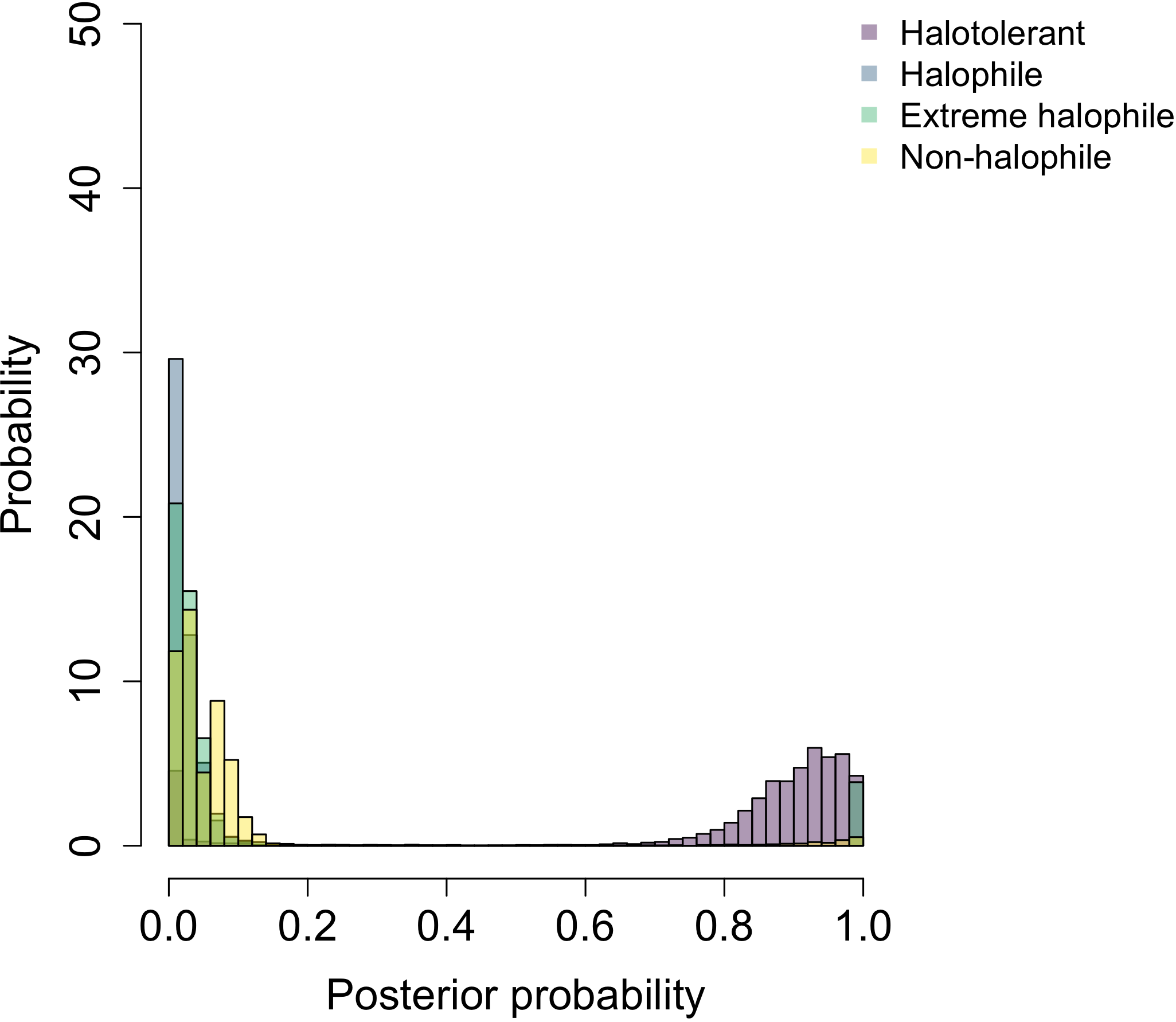

**pH**

**NaCl**

**b) LBCA**

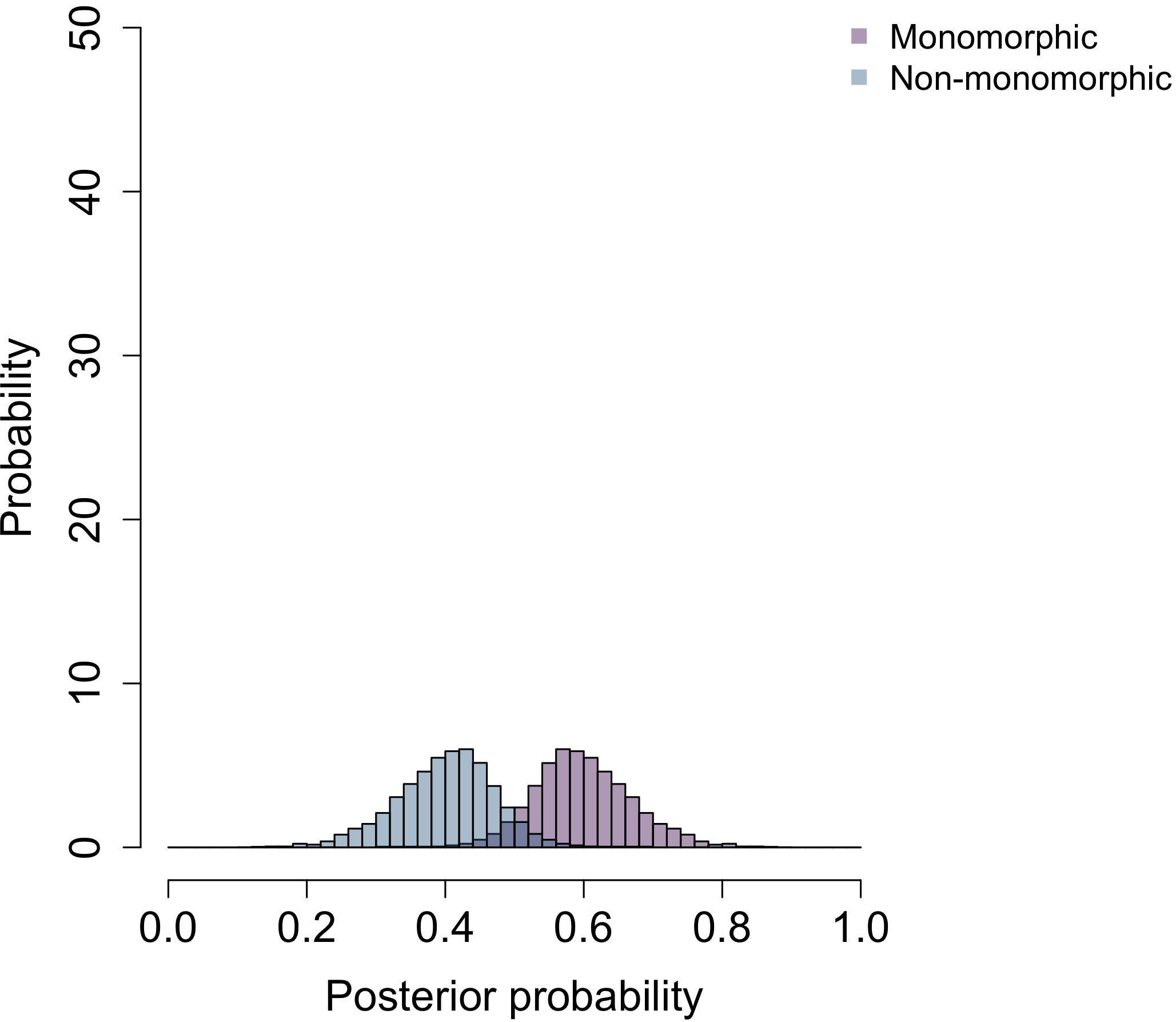

**Pleomorphism**

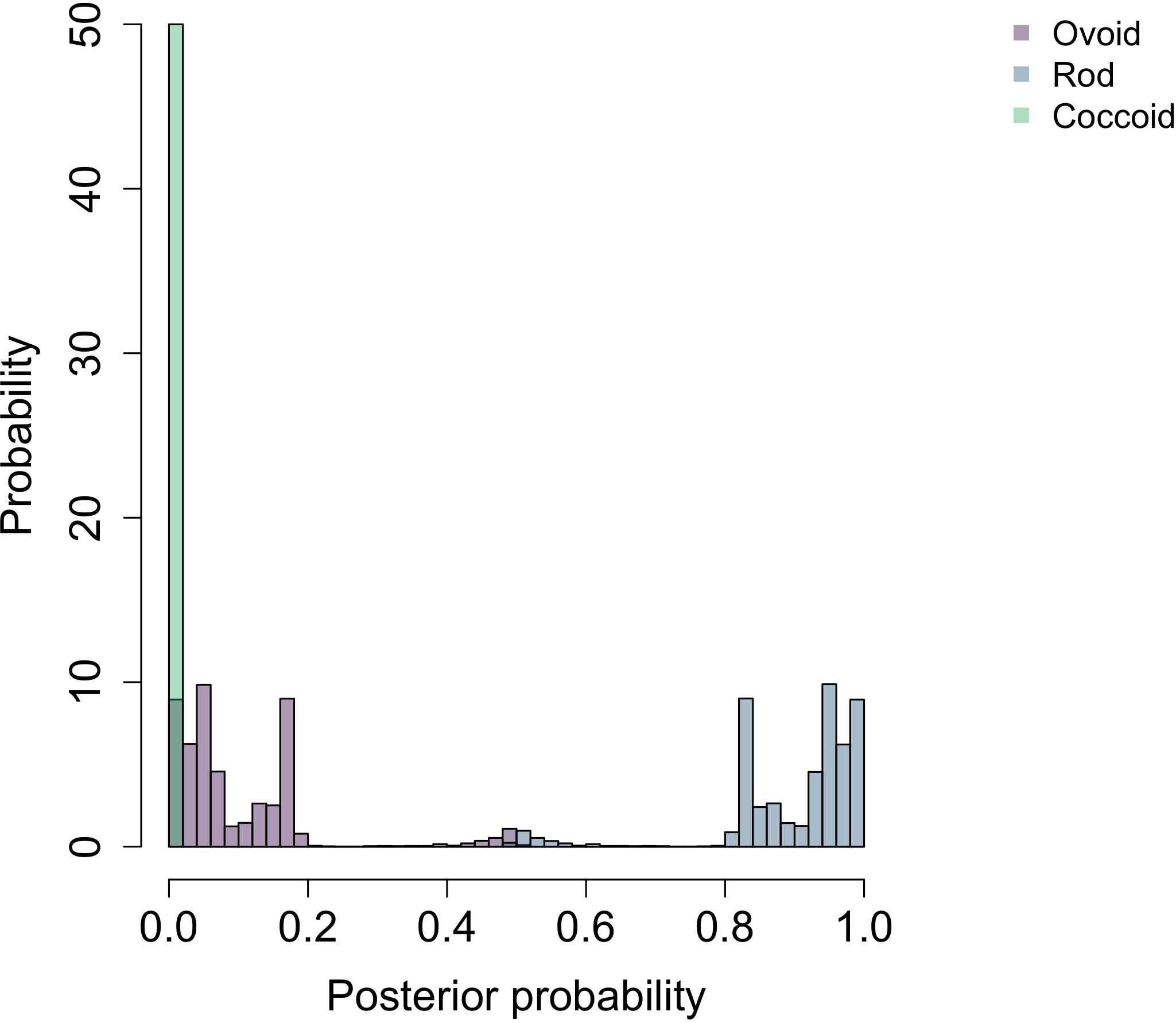

**Shape**

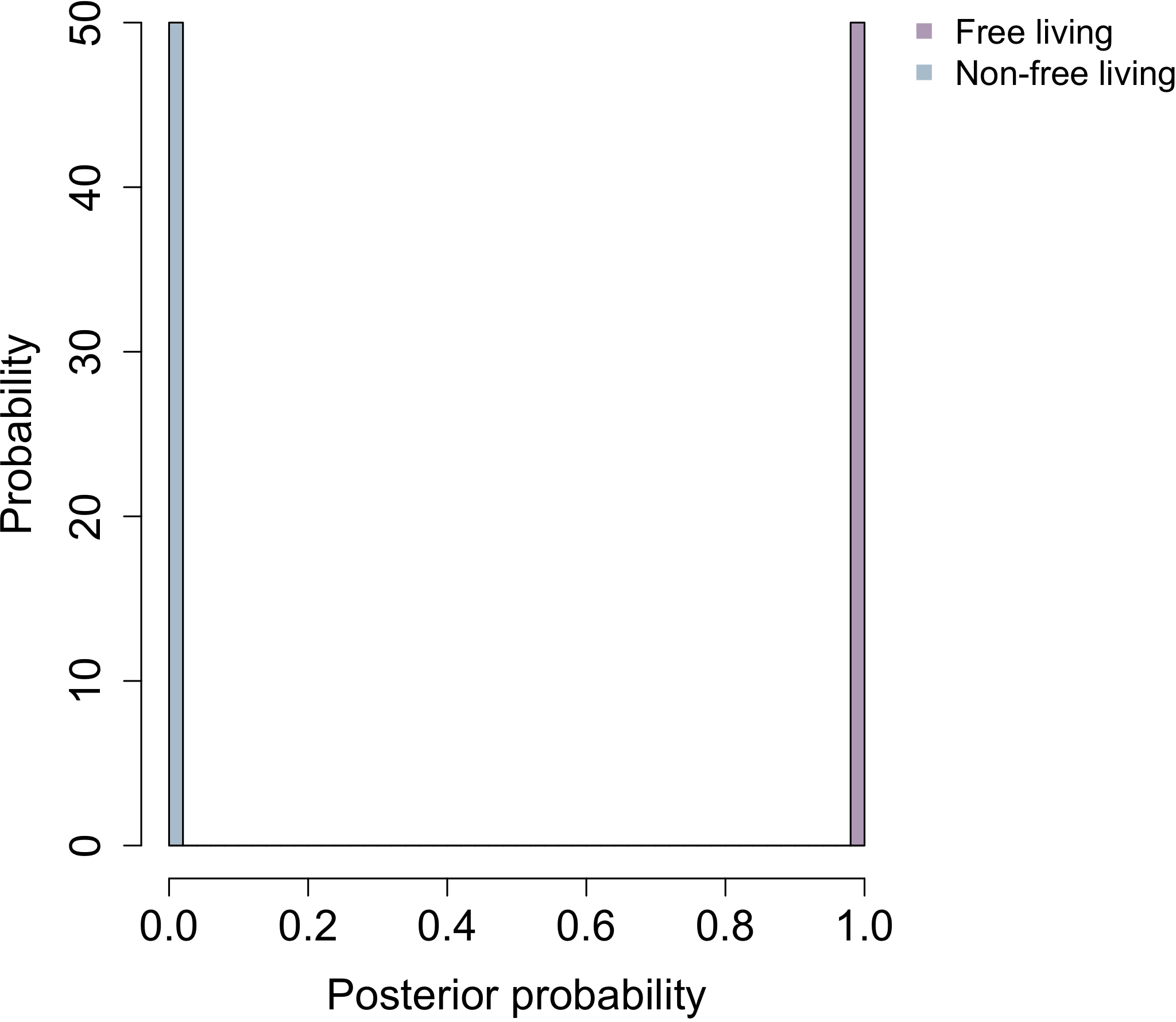

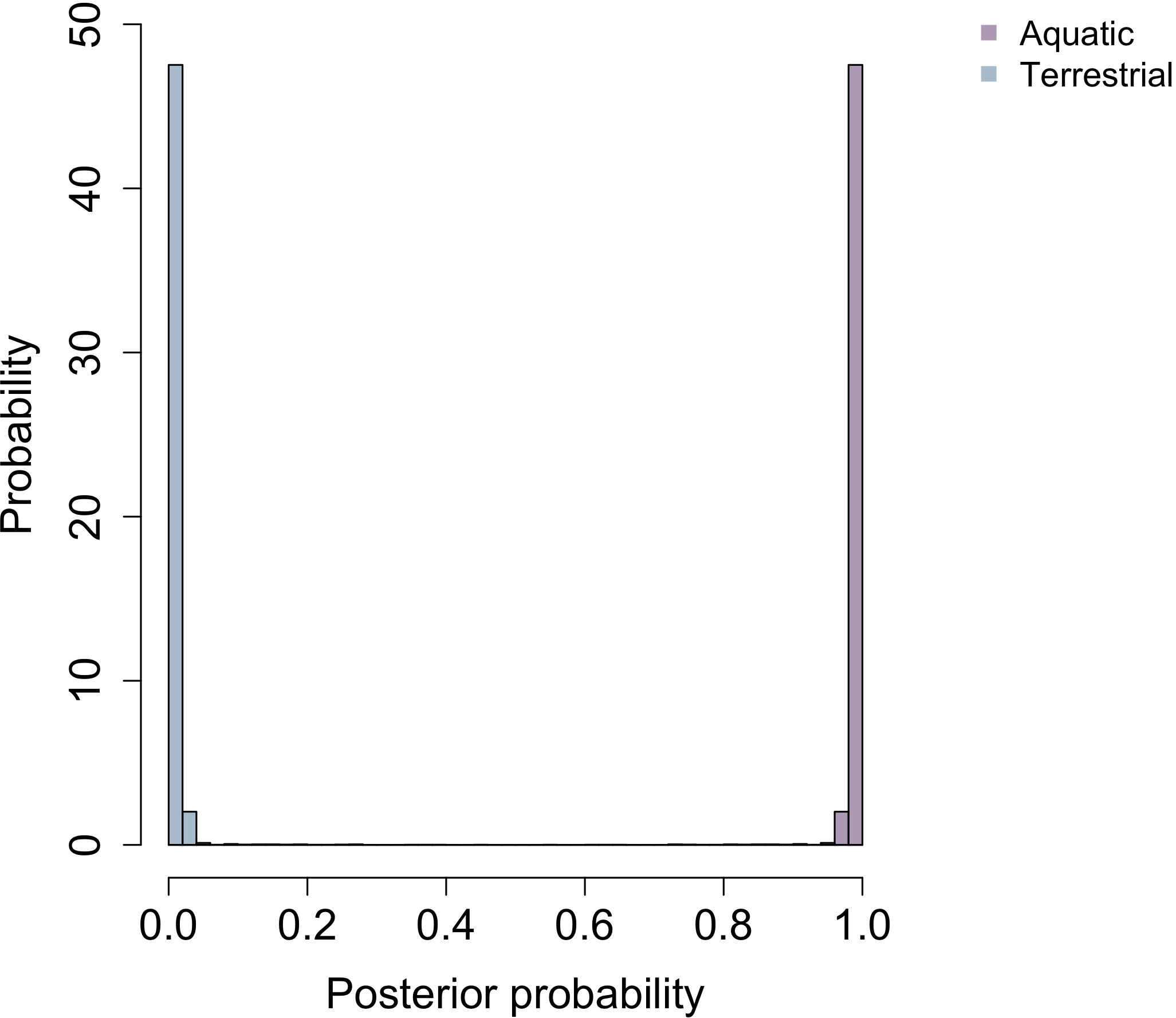

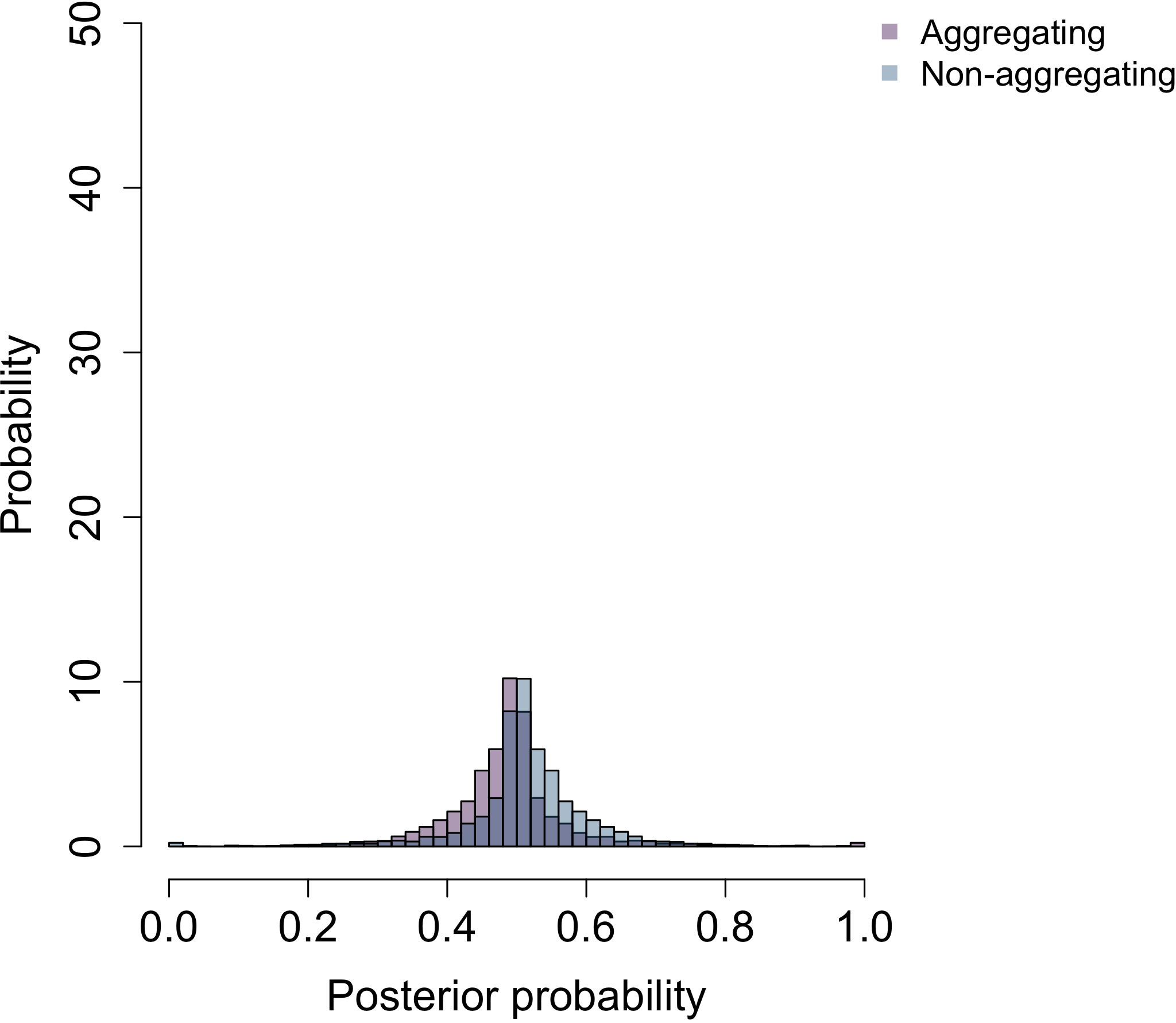

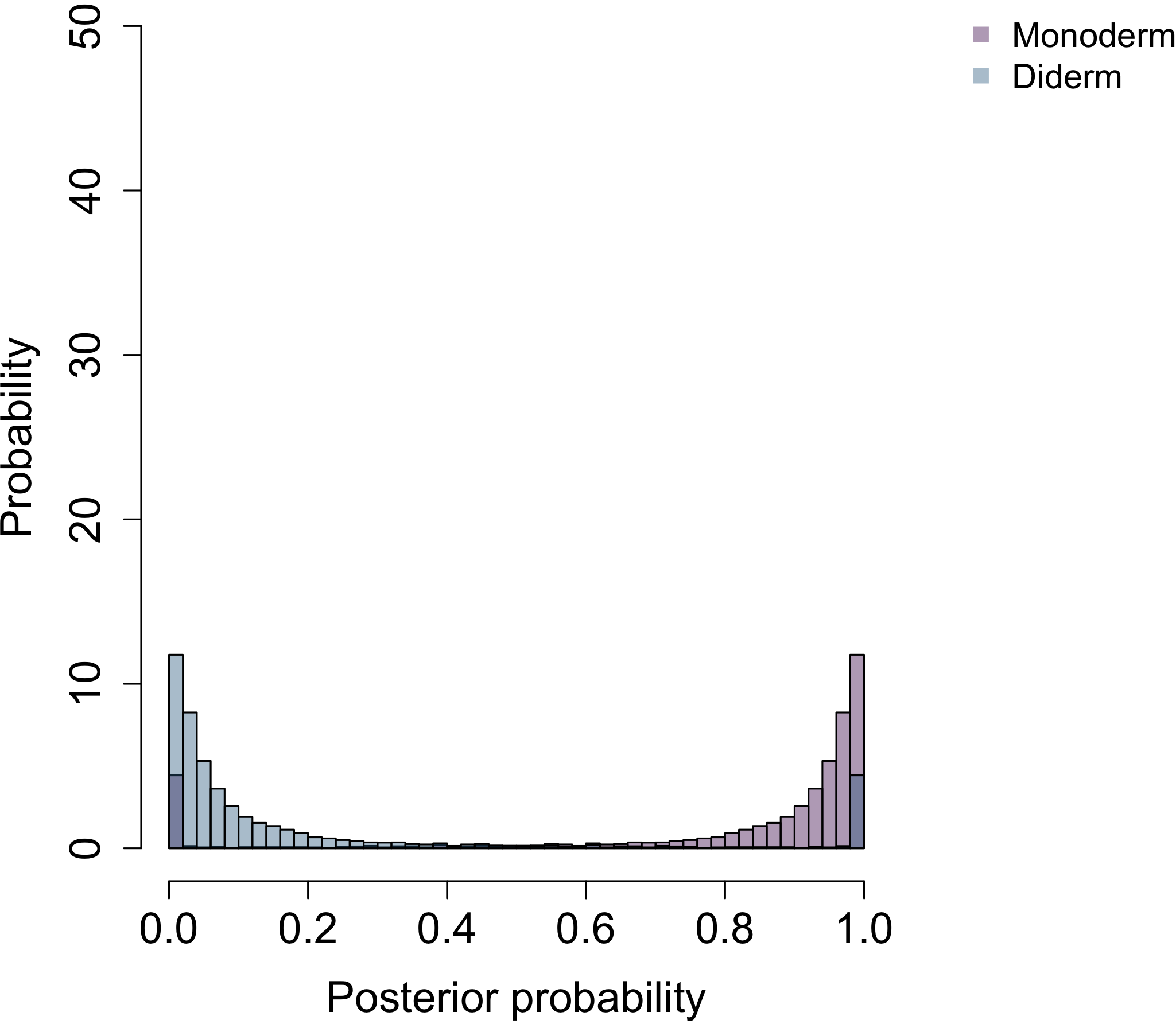

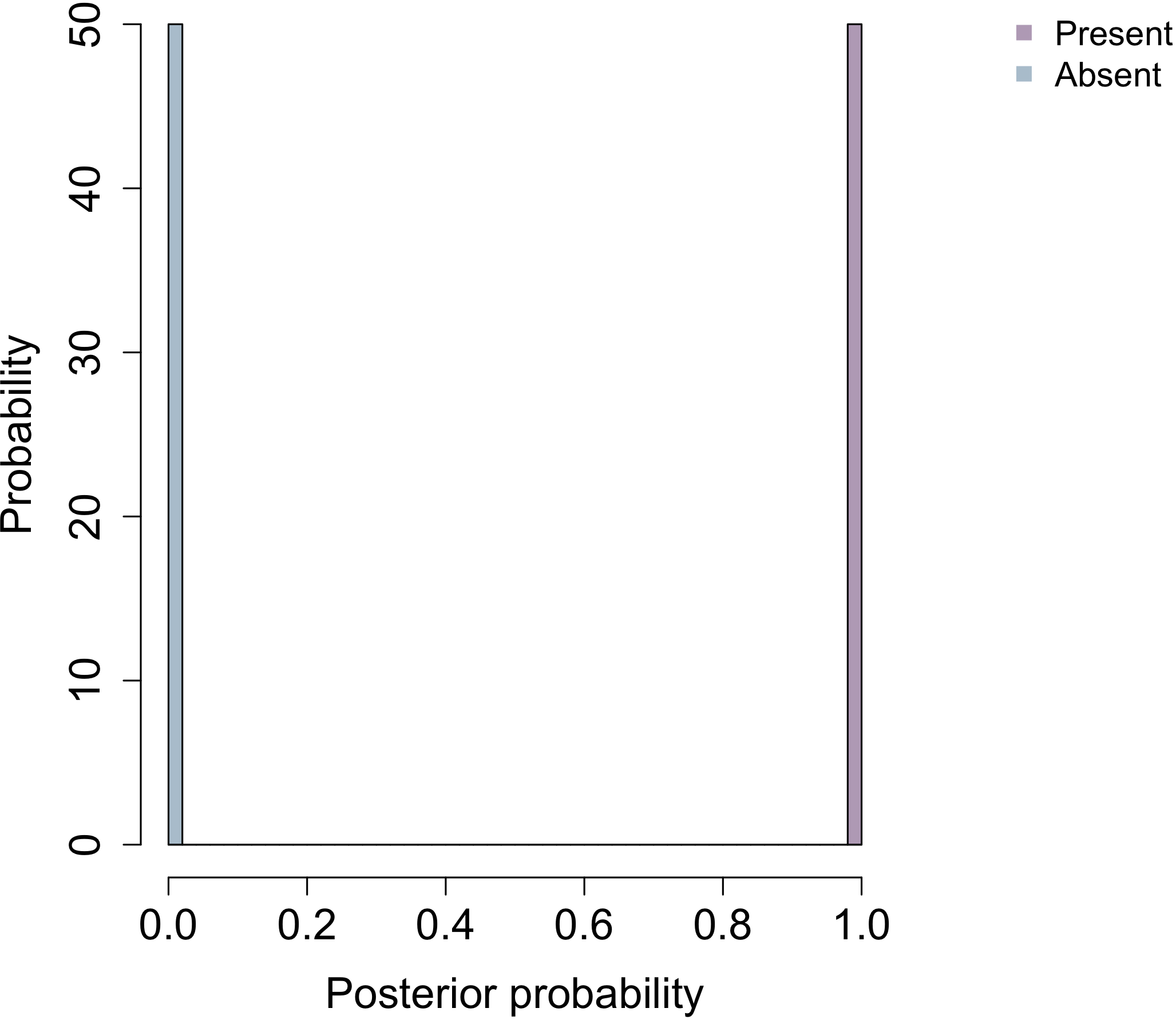

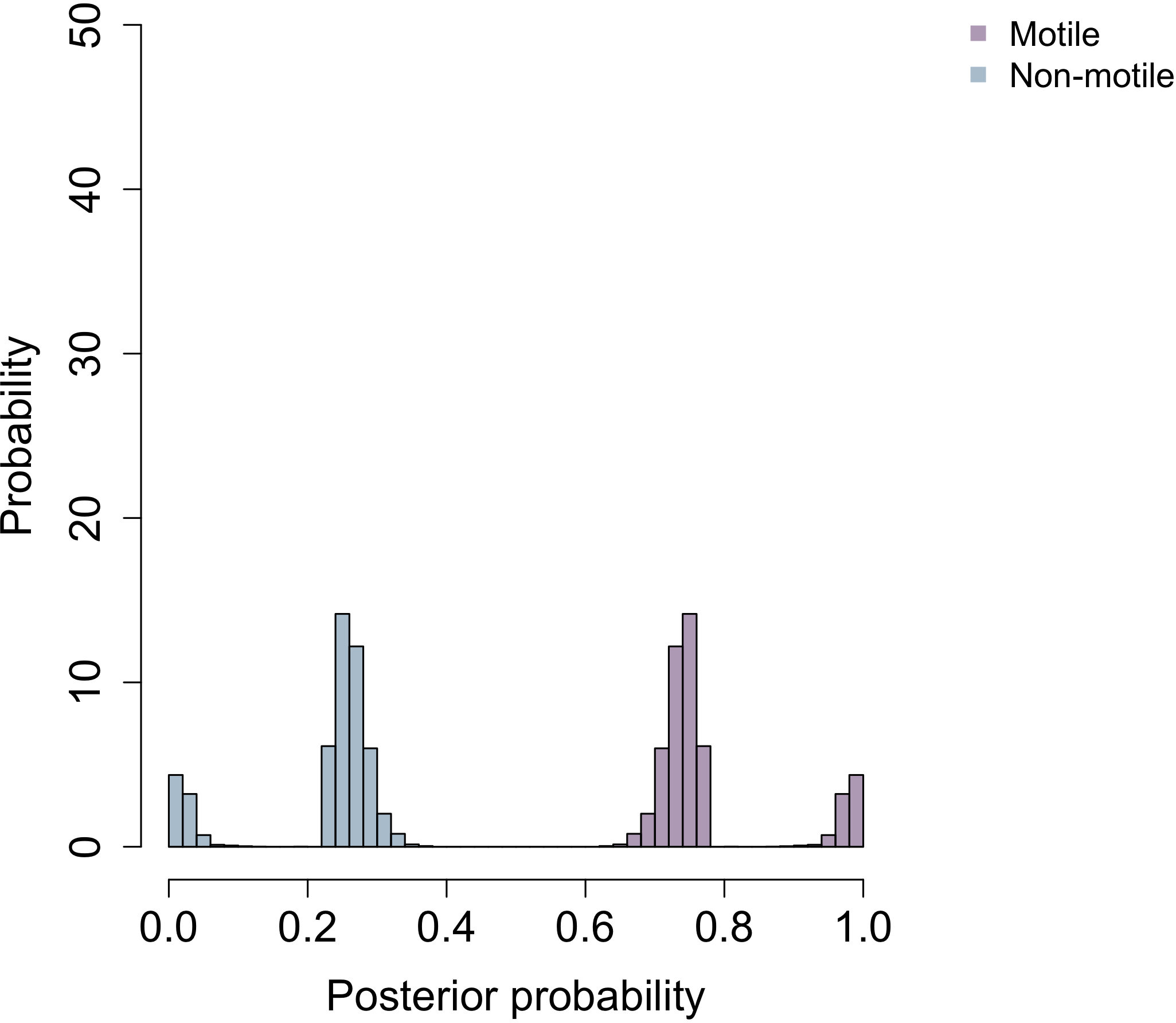

**Habitat b**

**Habitat a**

**Cell aggregation**

**Cell plan**

**Cell wall**

**Motile**

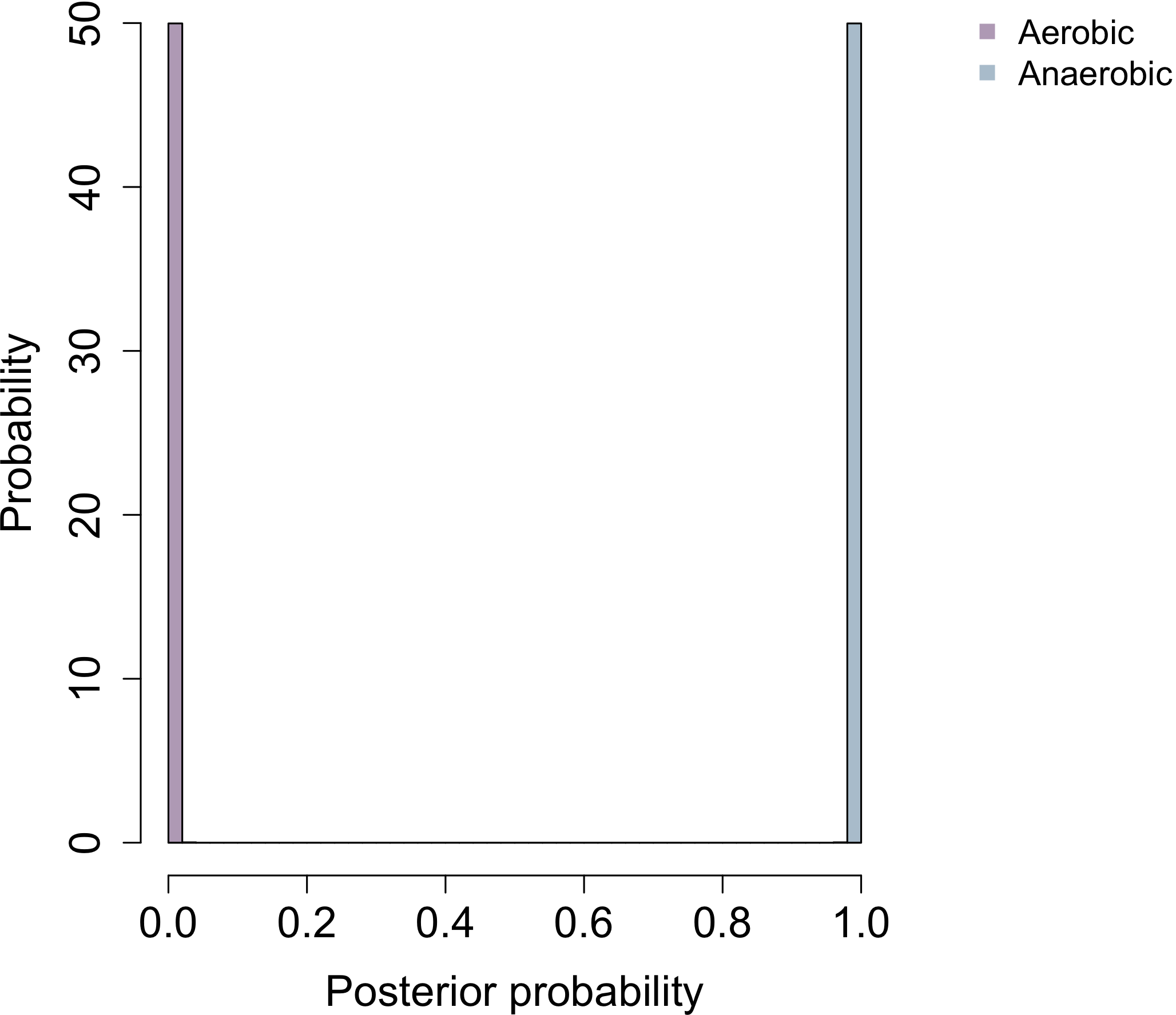

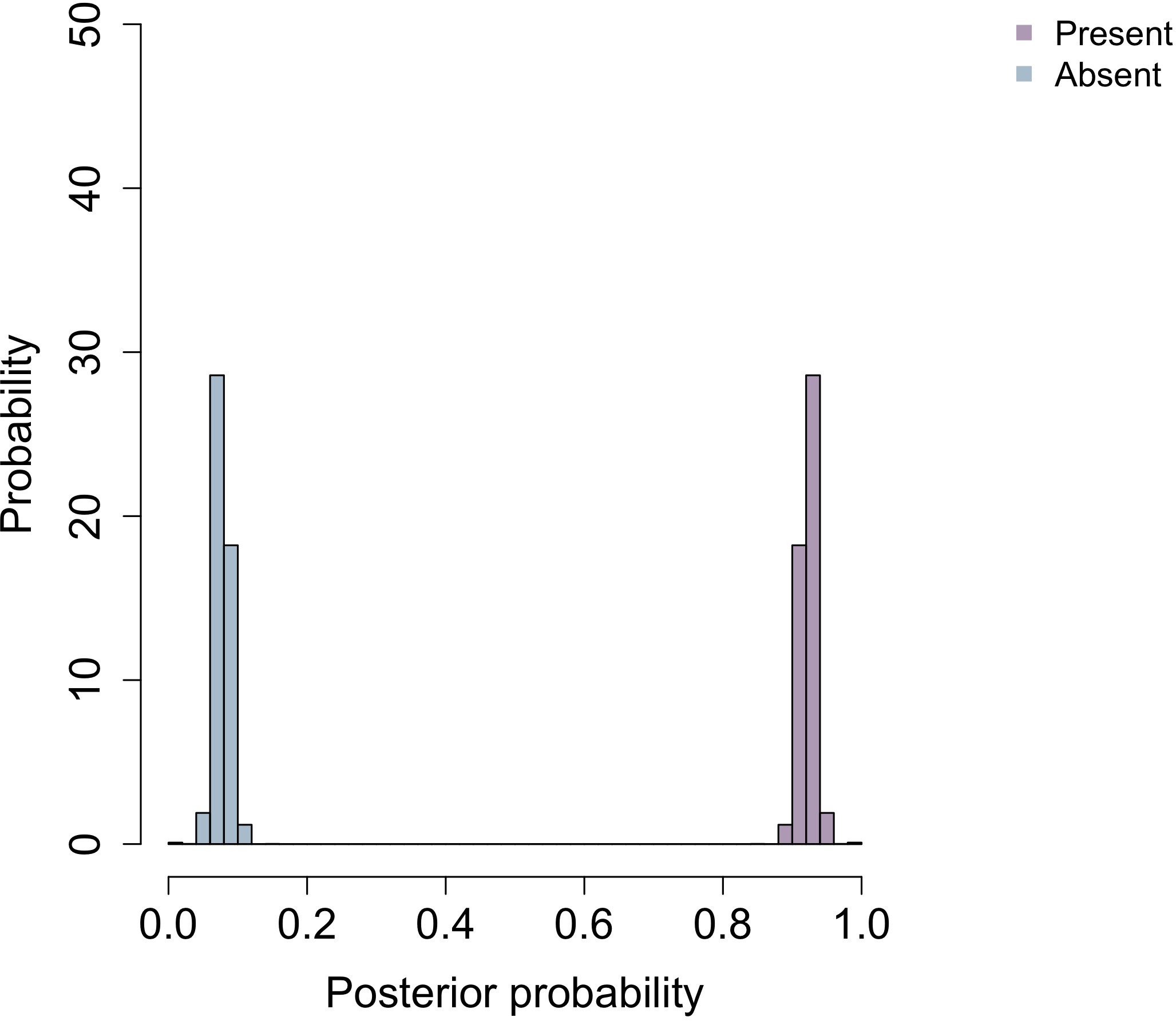

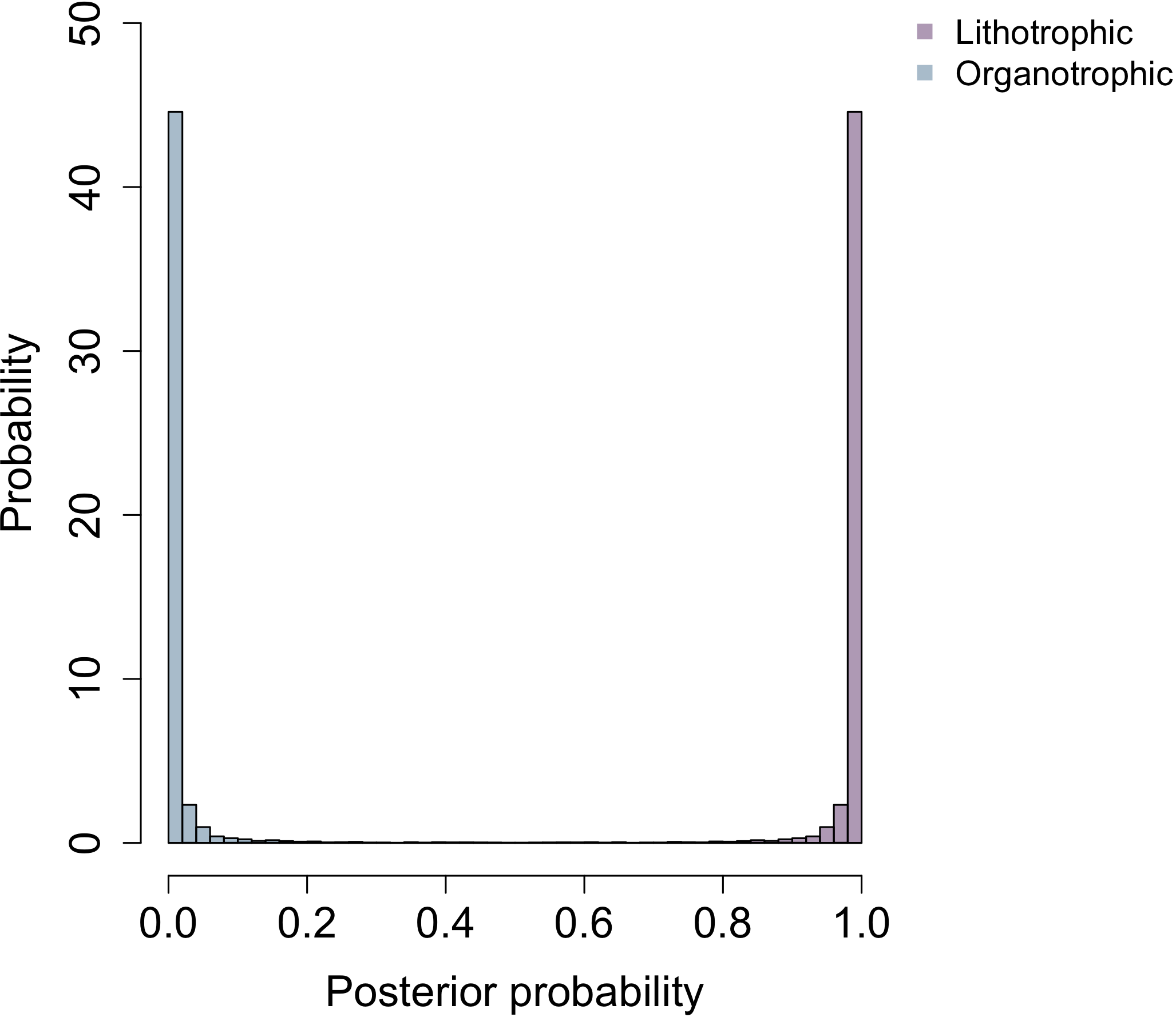

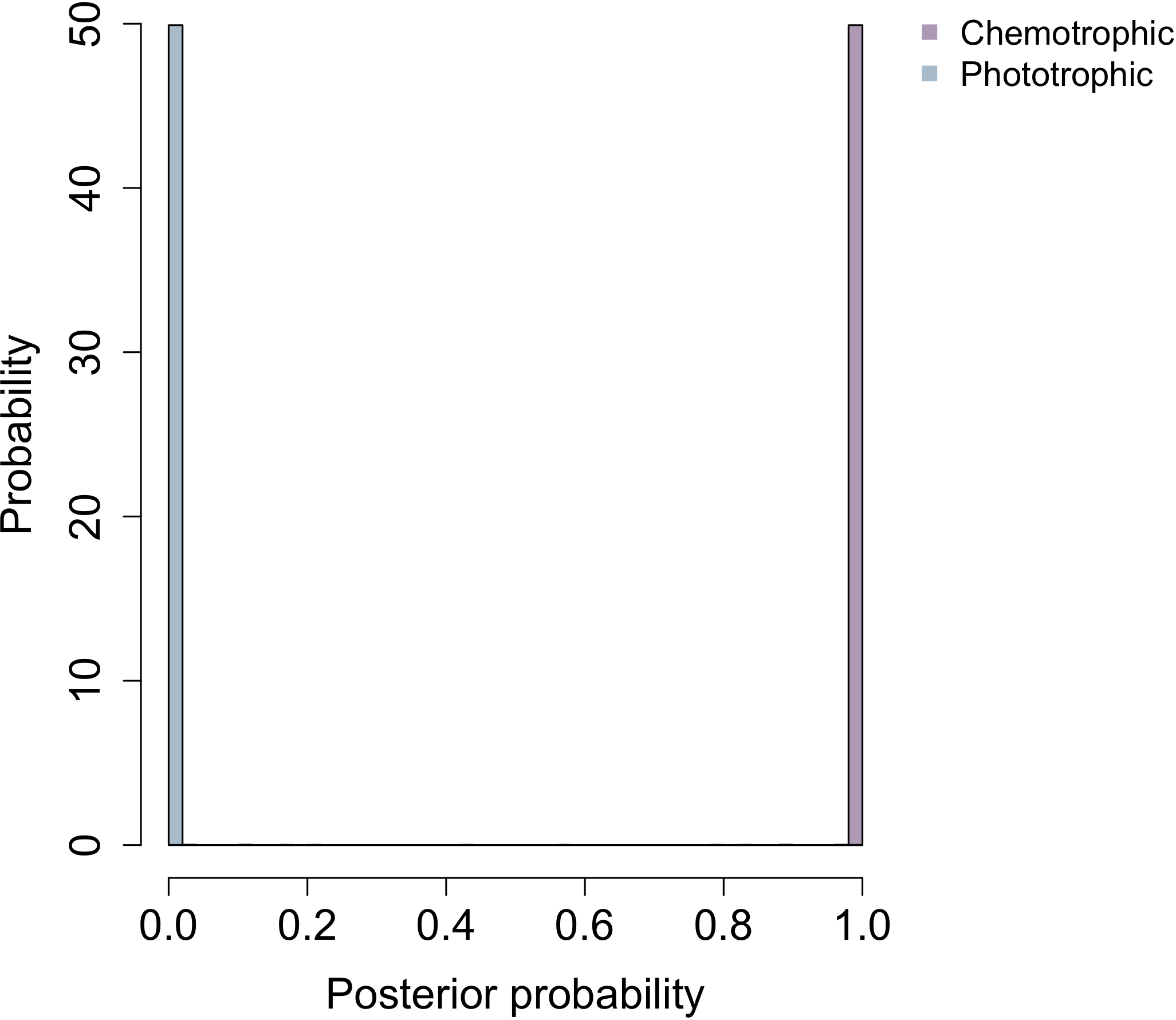

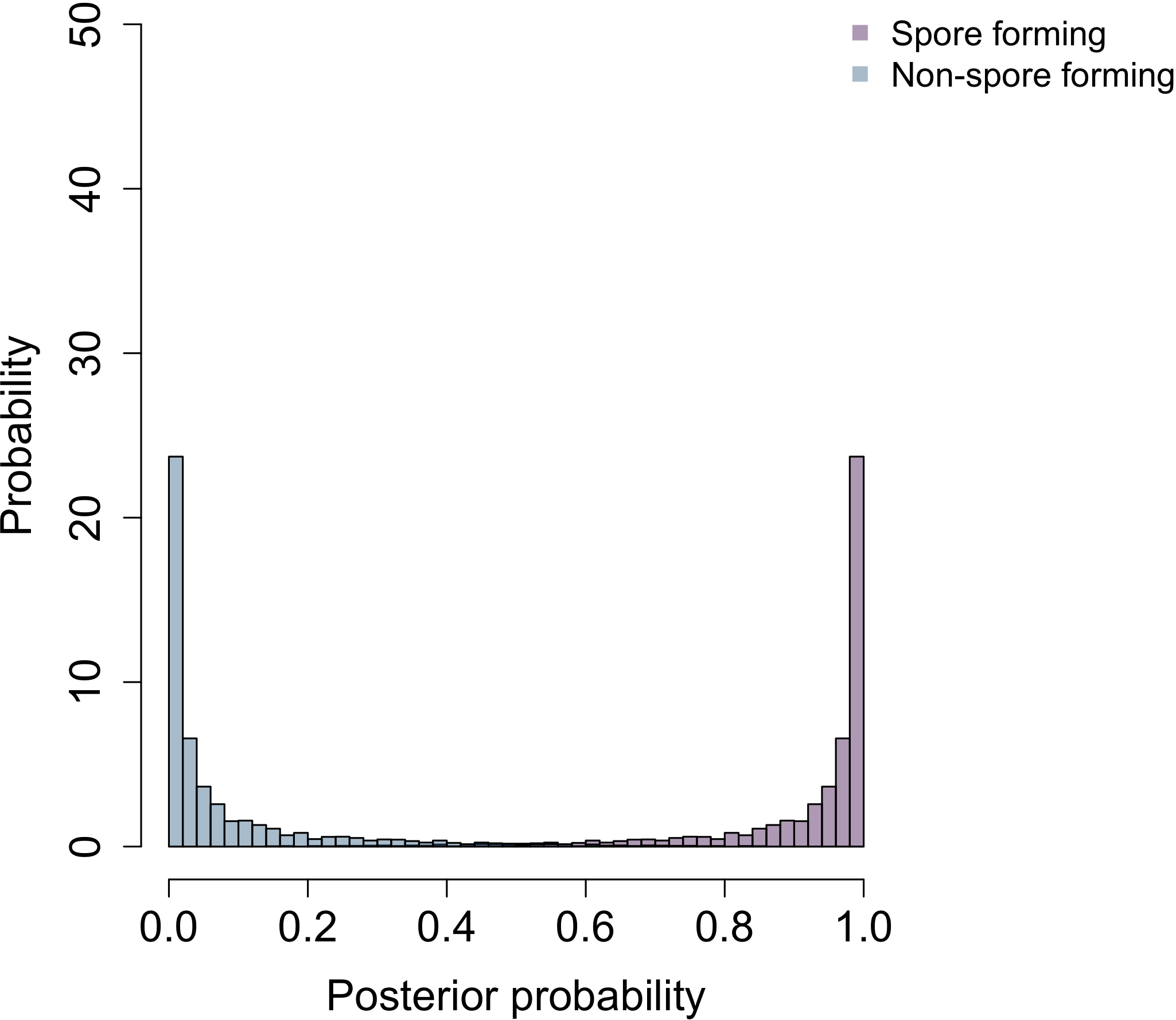

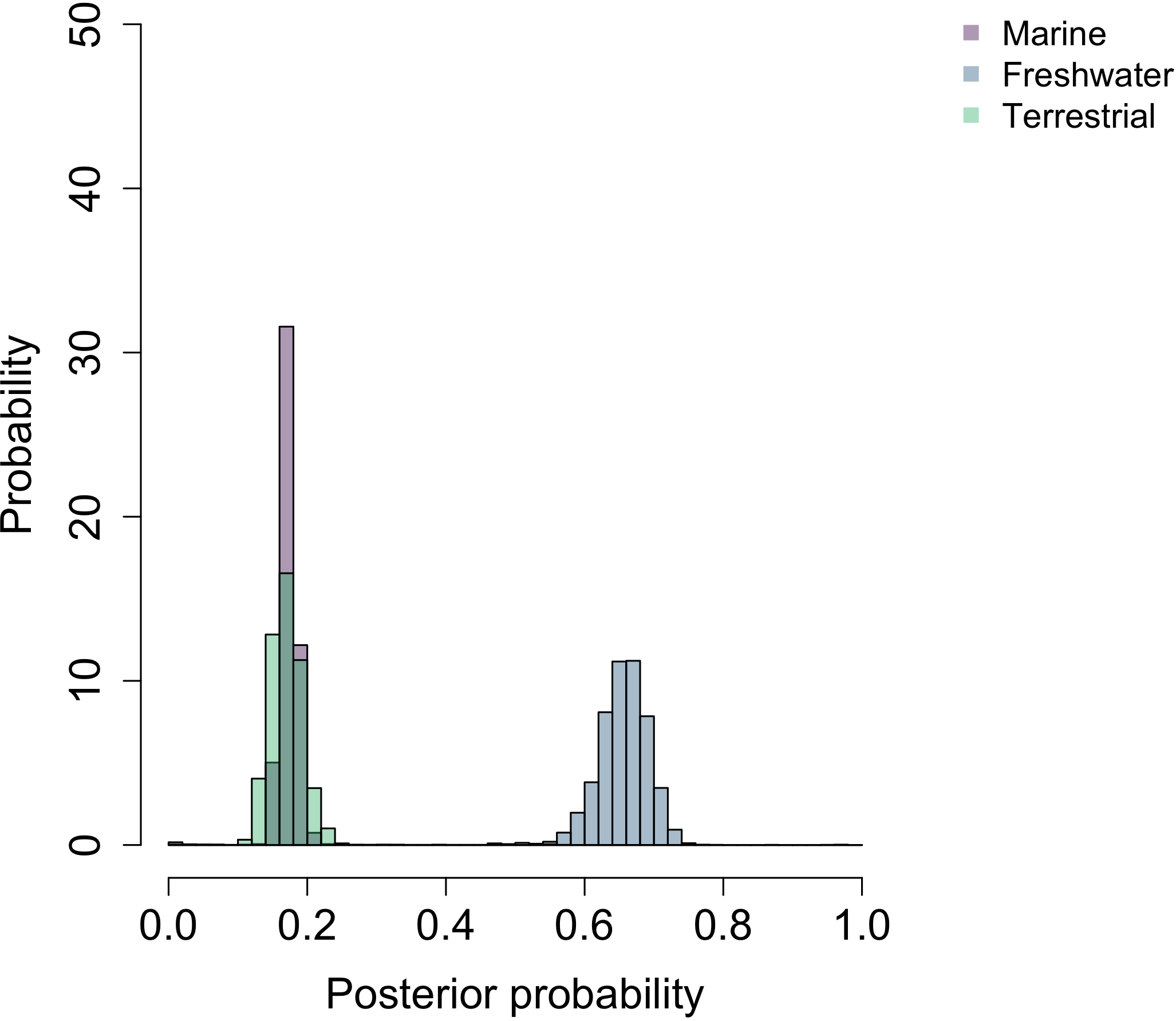

**Oxygen requirement**

**Energy source**

**Catalase**

**Electron donor**

**Sporulation**

**Habitat c**

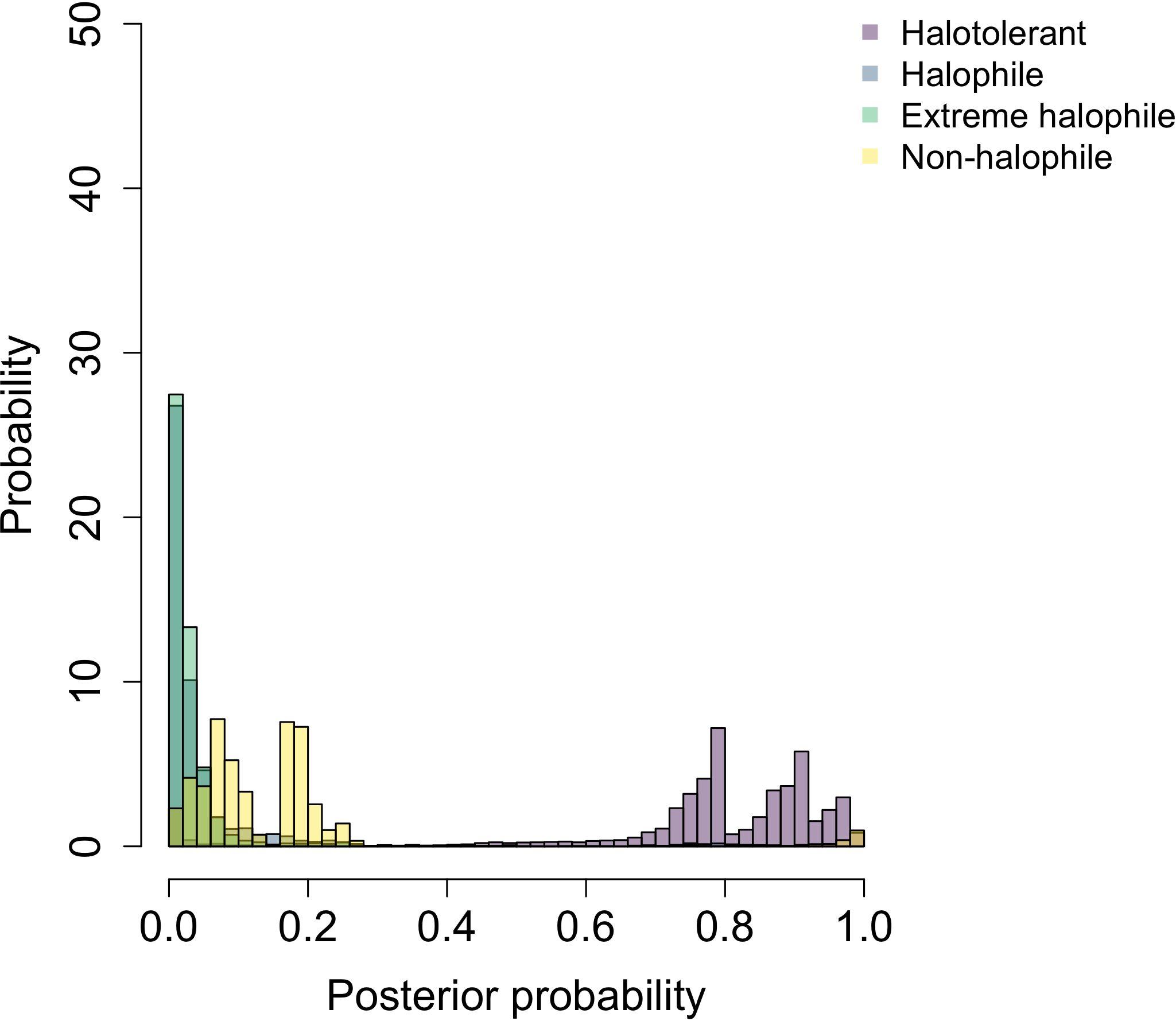

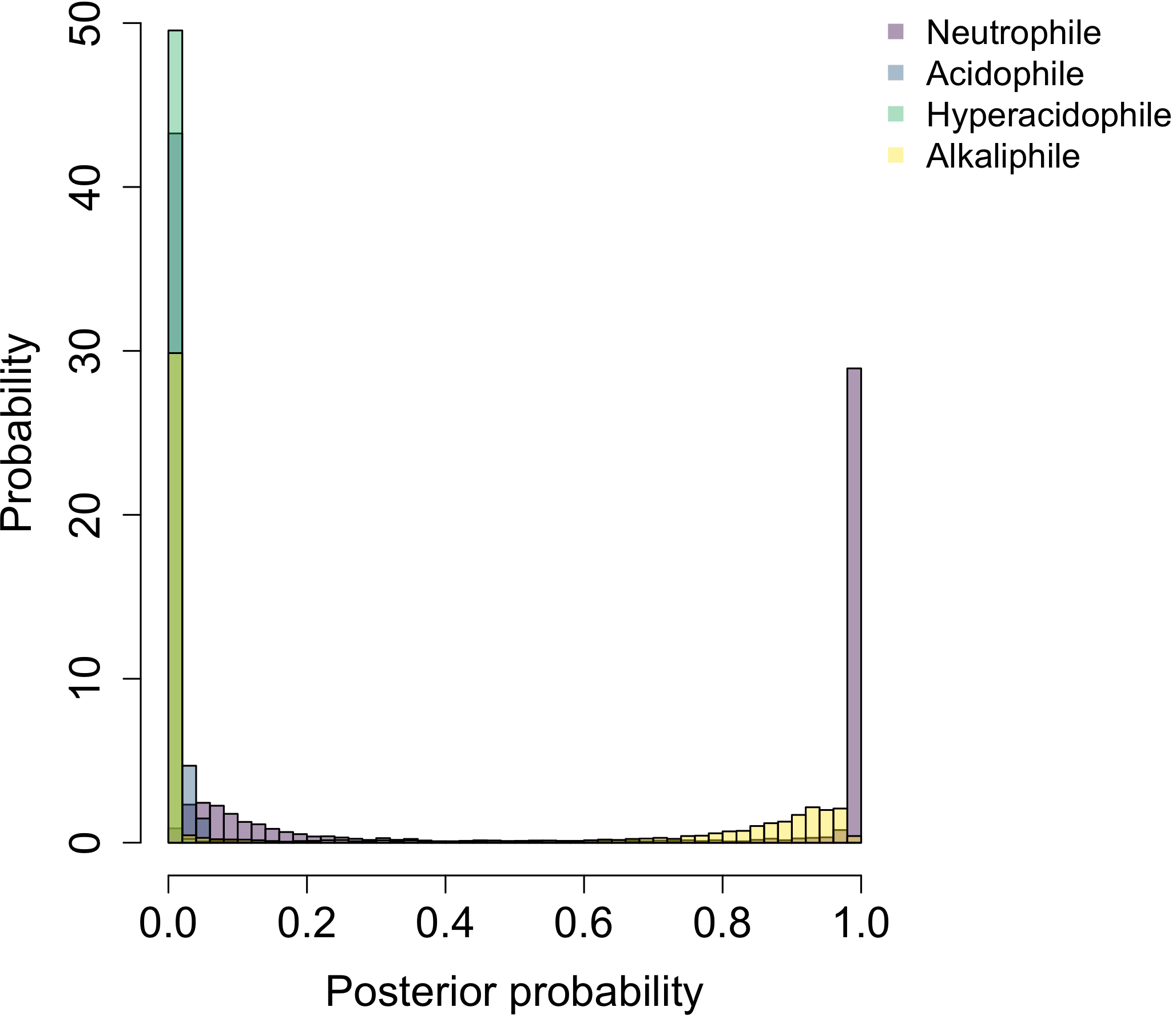

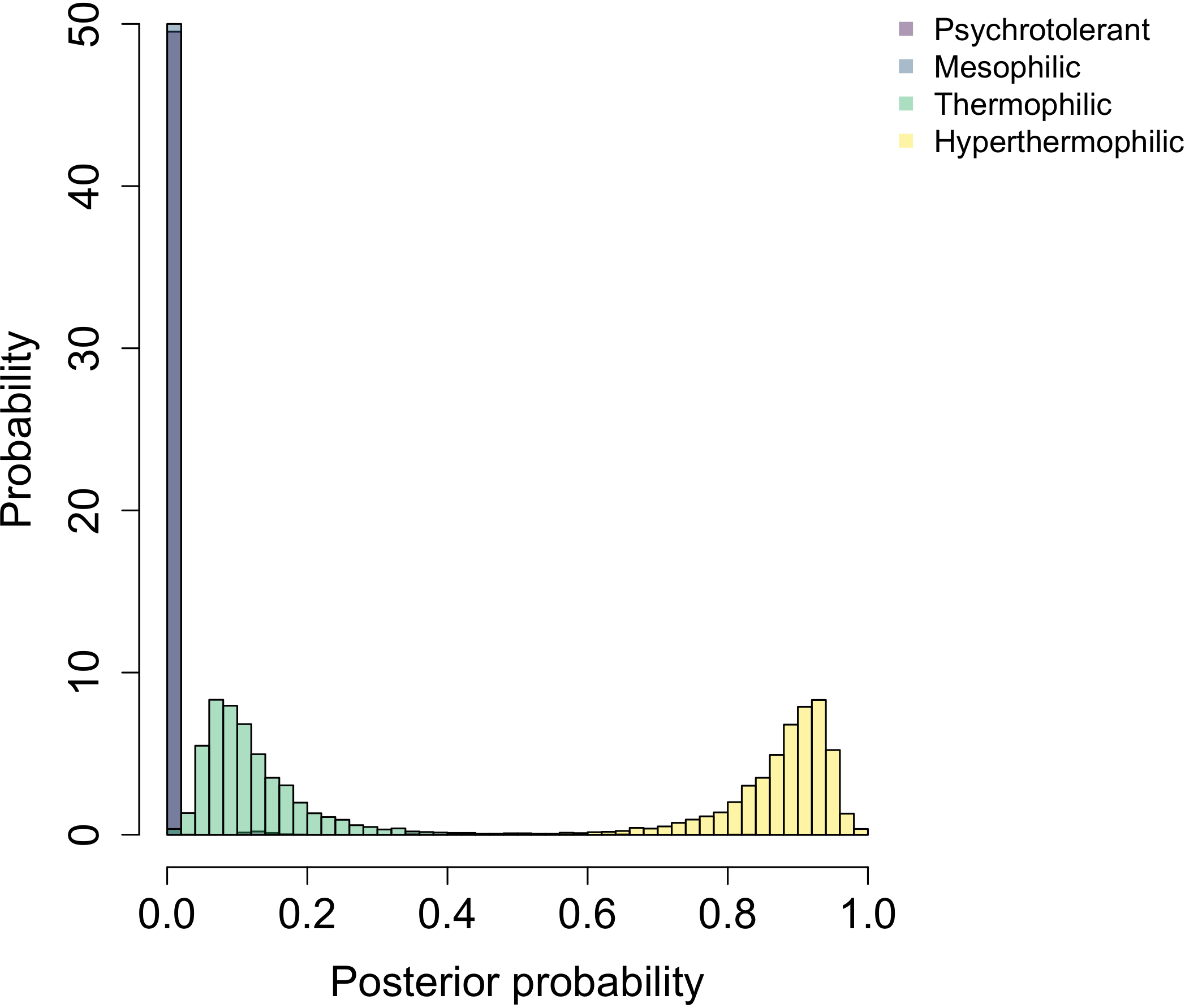

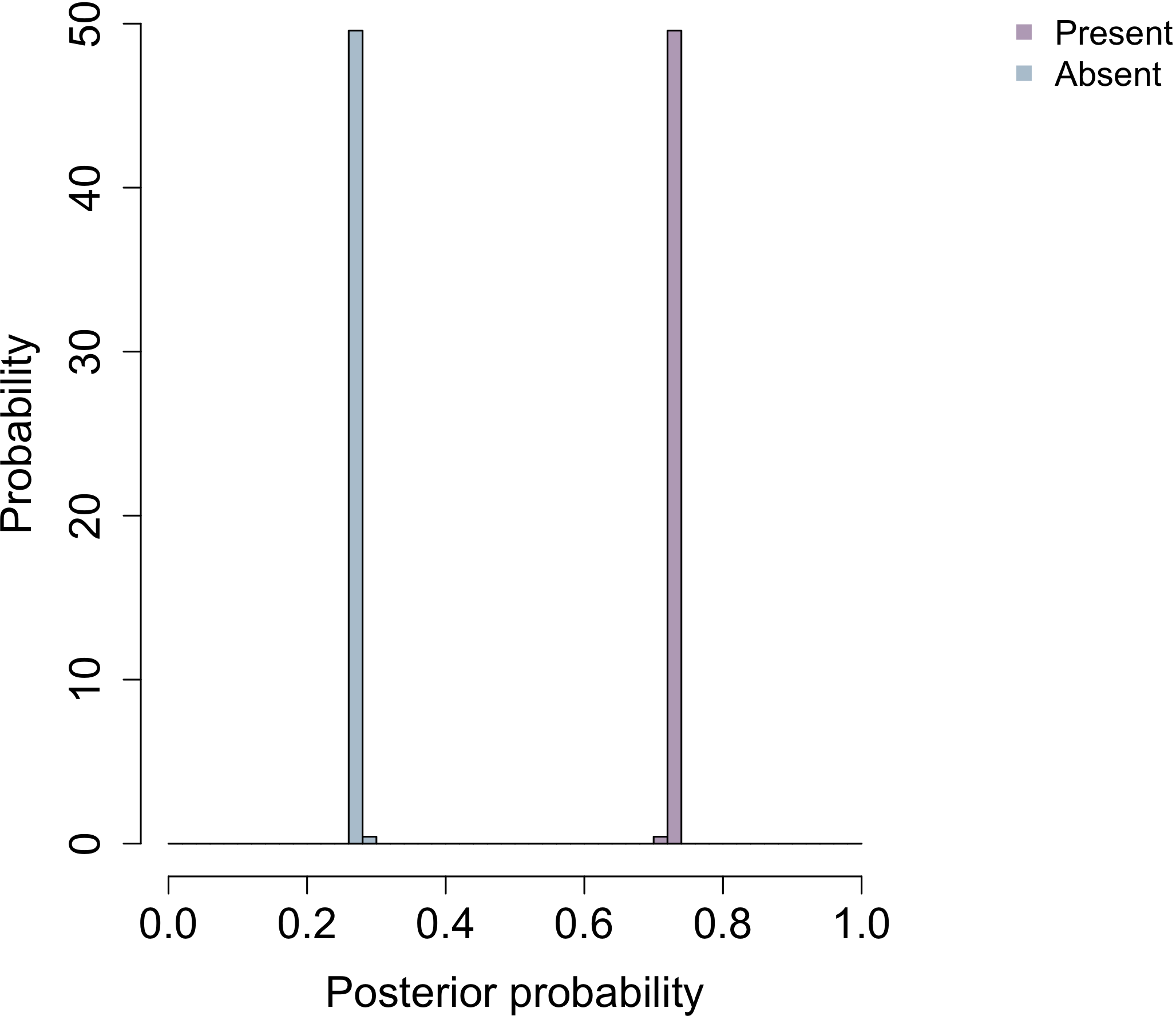

**pH**

**NaCl**

**Temperature**

**Catalase**

**c) LACA**

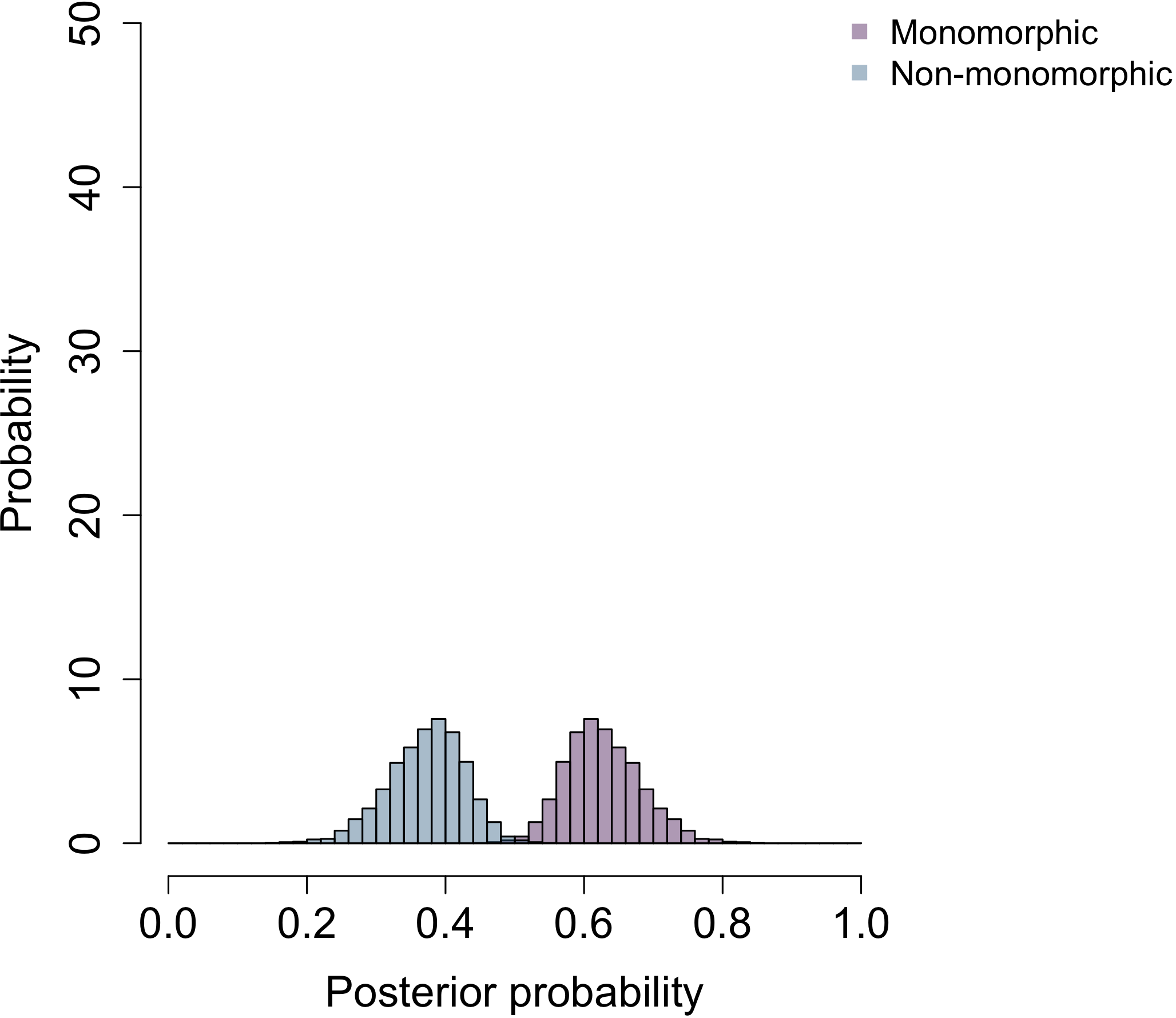

**Pleomorphism**

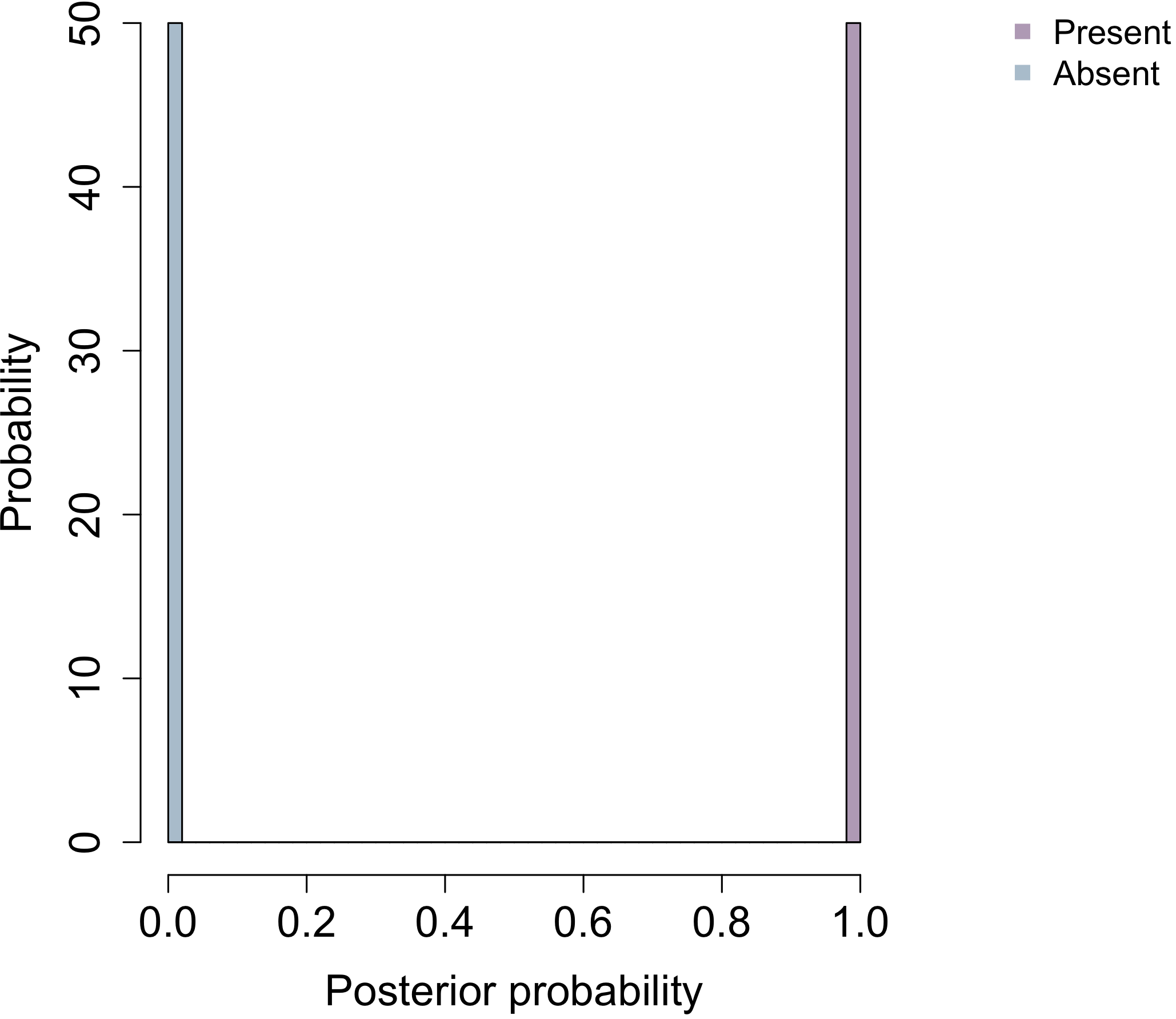

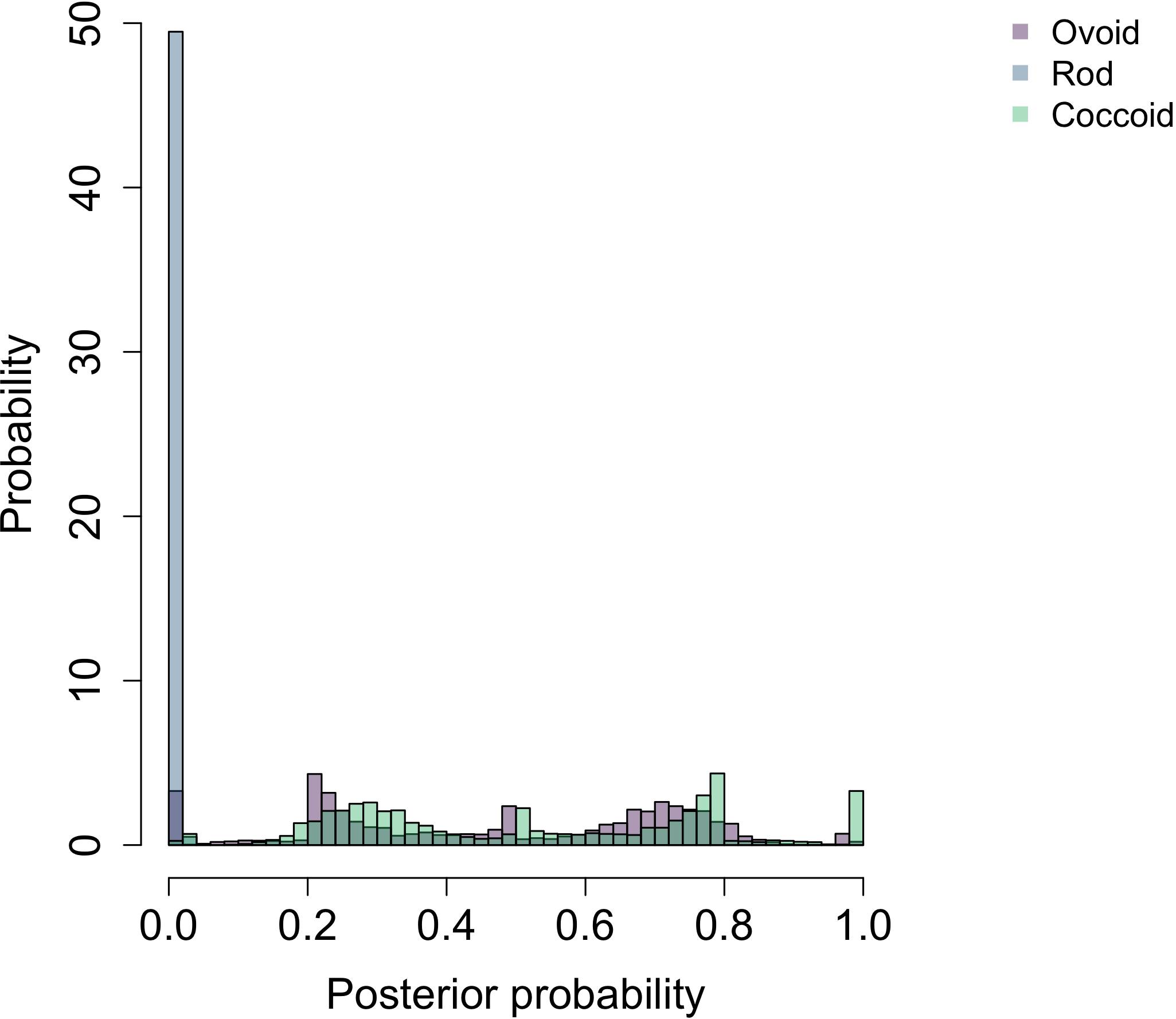

**Cell wall**

**Shape**

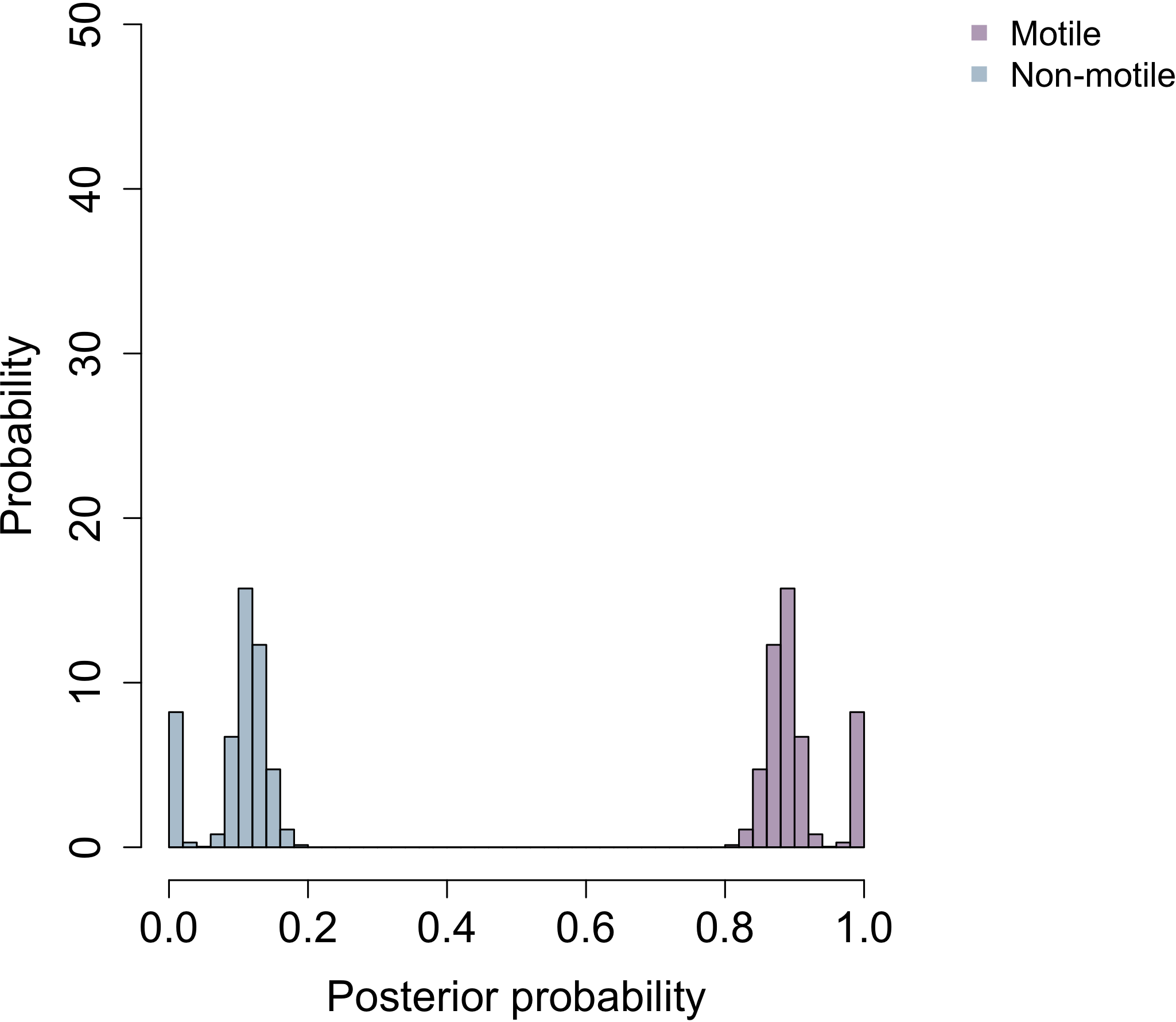

**Motility**

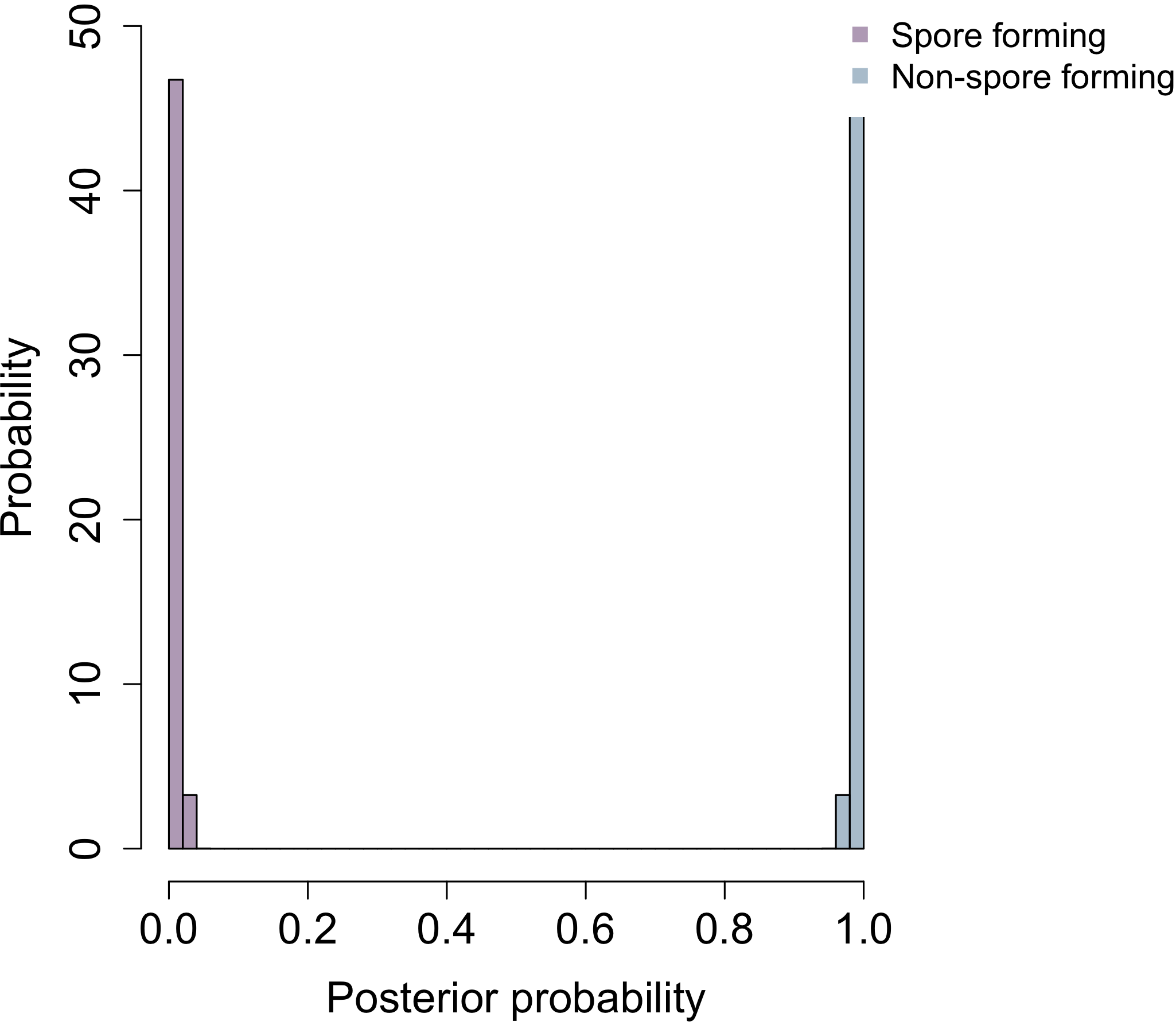

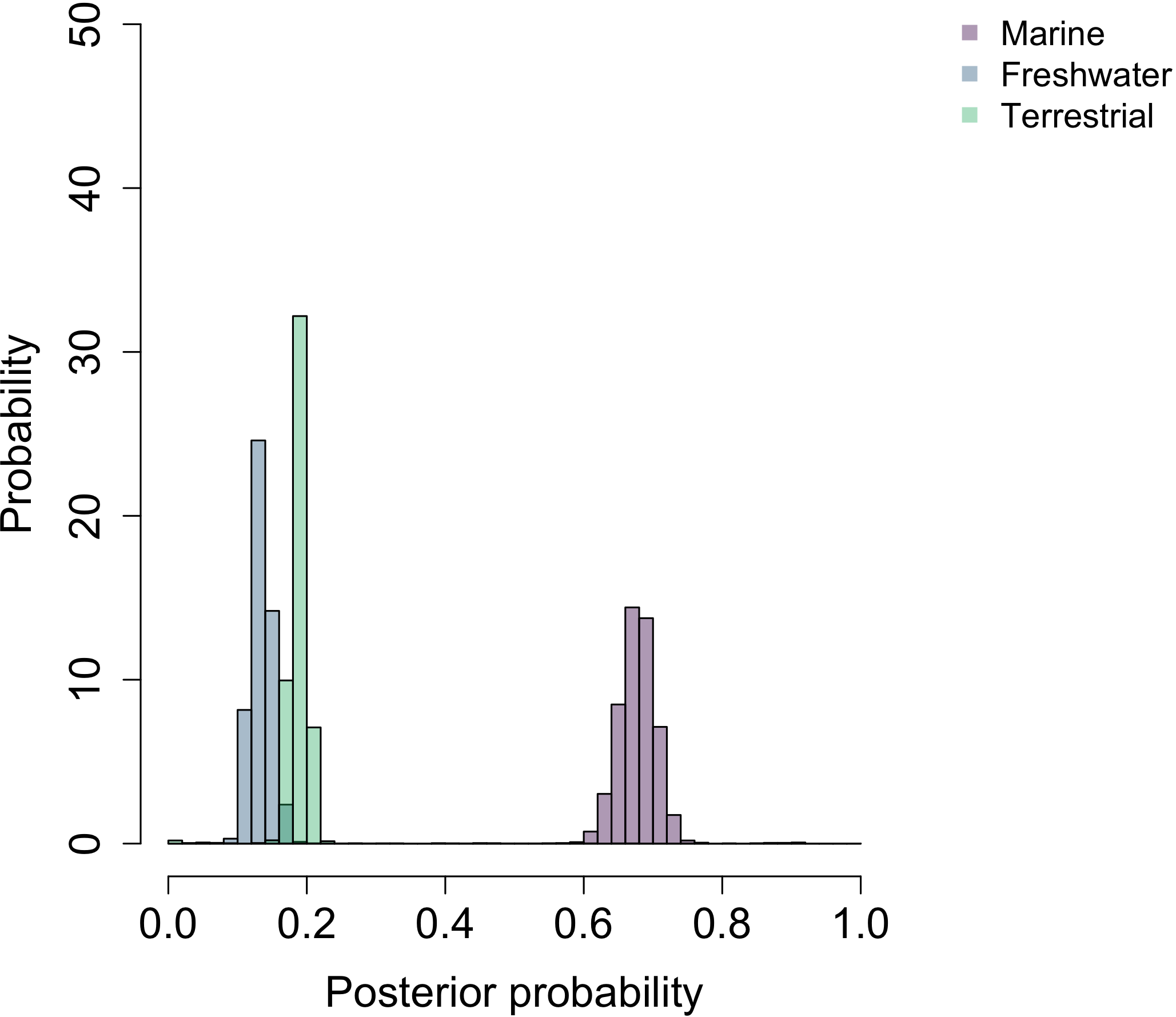

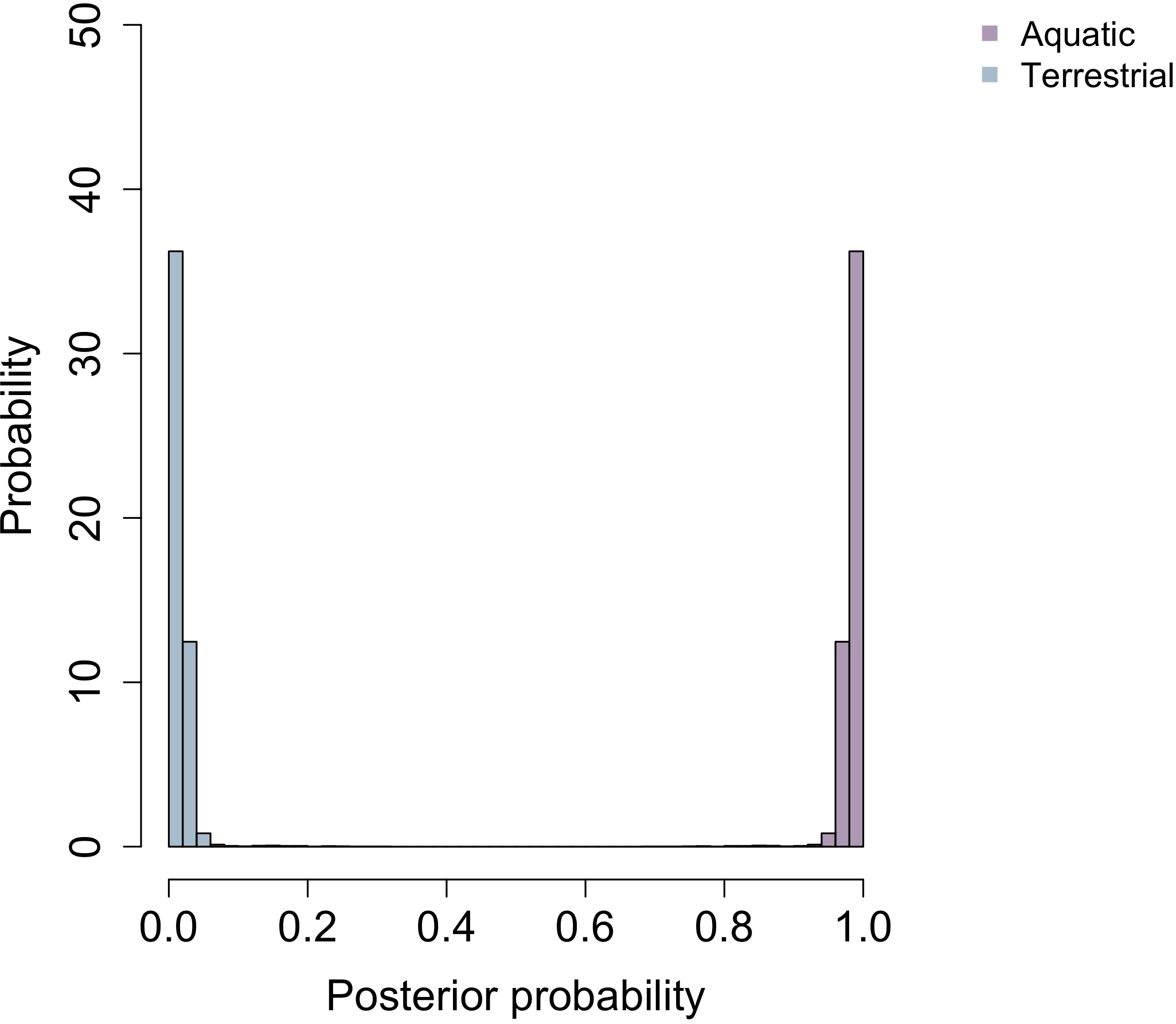

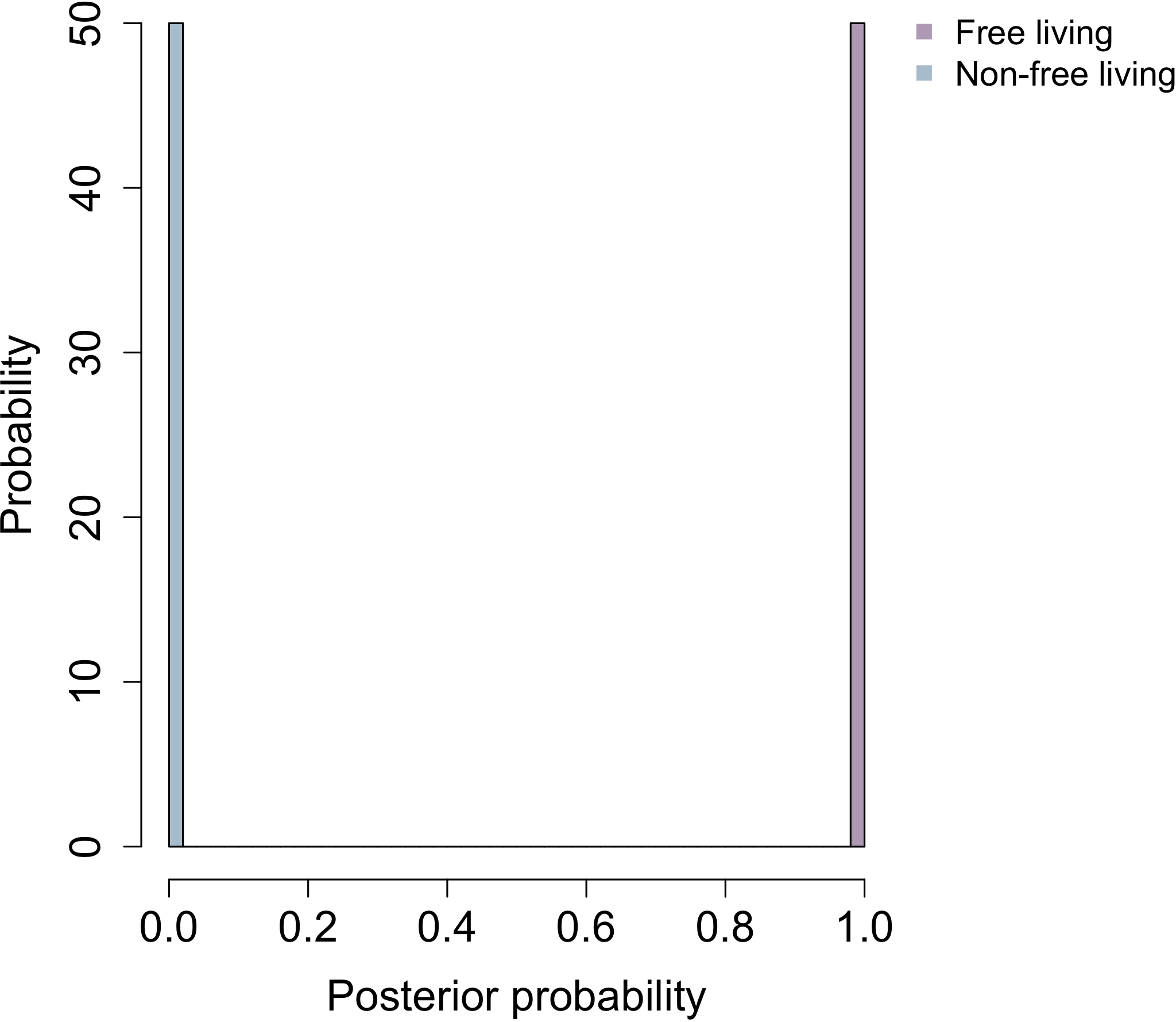

**Cell aggregation**

**Sporulation**

**Habitat c**

**Habitat b**

**Habitat a**

**Cell plan**

**Temperature**

**Oxidase**

**Oxygen requirement**

**Catalase**

**Energy source**

**Electron donor**

**pH**

**NaCl**

**Table S1. Bayesian estimates of the ancestral state for categorical traits from Chai et al’s tree.**

Posterior probabilities (PP) for all the characters from Chai et al’s tree are shown for LUCA, LBCA and LACA in the form of P (X) = value (0.00-1.00). Abbreviations: LUCA - last universal common ancestor; LBCA - last bacterial common ancestor; LACA - last archaeal common ancestor; P (X) - PP for character state X; HPD - Highest Posterior Density; 95% PI - 95% Probability Interval.

| **Character** | **Number of species** | **LUCA** | | | | | |
| --- | --- | --- | --- | --- | --- | --- | --- |
|  |  | **Mean PP** | **Median PP** | **Lower HPD** | **Upper HPD** | **Lower 95% PI** | **Upper 95% PI** |
| **Shape** | 2812 |  | | | | | |
| Ovoid (O) |  | P (O) = 0.64 | P (O) = 0.87 | 0.17 | 0.96 | 0.08 | 0.95 |
| Rod (R) |  | P (R) = 0.08 | P (R) = 0.04 | 0.00 | 0.48 | 0.01 | 0.54 |
| Coccoid (C) |  | P (C) = 0.28 | P (C) = 0.07 | 0.01 | 0.75 | 0.02 | 0.78 |
| **Pleomorphism** | 2843 |  | | | | | |
| Monomorphic (M) |  | P (M) = 0.43 | P (M) = 0.44 | 0.38 | 0.48 | 0.37 | 0.47 |
| Non-monomorphic (N) |  | P (N) = 0.57 | P (N) = 0.56 | 0.52 | 0.62 | 0.53 | 0.63 |
| **Motility** | 2586 |  | | | | | |
| Motile (M) |  | P (M) = 0.94 | P (M) = 0.98 | 0.86 | 0.99 | 0.85 | 0.99 |
| Non-motile (N) |  | P (N) = 0.06 | P (N) = 0.02 | 0.01 | 0.14 | 0.01 | 0.15 |
| **Cell wall** | 2761 |  | | | | | |
| Present (P) |  | P (P) = 1.00 | P (P) = 1.00 | 1.00 | 1.00 | 1.00 | 1.00 |
| Absent (A) |  | P (A) = 0.00 | P (A) = 0.00 | 0.00 | 0.00 | 0.00 | 0.00 |
| **Cell plan** | 2614 |  | | | | | |
| Monoderm (M) |  | P (M) = 1.00 | P (M) = 1.00 | 1.00 | 1.00 | 1.00 | 1.00 |
| Diderm (D) |  | P (D) = 0.00 | P (D) = 0.00 | 0.00 | 0.01 | 0.00 | 0.01 |
| **Cell aggregation** | 1363 |  | | | | | |
| Aggregating (A) |  | P (A) = 0.31 | P (A) = 0.32 | 0.15 | 0.45 | 0.15 | 0.45 |
| Non-aggregating (N) |  | P (N) = 0.69 | P (N) = 0.68 | 0.55 | 0.85 | 0.55 | 0.85 |
| **Habitat a** | 2752 |  | | | | | |
| Free-living (F) |  | P (F) = 0.98 | P (F) = 0.98 | 0.98 | 1.00 | 0.97 | 1.00 |
| Non-free living (N) |  | P (N) = 0.02 | P (N) = 0.02 | 0.00 | 0.02 | 0.00 | 0.03 |
| **Habitat b** | 1495 |  | | | | | |
| Aquatic (A) |  | P (A) = 0.95 | P (A) = 0.97 | 0.85 | 0.99 | 0.82 | 0.98 |
| Terrestrial (T) |  | P (T) = 0.05 | P (T) = 0.03 | 0.01 | 0.15 | 0.02 | 0.18 |
| **Habitat c** | 1486 |  | | | | | |
| Marine (M) |  | P (M) = 0.55 | P (M) = 0.56 | 0.39 | 0.62 | 0.38 | 0.62 |
| Freshwater (F) |  | P (F) = 0.27 | P (F) = 0.25 | 0.22 | 0.46 | 0.23 | 0.49 |
| Terrestrial (T) |  | P (T) = 0.18 | P (T) = 0.19 | 0.10 | 0.23 | 0.09 | 0.22 |
| **Sporulation** | 1943 |  | | | | | |
| Spore-forming (S) |  | P (S) = 0.01 | P (S) = 0.01 | 0.00 | 0.01 | 0.00 | 0.01 |
| Non-spore forming (N) |  | P (N) = 0.99 | P (N) = 0.99 | 0.99 | 1.00 | 0.99 | 1.00 |
| **Nutritional mode: Electron donor** | 932 |  | | | | | |
| Lithotrophic (L) |  | P (L) = 1.00 | P (L) = 1.00 | 1.00 | 1.00 | 0.99 | 1.00 |
| Organotrophic (O) |  | P (O) = 0.00 | P (O) = 0.00 | 0.00 | 0.00 | 0.00 | 0.00 |
| **Nutritional mode: Energy source** | 1036 |  | | | | | |
| Chemotrophic (C) |  | P (C) = 1.00 | P (C) = 1.00 | 1.00 | 1.00 | 1.00 | 1.00 |
| Phototrophic (P) |  | P (P) = 0.00 | P (P) = 0.00 | 0.00 | 0.00 | 0.00 | 0.00 |
| **Oxygen requirement** | 2532 |  | | | | | |
| Aerobic (A) |  | P (A) = 0.00 | P (A) = 0.00 | 0.00 | 0.01 | 0.00 | 0.01 |
| Anaerobic (N) |  | P (N) = 1.00 | P (N) = 1.00 | 0.99 | 1.00 | 0.99 | 1.00 |
| **Enzymes: catalase** | 1424 |  | | | | | |
| Present (P) |  | P (P) = 0.94 | P (P) = 0.93 | 0.89 | 1.00 | 0.89 | 1.00 |
| Absent (A) |  | P (A) = 0.06 | P (A) = 0.07 | 0.00 | 0.11 | 0.00 | 0.11 |
| **Enzymes: oxidase** | 1036 |  | | | | | |
| Present (P) |  | P (P) = 0.84 | P (P) = 0.84 | 0.78 | 0.90 | 0.77 | 0.90 |
| Absent (A) |  | P (A) = 0.16 | P (A) = 0.16 | 0.10 | 0.22 | 0.10 | 0.23 |
| **Temperature (optimum)** | 1838 |  | | | | | |
| Psychrotolerant (P) |  | P (P) = 0.01 | P (P) = 0.00 | 0.00 | 0.06 | 0.00 | 0.13 |
| Mesophilic (M) |  | P (M) = 0.00 | P (M) = 0.00 | 0.00 | 0.01 | 0.00 | 0.02 |
| Thermophilic (T) |  | P (T) = 0.10 | P (T) = 0.08 | 0.01 | 0.20 | 0.03 | 0.28 |
| Hyperthermophilic (H) |  | P (H) = 0.89 | P (H) = 0.92 | 0.72 | 1.00 | 0.53 | 0.97 |
| **NaCl (optimum)** | 524 |  | | | | | |
| Halotolerant (T) |  | P (T) = 0.74 | P (T) = 0.75 | 0.65 | 0.82 | 0.64 | 0.81 |
| Halophile (P) |  | P (P) = 0.08 | P (P) = 0.08 | 0.06 | 0.11 | 0.06 | 0.11 |
| Extreme halophile (E) |  | P (E) = 0.08 | P (E) = 0.07 | 0.05 | 0.11 | 0.05 | 0.11 |
| Non_halophile (N) |  | P (N) = 0.10 | P (N) = 0.10 | 0.08 | 0.13 | 0.08 | 0.13 |
| **pH (optimum)** | 1053 |  | | | | | |
| Neutrophile (N) |  | P (N) = 0.59 | P (N) = 0.62 | 0.28 | 0.81 | 0.17 | 0.76 |
| Acidophile (A) |  | P (A) = 0.27 | P (A) = 0.24 | 0.00 | 0.55 | 0.08 | 0.76 |
| Hyperacidophile (H) |  | P (H) = 0.01 | P (H) = 0.00 | 0.00 | 0.05 | 0.00 | 0.10 |
| Alkaliphile (K) |  | P (K) = 0.13 | P (K) = 0.13 | 0.01 | 0.24 | 0.03 | 0.25 |
| **Character** | **Number of species** | **LBCA** | | | | | |
|  |  | **Mean PP** | **Median PP** | **Lower HPD** | **Upper HPD** | **Lower 95% PI** | **Upper 95% PI** |
| **Shape** | 2812 |  | | | | | |
| Ovoid (O) |  | P (O) = 0.19 | P (O) = 0.20 | 0.00 | 0.58 | 0.00 | 0.60 |
| Rod (R) |  | P (R) = 0.76 | P (R) = 0.79 | 0.35 | 1.00 | 0.34 | 1.00 |
| Coccoid (C) |  | P (C) = 0.05 | P (C) = 0.01 | 0.00 | 0.21 | 0.00 | 0.24 |
| **Pleomorphism** | 2843 |  | | | | | |
| Monomorphic (M) |  | P (M) = 0.50 | P (M) = 0.50 | 0.48 | 0.52 | 0.47 | 0.52 |
| Non-monomorphic (N) |  | P (N) = 0.50 | P (N) = 0.50 | 0.48 | 0.52 | 0.48 | 0.53 |
| **Motility** | 2586 |  | | | | | |
| Motile (M) |  | P (M) = 0.94 | P (M) = 0.97 | 0.87 | 0.99 | 0.86 | 0.99 |
| Non-motile (N) |  | P (N) = 0.06 | P (N) = 0.03 | 0.01 | 0.13 | 0.01 | 0.01 |
| **Cell wall** | 2761 |  | | | | | |
| Present (P) |  | P (P) = 0.93 | P (P) = 0.93 | 0.93 | 0.93 | 0.93 | 0.93 |
| Absent (A) |  | P (A) = 0.07 | P (A) = 0.07 | 0.07 | 0.07 | 0.07 | 0.07 |
| **Cell plan** | 2614 |  | | | | | |
| Monoderm (M) |  | P (M) = 0.83 | P (M) = 0.97 | 0.43 | 1.00 | 0.38 | 1.00 |
| Diderm (D) |  | P (D) = 0.17 | P (D) = 0.03 | 0.00 | 0.57 | 0.00 | 0.62 |
| **Cell aggregation** | 1363 |  | | | | | |
| Aggregating (A) |  | P (A) = 0.40 | P (A) = 0.41 | 0.28 | 0.50 | 0.28 | 0.51 |
| Non-aggregating (N) |  | P (N) = 0.60 | P (N) = 0.59 | 0.50 | 0.72 | 0.49 | 0.72 |
| **Habitat a** | 2752 |  | | | | | |
| Free-living (F) |  | P (F) = 0.85 | P (F) = 0.82 | 0.81 | 0.93 | 0.81 | 0.93 |
| Non-free living (N) |  | P (N) = 0.15 | P (N) = 0.18 | 0.07 | 0.19 | 0.07 | 0.19 |
| **Habitat b** | 1495 |  | | | | | |
| Aquatic (A) |  | P (A) = 0.87 | P (A) = 0.89 | 0.74 | 0.92 | 0.70 | 0.91 |
| Terrestrial (T) |  | P (T) = 0.13 | P (T) = 0.11 | 0.08 | 0.26 | 0.09 | 0.30 |
| **Habitat c** | 1486 |  | | | | | |
| Marine (M) |  | P (M) = 0.36 | P (M) = 0.36 | 0.27 | 0.40 | 0.27 | 0.39 |
| Freshwater (F) |  | P (F) = 0.38 | P (F) = 0.36 | 0.36 | 0.50 | 0.36 | 0.53 |
| Terrestrial (T) |  | P (T) = 0.26 | P (T) = 0.27 | 0.16 | 0.30 | 0.15 | 0.29 |
| **Sporulation** | 1943 |  | | | | | |
| Spore-forming (S) |  | P (S) = 0.06 | P (S) = 0.05 | 0.03 | 0.10 | 0.03 | 0.10 |
| Non-spore forming (N) |  | P (N) = 0.94 | P (N) = 0.95 | 0.91 | 0.97 | 0.90 | 0.97 |
| **Nutritional mode: Electron donor** | 932 |  | | | | | |
| Lithotrophic (L) |  | P (L) = 0.94 | P (L) = 0.99 | 0.71 | 1.00 | 0.62 | 1.00 |
| Organotrophic (O) |  | P (O) = 0.06 | P (O) = 0.01 | 0.00 | 0.30 | 0.00 | 0.38 |
| **Nutritional mode: Energy source** | 1036 |  | | | | | |
| Chemotrophic (C) |  | P (C) = 1.00 | P (C) = 1.00 | 1.00 | 1.00 | 1.00 | 1.00 |
| Phototrophic (P) |  | P (P) = 0.00 | P (P) = 0.00 | 0.00 | 0.00 | 0.00 | 0.00 |
| **Oxygen requirement** | 2532 |  | | | | | |
| Aerobic (A) |  | P (A) = 0.03 | P (A) = 0.03 | 0.01 | 0.04 | 0.02 | 0.05 |
| Anaerobic (N) |  | P (N) = 0.97 | P (N) = 0.97 | 0.96 | 0.99 | 0.95 | 0.98 |
| **Enzymes: catalase** | 1424 |  | | | | | |
| Present (P) |  | P (P) = 0.88 | P (P) = 0.83 | 0.80 | 1.00 | 0.80 | 1.00 |
| Absent (A) |  | P (A) = 0.12 | P (A) = 0.17 | 0.00 | 0.20 | 0.00 | 0.20 |
| **Enzymes: oxidase** | 1036 |  | | | | | |
| Present (P) |  | P (P) = 0.80 | P (P) = 0.80 | 0.73 | 0.87 | 0.73 | 0.87 |
| Absent (A) |  | P (A) = 0.20 | P (A) = 0.20 | 0.13 | 0.27 | 0.13 | 0.27 |
| **Temperature (optimum)** | 1838 |  | | | | | |
| Psychrotolerant (P) |  | P (P) = 0.12 | P (P) = 0.07 | 0.00 | 0.43 | 0.00 | 0.48 |
| Mesophilic (M) |  | P (M) = 0.17 | P (M) = 0.13 | 0.02 | 0.47 | 0.03 | 0.50 |
| Thermophilic (T) |  | P (T) = 0.68 | P (T) = 0.76 | 0.12 | 0.92 | 0.01 | 0.90 |
| Hyperthermophilic (H) |  | P (H) = 0.03 | P (H) = 0.02 | 0.00 | 0.07 | 0.00 | 0.12 |
| **NaCl (optimum)** | 524 |  | | | | | |
| Halotolerant (T) |  | P (T) = 0.74 | P (T) = 0.75 | 0.62 | 0.85 | 0.60 | 0.84 |
| Halophile (P) |  | P (P) = 0.06 | P (P) = 0.06 | 0.03 | 0.10 | 0.03 | 0.10 |
| Extreme halophile (E) |  | P (E) = 0.08 | P (E) = 0.08 | 0.04 | 0.13 | 0.04 | 0.13 |
| Non_halophile (N) |  | P (N) = 0.12 | P (N) = 0.12 | 0.08 | 0.16 | 0.10 | 0.17 |
| **pH (optimum)** | 1053 |  | | | | | |
| Neutrophile (N) |  | P (N) = 0.55 | P (N) = 0.54 | 0.41 | 0.70 | 0.40 | 0.70 |
| Acidophile (A) |  | P (A) = 0.03 | P (A) = 0.02 | 0.00 | 0.06 | 0.01 | 0.15 |
| Hyperacidophile (H) |  | P (H) = 0.00 | P (H) = 0.00 | 0.00 | 0.02 | 0.00 | 0.03 |
| Alkaliphile (K) |  | P (K) = 0.42 | P (K) = 0.43 | 0.24 | 0.58 | 0.23 | 0.57 |
| **Character** | **Number of species** | **LACA** | | | | | |
|  |  | **Mean PP** | **Median PP** | **Lower HPD** | **Upper HPD** | **Lower 95% PI** | **Upper 95% PI** |
| **Shape** | 2812 |  | | | | | |
| Ovoid (O) |  | P (O) = 0.54 | P (O) = 0.61 | 0.18 | 0.80 | 0.20 | 0.81 |
| Rod (R) |  | P (R) = 0.01 | P (R) = 0.01 | 0.00 | 0.02 | 0.00 | 0.03 |
| Coccoid (C) |  | P (C) = 0.45 | P (C) = 0.39 | 0.20 | 0.82 | 0.18 | 0.80 |
| **Pleomorphism** | 2843 |  | | | | | |
| Monomorphic (M) |  | P (M) = 0.42 | P (M) = 0.43 | 0.37 | 0.47 | 0.37 | 0.46 |
| Non-monomorphic (N) |  | P (N) = 0.58 | P (N) = 0.57 | 0.53 | 0.63 | 0.54 | 0.63 |
| **Motility** | 2586 |  | | | | | |
| Motile (M) |  | P (M) = 0.84 | P (M) = 0.90 | 0.69 | 0.96 | 0.69 | 0.96 |
| Non-motile (N) |  | P (N) = 0.16 | P (N) = 0.10 | 0.04 | 0.31 | 0.04 | 0.31 |
| **Cell wall** | 2761 |  | | | | | |
| Present (P) |  | P (P) = 1.00 | P (P) = 1.00 | 1.00 | 1.00 | 1.00 | 1.00 |
| Absent (A) |  | P (A) = 0.00 | P (A) = 0.00 | 0.00 | 0.00 | 0.00 | 0.00 |
| **Cell plan** | 2614 |  | | | | | |
| Monoderm (M) |  | P (M) = 1.00 | P (M) = 1.00 | 1.00 | 1.00 | 1.00 | 1.00 |
| Diderm (D) |  | P (D) = 0.00 | P (D) = 0.00 | 0.00 | 0.00 | 0.00 | 0.00 |
| **Cell aggregation** | 1363 |  | | | | | |
| Aggregating (A) |  | P (A) = 0.34 | P (A) = 0.35 | 0.21 | 0.45 | 0.20 | 0.45 |
| Non-aggregating (N) |  | P (N) = 0.66 | P (N) = 0.65 | 0.55 | 0.79 | 0.55 | 0.80 |
| **Habitat a** | 2752 |  | | | | | |
| Free-living (F) |  | P (F) = 0.98 | P (F) = 0.98 | 0.97 | 0.99 | 0.97 | 0.99 |
| Non-free living (N) |  | P (N) = 0.02 | P (N) = 0.02 | 0.02 | 0.02 | 0.01 | 0.03 |
| **Habitat b** | 1495 |  | | | | | |
| Aquatic (A) |  | P (A) = 0.92 | P (A) = 0.94 | 0.80 | 0.97 | 0.77 | 0.96 |
| Terrestrial (T) |  | P (T) = 0.06 | P (T) = 0.08 | 0.03 | 0.20 | 0.04 | 0.23 |
| **Habitat c** | 1486 |  | | | | | |
| Marine (M) |  | P (M) = 0.56 | P (M) = 0.56 | 0.47 | 0.60 | 0.46 | 0.60 |
| Freshwater (F) |  | P (F) = 0.24 | P (F) = 0.23 | 0.21 | 0.38 | 0.22 | 0.40 |
| Terrestrial (T) |  | P (T) = 0.20 | P (T) = 0.21 | 0.13 | 0.23 | 0.12 | 0.23 |
| **Sporulation** | 1943 |  | | | | | |
| Spore-forming (S) |  | P (S) = 0.01 | P (S) = 0.01 | 0.00 | 0.01 | 0.00 | 0.02 |
| Non-spore forming (N) |  | P (N) = 0.99 | P (N) = 0.99 | 0.99 | 1.00 | 0.98 | 1.00 |
| **Nutritional mode: Electron donor** | 932 |  | | | | | |
| Lithotrophic (L) |  | P (L) = 0.1 | P (L) = 1.00 | 1.00 | 1.00 | 0.99 | 1.00 |
| Organotrophic (O) |  | P (O) = 0.00 | P (O) = 0.00 | 0.00 | 0.00 | 0.00 | 0.01 |
| **Nutritional mode: Energy source** | 1036 |  | | | | | |
| Chemotrophic (C) |  | P (C) = 1.00 | P (C) = 1.00 | 1.00 | 1.00 | 1.00 | 1.00 |
| Phototrophic (P) |  | P (P) = 0.00 | P (P) = 0.00 | 0.00 | 0.00 | 0.00 | 0.00 |
| **Oxygen requirement** | 2532 |  | | | | | |
| Aerobic (A) |  | P (A) = 0.06 | P (A) = 0.06 | 0.06 | 0.06 | 0.06 | 0.06 |
| Anaerobic (N) |  | P (N) = 0.94 | P (N) = 0.94 | 0.94 | 0.94 | 0.94 | 0.94 |
| **Enzymes: catalase** | 1424 |  | | | | | |
| Present (P) |  | P (P) = 1.00 | P (P) = 1.00 | 1.00 | 1.00 | 1.00 | 1.00 |
| Absent (A) |  | P (A) = 0.00 | P (A) = 0.00 | 0.00 | 0.00 | 0.00 | 0.00 |
| **Enzymes: oxidase** | 1036 |  | | | | | |
| Present (P) |  | P (P) = 1.00 | P (P) = 1.00 | 1.00 | 1.00 | 1.00 | 1.00 |
| Absent (A) |  | P (A) = 0.00 | P (A) = 0.00 | 0.00 | 0.00 | 0.00 | 0.00 |
| **Temperature (optimum)** | 1838 |  | | | | | |
| Psychrotolerant (P) |  | P (P) = 0.01 | P (P) = 0.00 | 0.00 | 0.03 | 0.00 | 0.03 |
| Mesophilic (M) |  | P (M) = 0.00 | P (M) = 0.00 | 0.00 | 0.00 | 0.00 | 0.00 |
| Thermophilic (T) |  | P (T) = 0.02 | P (T) = 0.02 | 0.01 | 0.05 | 0.01 | 0.08 |
| Hyperthermophilic (H) |  | P (H) = 0.97 | P (H) = 0.98 | 0.93 | 0.99 | 0.90 | 0.99 |
| **NaCl (optimum)** | 524 |  | | | | | |
| Halotolerant (T) |  | P (T) = 0.55 | P (T) = 0.55 | 0.50 | 0.57 | 0.48 | 0.57 |
| Halophile (P) |  | P (P) = 0.15 | P (P) = 0.15 | 0.15 | 0.17 | 0.15 | 0.17 |
| Extreme halophile (E) |  | P (E) = 0.13 | P (E) = 0.13 | 0.11 | 0.18 | 0.10 | 0.17 |
| Non-halophile (N) |  | P (N) = 0.17 | P (N) = 0.17 | 0.17 | 0.18 | 0.17 | 0.18 |
| **pH (optimum)** | 1053 |  | | | | | |
| Neutrophile (N) |  | P (N) = 0.33 | P (N) = 0.33 | 0.08 | 0.52 | 0.10 | 0.55 |
| Acidophile (A) |  | P (A) = 0.60 | P (A) = 0.60 | 0.40 | 0.90 | 0.34 | 0.86 |
| Hyperacidophile (H) |  | P (H) = 0.02 | P (H) = 0.00 | 0.00 | 0.10 | 0.00 | 0.21 |
| Alkaliphile (K) |  | P (K) = 0.05 | P (K) = 0.05 | 0.01 | 0.11 | 0.01 | 0.13 |

Table S2. Bayesian estimates of the ancestral state for categorical traits from Segata et al’s tree.

Posterior probabilities (PP) for all the characters from Segata et al’s tree are shown for LUCA, LBCA and LACA in the form of P (X) = value (0.00-1.00). Abbreviations: LUCA - last universal common ancestor; LBCA -last bacterial common ancestor; LACA - last archaeal common ancestor; P (X) - PP for character state X; HPD - Highest Posterior Density; 95% PI - 95% Probability Interval.

| **Character** | **Number of species** | **LUCA** | | | | | |
| --- | --- | --- | --- | --- | --- | --- | --- |
|  |  | **Mean PP** | **Median PP** | **Lower HPD** | **Upper HPD** | **Lower 95% PI** | **Upper 95% PI** |
| **Shape** | 1372 |  | | | | | |
| Ovoid (O) |  | P (O) = 0.65 | P (O) = 0.79 | 0.03 | 0.89 | 0.01 | 0.88 |
| Rod (R) |  | P (R) = 0.22 | P (R) = 0.14 | 0.06 | 0.67 | 0.07 | 0.71 |
| Coccoid (C) |  | P (C) = 0.13 | P (C) = 0.06 | 0.01 | 0.43 | 0.02 | 0.48 |
| **Pleomorphism** | 1385 |  | | | | | |
| Monomorphic (M) |  | P (M) = 0.63 | P (M) = 0.63 | 0.50 | 0.78 | 0.50 | 0.79 |
| Non-monomorphic (N) |  | P (N) = 0.37 | P (N) = 0.37 | 0.22 | 0.50 | 0.21 | 0.50 |
| **Motility** | 1251 |  | | | | | |
| Motile (M) |  | P (M) = 0.92 | P (M) = 0.90 | 0.89 | 1.00 | 0.89 | 1.00 |
| Non-motile (N) |  | P (N) = 0.08 | P (N) = 0.10 | 0.00 | 0.11 | 0.00 | 0.11 |
| **Cell wall** | 1331 |  | | | | | |
| Present (P) |  | P (P) = 1.00 | P (P) = 1.00 | 1.00 | 1.00 | 1.00 | 1.00 |
| Absent (A) |  | P (A) = 0.00 | P (A) = 0.00 | 0.00 | 0.00 | 0.00 | 0.00 |
| **Cell plan** | 1263 |  | | | | | |
| Monoderm (M) |  | P (M) = 0.95 | P (M) = 1.00 | 0.44 | 1.00 | 0.44 | 1.00 |
| Diderm (D) |  | P (D) = 0.05 | P (D) = 0.00 | 0.00 | 0.56 | 0.00 | 0.56 |
| **Cell aggregation** | 762 |  | | | | | |
| Aggregating (A) |  | P (A) = 0.48 | P (A) = 0.49 | 0.25 | 0.66 | 0.23 | 0.65 |
| Non-aggregating (N) |  | P (N) = 0.52 | P (N) = 0.51 | 0.34 | 0.75 | 0.35 | 0.77 |
| **Habitat a** | 1339 |  | | | | | |
| Free-living (F) |  | P (F) = 1.00 | P (F) = 1.00 | 0.99 | 1.00 | 0.99 | 1.00 |
| Non-free living (N) |  | P (N) = 0.00 | P (N) = 0.00 | 0.00 | 0.01 | 0.00 | 0.01 |
| **Habitat b** | 654 |  | | | | | |
| Aquatic (A) |  | P (A) = 0.98 | P (A) = 0.98 | 0.97 | 0.99 | 0.97 | 0.99 |
| Terrestrial (T) |  | P (T) = 0.02 | P (T) = 0.02 | 0.01 | 0.03 | 0.01 | 0.03 |
| **Habitat c** | 646 |  | | | | | |
| Marine (M) |  | P (M) = 0.43 | P (M) = 0.43 | 0.40 | 0.45 | 0.40 | 0.45 |
| Freshwater (F) |  | P (F) = 0.40 | P (F) = 0.40 | 0.38 | 0.41 | 0.38 | 0.41 |
| Terrestrial (T) |  | P (T) = 0.17 | P (T) = 0.17 | 0.14 | 0.22 | 0.14 | 0.22 |
| **Sporulation** | 937 |  | | | | | |
| Spore-forming (S) |  | P (S) = 0.75 | P (S) = 0.86 | 0.19 | 1.00 | 0.13 | 0.99 |
| Non-spore forming (N) |  | P (N) = 0.25 | P (N) = 0.14 | 0.00 | 0.81 | 0.01 | 0.87 |
| **Nutritional mode: Electron donor** | 434 |  | | | | | |
| Lithotrophic (L) |  | P (L) = 1.00 | P (L) = 1.00 | 1.00 | 1.00 | 1.00 | 1.00 |
| Organotrophic (O) |  | P (O) = 0.00 | P (O) = 0.00 | 0.00 | 0.00 | 0.00 | 0.00 |
| **Nutritional mode: Energy source** | 492 |  | | | | | |
| Chemotrophic (C) |  | P (C) = 1.00 | P (C) = 1.00 | 1.00 | 1.00 | 1.00 | 1.00 |
| Phototrophic (P) |  | P (P) = 0.00 | P (P) = 0.00 | 0.00 | 0.00 | 0.00 | 0.00 |
| **Oxygen requirement** | 1233 |  | | | | | |
| Aerobic (A) |  | P (A) = 0.01 | P (A) = 0.01 | 0.01 | 0.01 | 0.01 | 0.01 |
| Anaerobic (N) |  | P (N) = 0.99 | P (N) = 0.99 | 0.99 | 0.99 | 0.99 | 0.99 |
| **Enzymes: catalase** | 565 |  | | | | | |
| Present (P) |  | P (P) = 0.96 | P (P) = 0.96 | 0.93 | 0.98 | 0.93 | 0.98 |
| Absent (A) |  | P (A) = 0.04 | P (A) = 0.04 | 0.02 | 0.07 | 0.02 | 0.07 |
| **Enzymes: oxidase** | 398 |  | | | | | |
| Present (P) |  | P (P) = 0.84 | P (P) = 0.84 | 0.80 | 0.90 | 0.79 | 0.89 |
| Absent (A) |  | P (A) = 0.16 | P (A) = 0.16 | 0.11 | 0.21 | 0.11 | 0.21 |
| **Temperature (optimum)** | 888 |  | | | | | |
| Psychrotolerant (P) |  | P (P) = 0.00 | P (P) = 0.00 | 0.00 | 0.00 | 0.00 | 0.03 |
| Mesophilic (M) |  | P (M) = 0.00 | P (M) = 0.00 | 0.00 | 0.00 | 0.00 | 0.00 |
| Thermophilic (T) |  | P (T) = 0.00 | P (T) = 0.04 | 0.00 | 0.00 | 0.00 | 0.01 |
| Hyperthermophilic (H) |  | P (H) = 1.00 | P (H) = 1.00 | 0.99 | 1.00 | 0.97 | 1.00 |
| **NaCl (optimum)** | 191 |  | | | | | |
| Halotolerant (T) |  | P (T) = 0.80 | P (T) = 0.90 | 0.00 | 1.00 | 0.00 | 1.00 |
| Halophile (P) |  | P (P) = 0.02 | P (P) = 0.01 | 0.00 | 0.06 | 0.00 | 0.07 |
| Extreme halophile (E) |  | P (E) = 0.10 | P (E) = 0.03 | 0.00 | 1.00 | 0.00 | 1.00 |
| Non-halophile (N) |  | P (N) = 0.08 | P (N) = 0.04 | 0.00 | 0.25 | 0.00 | 0.92 |
| **pH (optimum)** | 481 |  | | | | | |
| Neutrophile (N) |  | P (N) = 0.57 | P (N) = 0.64 | 0.14 | 1.00 | 0.10 | 0.99 |
| Acidophile (A) |  | P (A) = 0.21 | P (A) = 0.22 | 0.00 | 0.36 | 0.00 | 0.38 |
| Hyperacidophile (H) |  | P (H) = 0.01 | P (H) = 0.00 | 0.00 | 0.07 | 0.00 | 0.11 |
| Alkaliphile (K) |  | P (K) = 0.21 | P (K) = 0.07 | 0.00 | 0.68 | 0.00 | 0.75 |
| **Character** | **Number of species** | **LBCA** | | | | | |
|  |  | **Mean PP** | **Median PP** | **Lower HPD** | **Upper HPD** | **Lower 95% PI** | **Upper 95% PI** |
| **Shape** | 1372 |  | | | | | |
| Ovoid (O) |  | P (O) = 0.09 | P (O) = 0.06 | 0.00 | 0.40 | 0.00 | 0.48 |
| Rod (R) |  | P (R) = 0.91 | P (R) = 0.94 | 0.60 | 1.00 | 0.52 | 1.00 |
| Coccoid (C) |  | P (C) = 0.00 | P (C) = 0.00 | 0.00 | 0.00 | 0.00 | 0.00 |
| **Pleomorphism** | 1385 |  | | | | | |
| Monomorphic (M) |  | P (M) = 0.60 | P (M) = 0.60 | 0.47 | 0.75 | 0.47 | 0.75 |
| Non-monomorphic (N) |  | P (N) = 0.40 | P (N) = 0.40 | 0.25 | 0.53 | 0.25 | 0.53 |
| **Motility** | 1251 |  | | | | | |
| Motile (M) |  | P (M) = 0.78 | P (M) = 0.75 | 0.70 | 1.00 | 0.68 | 1.00 |
| Non-motile (N) |  | P (N) = 0.22 | P (N) = 0.25 | 0.01 | 0.30 | 0.01 | 0.32 |
| **Cell wall** | 1331 |  | | | | | |
| Present (P) |  | P (P) = 1.00 | P (P) = 1.00 | 1.00 | 1.00 | 1.00 | 1.00 |
| Absent (A) |  | P (A) = 0.00 | P (A) = 0.00 | 0.00 | 0.00 | 0.00 | 0.00 |
| **Cell plan** | 1263 |  | | | | | |
| Monoderm (M) |  | P (M) = 0.81 | P (M) = 0.94 | 0.00 | 0.99 | 0.00 | 0.99 |
| Diderm (D) |  | P (D) = 0.19 | P (D) = 0.06 | 0.01 | 1.00 | 0.00 | 1.00 |
| **Cell aggregation** | 762 |  | | | | | |
| Aggregating (A) |  | P (A) = 0.44 | P (A) = 0.46 | 0.29 | 0.55 | 0.26 | 0.54 |
| Non-aggregating (N) |  | P (N) = 0.56 | P (N) = 0.54 | 0.45 | 0.71 | 0.46 | 0.74 |
| **Habitat a** | 1339 |  | | | | | |
| Free-living (F) |  | P (F) = 1.00 | P (F) = 1.00 | 1.00 | 1.00 | 1.00 | 1.00 |
| Non-free living (N) |  | P (N) = 0.00 | P (N) = 0.00 | 0.00 | 0.00 | 0.00 | 0.00 |
| **Habitat b** | 654 |  | | | | | |
| Aquatic (A) |  | P (A) = 0.98 | P (A) = 0.98 | 0.97 | 1.00 | 0.96 | 0.99 |
| Terrestrial (T) |  | P (T) = 0.02 | P (T) = 0.02 | 0.00 | 0.03 | 0.01 | 0.04 |
| **Habitat c** | 646 |  | | | | | |
| Marine (M) |  | P (M) = 0.68 | P (M) = 0.68 | 0.62 | 0.72 | 0.62 | 0.73 |
| Freshwater (F) |  | P (F) = 0.14 | P (F) = 0.13 | 0.11 | 0.17 | 0.11 | 0.17 |
| Terrestrial (T) |  | P (T) = 0.18 | P (T) = 0.19 | 0.17 | 0.21 | 0.17 | 0.21 |
| **Sporulation** | 937 |  | | | | | |
| Spore-forming (S) |  | P (S) = 0.92 | P (S) = 0.98 | 0.60 | 1.00 | 0.47 | 1.00 |
| Non-spore forming (N) |  | P (N) = 0.08 | P (N) = 0.02 | 0.00 | 0.38 | 0.00 | 0.49 |
| **Nutritional mode: Electron donor** | 434 |  | | | | | |
| Lithotrophic (L) |  | P (L) = 0.99 | P (L) = 1.00 | 0.95 | 1.00 | 0.89 | 1.00 |
| Organotrophic (O) |  | P (O) = 0.01 | P (O) = 0.00 | 0.00 | 0.05 | 0.00 | 0.11 |
| **Nutritional mode: Energy source** | 492 |  | | | | | |
| Chemotrophic (C) |  | P (C) = 1.00 | P (C) = 1.00 | 1.00 | 1.00 | 1.00 | 1.00 |
| Phototrophic (P) |  | P (P) = 0.00 | P (P) = 0.00 | 0.00 | 0.00 | 0.00 | 0.00 |
| **Oxygen requirement** | 1233 |  | | | | | |
| Aerobic (A) |  | P (A) = 0.00 | P (A) = 0.00 | 0.00 | 0.00 | 0.00 | 0.00 |
| Anaerobic (N) |  | P (N) = 1.00 | P (N) = 1.00 | 1.00 | 1.00 | 1.00 | 1.00 |
| **Enzymes: catalase** | 565 |  | | | | | |
| Present (P) |  | P (P) = 0.92 | P (P) = 0.92 | 0.90 | 0.94 | 0.90 | 0.94 |
| Absent (A) |  | P (A) = 0.08 | P (A) = 0.08 | 0.06 | 0.10 | 0.06 | 0.10 |
| **Enzymes: oxidase** | 398 |  | | | | | |
| Present (P) |  | P (P) = 0.73 | P (P) = 0.73 | 0.73 | 0.74 | 0.72 | 0.74 |
| Absent (A) |  | P (A) = 0.27 | P (A) = 0.27 | 0.26 | 0.27 | 0.26 | 0.28 |
| **Temperature (optimum)** | 888 |  | | | | | |
| Psychrotolerant (P) |  | P (P) = 0.00 | P (P) = 0.00 | 0.00 | 0.01 | 0.00 | 0.01 |
| Mesophilic (M) |  | P (M) = 0.00 | P (M) = 0.00 | 0.00 | 0.00 | 0.00 | 0.00 |
| Thermophilic (T) |  | P (T) = 0.13 | P (T) = 0.10 | 0.02 | 0.28 | 0.04 | 0.33 |
| Hyperthermophilic (H) |  | P (H) = 0.87 | P (H) = 0.90 | 0.71 | 0.98 | 0.66 | 0.96 |
| **NaCl (optimum)** | 191 |  | | | | | |
| Halotolerant (T) |  | P (T) = 0.76 | P (T) = 0.80 | 0.03 | 1.00 | 0.00 | 0.98 |
| Halophile (P) |  | P (P) = 0.05 | P (P) = 0.02 | 0.00 | 0.18 | 0.00 | 0.26 |
| Extreme halophile (E) |  | P (E) = 0.03 | P (E) = 0.02 | 0.00 | 0.08 | 0.00 | 0.16 |
| Non-halophile (N) |  | P (N) = 0.16 | P (N) = 0.11 | 0.00 | 0.36 | 0.01 | 0.97 |
| **pH (optimum)** | 481 |  | | | | | |
| Neutrophile (N) |  | P (N) = 0.68 | P (N) = 0.99 | 0.03 | 1.00 | 0.02 | 1.00 |
| Acidophile (A) |  | P (A) = 0.01 | P (A) = 0.00 | 0.00 | 0.04 | 0.00 | 0.05 |
| Hyperacidophile (H) |  | P (H) = 0.00 | P (H) = 0.00 | 0.00 | 0.01 | 0.00 | 0.01 |
| Alkaliphile (K) |  | P (K) = 0.31 | P (K) = 0.00 | 0.00 | 0.96 | 0.00 | 0.97 |
| **Character** | **Number of species** | **LACA** | | | | | |
|  |  | **Mean PP** | **Median PP** | **Lower HPD** | **Upper HPD** | **Lower 95% PI** | **Upper 95% PI** |
| **Shape** | 1372 |  | | | | | |
| Ovoid (O) |  | P (O) = 0.49 | P (O) = 0.54 | 0.00 | 0.82 | 0.00 | 0.85 |
| Rod (R) |  | P (R) = 0.00 | P (R) = 0.00 | 0.00 | 0.01 | 0.00 | 0.02 |
| Coccoid (C) |  | P (C) = 0.51 | P (C) = 0.46 | 0.18 | 1.00 | 0.15 | 1.00 |
| **Pleomorphism** | 1385 |  | | | | | |
| Monomorphic (M) |  | P (M) = 0.63 | P (M) = 0.62 | 0.53 | 0.74 | 0.53 | 0.74 |
| Non-monomorphic (N) |  | P (N) = 0.37 | P (N) = 0.38 | 0.26 | 0.47 | 0.26 | 0.47 |
| **Motility** | 1251 |  | | | | | |
| Motile (M) |  | P (M) = 0.90 | P (M) = 0.89 | 0.85 | 1.00 | 0.84 | 1.00 |
| Non-motile (N) |  | P (N) = 0.10 | P (N) = 0.11 | 0.00 | 0.15 | 0.00 | 0.16 |
| **Cell wall** | 1331 |  | | | | | |
| Present (P) |  | P (P) = 1.00 | P (P) = 1.00 | 1.00 | 1.00 | 1.00 | 1.00 |
| Absent (A) |  | P (A) = 0.00 | P (A) = 0.00 | 0.00 | 0.00 | 0.00 | 0.00 |
| **Cell plan** | 1263 |  | | | | | |
| Monoderm (M) |  | P (M) = 1.00 | P (M) = 1.00 | 1.00 | 1.00 | 1.00 | 1.00 |
| Diderm (D) |  | P (D) = 0.00 | P (D) = 0.00 | 0.00 | 0.00 | 0.00 | 0.00 |
| **Cell aggregation** | 762 |  | | | | | |
| Aggregating (A) |  | P (A) = 0.51 | P (A) = 0.51 | 0.31 | 0.69 | 0.29 | 0.69 |
| Non-aggregating (N) |  | P (N) = 0.49 | P (N) = 0.49 | 0.29 | 0.69 | 0.31 | 0.71 |
| **Habitat a** | 1339 |  | | | | | |
| Free-living (F) |  | P (F) = 1.00 | P (F) = 1.00 | 1.00 | 1.00 | 1.00 | 1.00 |
| Non-free living (N) |  | P (N) = 0.00 | P (N) = 0.00 | 0.00 | 0.00 | 0.00 | 0.00 |
| **Habitat b** | 654 |  | | | | | |
| Aquatic (A) |  | P (A) = 0.99 | P (A) = 0.99 | 0.98 | 1.00 | 0.98 | 1.00 |
| Terrestrial (T) |  | P (T) = 0.01 | P (T) = 0.01 | 0.00 | 0.02 | 0.00 | 0.02 |
| **Habitat c** | 646 |  | | | | | |
| Marine (M) |  | P (M) = 0.17 | P (M) = 0.17 | 0.15 | 0.20 | 0.15 | 0.20 |
| Freshwater (F) |  | P (F) = 0.66 | P (F) = 0.66 | 0.58 | 0.72 | 0.58 | 0.72 |
| Terrestrial (T) |  | P (T) = 0.17 | P (T) = 0.17 | 0.13 | 0.22 | 0.13 | 0.22 |
| **Sporulation** | 937 |  | | | | | |
| Spore-forming (S) |  | P (S) = 0.01 | P (S) = 0.01 | 0.00 | 0.02 | 0.00 | 0.02 |
| Non-spore forming (N) |  | P (N) = 0.99 | P (N) = 0.99 | 0.98 | 1.00 | 0.98 | 1.00 |
| **Nutritional mode: Electron donor** | 434 |  | | | | | |
| Lithotrophic (L) |  | P (L) = 1.00 | P (L) = 1.00 | 1.00 | 1.00 | 1.00 | 1.00 |
| Organotrophic (O) |  | P (O) = 0.00 | P (O) = 0.00 | 0.00 | 0.00 | 0.00 | 0.00 |
| **Nutritional mode: Energy source** | 492 |  | | | | | |
| Chemotrophic (C) |  | P (C) = 1.00 | P (C) = 1.00 | 1.00 | 1.00 | 1.00 | 1.00 |
| Phototrophic (P) |  | P (P) = 0.00 | P (P) = 0.00 | 0.00 | 0.00 | 0.00 | 0.00 |
| **Oxygen requirement** | 1233 |  | | | | | |
| Aerobic (A) |  | P (A) = 0.00 | P (A) = 0.00 | 0.00 | 0.00 | 0.00 | 0.00 |
| Anaerobic (N) |  | P (N) = 1.00 | P (N) = 1.00 | 1.00 | 1.00 | 1.00 | 1.00 |
| **Enzymes: catalase** | 565 |  | | | | | |
| Present (P) |  | P (P) = 1.00 | P (P) = 1.00 | 1.00 | 1.00 | 1.00 | 1.00 |
| Absent (A) |  | P (A) = 0.00 | P (A) = 0.00 | 0.00 | 0.00 | 0.00 | 0.00 |
| **Enzymes: oxidase** | 398 |  | | | | | |
| Present (P) |  | P (P) = 1.00 | P (P) = 1.00 | 1.00 | 1.00 | 1.00 | 1.00 |
| Absent (A) |  | P (A) = 0.00 | P (A) = 0.00 | 0.00 | 0.00 | 0.00 | 0.00 |
| **Temperature (optimum)** | 888 |  | | | | | |
| Psychrotolerant (P) |  | P (P) = 0.00 | P (P) = 0.00 | 0.00 | 0.00 | 0.00 | 0.04 |
| Mesophilic (M) |  | P (M) = 0.00 | P (M) = 0.00 | 0.00 | 0.00 | 0.00 | 0.00 |
| Thermophilic (T) |  | P (T) = 0.00 | P (T) = 0.00 | 0.00 | 0.00 | 0.00 | 0.00 |
| Hyperthermophilic (H) |  | P (H) = 1.00 | P (H) = 1.00 | 1.00 | 1.00 | 0.96 | 1.00 |
| **NaCl (optimum)** | 191 |  | | | | | |
| Halotolerant (T) |  | P (T) = 0.73 | P (T) = 0.78 | 0.00 | 0.98 | 0.00 | 1.00 |
| Halophile (P) |  | P (P) = 0.04 | P (P) = 0.01 | 0.00 | 0.13 | 0.00 | 0.15 |
| Extreme halophile (E) |  | P (E) = 0.15 | P (E) = 0.08 | 0.00 | 1.00 | 0.00 | 1.00 |
| Non-halophile (N) |  | P (N) = 0.08 | P (N) = 0.06 | 0.00 | 0.22 | 0.00 | 0.49 |
| **pH (optimum)** | 481 |  | | | | | |
| Neutrophile (N) |  | P (N) = 0.24 | P (N) = 0.17 | 0.01 | 0.76 | 0.07 | 0.92 |
| Acidophile (A) |  | P (A) = 0.69 | P (A) = 0.74 | 0.06 | 0.87 | 0.06 | 0.87 |
| Hyperacidophile (H) |  | P (H) = 0.03 | P (H) = 0.00 | 0.00 | 0.16 | 0.00 | 0.20 |
| Alkaliphile (K) |  | P (K) = 0.05 | P (K) = 0.04 | 0.00 | 0.11 | 0.00 | 0.18 |

**Table S3. Bayesian estimates of continuous traits for LUCA, LBCA and LACA from Segata et al’s tree.**

Ancestral state estimates for cell and genome sizes, optimal temperature, NaCl and pH from the best of two competing models - the random walk and directional models (See Methods). Phylogenetic signal (λ) is estimated for each trait. Abbreviations: LUCA – last universal common ancestor; LBCA – last bacterial common ancestor; LACA – last archaeal common ancestor. HPD - Highest Posterior Density; 95% PI - 95% Probability Interval.

|  | **Number of species** | **λ** | **λ**  **HPD**  **range** | **λ**  **95% PI range** | **LUCA PP** | **LUCA 95% PI range** | **LBCA**  **PP** | **LBCA**  **95% PI range** | **LACA**  **PP** | **LACA 95% PI range** |
| --- | --- | --- | --- | --- | --- | --- | --- | --- | --- | --- |
| **Cell size (µm)** | ƒ | | | | | | | | | |
| **Average width** | 841 | 1.00 | 0.99-1.00 | 0.99-1.00 | 0.55 | 0.44-0.68 | 0.55 | 0.44-0.68 | 0.54 | 0.41-0.70 |
| **Average length** | 818 | 0.99 | 0.99 | 0.99 | 2.18 | 1.59-2.98 | 2.19 | 1.65-2.91 | 2.45 | 1.65-3.65 |
| **Genome** |  | | | | | | | | | |
| **Average genome size (Mb)** | 1488 | 1.00 | 1.00 | 1.00 | 2.65 | 2.28-3.08 | 2.86 | 2.43-3.36 | 2.33 | 1.99-2.72 |
| **Average gene number** | 1486 | 1.00 | 1.00 | 1.00 | 2598 | 2216-3045 | 2578 | 2224-2989 | 2448 | 2057-2913 |
| **Temperature (°C)** |  | | | | | | | | | |
| **Optimum lower** | 684 | 1.00 | 1.00 | 1.00 | 72.6 | 62.9-83.7 | 63.6 | 54.1-74.7 | 80.8 | 71.7-91 |
| **Optimum upper** | 659 | 1.00 | 1.00 | 1.00 | 72.8 | 64.3-82.3 | 66.1 | 58.2-75.1 | 79.9 | 71.2-89.6 |
| **pH** |  | | | | | | | | | |
| **Optimum lower** | 398 | 1.00 | 1.00 | 1.00 | 7.0 | 6.7-7.3 | 7.1 | 6.8-7.4 | 6.9 | 6.5-7.2 |
| **Optimum upper** | 391 | 0.99 | 0.99 | 0.99 | 7.0 | 6.8-7.3 | 7.1 | 6.9-7.4 | 6.9 | 6.6-7.2 |
| **NaCl (% w/v)** |  | | | | | | | | | |
| **Optimum lower** | 176 | 0.95 | 0.95 | 0.95 | 1.80 | 1.38-2.31 | 1.82 | 1.36-2.38 | 1.78 | 1.31-2.35 |
| **Optimum upper** | 174 | 0.98 | 0.98-0.99 | 0.98-0.99 | 2.26 | 1.63-3.06 | 2.29 | 1.57-3.20 | 2.22 | 1.53-3.09 |

**Table S4. Physicochemical parameters estimate from the reduced categorical dataset from Chai et al’s tree.**

Posterior probabilities (PP) for optimum temperature pH and NaCl from Chai et al’s tree are shown for LUCA, LBCA and LACA in the form of P (X) = value (0.00-1.00). Abbreviations: LUCA - last universal common ancestor; LBCA -last bacterial common ancestor; LACA - last archaeal common ancestor; P (X) - PP for character state X; HPD - Highest Posterior Density; 95% PI - 95% Probability Interval.

| **Categorical character (Based on non-duplicated continuous data)** | **Number of species** | **LUCA** | | | | | |
| --- | --- | --- | --- | --- | --- | --- | --- |
|  |  | **Mean PP** | **Median PP** | **Lower HPD** | **Upper HPD** | **Lower 95% PI** | **Upper 95% PI** |
| **Temperature (optimum)** | 1533 |  | | | | | |
| Psychrotolerant (P) |  | P (P) = 0.00 | P (P) = 0.00 | 0.00 | 0.03 | 0.00 | 0.07 |
| Mesophilic (M) |  | P (M) = 0.00 | P (M) = 0.00 | 0.00 | 0.01 | 0.00 | 0.01 |
| Thermophilic (T) |  | P (T) = 0.14 | P (T) = 0.06 | 0.00 | 0.66 | 0.00 | 0.91 |
| Hyperthermophilic (H) |  | P (H) = 0.86 | P (H) = 0.93 | 0.31 | 1.00 | 0.06 | 0.99 |
| **NaCl (optimum)** | 491 |  | | | | | |
| Halotolerant (T) |  | P (T) = 0.68 | P (T) = 0.70 | 0.45 | 0.81 | 0.38 | 0.79 |
| Halophile (P) |  | P (P) = 0.10 | P (P) = 0.09 | 0.06 | 0.14 | 0.07 | 0.14 |
| Extreme halophile (E) |  | P (E) = 0.10 | P (E) = 0.09 | 0.05 | 0.21 | 0.06 | 0.23 |
| Non-halophile (N) |  | P (N) = 0.12 | P (N) = 0.11 | 0.08 | 0.25 | 0.08 | 0.28 |
| **pH (optimum)** | 929 |  | | | | | |
| Neutrophile (N) |  | P (N) = 0.55 | P (N) = 0.58 | 0.25 | 0.77 | 0.19 | 0.74 |
| Acidophile (A) |  | P (A) = 0.29 | P (A) = 0.26 | 0.00 | 0.57 | 0.05 | 0.72 |
| Hyperacidophile (H) |  | P (H) = 0.02 | P (H) = 0.00 | 0.00 | 0.09 | 0.00 | 0.17 |
| Alkaliphile (K) |  | P (K) = 0.14 | P (K) = 0.14 | 0.05 | 0.27 | 0.03 | 0.26 |
| **Categorical character (Based on non-duplicated continuous data)** | **Number of species** | **LBCA** | | | | | |
|  |  | **Mean PP** | **Median PP** | **Lower HPD** | **Upper HPD** | **Lower 95% PI** | **Upper 95% PI** |
| **Temperature (optimum)** | 1533 |  | | | | | |
| Psychrotolerant (P) |  | P (P) = 0.07 | P (P) = 0.02 | 0.00 | 0.28 | 0.00 | 0.33 |
| Mesophilic (M) |  | P (M) = 0.12 | P (M) = 0.09 | 0.00 | 0.33 | 0.01 | 0.39 |
| Thermophilic (T) |  | P (T) = 0.71 | P (T) = 0.78 | 0.26 | 0.91 | 0.16 | 0.89 |
| Hyperthermophilic (H) |  | P (H) = 0.10 | P (H) = 0.04 | 0.00 | 0.49 | 0.00 | 0.73 |
| **NaCl (optimum)** | 491 |  | | | | | |
| Halotolerant (T) |  | P (T) = 0.66 | P (T) = 0.68 | 0.45 | 0.83 | 0.38 | 0.80 |
| Halophile (P) |  | P (P) = 0.09 | P (P) = 0.08 | 0.04 | 0.14 | 0.05 | 0.15 |
| Extreme halophile (E) |  | P (E) = 0.11 | P (E) = 0.10 | 0.05 | 0.20 | 0.06 | 0.22 |
| Non_halophile (N) |  | P (N) = 0.14 | P (N) = 0.14 | 0.09 | 0.27 | 0.10 | 0.29 |
| **pH (optimum)** | 929 |  | | | | | |
| Neutrophile (N) |  | P (N) = 0.52 | P (N) = 0.52 | 0.42 | 0.65 | 0.42 | 0.65 |
| Acidophile (A) |  | P (A) = 0.03 | P (A) = 0.02 | 0.00 | 0.06 | 0.01 | 0.13 |
| Hyperacidophile (H) |  | P (H) = 0.01 | P (H) = 0.00 | 0.00 | 0.03 | 0.00 | 0.04 |
| Alkaliphile (K) |  | P (K) = 0.44 | P (K) = 0.44 | 0.30 | 0.57 | 0.28 | 0.55 |
| **Categorical character (Based on non-duplicated continuous data)** | **Number of species** | **LACA** | | | | | |
|  |  | **Mean PP** | **Median PP** | **Lower HPD** | **Upper HPD** | **Lower 95% PI** | **Upper 95% PI** |
| **Temperature (optimum)** | 1533 |  | | | | | |
| Psychrotolerant (P) |  | P (P) = 0.00 | P (P) = 0.00 | 0.00 | 0.02 | 0.00 | 0.04 |
| Mesophilic (M) |  | P (M) = 0.00 | P (M) = 0.00 | 0.00 | 0.00 | 0.00 | 0.00 |
| Thermophilic (T) |  | P (T) = 0.04 | P (T) = 0.02 | 0.00 | 0.10 | 0.01 | 0.40 |
| Hyperthermophilic (H) |  | P (H) = 0.96 | P (H) = 0.98 | 0.88 | 1.00 | 0.59 | 0.99 |
| **NaCl (optimum)** | 491 |  | | | | | |
| Halotolerant (T) |  | P (T) = 0.52 | P (T) = 0.54 | 0.36 | 0.57 | 0.32 | 0.57 |
| Halophile (P) |  | P (P) = 0.15 | P (P) = 0.15 | 0.15 | 0.18 | 0.14 | 0.18 |
| Extreme halophile (E) |  | P (E) = 0.15 | P (E) = 0.14 | 0.11 | 0.24 | 0.12 | 0.26 |
| Non-halophile (N) |  | P (N) = 0.18 | P (N) = 0.17 | 0.16 | 0.26 | 0.16 | 0.28 |
| **pH (optimum)** | 929 |  | | | | | |
| Neutrophile (N) |  | P (N) = 0.31 | P (N) = 0.31 | 0.07 | 0.50 | 0.10 | 0.55 |
| Acidophile (A) |  | P (A) = 0.60 | P (A) = 0.61 | 0.40 | 0.88 | 0.24 | 0.84 |
| Hyperacidophile (H) |  | P (H) = 0.03 | P (H) = 0.00 | 0.00 | 0.19 | 0.00 | 0.27 |
| Alkaliphile (K) |  | P (K) = 0.06 | P (K) = 0.05 | 0.01 | 0.12 | 0.02 | 0.14 |

**Table S5. Physicochemical parameters estimate from the reduced categorical dataset from Segata et al’s tree.**

Posterior probabilities (PP) for optimum temperature pH and NaCl from Segata et al’s tree are shown for LUCA, LBCA and LACA in the form of P (X) = value (0.00-1.00). Abbreviations: LUCA - last universal common ancestor; LBCA -last bacterial common ancestor; LACA - last archaeal common ancestor; P (X) - PP for character state X; HPD - Highest Posterior Density; 95% PI - 95% Probability Interval.

| **Character (Based on non-duplicated continuous data)** | **Number of species** | **LUCA** | | | | | |
| --- | --- | --- | --- | --- | --- | --- | --- |
|  |  | **Mean PP** | **Median PP** | **Lower HPD** | **Upper HPD** | **Lower 95% PI** | **Upper 95% PI** |
| **Temperature (optimum)** | 714 |  | | | | | |
| Psychrotolerant (P) |  | P (P) = 0.00 | P (P) = 0.00 | 0.00 | 0.03 | 0.00 | 0.07 |
| Mesophilic (M) |  | P (M) = 0.00 | P (M) = 0.00 | 0.00 | 0.00 | 0.00 | 0.00 |
| Thermophilic (T) |  | P (T) = 0.00 | P (T) = 0.00 | 0.00 | 0.00 | 0.00 | 0.01 |
| Hyperthermophilic (H) |  | P (H) = 1.00 | P (H) = 1.00 | 0.99 | 1.00 | 0.99 | 1.00 |
| **NaCl (optimum)** | 181 |  | | | | | |
| Halotolerant (T) |  | P (T) = 0.77 | P (T) = 0.90 | 0.00 | 0.99 | 0.00 | 0.99 |
| Halophile (P) |  | P (P) = 0.03 | P (P) = 0.02 | 0.00 | 0.08 | 0.00 | 0.10 |
| Extreme halophile (E) |  | P (E) = 0.10 | P (E) = 0.03 | 0.00 | 1.00 | 0.00 | 1.00 |
| Non_halophile (N) |  | P (N) = 0.10 | P (N) = 0.03 | 0.00 | 0.74 | 0.00 | 0.94 |
| **pH (optimum)** | 403 |  | | | | | |
| Neutrophile (N) |  | P (N) = 0.54 | P (N) = 0.61 | 0.19 | 1.00 | 0.14 | 0.98 |
| Acidophile (A) |  | P (A) = 0.23 | P (A) = 0.23 | 0.00 | 0.39 | 0.00 | 0.42 |
| Hyperacidophile (H) |  | P (H) = 0.02 | P (H) = 0.00 | 0.00 | 0.09 | 0.00 | 0.16 |
| Alkaliphile (K) |  | P (K) = 0.21 | P (K) = 0.08 | 0.00 | 0.58 | 0.00 | 0.66 |
| **Character (Based on non-duplicated continuous data)** | **Number of species** | **LBCA** | | | | | |
|  |  | **Mean PP** | **Median PP** | **Lower HPD** | **Upper HPD** | **Lower 95% PI** | **Upper 95% PI** |
| **Temperature (optimum)** | 714 |  | | | | | |
| Psychrotolerant (P) |  | P (P) = 0.00 | P (P) = 0.00 | 0.00 | 0.00 | 0.00 | 0.01 |
| Mesophilic (M) |  | P (M) = 0.00 | P (M) = 0.00 | 0.00 | 0.00 | 0.00 | 0.00 |
| Thermophilic (T) |  | P (T) = 0.11 | P (T) = 0.10 | 0.02 | 0.24 | 0.03 | 0.28 |
| Hyperthermophilic (H) |  | P (H) = 0.89 | P (H) = 0.90 | 0.75 | 0.98 | 0.72 | 0.97 |
| **NaCl (optimum)** | 181 |  | | | | | |
| Halotolerant (T) |  | P (T) = 0.75 | P (T) = 0.81 | 0.01 | 0.99 | 0.00 | 0.98 |
| Halophile (P) |  | P (P) = 0.06 | P (P) = 0.06 | 0.00 | 0.23 | 0.00 | 0.29 |
| Extreme halophile (E) |  | P (E) = 0.03 | P (E) = 0.02 | 0.00 | 0.11 | 0.00 | 0.19 |
| Non-halophile (N) |  | P (N) = 0.16 | P (N) = 0.10 | 0.00 | 0.84 | 0.01 | 0.98 |
| **pH (optimum)** | 403 |  | | | | | |
| Neutrophile (N) |  | P (N) = 0.65 | P (N) = 0.95 | 0.07 | 1.00 | 0.40 | 1.00 |
| Acidophile (A) |  | P (A) = 0.02 | P (A) = 0.01 | 0.00 | 0.06 | 0.00 | 0.07 |
| Hyperacidophile (H) |  | P (H) = 0.00 | P (H) = 0.00 | 0.00 | 0.01 | 0.00 | 0.02 |
| Alkaliphile (K) |  | P (K) = 0.33 | P (K) = 0.02 | 0.00 | 0.92 | 0.00 | 0.95 |
| **Character (Based on non-duplicated continuous data)** | **Number of species** | **LACA** | | | | | |
|  |  | **Mean PP** | **Median PP** | **Lower HPD** | **Upper HPD** | **Lower 95% PI** | **Upper 95% PI** |
| **Temperature (optimum)** | 714 |  | | | | | |
| Psychrotolerant (P) |  | P (P) = 0.00 | P (P) = 0.00 | 0.00 | 0.00 | 0.00 | 0.00 |
| Mesophilic (M) |  | P (M) = 0.00 | P (M) = 0.00 | 0.00 | 0.00 | 0.00 | 0.00 |
| Thermophilic (T) |  | P (T) = 0.00 | P (T) = 0.00 | 0.00 | 0.00 | 0.00 | 0.00 |
| Hyperthermophilic (H) |  | P (H) = 1 | P (H) = 1 | 1.00 | 1.00 | 1.00 | 1.00 |
| **NaCl (optimum)** | 181 |  | | | | | |
| Halotolerant (T) |  | P (T) = 0.71 | P (T) = 0.79 | 0.00 | 0.99 | 0.00 | 0.99 |
| Halophile (P) |  | P (P) = 0.06 | P (P) = 0.01 | 0.00 | 0.16 | 0.00 | 0.18 |
| Extreme halophile (E) |  | P (E) = 0.15 | P (E) = 0.06 | 0.00 | 1.00 | 0.00 | 1.00 |
| Non-halophile (N) |  | P (N) = 0.08 | P (N) = 0.04 | 0.00 | 0.33 | 0.00 | 0.49 |
| **pH (optimum)** | 403 |  | | | | | |
| Neutrophile (N) |  | P (N) = 0.24 | P (N) = 0.19 | 0.02 | 0.82 | 0.06 | 0.92 |
| Acidophile (A) |  | P (A) = 0.67 | P (A) = 0.72 | 0.06 | 0.86 | 0.05 | 0.86 |
| Hyperacidophile (H) |  | P (H) = 0.03 | P (H) = 0.00 | 0.00 | 0.18 | 0.00 | 0.28 |
| Alkaliphile (K) |  | P (K) = 0.06 | P (K) = 0.05 | 0.00 | 0.18 | 0.00 | 0.29 |

**Table S6. Test for phylogenetic signal in the 18 phenotypic traits from both trees.** For each trait P (0), P (1), P (2) and P (3) represent the posterior probability for states 0, 1, 2 and 3 respectively (See Methods). **RND**: randomized data result (see Methods)

| **Traits from Segata’s tree** | **Number of States** | **Log likelihood (Lh)** | **P (0)** | **P (1)** | **P (2)** | **P (3)** |
| --- | --- | --- | --- | --- | --- | --- |
| Catalase | 2 | -151.566432 | 0.004425 | 0.995575 | NA | NA |
| Catalase RND | 2 | -358.318775 | 0.5 | 0.5 | NA | NA |
| Cell Aggregation | 2 | -212.867937 | 0.49452 | 0.50548 | NA | NA |
| Cell Aggregation RND | 2 | -225.011069 | 0.5 | 0.5 | NA | NA |
| Cell plan | 2 | -151.247436 | 0.999975 | 0.000025 | NA | NA |
| Cell plan RND | 2 | -853.001253 | 0.5 | 0.5 | NA | NA |
| Cell wall | 2 | -33.111123 | 0 | 1 | NA | NA |
| Cell wall RND | 2 | -165.002997 | 0.5 | 0.5 | NA | NA |
| Electron donor | 2 | -62.181108 | 1 | 0 | NA | NA |
| Electron donor RND | 2 | -129.80615 | 0.5 | 0.5 | NA | NA |
| Energy source | 2 | -38.294749 | 0 | 1 | NA | NA |
| Energy source RND | 2 | -80.250359 | 0.5 | 0.5 | NA | NA |
| Habitat a | 2 | -446.061496 | 0.002267 | 0.997733 | NA | NA |
| Habitat a RND | 2 | -823.652364 | 0.5 | 0.5 | NA | NA |
| Habitat b | 2 | -241.107119 | 0.936058 | 0.063942 | NA | NA |
| Habitat b RND | 2 | -317.362116 | 0.5 | 0.5 | NA | NA |
| Habitat c | 3 | -451.195209 | 0.452956 | 0.047519 | 0.499525 | NA |
| Habitat c RND | 3 | -565.620046 | 0.333333 | 0.333333 | 0.333333 | NA |
| Motility | 2 | -540.704001 | 0.004321 | 0.995679 | NA | NA |
| Motility RND | 2 | -866.130452 | 0.5 | 0.5 | NA | NA |
| NaCl | 4 | -147.656713 | 0.2108 | 0.0016 | 0.000033 | 0.787568 |
| NaCl RND | 4 | -189.272762 | 0.249993 | 0.249999 | 0.249981 | 0.250026 |
| Oxidase | 2 | -127.472032 | 0.092247 | 0.907753 | NA | NA |
| Oxidase RND | 2 | -272.586531 | 0.5 | 0.5 | NA | NA |
| Oxygen | 2 | -186.293445 | 0.996422 | 0.003578 | NA | NA |
| Oxygen RND | 2 | -625.117046 | 0.5 | 0.5 | NA | NA |
| pH | 4 | -333.056722 | 0.553116 | 0.270402 | 0 | 0.176482 |
| pH RND | 4 | -427.399762 | 0.25 | 0.25 | 0.25 | 0.25 |
| Pleomorphism | 2 | -523.597604 | 0.374752 | 0.625248 | NA | NA |
| Pleomorphism RND | 2 | -604.011942 | 0.5 | 0.5 | NA | NA |
| Shape | 3 | -399.226776 | 0.679857 | 0.285529 | 0.034614 | NA |
| Shape RND | 3 | -732.786761 | 0.333333 | 0.333333 | 0.333333 | NA |
| Sporulation | 2 | -150.985422 | 0.062159 | 0.937841 | NA | NA |
| Sporulation RND | 2 | -421.373437 | 0.5 | 0.5 | NA | NA |
| Temperature | 4 | -279.117382 | 0 | 0 | 0.006667 | 0.993332 |
| Temperature RND | 4 | -571.85098 | 0.25 | 0.25 | 0.25 | 0.25 |
| **Traits from Chai’s tree** | **Number of States** | **Log likelihood (Lh)** | **P (0)** | **P (1)** | **P (2)** | **P (3)** |
| Catalase | 2 | -347.862197 | 0.002633 | 0.997367 | NA | NA |
| Catalase RND | 2 | -810.69535 | 0.5 | 0.5 | NA | NA |
| Cell Aggregation | 2 | -409.613607 | 0.711732 | 0.288268 | NA | NA |
| Cell Aggregation RND | 2 | -444.061443 | 0.5 | 0.5 | NA | NA |
| Cell plan | 2 | -225.647757 | 0.999945 | 0.000055 | NA | NA |
| Cell plan RND | 2 | -1780.236585 | 0.5 | 0.5 | NA | NA |
| Cell wall | 2 | -42.379022 | 0.000662 | 0.999338 | NA | NA |
| Cell wall RND | 2 | -277.475041 | 0.5 | 0.5 | NA | NA |
| Electron donor | 2 | -99.361806 | 0.999992 | 0.000008 | NA | NA |
| Electron donor RND | 2 | -241.145059 | 0.5 | 0.5 | NA | NA |
| Energy source | 2 | -60.360516 | 0 | 1 | NA | NA |
| Energy source RND | 2 | -112.84899 | 0.5 | 0.5 | NA | NA |
| Habitat a | 2 | -1123.576693 | 0.007233 | 0.992767 | NA | NA |
| Habitat a RND | 2 | -1737.047742 | 0.5 | 0.5 | NA | NA |
| Habitat b | 2 | -569.212313 | 0.879962 | 0.120038 | NA | NA |
| Habitat b RND | 2 | -797.508744 | 0.5 | 0.5 | NA | NA |
| Habitat c | 3 | -995.827125 | 0.784805 | 0.048754 | 0.16644 | NA |
| Habitat c RND | 3 | -1335.956717 | 0.333333 | 0.333333 | 0.333333 | NA |
| Motility | 2 | -1119.923143 | 0.015955 | 0.984045 | NA | NA |
| Motility RND | 2 | -1807.14771 | 0.5 | 0.5 | NA | NA |
| NaCl | 4 | -430.453899 | 0.431519 | 0.024802 | 0.015686 | 0.527993 |
| NaCl RND | 4 | -588.919942 | 0.25 | 0.25 | 0.25 | 0.25 |
| Oxidase | 2 | -362.297605 | 0.033355 | 0.966645 | NA | NA |
| Oxidase RND | 2 | -692.793618 | 0.5 | 0.5 | NA | NA |
| Oxygen | 2 | -296.24967 | 0.981483 | 0.018517 | NA | NA |
| Oxygen RND | 2 | -1238.112531 | 0.5 | 0.5 | NA | NA |
| pH | 4 | -760.702113 | 0.463424 | 0.417619 | 0 | 0.118957 |
| pH RND | 4 | -937.777142 | 0.25 | 0.25 | 0.25 | 0.25 |
| Pleomorphism | 2 | -983.755507 | 0.625548 | 0.374452 | NA | NA |
| Pleomorphism RND | 2 | -1145.933157 | 0.5 | 0.5 | NA | NA |
| Shape | 3 | -682.392924 | 0.752887 | 0.022408 | 0.224705 | NA |
| Shape RND | 3 | -1349.395684 | 0.333333 | 0.333333 | 0.333333 | NA |
| Sporulation | 2 | -227.854428 | 0.993748 | 0.006252 | NA | NA |
| Sporulation RND | 2 | -936.660441 | 0.5 | 0.5 | NA | NA |
| Temperature | 4 | -539.138796 | 0.000099 | 0.000459 | 0.235656 | 0.763786 |

**Table S6.1. Character state coding used in Table S6.**

| **Trait** | **State 0** | **State 1** | **State 2** | **State 3** |
| --- | --- | --- | --- | --- |
| Catalase | Absent | Present | NA | NA |
| Cell Aggregation | Non-aggregating | Aggregating | NA | NA |
| Cell plan | Monoderm | Diderm | NA | NA |
| Cell wall | Absent | Present | NA | NA |
| Electron donor | Lithotrophic | Organotrophic | NA | NA |
| Energy source | Phototroph | Chemotroph | NA | NA |
| Habitat a | Non-free living | Free living | NA | NA |
| Habitat b | Aquatic | Terrestrial | NA | NA |
| Habitat c | Marine | Terrestrial | Fresh-water | NA |
| Motility | Non-motile | Motile | NA | NA |
| Nacl | Non-halophile | Halophile | Extreme-halophile | Halotolerant |
| Oxidase | Absent | Present | NA | NA |
| Oxygen | Anaerobic | Aerobic | NA | NA |
| pH | Neutrophile | Acidophile | Hyperacidophile | Alkaliphile |
| Pleomorphism | Monomorphic | Non-monomorphic | NA | NA |
| Shape | Ovoid | Rod | Coccoid | NA |
| Sporulation | Non-spore forming | Spore forming | NA | NA |
| Temperature | Psychrotolerant | Mesophilic | Thermophilic | Hyperthermophilic |

**Table S7. Numbers of species scored for each trait.**

|  | **Number of species scored** | |
| --- | --- | --- |
| **Character trait** | Large tree^1^ | Small tree^8^ |
| Shape | 2812 | 1372 |
| Pleomorphism | 2843 | 1385 |
| Motility | 2586 | 1251 |
| Cell wall | 2761 | 1331 |
| Cell plan | 2614 | 1263 |
| Cell aggregation | 1363 | 762 |
| Habitat a | 2752 | 1339 |
| Habitat b | 1495 | 654 |
| Habitat c | 1486 | 646 |
| Sporulation | 1943 | 937 |
| Nutritional mode: Electron donor | 932 | 434 |
| Nutritional mode: Energy source | 1036 | 492 |
| Oxygen requirement | 2532 | 1233 |
| Enzymes: catalase | 1424 | 565 |
| Enzymes: oxidase | 1036 | 398 |
| Temperature (optimum) | 1838 | 888 |
| NaCl (optimum) | 524 | 191 |
| pH (optimum) | 1053 | 481 |
| Cell width | 1705 | 841 |
| Cell length | 1656 | 818 |
| Genome size | 2978 | 1488 |
| Gene number | 2965 | 1486 |
| Optimum lower temperature | 1448 | 684 |
| Optimum upper temperature | 1432 | 659 |
| Optimum lower pH | 896 | 398 |
| Optimum upper pH | 879 | 391 |
| Optimum lower NaCl | 470 | 176 |
| Optimum upper NaCl | 479 | 174 |

**Bacterial and Archaeal Phenotypic Database (BAPdb)**

**(separate file)**

doi: https://doi.org/10.6084/m9.figshare.12987509.v1

**References**.

1.	Chai, J., Kora, G., Ahn, T. H., Hyatt, D. & Pan, C. Functional phylogenomics analysis of bacteria and archaea using consistent genome annotation with UniFam. *BMC Evol. Biol.* **14**, 1–13 (2014).

2.	Kuever, J., Rainey, F. A. & Widdel, F. Bergey’s Manual of Systematic Bacteriology Volume 1: The Archaea and the Deeply Branching and Phototrophic Bacteria. in *Bergey’s Manual of Systematic Bacteriology* (2005). doi:10.1007/978-0-387-21609-6

3.	Brenner, D. J., Krieg, N. R. & Staley, J. T. *Bergey’s Manual of Systematic Bacteriology - Vol 2: The Proteobacteria. Part C The Alpha-, Beta-, Delta-, and Epsilonproteobacteria*. *Springer-Verlag New York Inc.* (2005). doi:10.1007/0-387-29298-5

4.	Vos, P. *et al.* *Bergey’s Manual of Systematic Bacteriology - Vol 3: The Firmicutes*. *Springer-Verlag New York Inc.* (2009). doi:10.1007/b92997

5.	Krieg, N. R. *et al.* *Bergey’s Manual of Systematic Bacteriology Volume Four*. *Bergey’s Manual of Systematic Bacteriology, Volume 4* (2010). doi:10.1007/978-0-387-68572-4

6.	Goodfellow, M. *et al.* Bergey’s Manual of Systematic Bacteriology - Second Edition, Volume 5. in *Bergey’s Manual* (2012). doi:10.1007/978-0-387-68233-4

7.	Parte, A. C. LPSN - List of prokaryotic names with standing in nomenclature (Bacterio.net), 20 years on. *International Journal of Systematic and Evolutionary Microbiology* (2018). doi:10.1099/ijsem.0.002786

8.	Segata, N., Börnigen, D., Morgan, X. C. & Huttenhower, C. PhyloPhlAn is a new method for improved phylogenetic and taxonomic placement of microbes. *Nat. Commun.* **4**, 2304 (2013).

9.	Monciardini, P., Cavaletti, L., Schumann, P., Rohde, M. & Donadio, S. Conexibacter woesei gen. nov., sp. nov., a novel representative of a deep evolutionary line of descent within the class Actinobacteria. *Int. J. Syst. Evol. Microbiol.* **53**, 569–576 (2003).
